# The role of evolving niche choice in herbivore adaptation to host plants

**DOI:** 10.1101/2024.05.02.592156

**Authors:** Peter Nabutanyi, Alitha Edison, Peter Czuppon, Shuqing Xu, Meike J. Wittmann

**Author notes:** **Corresponding author:** Peter Nabutanyi, Department of Theoretical Biology, Faculty of Biology, Bielefeld University, Universitätsstraße 25, 33615 Bielefeld, Germany. **Authors’ e-mail addresses:**.

## Abstract

Individuals living in heterogeneous environments often choose microenvironments that provide benefits to their fitness. Theory predicts that such “niche choice” can promote rapid adaptation to novel environments and help maintain genetic diversity. An open question of large applied importance is how niche choice and niche choice evolution affect the evolution of insecticide resistance in phytophagous insects. We, therefore, developed an individual-based model based on phytophagous insects to examine the evolution of insecticide resistance and host plant choice via oviposition preferences. To find biologically realistic parameter ranges, we performed an empirical literature survey on insecticide resistance in major agricultural pests and also conducted a density-dependent survival experiment using potato beetles. We find that, in comparison to a scenario where individuals randomly oviposit eggs on toxic or non-toxic plants, the evolution of niche choice generally leads to slower evolution of resistance and facilitates the coexistence of different phenotypes. Our simulations also reveal that recombination rate and dominance effects can influence the evolution of both niche choice and resistance. Thus, this study provides new insights into the effects of niche choice on resistance evolution and highlights the need for more studies on the genetic basis of resistance and choice.

## 1 Introduction

Individuals of many animal species living in heterogeneous environments use various cues to choose habitats to feed, live and reproduce. This process is termed habitat choice or niche choice (Edelaar et al., 2008; Akcali & Porter, 2017; Edelaar et al., 2017; Camacho & Hendry, 2020; Mortier & Bonte, 2020; Trappes et al., 2022). The choice of habitat can substantially impact the survival and reproductive success of an individual, thereby influencing ecological and evolutionary processes (Morris, 2003; Berner & Thibert-Plante, 2015). For example, many phytophagous insects are specialized and adapted to a few host plant species. Even within plant species, food quality and quantity vary among individuals, such that the choice of suitable individual host plants for feeding and oviposition is critical for the insects’ fitness. Studies have shown that host plant choice can facilitate adaptation to the host plant (Levins & MacArthur, 1969; Jaenike, 1978; Thompson, 1988). In addition, such adaptive dispersal can also facilitate the maintenance of genetic diversity in a heterogeneous environment (Edelaar et al., 2008; Ravigné et al., 2009; Bolnick & Otto, 2013; Czuppon et al., 2021). Although niche choice is generally perceived to increase individual fitness (Trappes et al., 2022), research also shows that many animals make habitat choices that do not always match their phenotype (Cotter & Edwards, 2006; Camacho et al., 2015, 2020). A meta-analysis showed that among insect herbivores, the correlations between preference and fitness are highly variable and can even be negative (Mayhew, 1997; Gripenberg et al., 2010). Thus, it remains open how host plant choices in insect herbivores influence individual fitness and trait evolution.

Host plant adaptation and specialization in phytophagous insects can be very rapid, especially in the face of agricultural practices. A classic example is the rapid evolution of insecticide resistance in agricultural pests. Because insecticide applications in agricultural fields create heterogeneous environments with varying concentrations and combinations of toxins spread over space and time (Pose-Juan et al., 2015; Kosubová et al., 2020; Liu et al., 2023), insect pests can choose non-toxic hosts and/or adapt to toxins. Such host preference by insects can influence the evolution of dispersal and specialization (Ravigné et al., 2024). In these heterogeneous environments and given the short generation time of many insect pests, niche choice as a trait can evolve over time (Jaenike & Holt, 1991). The evolution of niche choice can be facilitated by a number of factors, such as differences in selection pressures between habitats and local dispersal (Edelaar et al., 2008, 2017; Camacho & Hendry, 2020). Models show that the joint evolution of niche choice and adaptation to the environmental conditions can alter evolutionary outcomes under certain conditions (Ravigné et al., 2009; Berner & Thibert-Plante, 2015; Kisdi et al., 2020). The evolution of niche choice can also facilitate diversification and local adaptation to a new habitat. For example, generalists can be favoured under fixed niche choice, but specialists are favoured when both niche choice and local adaptation evolve (Ravigné et al., 2009).

While studies have shown that agricultural pests can show different behavioural responses to insecticides or toxins (Zhao et al., 2016; Luong et al., 2016; Nansen et al., 2016; Edison et al., 2024), we still know little about the role of host choice on the evolution of insecticide resistance in agricultural pests. In particular, it is unclear whether niche choice and the evolution of niche choice will speed up or slow down the evolution of insecticide resistance. Previous theoretical studies have demonstrated that evolutionary outcomes depend on the nature of fitness functions, habitat choice mechanisms, and parameter ranges (Ravigné et al., 2009; Berner & Thibert-Plante, 2015). Thus, there is a need to find out where most pest insect systems fall and bridge the gap between theoretical models and empirical studies, for example, by parameterizing models based on empirical data. In addition, previous studies have focused on the role of ecological parameters, but it remains unclear how genetic factors such as recombination rates between loci and allele dominance coefficients, i.e., genetic architecture, affect the joint evolution of niche choice and host adaptation. Furthermore, most previous studies on the evolution of habitat choice have focused on matching habitat choice (Edelaar et al., 2008; Ravigné et al., 2009; Camacho et al., 2015; Berner & Thibert-Plante, 2015; Akcali & Porter, 2017; Edelaar et al., 2017; Mortier & Bonte, 2020; Camacho & Hendry, 2020; Ravigné et al., 2024). However, oviposition behaviour in some phytophagous insects has been linked to direct genetic habitat choice where individual choices are determined by genetic alleles and not influenced by fitness expectations (matching habitat choice) or learning (induced habitat choice), i.e., individual performance and habitat choices are independent (Jaenike, 1987; Mortier & Bonte, 2020; Fanara et al., 2023; Álvarez-Ocaña et al., 2023). So far, there is still little research on the evolution of choice behaviour based on direct genetic habitat choice (Berner & Thibert-Plante, 2015), in particular for its effect on resistance evolution in insects.

To address these challenges, we developed an eco-evolutionary individual-based model to investigate how niche choice evolves and affects the evolutionary dynamics of resistance. In our model, we consider a population of insect herbivores living in a heterogeneous habitat with toxic and non-toxic host plants where individual behaviour in choosing their hosts to oviposit affects individual fitness and resistance evolution at the population level. We focus on the so-called direct genetic habitat choice. We simulated populations where choice and resistance traits are determined by linked or unlinked loci evolving over time and compared them to populations with fixed host choice. To place our model into a biologically realistic parameter space and explore the impact of host plant choice on the evolution of agricultural pests, we selected ten herbivorous agricultural pests and carried out a literature survey to estimate the distributions of key model parameters. In addition, we carried out an experiment with one of the pest species, the Colorado potato beetle, to estimate the local density-dependent survival that we use in our model simulation. Together, we use our model to study the role of host choice evolution on resistance evolution in agricultural pests.

## 2 Methods

### Model overview

We model a diploid and, for simplicity, hermaphroditic population. The population lives in a habitat consisting of toxic and non-toxic microenvironments (Figure 1). We consider two diallelic loci that are either unlinked or linked with a given per-generation recombination probability between them. One locus, the resistance locus, codes for the resistance trait that might also confer costs or benefits on either offspring viability, fecundity, or both. The other locus, the choice locus, codes for the choice trait that determines the probability with which an individual chooses a non-toxic microenvironment. Thus, the two traits jointly determine individual fitness. Our model captures the role of key ecological and genetic parameters and processes, such as local density regulation, host mating preference, resistance costs and benefits on fitness components and micro-habitat differences in selection pressure.

**Figure 1:**
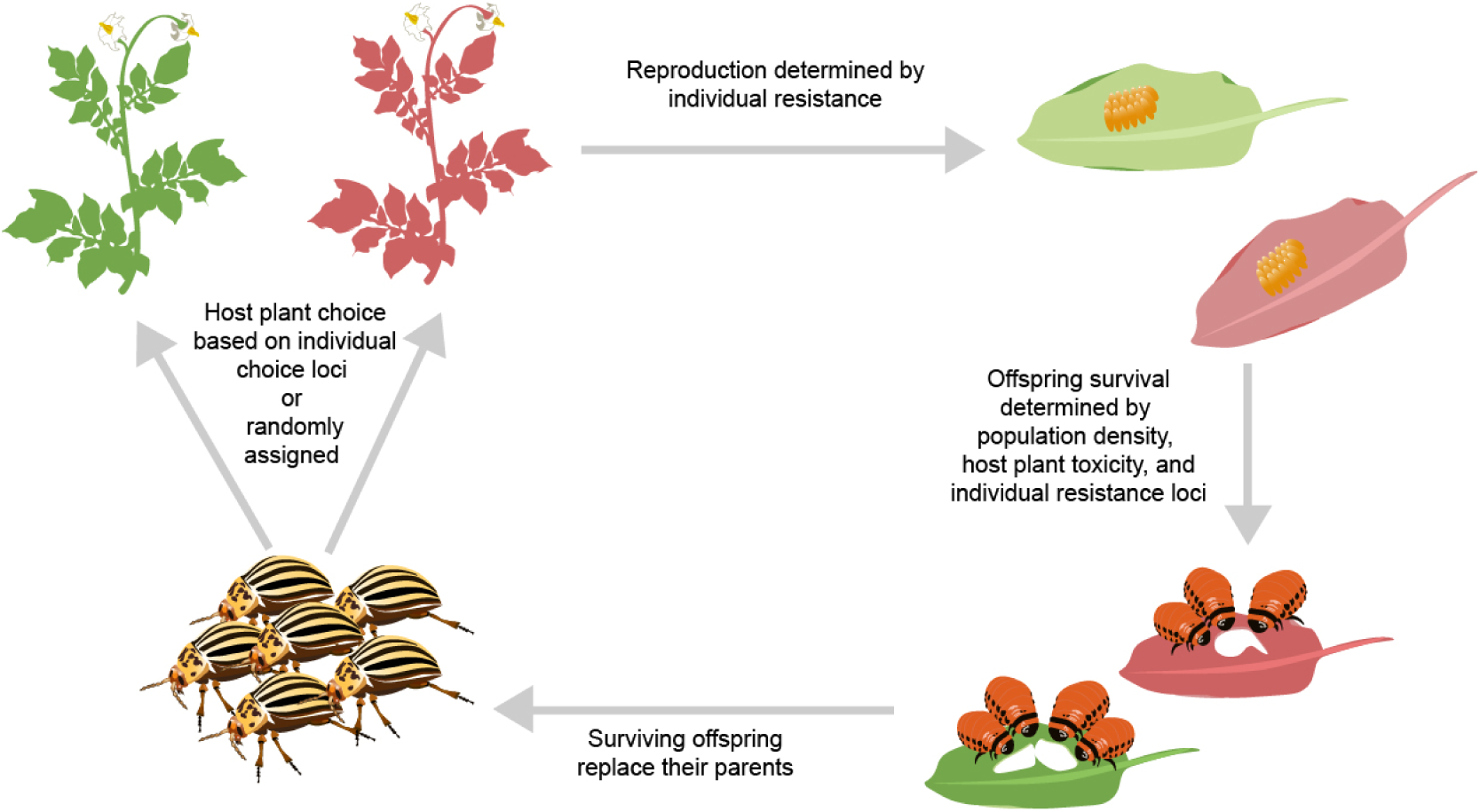
Flow chart showing the major steps of the model. We assume that the red plant is more toxic than the green one. Adult individuals either choose their habitats based on their genetically determined niche choice traits (evolving choice scenario) or choose randomly based on a fixed probability irrespective of their genetically determined choice traits (fixed choice scenario). At the reproduction stage, each individual then determines whether to mate with an individual from the same host plant or the other, depending on the level of host mating preference. We assume that each mating results in one offspring, and the mating decisions are made separately and independently for each offspring. The total number of offspring per individual is drawn from a Poisson distribution with a mean parameter influenced by individual resistance. We apply a global density regulation by randomly selecting a maximum of *K* offspring whenever the total number of offspring is above *K*. The offspring stays at the host plant of the focal parent (i.e., the “mother’s” host plant). Each offspring’s viability depends on the host plant’s toxicity, offspring resistance, and local larval density on the host plant. All surviving offspring are then pooled to replace the parent generation.

### Individual-based model

#### Composition of the initial population

The habitat consists of two equally frequent micro-environments, i.e., two types of host plant. We assume host plant 1 to be non-toxic, with toxicity *τ*_1_ = 0, and plant 2 to be toxic, 0 *< τ*_2_ *≤* 1. We start the population with *N*_0_ herbivore individuals. The allelic state for each allele copy in every individual is independently drawn from a Bernoulli distribution with the desired initial frequency of resistance alleles *p_r,_*_0_ and choice alleles *p_c,_*_0_.

#### Life cycle

In every generation, *t*, the current population size *N_t_*is determined. If the population is extinct, *N_t_* = 0, the simulation stops. Otherwise, each individual independently chooses one of the host plants. The non-toxic host plant is chosen with probability *z_c_*, where *z_c_*is a host plant choice trait that is either fixed for all individuals (*z_c,fixed_*, under fixed random choice) or genetically determined by a choice locus (under evolving niche choice). For genetically determined choice, we assume that the choice trait is determined by 1 locus with two alleles: 0 (preference for the toxic plant) and 1 (preference for the non-toxic plant). Homozygotes of type 11 always go to the non-toxic plant (*z_c_* = 1), and 00 homozygotes always go to the toxic plant (*z_c_* = 0). Heterozygotes denoted by 10 or 01 go to the non-toxic plant with probability *z_c_* = *h_c_*, where *h_c_* is the dominance coefficient of allele 1 at the choice locus.

Once all individuals have chosen their host plant, we determine the number of individuals on the two host plants, *N_t,_*_1_ and *N_t,_*_2_. Now, for each individual, we draw the number of offspring from a Poisson distribution with mean

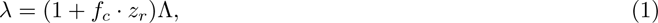

where *z_r_* is the individual’s resistance trait computed using analogous assumptions as in the choice trait. Based on our literature survey, the resistance trait is determined by a single locus (details in Subsection “Quantitative literature survey and model parameterization” below). We assume the locus has two alleles: 1 for the resistance allele and 0 for the non-resistance allele, such that 11 homozygotes are fully resistant, *z_r_* = 1, 00 homozygotes are fully susceptible, *z_r_* = 0, and heterozygotes have intermediate resistance *z_r_* = *h_r_*, where *h_r_* is the dominance coefficient of the resistance allele. Λ is the fecundity of non-resistant individuals (*z_r_*= 0), and *f_c_ ≥ −*1 is the fecundity cost or benefit associated with resistance, such that, positive and negative values imply resistance benefit and cost to fecundity, respectively.

To determine the offspring’s genotype, we assume for simplicity that mating is promiscuous such that, as an approximation, every offspring produced by an individual is independently assigned a second parent. For each offspring, the focal parent decides, using a host mating preference parameter *−*1 *≤ M_a_ ≤* 1, which host plant to choose the mating partner from. For *M_a_ >* 0, the mating partner is picked from the same host plant of the focal parent with probability *M_a_* and randomly from all host plants with probability 1 *− M_a_*. Consequently, for a focal parent at host plant *i*, the overall probability that a mate comes from the same host plant is 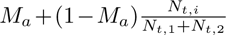. This tendency for a focal individual to choose a mate from the same host plant is referred to as assortative mating. If *M_a_* = 0, the mating partner is randomly chosen from the host plants. However, if *M_a_ <* 0, then *|M_a_|* represents the probability of picking a mate from a different host plant *j*, i.e., disassortative mating. The total probability that a mate comes from a different host plant is thus 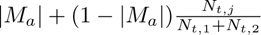.

Each offspring receives one allele at each locus from each of its two parents. The alleles are chosen randomly and independently under free recombination (recombination rate *R* = 0.5). Otherwise, they are partially or completely linked (0 *≤ R <* 0.5). For this, every individual has two chromosomes, and with probability 1 *− R*, resistance and choice alleles on the same chromosome are inherited together. The offspring stays on the host plant of the focal parent. We assume that no new mutations happen at the time scale of our simulations.

If the total number of offspring exceeds the global or environmental carrying capacity, *K*, the offspring population is randomly down-sampled to *K*. This step, therefore, implements a global density regulation. Furthermore, offspring survival depends on the plant’s toxicity, the resistance trait, and how crowded the plant is (i.e., local density regulation). That is, for an offspring on host plant *i* with resistance trait *z_r_*, the probability of surviving to adulthood is

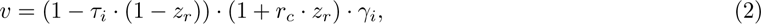

where the first factor means that the toxicity of the host plant is reduced in proportion to the resistance trait. The second factor gives the effect of resistance on offspring viability *r_c_*, such that positive and negative values imply resistance benefit and cost to viability, respectively. Negative values of this factor were set to 0. The last factor is the local density-dependent survival probability at host plant *i*, estimated from an experiment using potato beetles (as detailed in Subsection “Estimating local density regulation” below), and is given by

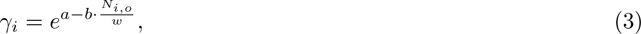

where *a* = 0.01148851 and *b* = 0.03404447 are the statistically estimated parameters, *N_i,o_* is the number of offspring produced at host plant *i*, and *w* is a scaling parameter. Equation (3) was restricted to be at most 1 by setting it to 1 whenever *γ_i_ >* 1.

Finally, the parent generation is removed, and all the surviving offspring form the new population form the next generation. The model is run for *t_max_* = 200 generations. We track and report the population sizes and frequencies of the resistance and choice alleles or traits after the offspring survival stage.

### Quantitative literature survey and model parameterization

From the Arthropod Pesticide Resistance Database (Mota-Sanchez & Wise, 2023), we made a list of the top 10 agricultural pests with the most cases of pesticide resistance and ranked them according to the number of compounds they were reported to be resistant against (Table S1.1). This top 10 list contains four moth species (the diamondback moth *Plutella xylostella*, the cotton bollworm *Helicoverpa armigera*, the fall armyworm *Spodoptera frugiperda*, and the beet armyworm *Spodoptera exigua*), two mite species (the two-spotted spider mite *Tetranychus urticae* and the European red mite *Panonychus ulmi*), two aphids (the green peach aphid *Myzus persicae* and the cotton aphid *Aphis gossypii*), as well as the sweet potato whitefly (*Bemisia tabaci*) and the Colorado potato beetle (*Leptinotarsa decemlineata*). We then conducted a literature review on the Web of Science using topic- and species-specific search strings and performed a quantitative survey to estimate the range and the distribution of the key model parameters (Table 1). We screened 802 articles and extracted 1552 values from 229 articles (details are given in Subsection S1). From the review, we obtained distributions of host plant toxicity, costs of resistance on viability and fecundity, the mean number of offspring, level of host mating preference, and frequency of the resistance alleles (Subsection S1). The ranges of parameters from the literature were adjusted to align with our model interpretation where necessary. For example, from the literature survey, both plants had some level of toxicity ranging between 0 and 1, but the distribution was converted such that relative toxicity was used where one plant was totally non-toxic while the other plant was toxic (details in Subsection S1.3). For parameters with inadequate information from the review, we chose values from a reasonable parameter range (Table 1).

**Table 1:**
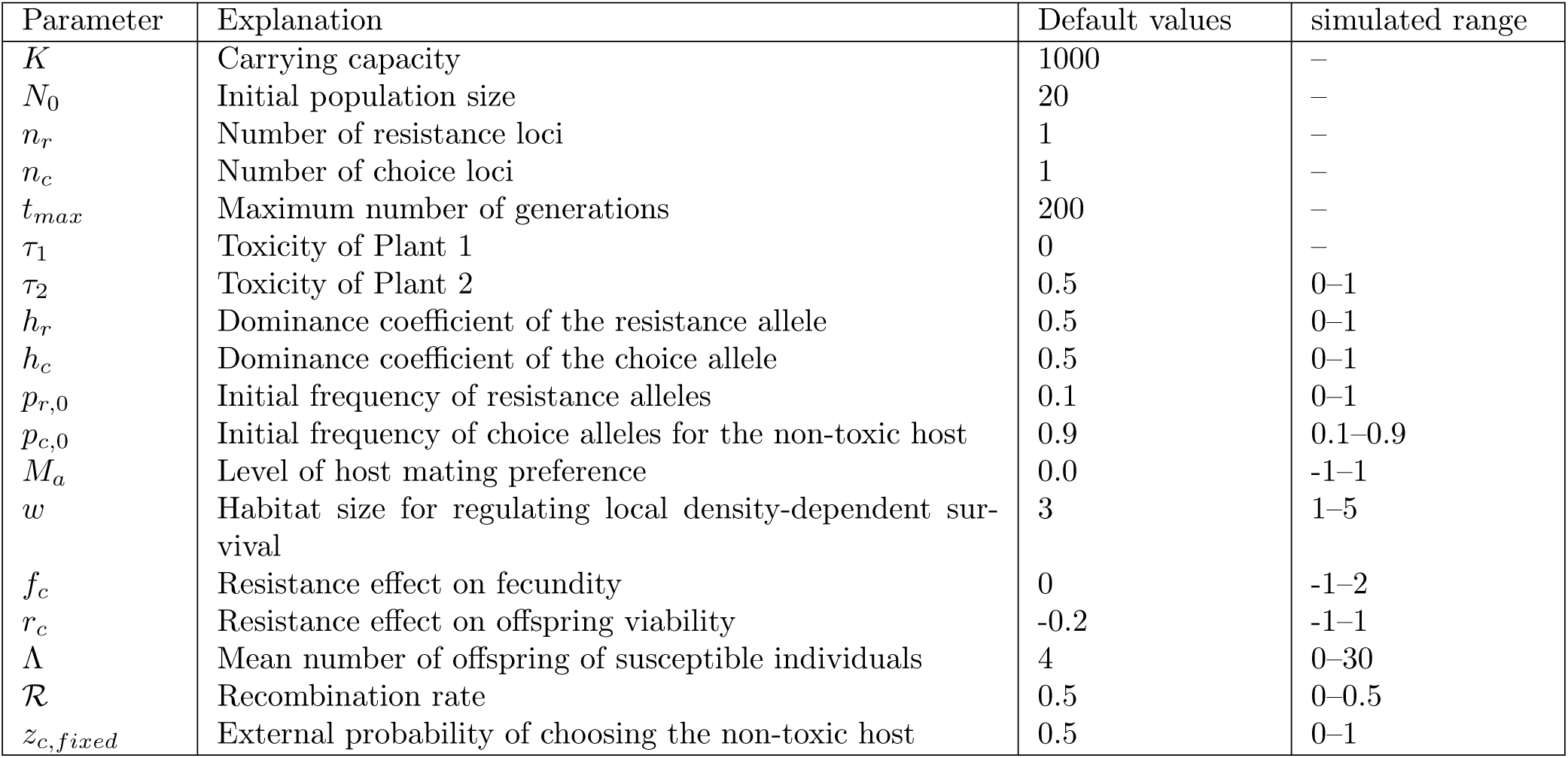
Explanation of all parameters used in the model. Unless specified otherwise, in all results plotted, the parameters are at their default values given in the third column. The default values are close to the means of the empirical parameter distributions from the literature survey.

| Parameter | Explanation | Default values | simulated range |
| --- | --- | --- | --- |
| $K$ | Carrying capacity | 1000 | – |
| $N_0$ | Initial population size | 20 | – |
| $n_r$ | Number of resistance loci | 1 | – |
| $n_c$ | Number of choice loci | 1 | – |
| $t_{max}$ | Maximum number of generations | 200 | – |
| $\tau_1$ | Toxicity of Plant 1 | 0 | – |
| $\tau_2$ | Toxicity of Plant 2 | 0.5 | 0–1 |
| $h_r$ | Dominance coefficient of the resistance allele | 0.5 | 0–1 |
| $h_c$ | Dominance coefficient of the choice allele | 0.5 | 0–1 |
| $p_{r,0}$ | Initial frequency of resistance alleles | 0.1 | 0–1 |
| $p_{c,0}$ | Initial frequency of choice alleles for the non-toxic host | 0.9 | 0.1–0.9 |
| $M_a$ | Level of host mating preference | 0.0 | –1–1 |
| $w$ | Habitat size for regulating local density-dependent survival | 3 | 1–5 |
| $f_c$ | Resistance effect on fecundity | 0 | –1–2 |
| $r_c$ | Resistance effect on offspring viability | –0.2 | –1–1 |
| $\Lambda$ | Mean number of offspring of susceptible individuals | 4 | 0–30 |
| $\mathcal{R}$ | Recombination rate | 0.5 | 0–0.5 |
| $z_{c,fixed}$ | External probability of choosing the non-toxic host | 0.5 | 0–1 |

### Estimating local density regulation

Our literature survey did not provide enough information on density regulation. Since this is a key parameter for the evolution of niche choice and local adaptation (Ravigné et al., 2004, 2009), we empirically estimated density-dependent survival probability. To obtain a realistic density-dependent fitness response as the starting point, we carried out an experiment using one of our 10 insect pest species from the survey, the Colorado potato beetles, *Leptinotarsa decemlineata*. We set up cages with different initial population sizes, i.e., numbers of first instar larvae, and quantified the survival until the pupation stage. Each cage was supplied with the same fixed amount of non-toxic potato plants as food resources. A detailed description is in Section S2. The observed relationship between initial beetle density and fitness (per-capita survival) was estimated by a linear model after log transformation. From the experiment, we obtain the density-dependent survival probability at the host plant *i* as given in Equation (3). To make our model applicable to different habitat sizes and population densities, we introduced a scaling parameter, *w ≥* 0, referred to as habitat size. The experimental data gives *w* = 1, which is the strongest density regulation we consider in our simulations. The higher the value of *w*, the weaker the local density regulation, i.e., higher offspring survival. The parameter, *w*, can also be interpreted as the number of plants available, where higher values imply more plants available. Therefore, Equation (3) generally implements local density regulation by indirectly setting local carrying capacities. As density regulation may take different forms in other phytophagous insects, our density regulation function and parameter distributions act as best guesses given the currently available empirical studies.

### Simulations and analysis

We simulate populations either under a fixed host plant choice, where a host plant has an externally fixed probability of being chosen by an individual irrespective of individual choice trait (random choice for probability of 0.5) or under the evolving host plant choice, where the probability of choosing a host plant depends on the choice trait of the individual genetically determined by the choice locus. The model is implemented in R (R Core Team, 2023).

To understand the role of model parameters and the evolution of host plant choice on the evolution of resistance, we examine the speed of trait evolution, trait means, and trait variances within populations by varying a given parameter while fixing the rest of the parameters to their default values (Table 1). The default values represent relatively large habitats, *w* = 3, that maintain large populations. For comparison, results under small habitats, *w* = 1, that maintain small populations are provided in the supplement (Subsection S4). Furthermore, the default settings are codominance of choice and resistance alleles, low initial frequency of resistance alleles, high frequency of choice alleles (for the non-toxic host), random mating, free recombination, no resistance effects on fecundity, and substantial resistance cost on viability. In addition, we analyse the distribution of the traits in the final generations for simulations where the parameters are randomly picked from the empirical distributions. Here, we draw a total of 500 parameter sets from the empirical parameter distributions or a uniform distribution for the case of parameters without sufficient information from the literature survey (Figure 2, Table 1). We do the sampling both separately for different species and jointly for artificially created species where the species distributions are combined.

**Figure 2:**
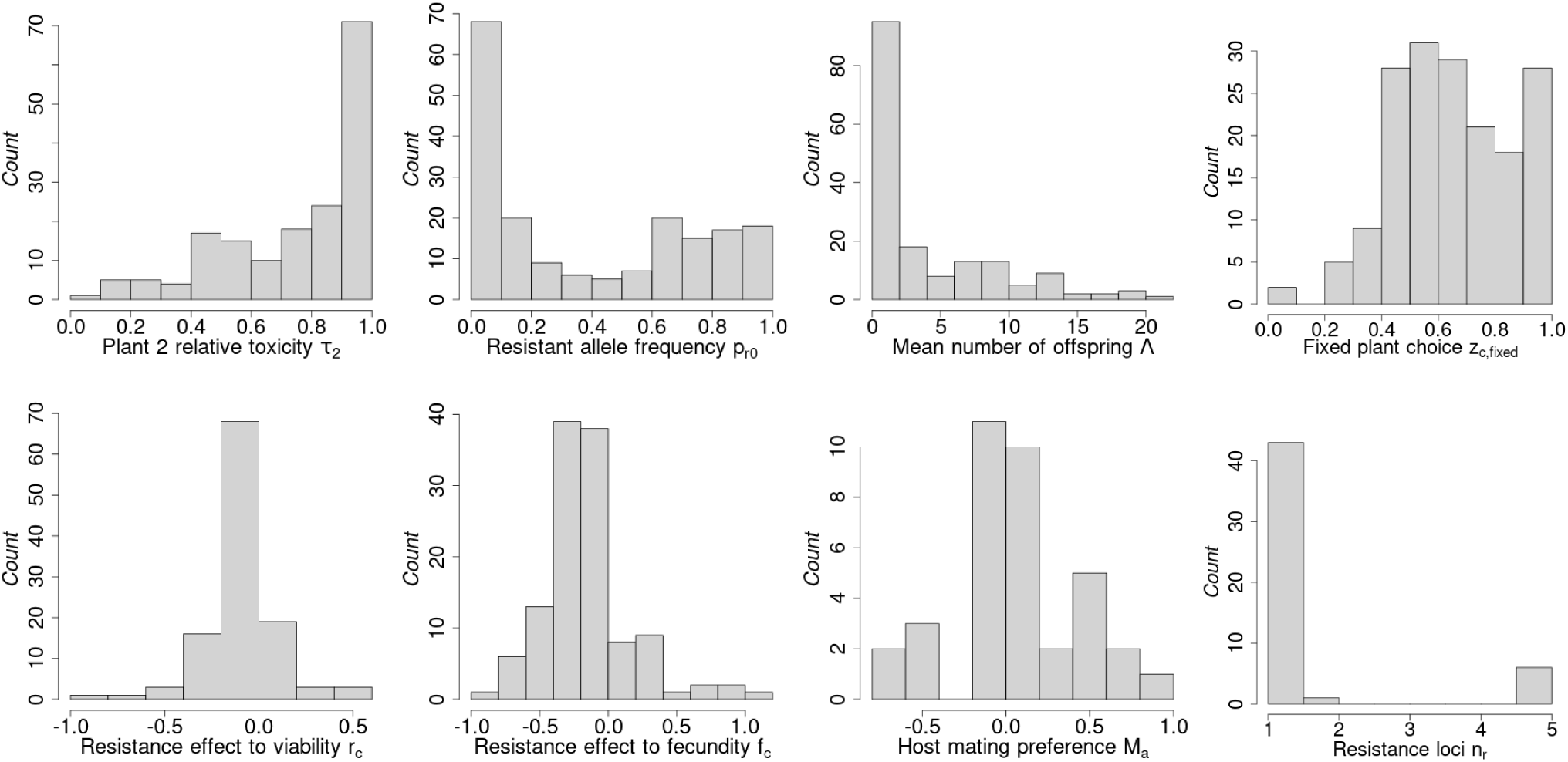
Distributions of parameters estimated from the literature survey across all 10 species. Toxicity at plant 2 assumes that toxicity at plant 1 is 0 (i.e., we show relative toxicity). The fixed plant choice gives the probability of choosing the toxic host (under fixed random choice behaviour). For host mating preference, the magnitude of values indicates the tendency to choose a mate from the same host plant (assortative mating - positive values), a different host plant (disassortative mating - negative values), or randomly among the available host plants (random mating - 0). Figure S1.1 shows the distributions at the species level. The details on how the distributions were obtained are in Subsection S1.

## 3 Results

### Empirical distributions from the literature survey

We first highlight the key results from the parameter distributions obtained from our literature survey (Figure 2). Most of the parameter distributions were highly skewed. The habitat toxicity is very high, with a median of 0.87 across all the data points included in the literature survey. Alleles associated with insecticide resistance were rare in populations, with median resistance allele frequencies of 0.22. The mean number of offspring produced per female varies between species, with the highest numbers reported for some of the lepidopteran species. The distribution of the plant choice shows that individuals were biased towards the non-toxic habitat for oviposition. For both survival and fecundity, we find more fitness costs than benefits associated with insecticide resistance. Data on mating preferences were limited to three lepidopteran species, namely, *Plutella xylostella*, *Helicoverpa armigera*, and *Spodoptera frugiperda* (Figure S1.1). Mating is in most cases close to random, but both assortative and disassortative mating do occur in these species. In the text, assortative mating refers to positive values of host mating preference, while disassortative mating refers to negative values of host mating preference. Most of the studies showed that only one locus was associated with insecticide resistance, which is the value we chose for our simulations. Our survey hardly obtained genetic information regarding species’ habitat choices. Therefore, we randomly chose the parameters from a feasible range for our simulations (Table 1).

### Population dynamics and evolution over time

In the presentation of the remaining results, we systematically compare scenarios where niche choice genetically evolves (“evolving choice behaviour”) to scenarios where niche choice is fixed and random, where individuals randomly move to each plant with equal probability (“fixed random choice behaviour”).

Our simulated herbivore populations appear to reach equilibrium population sizes by the end of 200 generations (Figure 3). In contrast to fixed random choice, where the population size at the two host plants was the same due to a fixed equal probability of choosing either plant, under evolving host plant choice toxic host plants had fewer individuals than non-toxic plants. The final population sizes increased with habitat size under both evolving and fixed host plant choice scenarios (compare upper to lower row in Figure 3, also Figure S3.1). However, strong local density regulation resulted in population sizes far below *K* = 1000, i.e., the set global carrying capacity.

**Figure 3:**
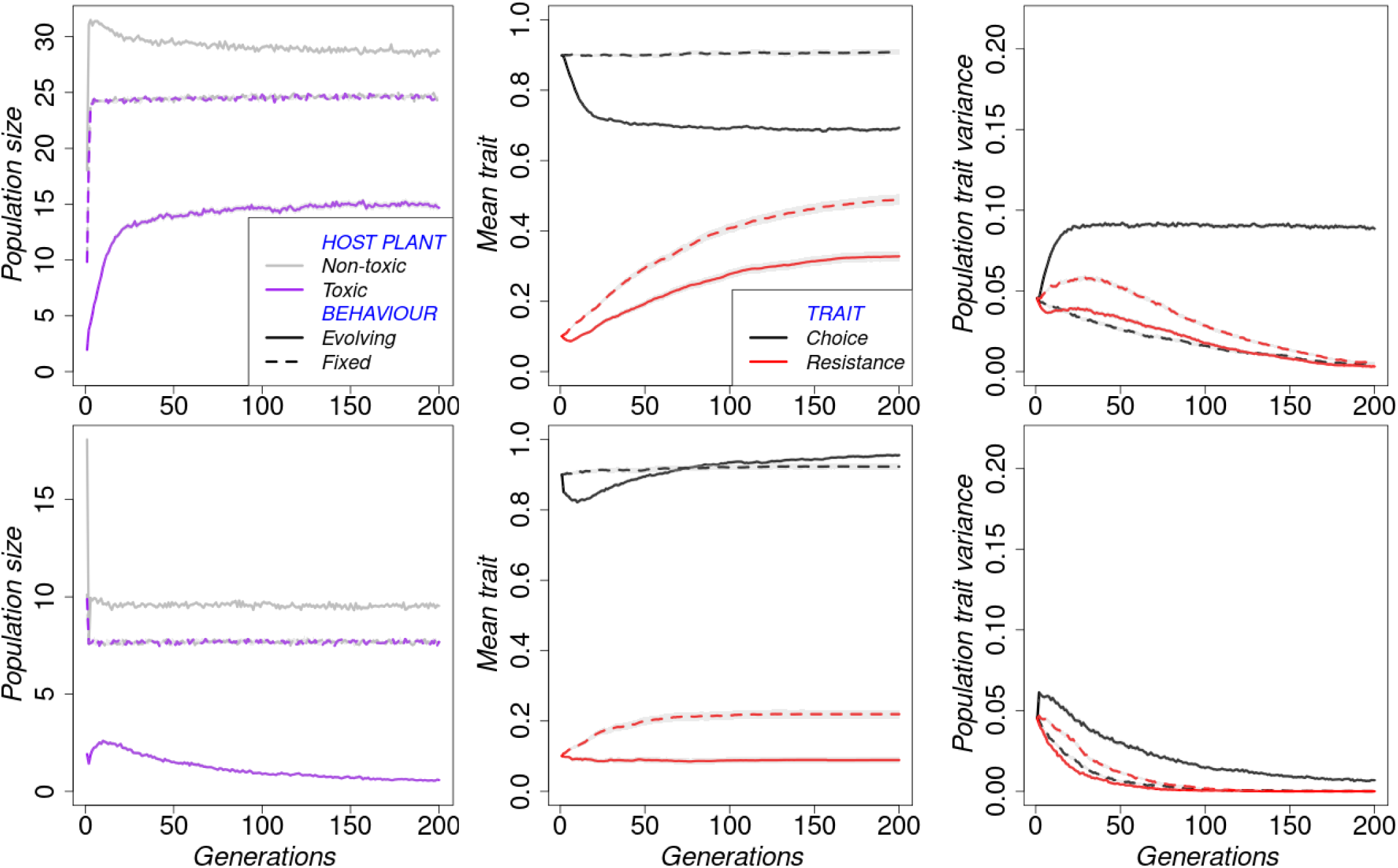
Trajectories of population sizes at the host plants, the population mean trait values and trait variances averaged across 1000 replicates. Notice the overlap of population sizes at the toxic and non-toxic plants under random choice behaviour. Upper row: large habitats *w* = 3; Lower row: small habitats *w* = 1. The grey regions represent the standard error (although too small to be seen in most plots).

In both behavioural scenarios, populations in large habitats became more resistant over time on average, but this increase was faster and led to a higher population mean resistance when individuals randomly chose their habitats than when they chose a habitat based on their genetically determined choice trait (Figure 3 middle column). In small habitats, resistance slightly increased under random choice of host plant but remained constant under evolving host plant choice. The average variance in resistance among individuals in a population decreased towards zero over the course of the simulation under both choice behaviours, indicating that in most replicate simulations, either the resistant or the sensitive allele went to fixation (Figure 3 right column).

On the other hand, the mean choice trait initially decreased under evolving host plant choice but remained constant under fixed random choice. As expected, the choice trait has no effect in the fixed random choice scenario, and therefore, the choice trait evolves neutrally. In addition, differences in habitat size led to qualitative changes in the dynamics of the choice trait (Figures 3). In large habitats, the choice trait rapidly settled to around 0.7, which roughly reflects the proportion of individuals on the non-toxic host plant. In small habitats, the initial rapid decrease was followed by a gradual increase to a constant value higher than the initial choice trait. The variance in choice trait among individuals was higher when niche choice was allowed to evolve than when individual choice of habitat was fixed and random. Populations in large habitats also maintained more variable phenotypes than populations in small habitats.

### Effects of model parameters on trait means and trait variances

Figure 4 shows final trait values simulated under the two choice behaviours. Given the default initial allele frequencies in Table 1, the initial mean resistance trait was 0.1, while the initial mean choice trait was 0.9 (except for the first two panels where the respective allele frequencies were varied). Final trait values reported in various figures that deviate from these initial traits indicate an evolved trait. For example, a lower choice trait value compared to the initial value indicates an increased preference for the toxic host plant, while a higher resistance trait implies an increased tolerance to toxins.

**Figure 4:**
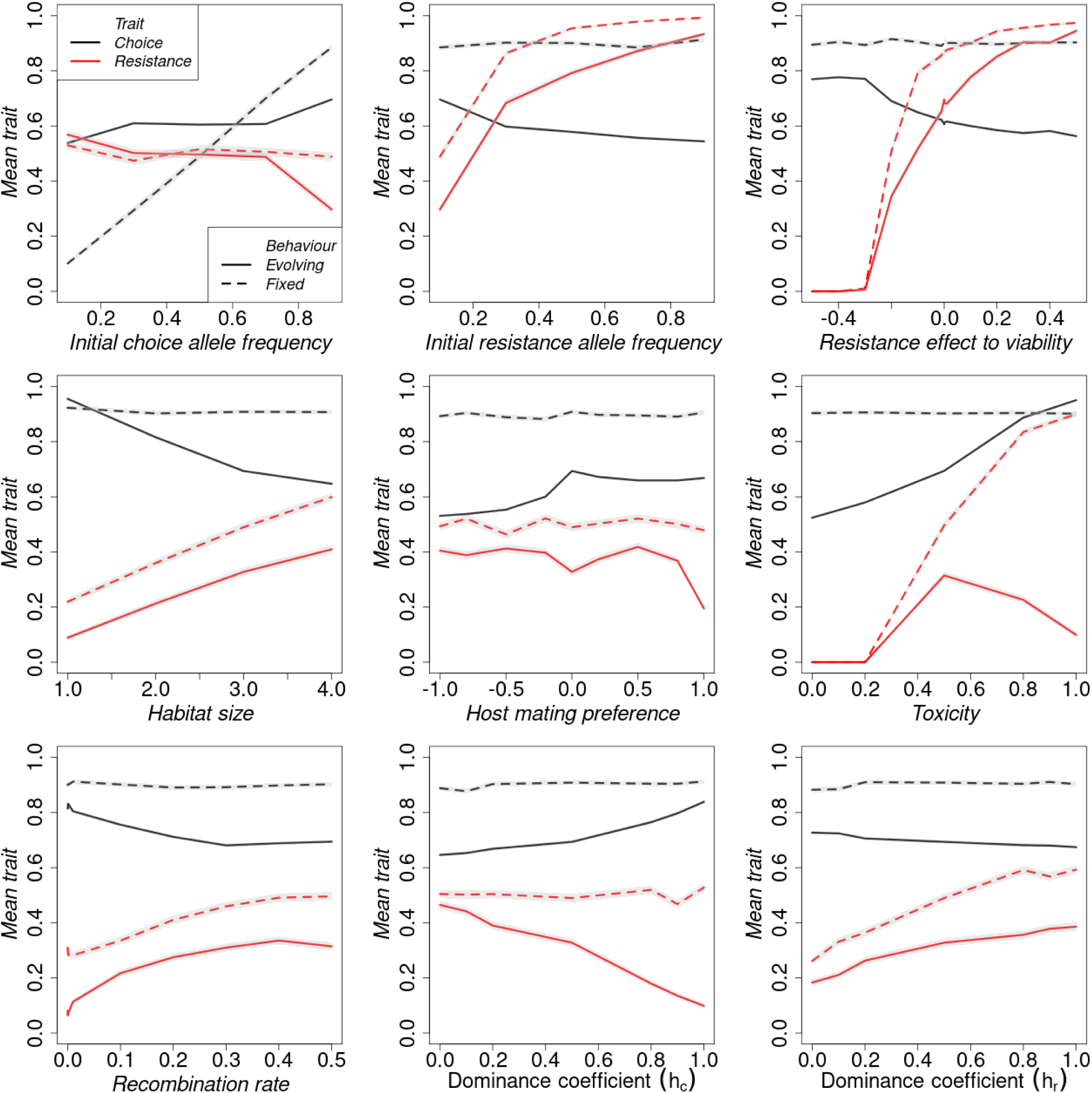
Final trait means: Effect of model parameters on the average resistance trait (red lines) and choice trait (black lines) at the end of the simulation under fixed random choice (dashed lines) and evolving host plant choice (continuous lines). The grey area shows the standard error around the mean trait across 1000 replicates.

As expected, the mean choice trait remains constant and is determined by the initial choice allele frequency under fixed random choice. However, under evolving plant choice, the niche choice trait evolved under a broad range of parameter values (Figures 4 and S4.2). While the final mean choice trait was generally above 0.5, and thus more individuals still chose the non-toxic plant than under fixed random choice, the final mean choice trait was lower than the initial mean choice trait for most of the parameter combinations (black solid lines below black dashed lines). Thus, at the end of the simulation more individuals chose the toxic habitat than at the beginning. There were also a few exceptions, however: individuals evolved larger avoidance of the toxic habitats (black solid lines above black dashed lines) when the initial choice allele frequency was low, when habitats were very small, or when the toxicity of the toxic plant was extremely high. In small habitats, the direction of choice evolution was not influenced by the initial choice allele frequency but by the initial frequency of the resistance allele, the resistance effect on fitness components, and host mating preference (Figures S3.3 and S4.2).

Consistent with the time series results, across broad ranges of the various model parameters, the evolution of host plant choice led to lower mean resistance at the end of the simulation than under fixed random choice, except when the initial choice allele frequency is low (Figures 4, S4.2, and S5.2). Furthermore, the speed of evolution of resistance is lower when niche choice evolves than when niche choice is fixed, except when the initial choice allele frequency for the non-toxic plant is low (Figures S3.2, S4.1). In small habitats, many model parameters had a weaker influence on the trait evolution (Subsection S4). When the initial frequency of the choice allele was low, the rate of resistance trait evolution was higher (resulting in a slightly higher mean resistance) under evolving niche choice than under fixed random choice (Figures S3.2 and S5.1).

Under all parameter ranges, populations under fixed random choice lost almost all the variation in both traits (Figures 5, S4.3, and S5.3). However, populations where plant choice evolved maintained variation in the choice trait up to the end of the simulation. For the resistance trait, variation was only maintained at very low recombination rates and high assortative mating.

**Figure 5:**
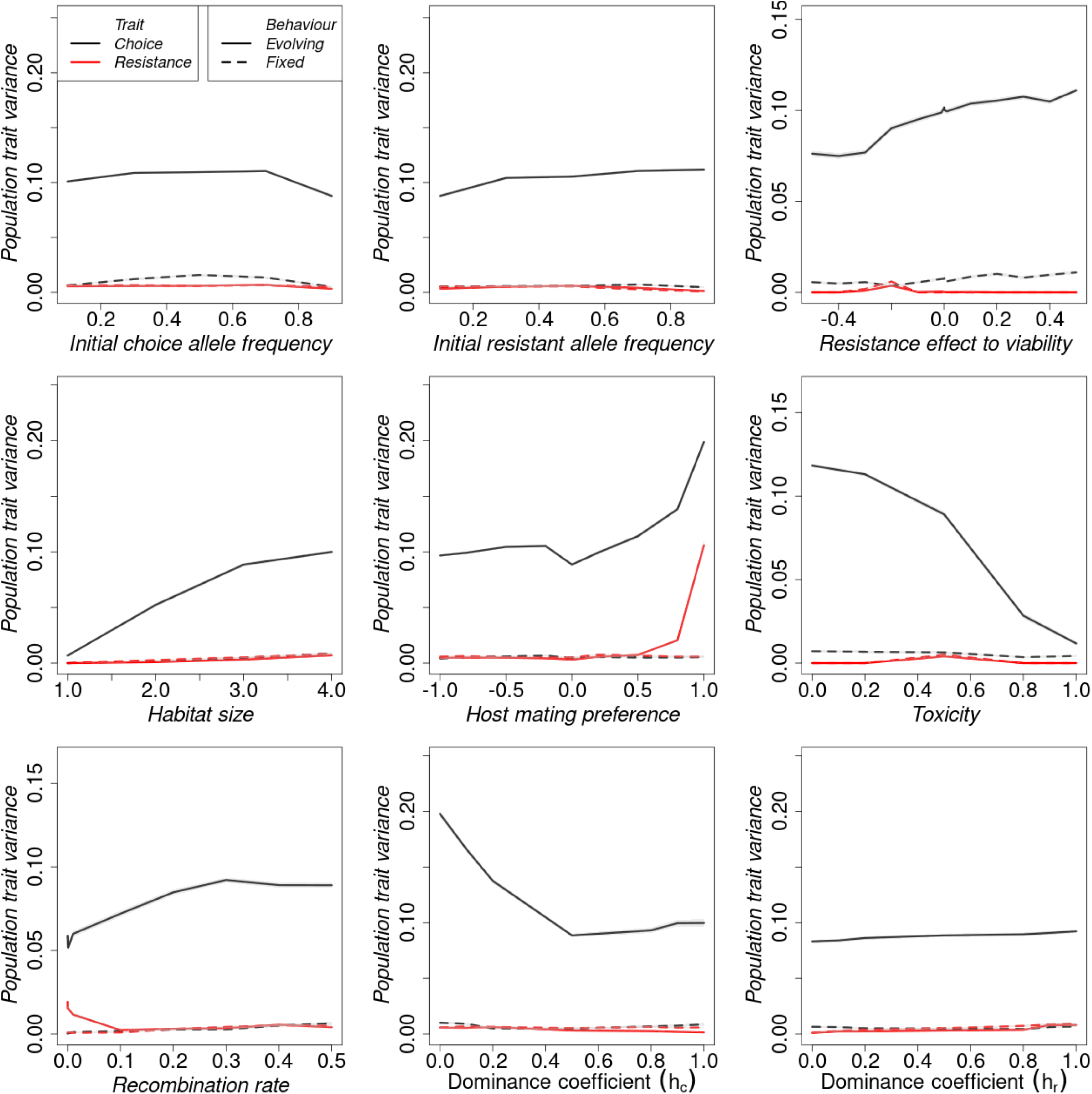
Final trait variances: Effect of model parameters on the population trait variance in resistance trait (red lines) and choice trait (black lines) between populations at the end of the simulation under fixed random choice (dashed lines) and evolving host plant choice (continuous lines). The grey area (which is too small to be seen) represents the standard error in the mean trait variance across 1000 replicate populations.

### Results for parameter values drawn from the quantitative literature survey

The above results were obtained by averaging results from 1000 populations (replicates), all of which had the same starting conditions and parameter values. However, parameters for most populations are not precisely known and vary between species. To get a better picture of the effect of niche choice evolution in phytophagous insects, for each of the 10 pest species included in the literature survey, we drew 500 parameter sets from the obtained species’ empirical parameter distributions (Figure S1.1) or from the uniform distribution (if there was inadequate data available for a given parameter). We also drew parameter sets from pooled species distributions for each parameter to artificially define “species 11” (corresponding to the distributions in Figure 2). We ran 100 replicates of the model for each parameter set under fixed random and evolving niche choice. Note that under fixed random choice behaviour, the probability of choosing a toxic plant was no longer 0.5, but also sampled from the empirical plant choice distribution. The mean variable at the end of the simulation across replicates counted as one outcome or data point in the distribution.

Consistent with observations in the previous subsections, the mean resistance and speed of resistance evolution of the species under fixed niche choice were for most species higher than under evolving niche choice (Figures 6 and S6.2), with the other three species (6, 8, and 9) showing almost no difference between the two choice scenarios. In most cases, the evolution of niche choice led to a higher variance in both the resistance trait and choice trait than when niche choice was fixed (Figures 6 and S6.3).

**Figure 6:**
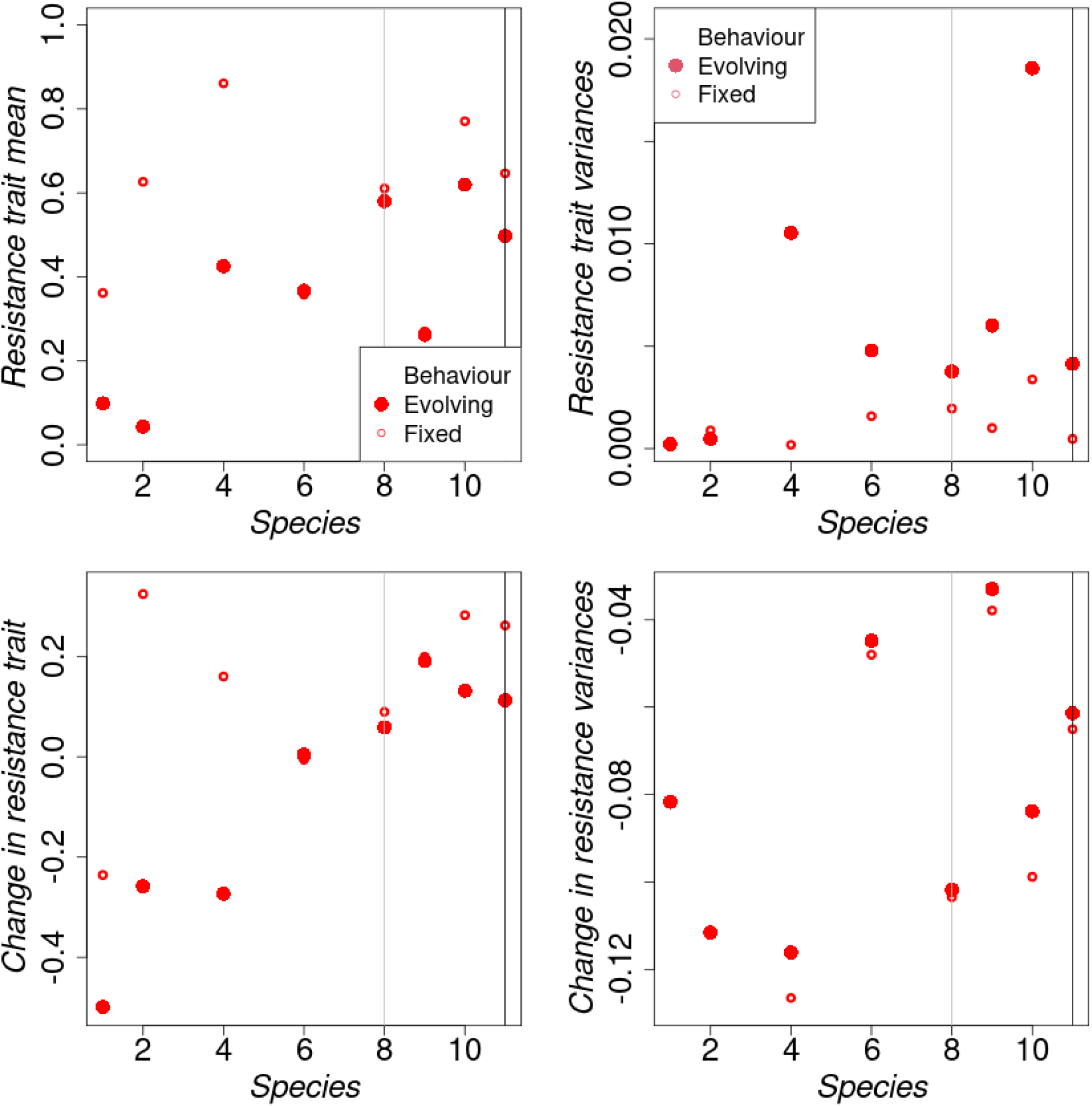
The resistance trait means and trait variances obtained in generation 200 from the simulated empirical distributions. Row 2 gives the change in trait means/variances between the initial generation and the 200th generation (negative/positive change implies a decrease/increase in trait mean or variance). The actual distributions from which these statistics were obtained are given in Figure S6.1. We omitted observations from species 3, 5, and 7 because their sampled parameter distributions had very few data points, and the number of offspring was below 1 for species 3 and 7, leading to population extinction in all (or most of) the replicates. Also, because of missing information on parameters, species 8 (grey vertical line) has all the parameters from the uniform distribution with the fixed random choice of 0.52 (based on the data for this species). The observations shown for “species 11” (black vertical line) come from simulations where the species parameter distributions are pooled together for sampling. The species names are: 1: *Plutella xylostella*, 2: *Tetranychus urticae*, 3: *Myzus persicae*, 4: *Bemisia tabaci*, 5: *Leptinotarsa decemlineata*, 6: *Helicoverpa armigera*, 7: *Aphis gossypii*, 8: *Panonychus ulmi*, 9: *Spodoptera frugiperda*, and 10: *Spodoptera exigua*.

Due to different parameter distributions (Figure S1.1), species differed in initial trait values and trait variances. We, therefore, computed the change in trait means over the course of the simulation (lower row in Figure 6). Some species became more resistant by the end of the simulation under both behaviours. However, three species, 1, 2, and 4, show a strong decrease in resistance under evolving plant choice behaviour. Niche choice also evolved, with all species showing a higher preference for the non-toxic plant by the end of the simulation (Figure S6.3). Trait variances decreased over the course of the simulation for all species under both plant choice behaviours.

## 4 Discussion

The evolution of resistance by various species to hostile environments has been observed in many fields of research, such as increased antibiotic resistance in medicine and pesticide or insecticide resistance in agriculture. In agriculture, the application of insecticides provides heterogeneous environments to pests as plants may differ in toxin concentration levels. In such micro-habitats, habitat choice or niche choice is one mechanism that can influence the fitness of individuals. This, in turn, can have a substantial influence on ecological and evolutionary processes (Morris, 2003; Berner & Thibert-Plante, 2015). Additionally, niche choice can evolve over time as a trait. However, it remains unclear how the evolution of habitat choice affects host adaptation. We simulated populations where niche choice is either random and fixed or evolves over time. We compared population dynamics from the two behavioural scenarios to evaluate the role of niche choice evolution in the evolution of toxin resistance.

We find that the joint evolution of niche choice and resistance maintains more genetic diversity within populations than when niche choice is fixed and random. This is in line with other studies with adaptive dispersal, including those considering other mechanisms of habitat choice such as matching habitat choice (Edelaar et al., 2008; Ravigné et al., 2009; Bolnick & Otto, 2013). Our study further shows that the evolution of niche choice hampers the evolution of resistance if the choice for the non-toxic habitat is initially predominant. When the choice for the toxic habitat is predominant initially, resistance evolution under evolving niche choice behaviour is more or less similar to the fixed random choice behaviour, but faster. Essentially, this means that the more individuals are exposed to the challenging habitat, the faster the populations become resistant. This result confirms and extends previous findings from models of evolutionary rescue and source-sink dynamics where a larger number of individuals exposed to the challenging habitat results in a higher probability of local adaptation (Czuppon et al., 2021; Tomasini & Peischl, 2022).

Our model is close to one of the scenarios in Berner & Thibert-Plante (2015), genetic habitat preference with a dispersal probability of 1. However, in their model, variation arises through mutations. They find that with random dispersal, choice trait evolution requires large dispersal probabilities, asymmetric carrying capacities and large selective differences between habitats. In our model, the number of individuals showing preference for the toxic plant increased over time under a wide range of model parameters when habitat choice was genetically determined. The individuals in our model had 100% dispersal probability, and choice evolution generally required conditions that maintain similar population sizes in both habitats (Figure S3.1). That is, conditions that increase survival in the toxic habitat, such as large habitats, high fitness benefits associated with resistance, high recombination rates, low toxicity levels in the toxic habitat, and low dominance of the choice allele.

Our study offers a unique contribution by incorporating genetic realism through the consideration of different recombination rates and dominance coefficients that were not taken into account by previous research. Recombination plays a crucial role in shaping genetic diversity and influencing natural selection by breaking existing gene combinations and forming new ones. This reduces interference between selected loci (Dumont, 2020). High recombination rates can also decrease the loss of genetic variance and, therefore, increase the overall response to selection (Moradigaravand et al., 2014; Battagin et al., 2016). We find that the recombination rate can influence the rate of niche choice evolution, population trait variance, and population resistance. By increasing the recombination rate between choice locus and resistance locus, populations become more resistant. We conjecture that this is because, at a high recombination rate, sensitive types that choose to go to the non-toxic habitat cannot easily form through linkage. That is, every gene copy at the resistance locus will regularly find itself in an individual in the toxic habitat, which leads to higher effective selection pressures on the resistance allele.

Dominance coefficients can influence the fitness of individuals in different habitats. In the case of pesticide resistance, for example, even individuals with one copy of a dominant resistance allele will be resistant to the pesticide, increasing the frequency of the resistant individuals as they are more likely to survive and reproduce in habitats where the pesticide is present than susceptible phenotypes (Stratonovitch et al., 2014). Dominance coefficients of insecticide resistance alleles have been found to range from 0 (completely recessive) to 1 (completely dominant) (Bourguet et al., 2000). Pest management strategies can increase the fitness of pests in treated areas via the selection of dominance modifiers (Bourguet et al., 2000; and other references therein). In our model, we assume a uniform distribution of dominance coefficients over the range from 0 to 1 for both resistance and choice alleles due to inadequate information from the surveyed literature. As expected, we find that dominance for the resistance allele increases the average population resistance. Similarly, dominance for choice increases the average choice trait. However, dominance for choice also influences resistance evolution. With increasing dominance for choice, more heterozygotes go to the non-toxic plant. Thus, overall, fewer individuals are exposed to the toxic plant, leading to weaker selection for resistance and lower average resistance traits. On the other hand, under our default setting with a low initial frequency of the resistance allele, resistance alleles are quickly lost in the population due to genetic drift and the cost of resistance on survival. Mating between heterozygotes becomes less frequent by the end of the simulation, rendering the effect of resistance dominance weak as populations produce more susceptible homozygotes.

The evolution of resistance and choice also depended on ecological parameters such as the degree of assortative mating. Host mating preference reduces the exchange of genes between different phenotypes in a population and, in extreme cases, can result in speciation (Webster et al., 2012; Servedio & Boughman, 2017; Schumer et al., 2017). We find that strong assortative mating reduced mean population resistance. Strong assortative mating was also required for the maintenance of high variance in the resistance trait. In our model, strong assortative mating would lead to a higher proportion of allele copies at the resistance locus that rarely “see” the toxic environment and, therefore, lead to lower resistance levels in the population. Due to the restriction in the exchange of genes between habitats, the population can easily maintain both highly resistant and susceptible individuals. Since none of the processes we consider is sex-specific, we simulated hermaphroditic populations as a logistic assumption (Servedio et al., 2014). We do not expect qualitative changes in the results with separate sexes.

Our populations were subjected to both global density regulation (via the carrying capacity parameter) and local density regulation (via the habitat size parameter). However, for the parameter settings we investigated, local density regulation was generally strong, and population sizes (recorded after local density regulation) never reached the global density cap. We often observe allele fixations due to generally low population sizes maintained, but looking across the replicates, we also see that not always the same allele becomes fixed in one trait. Increasing habitat size results in weaker local density regulation and, therefore, habitats maintain larger populations. This increases phenotypic variation within populations and, consequently, the probability of genetic polymorphism at the choice locus. Ravigné et al. (2009) found that local density regulation results in evolutionary branching into two specialist types. They consider asexuals, which makes diversification or branching easier. For diploids, as already mentioned, Berner & Thibert-Plante (2015) also have a very similar setup to ours, with high assortative mating (mating after dispersal) and local density regulation. Our model setup compares best to their genetic dispersal mechanism with dispersal propensity 1, where they find very low divergence in that case.

### Insights from the quantitative literature survey and implications for the management of insect pests

Most of the articles included in our literature review are in the context of insecticide resistance. Given the continued use of insecticides and the ubiquitous nature of resistance, it is unsurprising that host plant toxicity is very high, with a median of 0.87. Alleles associated with insecticide resistance are thought to be rare in populations prior to selection (Roush & McKenzie, 1987; Tabashnik, 1994). We found that the median resistance allele frequency is much higher than what was found in early empirical studies (Gould et al., 1997; Génissel et al., 2003; Tabashnik et al., 2008, 2000), probably because strong selection pressures had already acted before the studies in our survey. However, the median resistance allele frequency in our survey is similar to or lower than the more recent findings (Freeman et al., 2021a; Kaduskar et al., 2022). In line with our literature survey, a recent meta-analysis found that about 60% of studies showed fitness costs associated with insecticide resistance (Freeman et al., 2021b). But for a clearer understanding of the nature of fitness costs and benefits, the effects of population density must be considered, and population sizes should be standardized. The genetic background and relatedness of the insects are also necessary information for a broader understanding of fitness costs in nature (ffrench-Constant & Bass, 2017).

Like in the main results where the choice for the non-toxic habitat is initially predominant, most populations under fixed random behaviour became more resistant than those under evolving random choice. Generally, trait evolution across species under fixed random choice behaviour from sampled parameter distributions showed increased resistance over time in almost all species. This could be due to the high relative toxicity obtained from the literature, as only resistant individuals in a population can survive at the toxic plant, although other species parameters like initial allele frequency and resistance effects on fitness could have played a substantial role. However, under evolving choice behaviour, some species showed reduced resistance, and others showed increased resistance at the end of the simulation. It is unclear why we get such mixed observations.

In agricultural fields, non-toxic plants, known as refuges, are commonly maintained to slow down the evolution of resistance (Gould, 1998). This strategy works under specific conditions like low initial resistance allele frequency, high fitness costs of resistance, and large refuge sizes (Carrière & Tabashnik, 2001; Crowder & Carrière, 2009; Jongsma et al., 2010). In our model, we do not focus on the effect of introducing a non-toxic plant *per se* but consider a field where both refuges and treated plants are equally present. In addition to the above conditions that slow down resistance evolution in the presence of refuge plants, our model also shows that strong local density regulation, high dominance of choice alleles, low dominance of resistance alleles and evolution of niche choice slow down population resistance. The success of the refuge strategy in the field has also been shown experimentally, although the effectiveness is not as dramatic as previously thought (Jin et al., 2015; Carrière et al., 2012; Huang et al., 2011). This highlights the importance of gathering more empirical and quantitative data on the ecology, behaviour, and genetic mechanisms of important agricultural pests to design the best control strategies.

### Limitations and future work

We considered only one mechanism for the joint evolution of niche choice and resistance, a so-called two-allele mechanism where separate loci encode choice and resistance. Other mechanisms, e.g., with one locus, with pleiotropic effects, or habitat imprinting, can have quite different results (Berner & Thibert-Plante, 2015; Bono et al., 2017). Our model maintained quite small population sizes and did not allow for recurrent mutations. Thus our results are directly applicable to small experimental populations on a short time scale, but more exploration is needed to see how the dynamics play out in large natural populations and on a longer time scale where new mutations can appear.

For many species, despite being important agricultural pests, we found little information on the genetic basis of choice. We considered simple genetics, such as a single bi-allelic locus per trait. Increasing the number of loci per trait would likely slow down the evolution of resistance evolution (Berner & Thibert-Plante, 2015). For dominance coefficients and recombination rates, we explored all the possible ranges of parameter values. We found both qualitative and quantitative effects of these parameters on the joint evolution of niche choice and resistance. Studies investigating choice behaviour and its genetic basis in the context of resistance are scarce, possibly due to the difficulties in accurately quantifying behavioural variations and the polygenic nature of most behavioural traits. Obtaining such information, e.g., via crossing experiments, is critical to better understanding the evolution of resistance and how it is influenced by niche choice.

### Conclusions

Our study agrees with existing literature that niche choice evolution maintains genetic diversity. Most importantly, our study shows that the evolution of niche choice slows down the evolution of resistance in heterogeneous habitats under a wide range of parameters. In addition, our study reveals the importance of recombination rates and choice dominance in shaping evolutionary outcomes. However, more empirical studies are needed to identify genetic factors influencing individual preferences in heterogeneous habitats.

## Supporting information

These are R scripts and data files used for generating results.

## Supplementary Material

### S1 Quantitative literature survey

**Table S1.1:** Top 10 agricultural pests as of February 2023 in the Arthropod Pesticide Resistance Database (APRD) (Mota-Sanchez & Wise, 2023). To create the list, we first used the search feature to browse through the APRD database by insect order. Then, a list of herbivore species with >100 global resistance cases (#Cases in APRD) was made. From that list, we took the top 10 species with the most resistance cases. Each species from that list was then individually examined in the database to extract the number of compounds against which resistance cases are reported (#Compounds). Finally, the species were ranked based on the descending order of the number of compounds.

| Rank | Scientific name | Common name | #Cases in APRD | #Compounds | Order |
| --- | --- | --- | --- | --- | --- |
| 1 | *Plutella xylostella* | Diamondback moth | 1010 | 101 | Lepidoptera |
| 2 | *Tetranychus urticae* | Two-spotted spider mite | 557 | 96 | Trombidiformes |
| 3 | *Myzus persicae* | Green peach aphid | 487 | 84 | Homoptera |
| 4 | *Bemisia tabaci* | Sweet potato whitefly | 697 | 68 | Homoptera |
| 5 | *Leptinotarsa decemlineata* | Colorado potato beetle | 306 | 56 | Coleoptera |
| 6 | *Helicoverpa armigera* | Cotton bollworm | 891 | 55 | Lepidoptera |
| 7 | *Aphis gossypii* | Cotton aphid | 302 | 51 | Homoptera |
| 8 | *Panonychus ulmi* | European red mite | 203 | 48 | Trombidiformes |
| 9 | *Spodoptera frugiperda* | Fall armyworm | 200 | 45 | Lepidoptera |
| 10 | *Spodoptera exigua* | Beet army worm | 675 | 43 | Lepidoptera |

#### S1.1 Literature review

Literature review was initially conducted on Web of Science using general topic related search strings and combinations, e.g.: survival insecticide, oviposition choice, fitness cost, mating choice, etc. Since this resulted in many unrelated articles passing the filter, an optimised search string was designed for each species as shown below:

TS=((host OR oviposition) AND choice AND (insecticide OR pesticide) OR fecundity cost OR resistance cost OR mating choice OR assortative mating OR ((number OR frequency) AND loci AND (resistance OR host choice))) AND (TI= (Species name) OR AB= (Species name) OR AK= (Species name))

In February 2023, the marked lists were created in Web of Science for each of the ten species. In total for all species and topics, 802 articles were found and screened. Out of these, data was extracted from 229 articles (28.68%).

#### S1.2 Criteria for inclusion of articles

Studies in which the key parameters were experimentally determined were considered. Experiments in which the pests were directly exposed to leaf discs or intact plants for feeding and/or oviposition were exclusively considered. Studies in which experiments were performed using artificial diet mixed with insecticide were not included. Reviews and book chapters were excluded. Articles which were written in languages other than English were not considered due to authors’ limitations. In cases where the full text was not available, the authors were contacted. If there was no response in due time, those studies were excluded. The reason for the exclusion of every excluded article is specified individually in the dataset (Subsection S7).

#### S1.3 Data extraction and parameter distributions

From all articles that passed the inclusion criteria, relevant raw data were extracted from the text, tables, or figures. Considering each article separately, the parameters were calculated from the raw data as shown below.

##### Toxicity

To calculate toxicity at each host plant *i*, 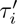, and the resistance cost on viability 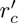, the percentage mortality of each genotype (susceptible or resistant) was obtained on each host plant. The toxicity at host plant *i* was taken as the percentage mortality of the susceptible genotype at the respective plant *i* (*i* = *{*1, 2*}*).

To align with the model where plant 1 is assumed to be non-toxic (*τ*_1_ = 0), we instead used relative toxicity by first getting the survival at both plants (i.e., 1-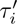). We divide survival at both host plants by survival at plant 1 (expected to be the highest since it is the non-toxic host). With this, we get relative survival at plant 1 as 1, while at plant 2, it is at most 1 (we removed cases with higher survival at plant 2 than at plant 1). We then subtract both values from 1 to get the relative mortality at both plants that act as toxicity (with *τ*_1_ = 0, 0 *≤ τ*_2_ *≤* 1).

##### Number of eggs to number of offspring

From the survey, we obtained the average number of eggs produced by a susceptible female on a non-toxic host plant, Λ*^t^*. We arbitrarily assumed only 1% of the eggs result in offspring. Therefore, we divided the number of eggs per female by 100 to get the mean number of offspring, 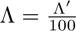.

##### Resistance costs on viability

We calculated the resistance effect on viability, 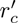, as

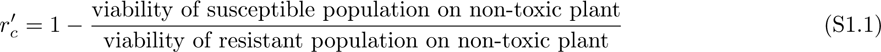

where negative values of 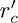 imply resistance cost on viability while positive values indicate resistance benefits on viability.

Without local density regulation in our model and assuming zero toxicity at the non-toxic host, each offspring at the non-toxic host is viable with probability *v* = (1 + *r_c_z_r_*), where *z_r_* is again the individual resistance trait and *r_c_* is the resistance effect on viability. Now, from Equation (S1.1) and assuming susceptibles have zero resistance while resistant individuals have *z_r_* = 1, we have

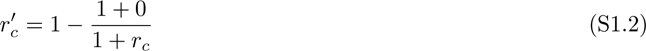

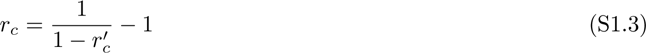

##### Resistance costs on fecundity

Similarly, to get the resistance effect on fecundity, 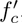, and the choice index, 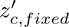, we extracted the number of eggs laid on each host plant and by each genotype separately. We obtain 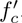 from

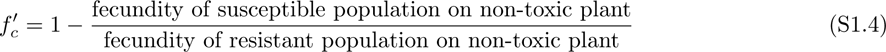

where negative values of 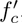 imply resistance cost on fecundity while positive values indicate resistance benefits on fecundity.

From our model, the expected number of offspring is given by *λ* = (1 + *f_c_ · z_r_*)Λ, where *f_c_* is the fecundity cost, *z_r_* is the resistance trait and Λ is the fecundity of a non-resistant individual. We assume that a resistant individual has full resistance of *z_r_* = 1 while a susceptible individual has *z_r_* = 0. With these assumptions, together with Equation (S1.4), we get

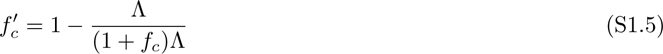

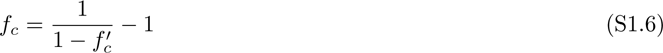

##### Choice index

The choice index that shows host plant preference by an individual was calculated using the number of eggs laid on toxic and non-toxic plants (both of which were available in the same amounts) and then averaged across all individuals, irrespective of genotype.

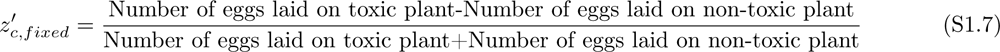

Here, positive values indicate a preference for the toxic host, while negative values indicate a preference for the non-toxic host. If the choice was reported as a proportion, it was converted to the choice index appropriately.

We use the choice index obtained from the data as the fixed random choice parameter for between the toxic and non-toxic niches by the individuals. To align with our model range, we therefore transform the given values from the interval (−1, 1) to (0, 1) by 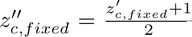. Since values of 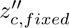 indicate the probability of choosing a toxic host, 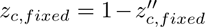 random host plant choice. indicates the probability of choosing a non-toxic host, with 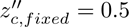 indicating

##### Host mating preference

In addition, the number of mates chosen from the same host and the other host were extracted to calculate the level of host mating preference, 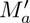, as

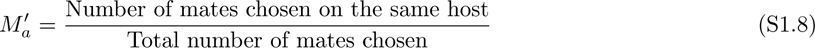

where 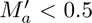 indicates a preference for mates on different host plants, 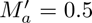 indicates random mating, and 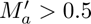 indicates a preference for the mates on the same host plant.

To align with the model implementation, we convert the values from the interval (0, 1) to (−1, 1) by 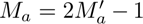 such that negative values of *M_a_* correspond to the tendency of an individual to choose a mate from a different host plant, positive values indicate the tendency to choose from the same host plant while a value of 0 indicates that individuals randomly choose their mates from the available host plants. This conversion assumes equal densities of individuals at the focal host plant and the other host plants.

##### Other parameters

Lastly, other parameters that were extracted include the frequency of the resistance allele, *p_r,_*_0_, and the number of resistance loci *n_r_*. No similar information was found for the corresponding choice parameters. The resultant parameter distributions are shown in Figure 2.

#### S1.4 Literature data distributions for each species

**Figure S1.1:**
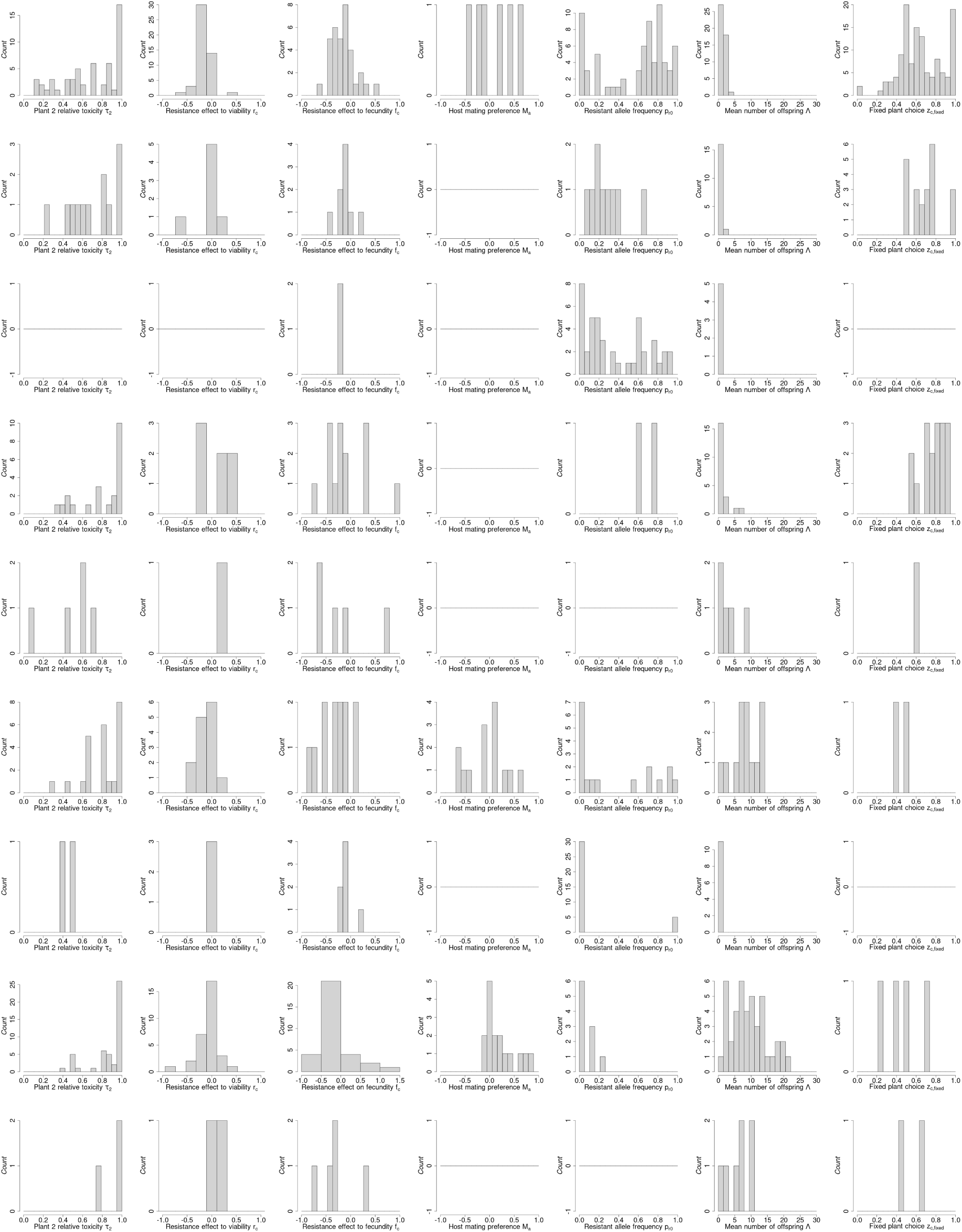
Parameter distributions for each species. Each row corresponds to one species.Also, row *i* corresponds to species with rank *i*, except species with rank 8 is omitted because there were no data points apart from one data point under the choice index. Toxicity at plant 2 assumes that toxicity at plant 1 is 0 (i.e., we show relative toxicity).

### S2 Quantifying density regulation

We did not find enough information about host density-dependent population regulation from the literature survey. We, therefore, ran a density-survival experiment using Colorado potato beetles (*Leptinotarsa decemlineata*). We measured survival at different larvae densities.

For this experiment, we used a strain collected from Spain in 2012 and maintained in the greenhouse since. The beetles were reared on potato plants (*Solanum tuberosum*). The beetles and the plants were maintained in a greenhouse at 24°C under a long day photoperiod (16h:8h L:D) following the protocol in (Edison et al., 2024). The experiment was conducted in insect cages of size 85cm x 45cm x 55cm. Two trays (39cm x 29cm x 7cm) of potato plants were placed in each cage. For assessing the effect of density on survival, first instar larvae were placed directly on potato leaves at varying densities (3, 10, 25, 45, 150, 300). The larvae were allowed to feed and develop up to pupation. The number of surviving individuals was recorded at the end of the experiment and used to fit a density-dependent survival fitness function (Equation (3) and Figure S2.1). The observed relationship between initial beetle density and fitness (per capital survival) was estimated by linear model after log transformation.

**Figure S2.1:**
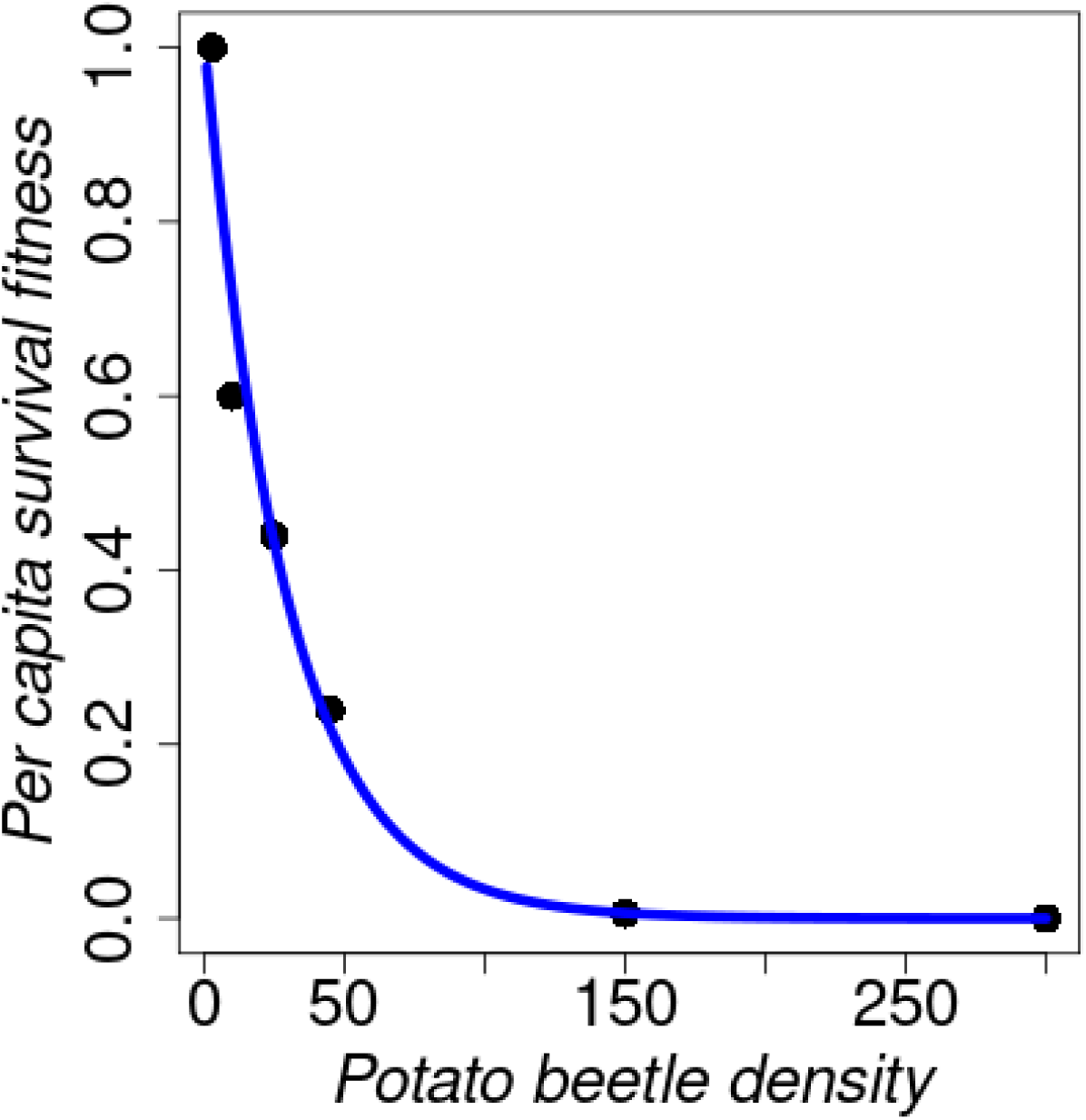
Data (black points) used for fitting the density-dependent survival fitness function (blue curve) in Equation (3).

### S3 Supplementary figures based on default parameters in Table 1

#### S3.1 Effect of the model parameters on population sizes

**Figure S3.1:**
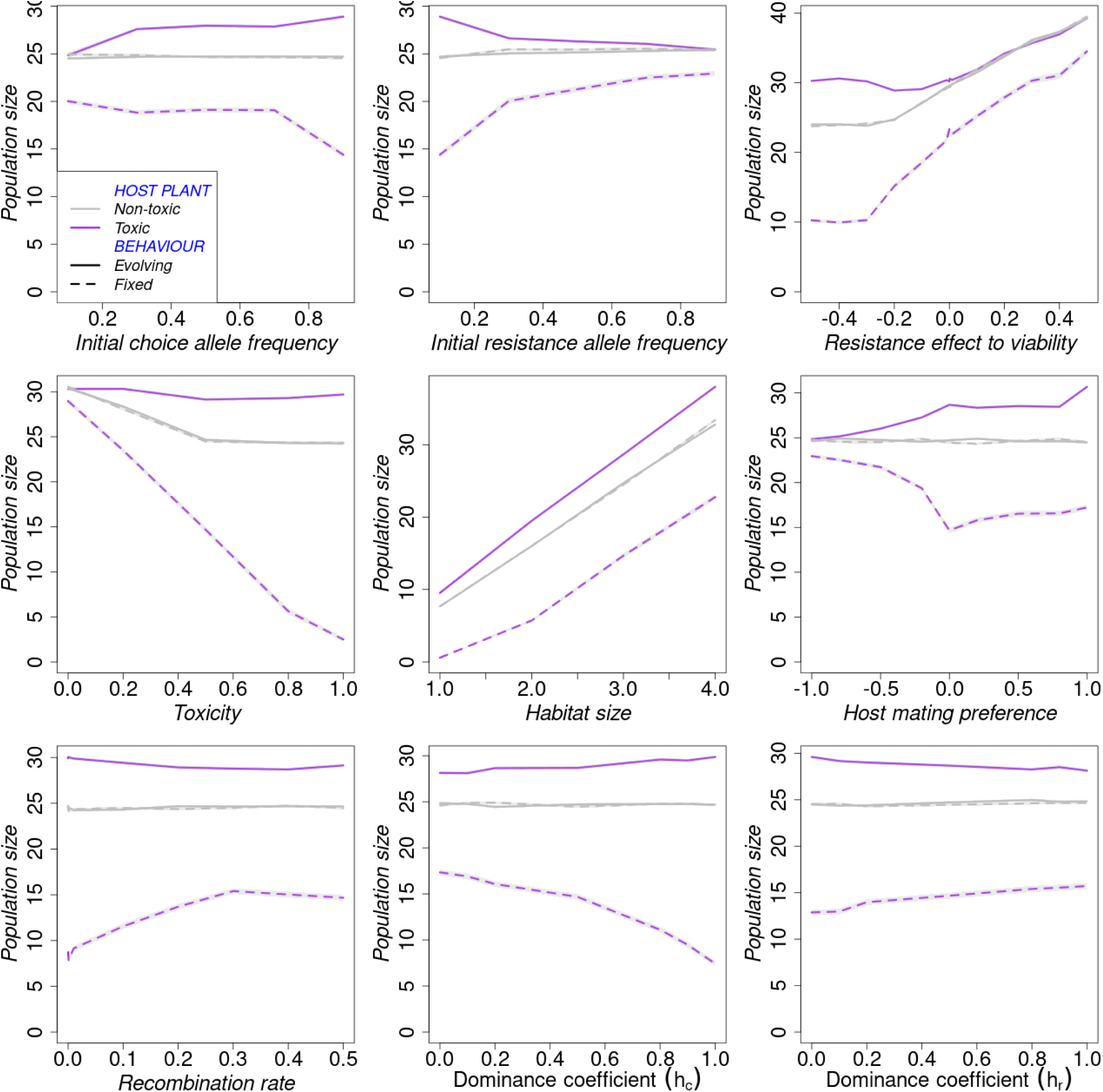
Final population sizes under default parameters: Effect of model parameters on the average population size at the end of the simulation.

#### S3.2 Effect of model parameters on the speed of trait evolution

**Figure S3.2:**
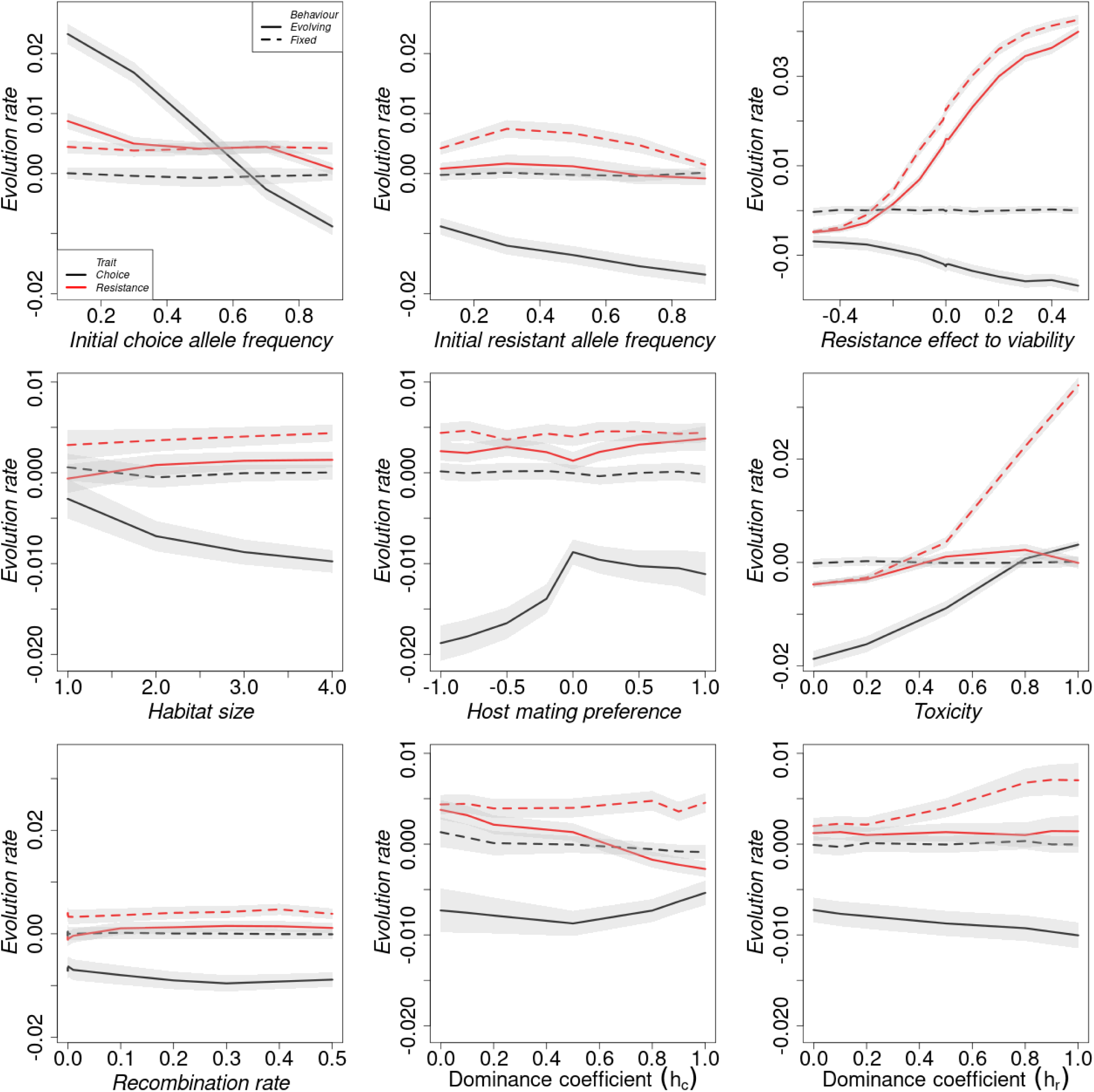
Rate of trait evolution under default parameters: The effect of model parameters on the evolution rate of the resistance trait (red lines) and choice trait (black lines) under fixed host plant choice (dashed lines) and evolving host plant choice (continuous lines). The grey-shaded area shows the standard error. The values indicate the average change in trait per generation between the 1st and 20th generations.

#### S3.3 Effect of resistance effect on fecundity

**Figure S3.3:**
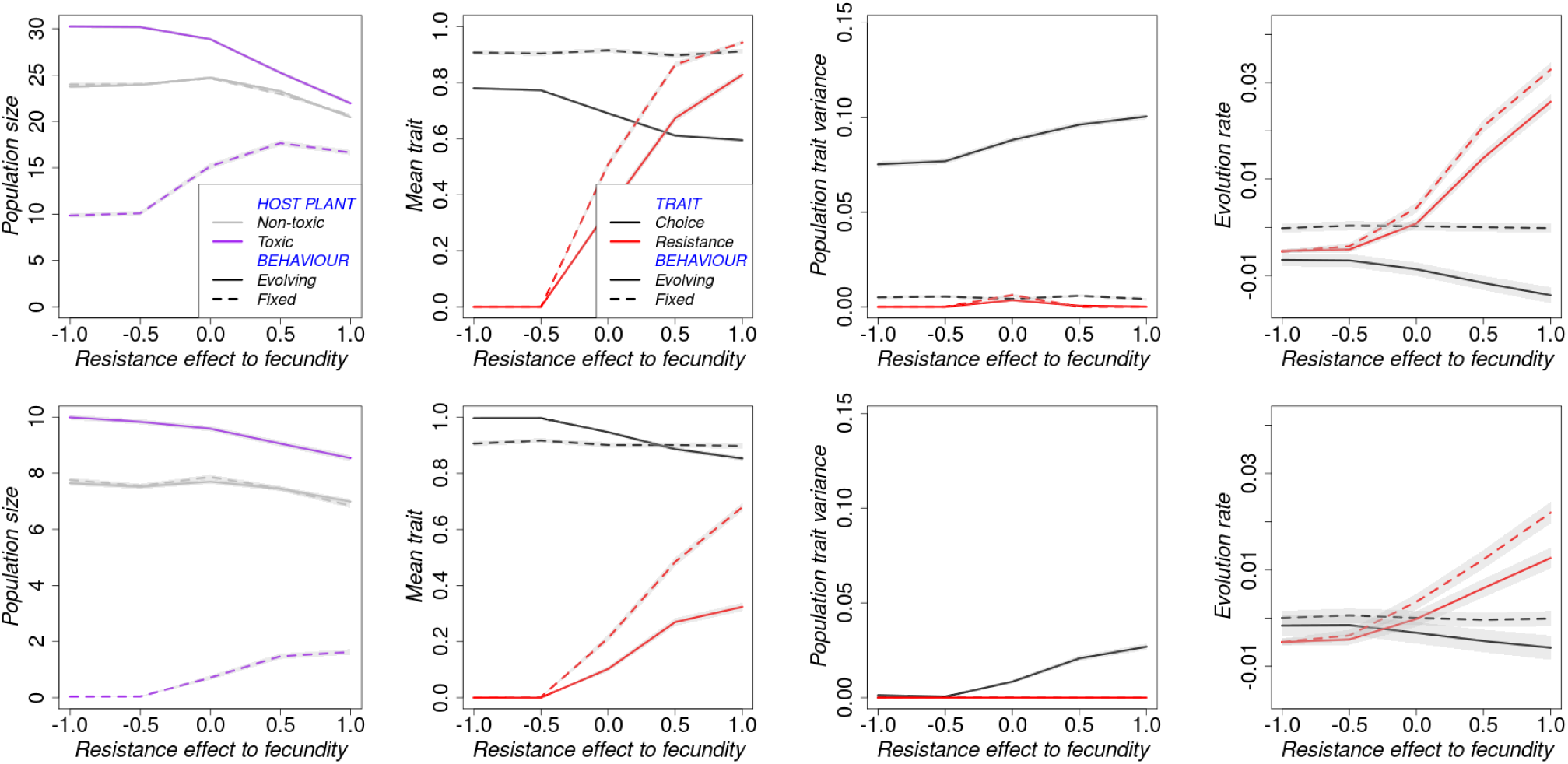
Effect of resistance effect on fecundity on population dynamics in large habitats, *w* = 3, (row 1) and in small habitats, *w* = 1, (row 2). Default parameters are as given in Table 1.

### S4 Parallel figures to those in the main section but with low habitat size, ***w* = 1**, as default

#### S4.1 Speed of adaptation

**Figure S4.1:**
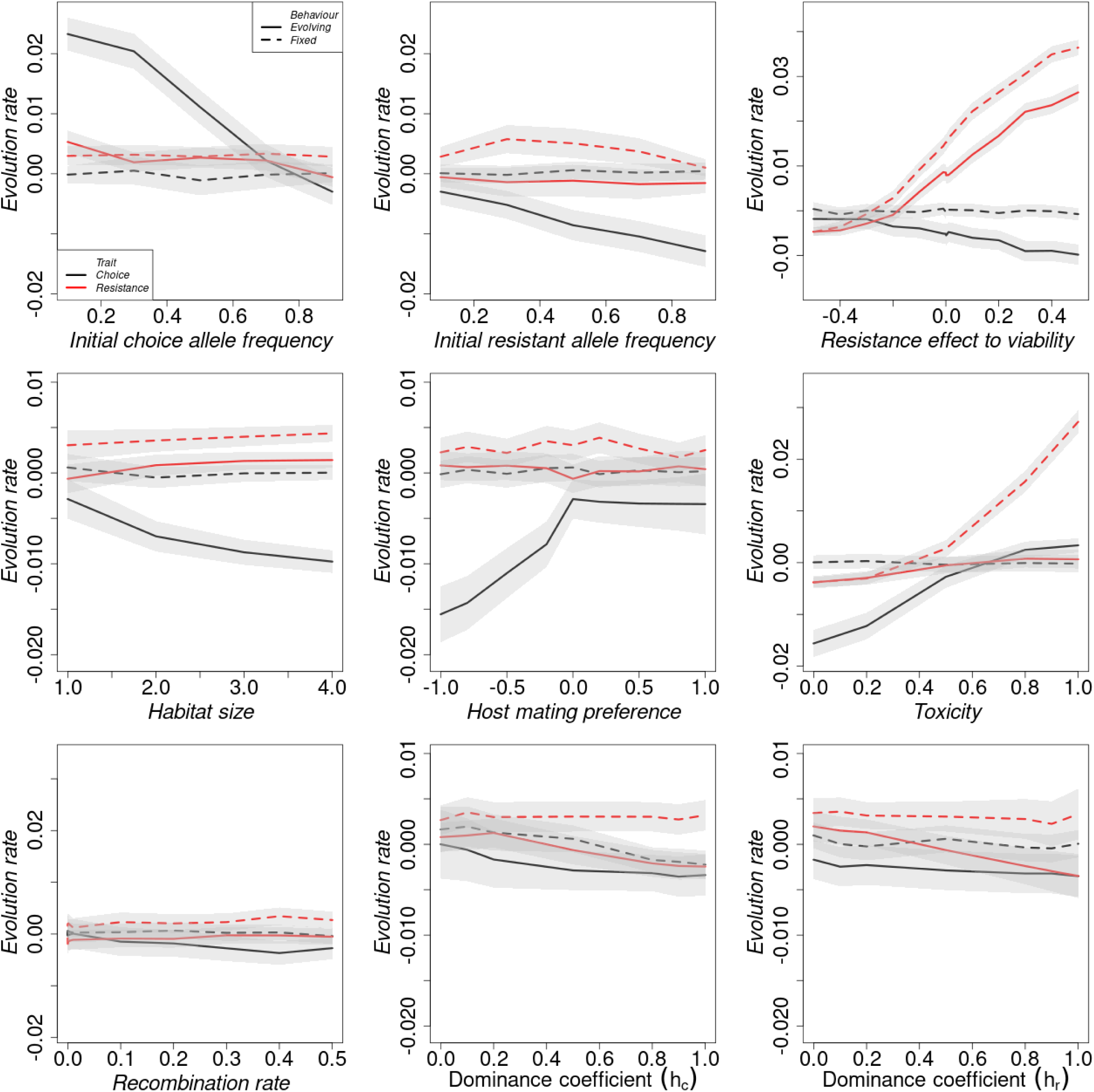
Rate of trait evolution in small habitats, *w* = 1: The effect of model parameters on the evolution rate of the resistance trait (red lines) and choice trait (black lines) under fixed host plant choice (dashed lines) and evolving host plant choice (continuous lines). The grey-shaded area shows the standard error. The values indicate the average change in trait per generation between the 1st and 20th generations.

#### S4.2 Mean traits

**Figure S4.2:**
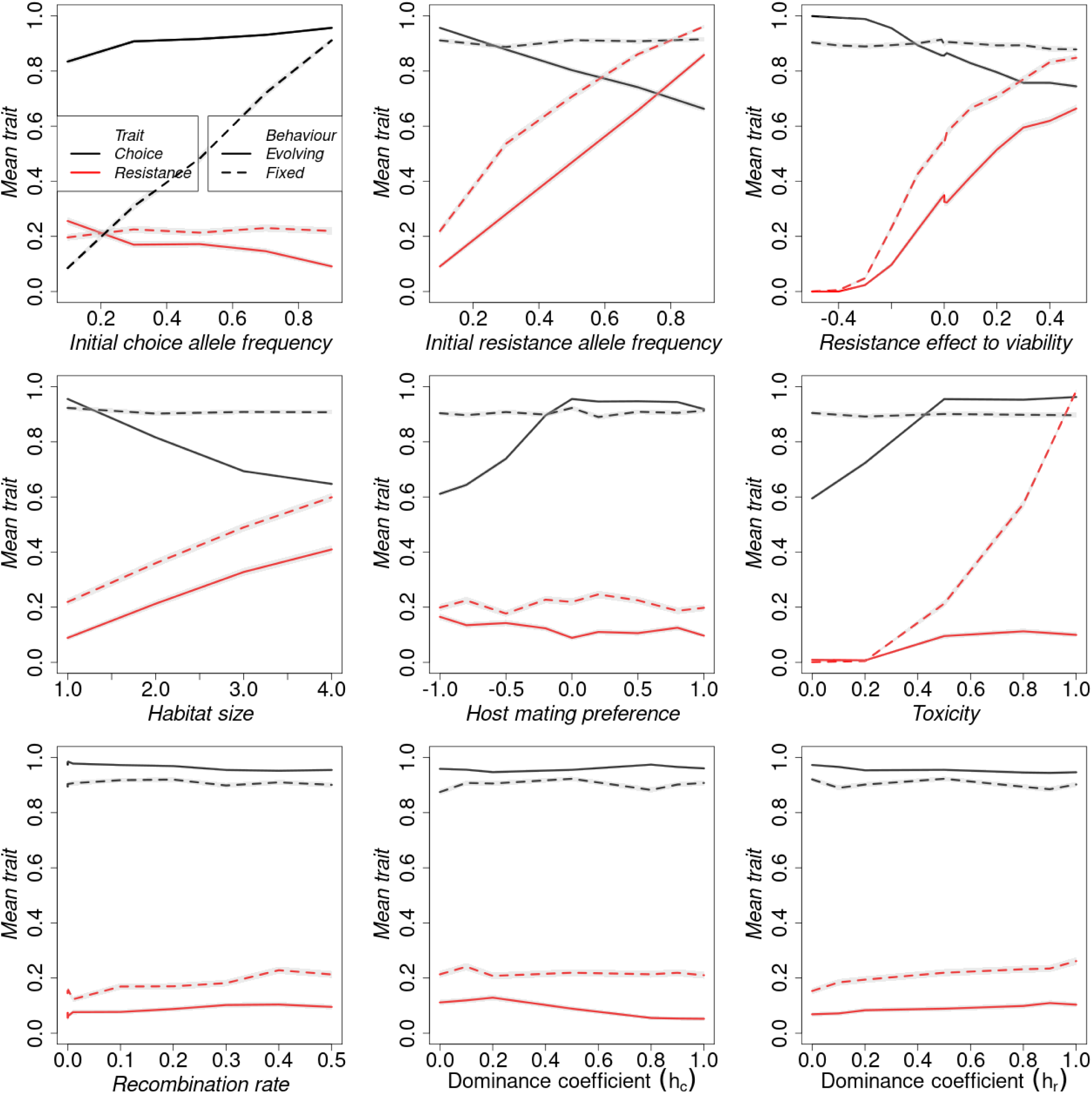
Final mean traits in small habitats, *w* = 1: Effect of model parameters on the average resistance trait (red lines) and choice trait (black lines) at the end of the simulation under fixed random choice (dashed lines) and evolving host plant choice (continuous lines).

#### S4.3 Population trait variance

**Figure S4.3:**
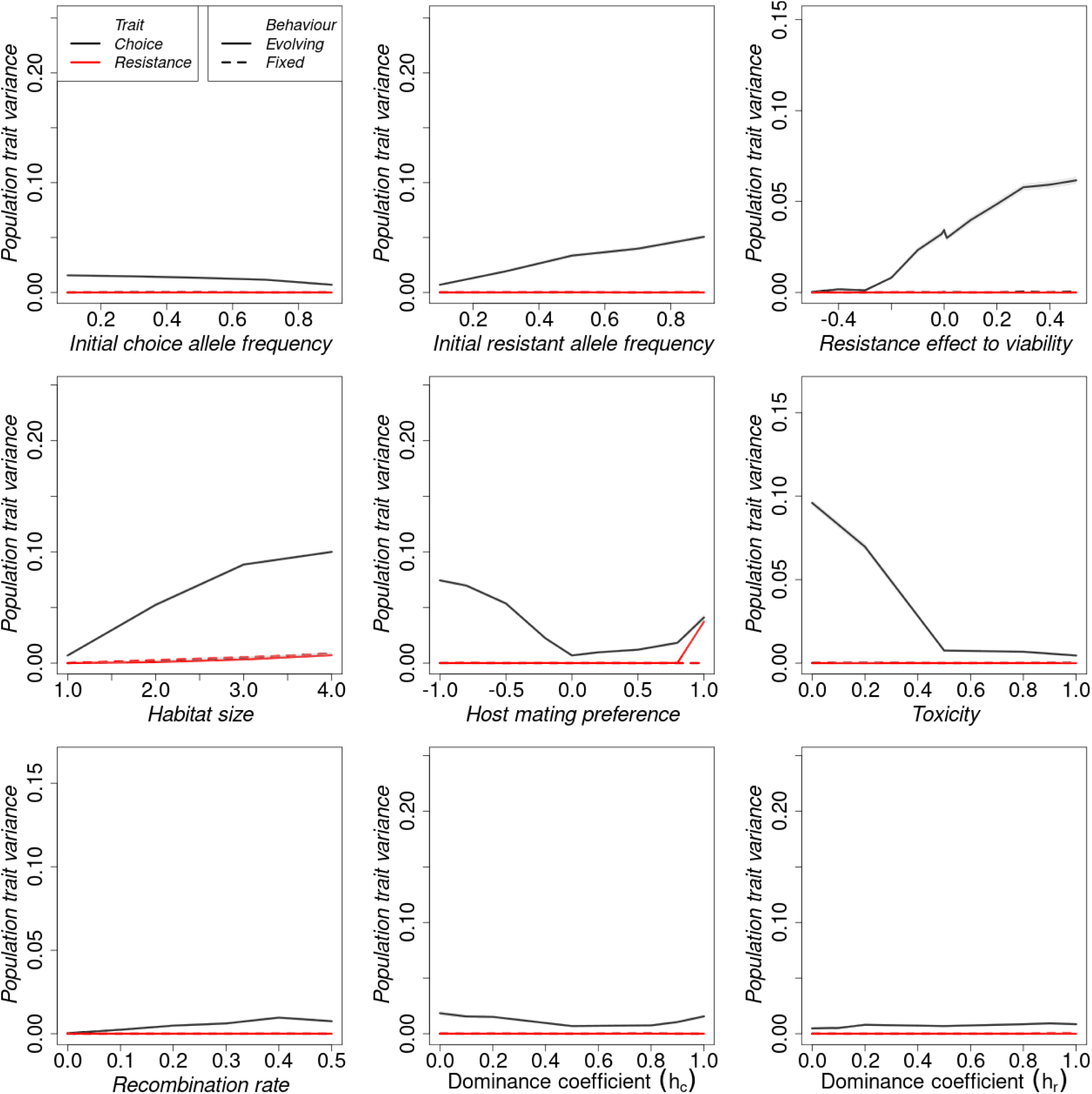
Final trait variance in small habitats, *w* = 1: Effect of model parameters on the population trait variance in resistance trait (red lines) and choice trait (black lines) between populations at the end of the simulation under fixed random choice (dashed lines) and evolving host plant choice (continuous lines). Plotted is the mean of trait variance across replicates.

### S5 Results from large habitats, ***w* = 3**, and low initial choice allele frequency for the non-toxic habitat, ***p_c,_*_0_ = 0.1**, as default

#### S5.1 Speed of adaptation

**Figure S5.1:**
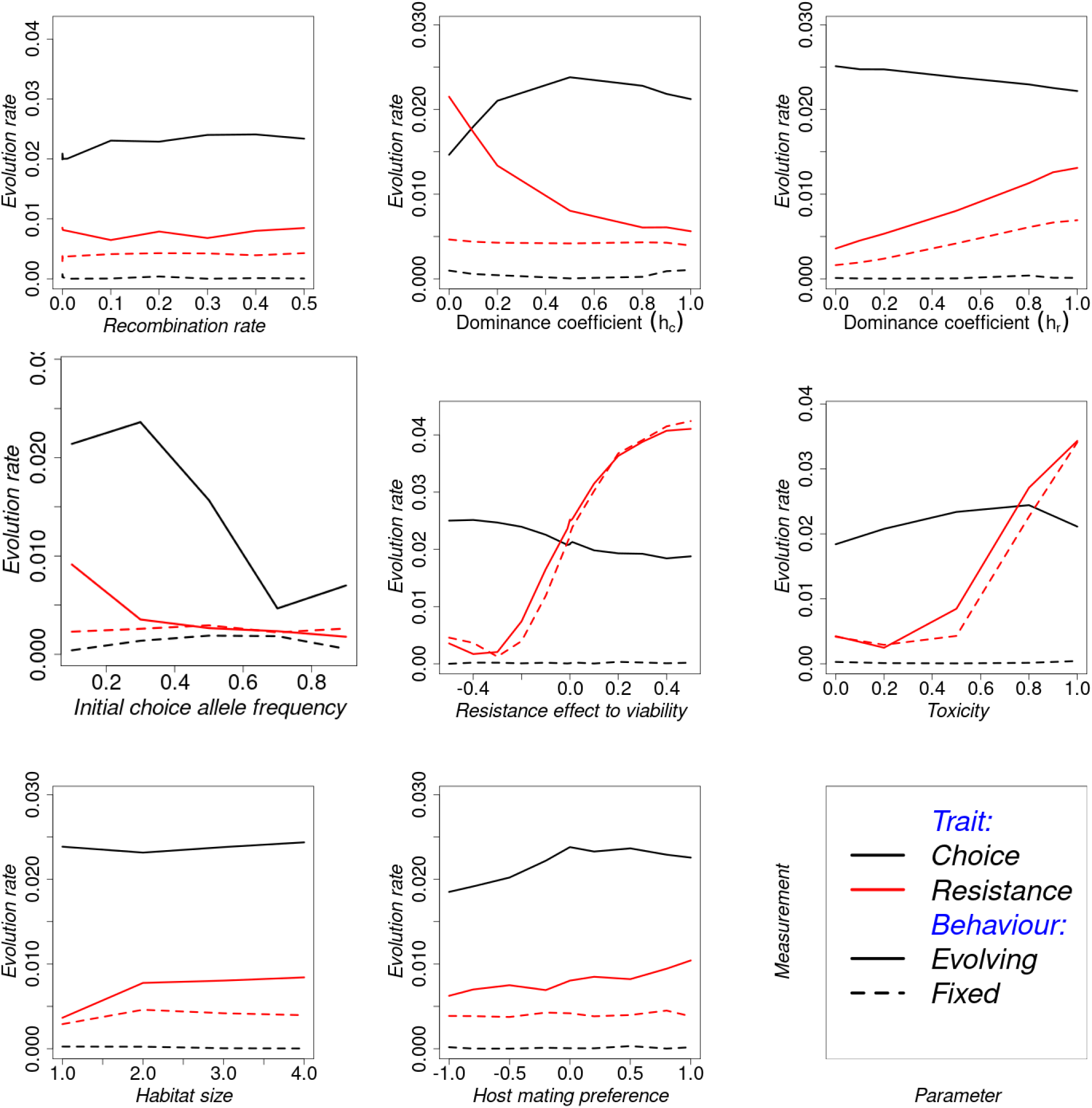
Rate of trait evolution in large habitats, *w* = 3 but low initial choice allele frequency *p_c,_*_0_ = 0.1: Effect of model parameters on the evolution rate of the resistance trait (red lines) and choice trait (black lines) under random behaviour (dashed lines) and choosy behaviour (continuous lines). The values indicate the (absolute) average change in trait per generation between the 1st and 20th generations.

#### S5.2 Mean traits

**Figure S5.2:**
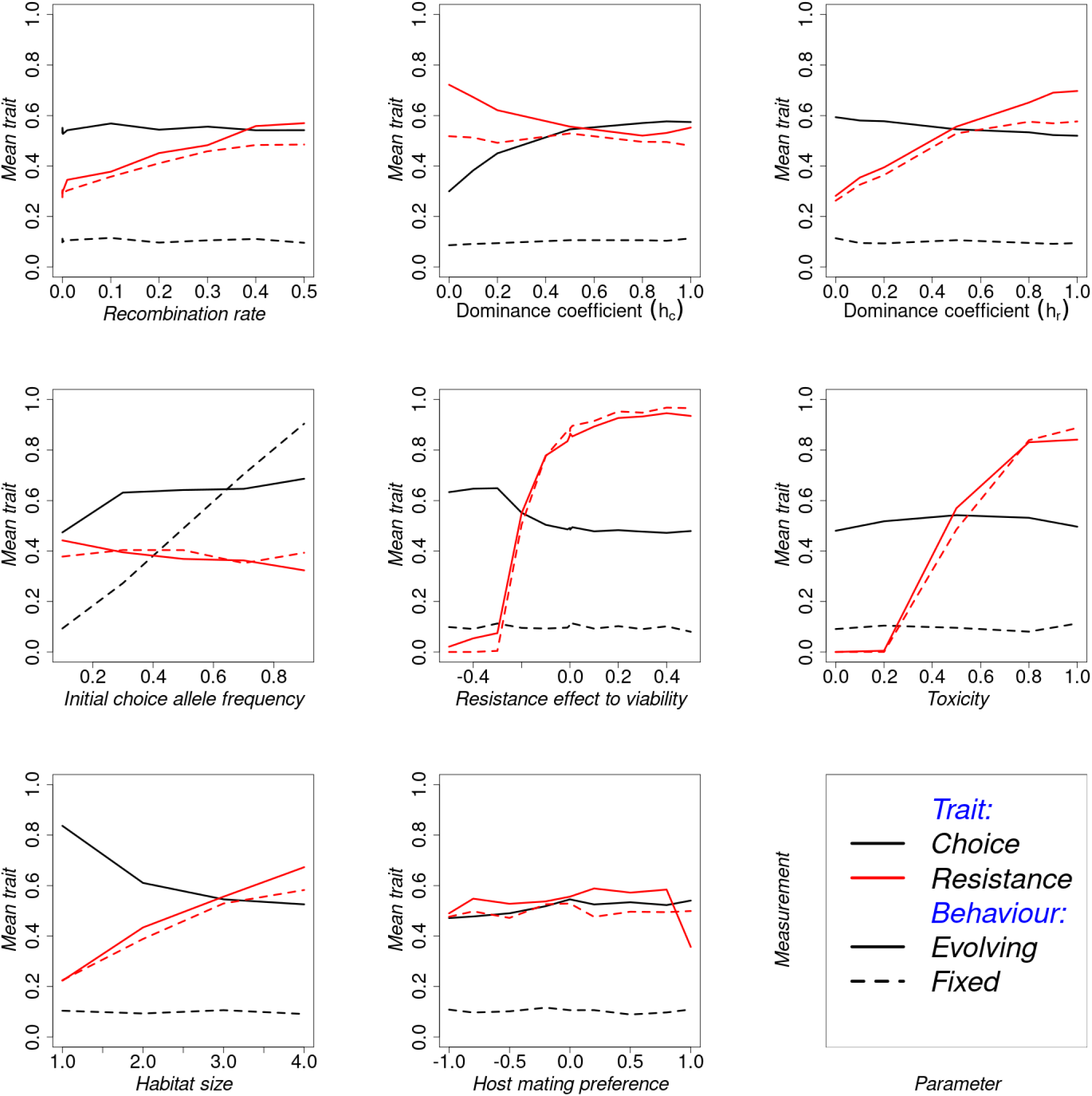
Final mean traits in large habitats, *w* = 3 but low initial choice allele frequency *p_c,_*_0_ = 0.1: Effect of model parameters on the average resistance trait (red lines) and choice trait (black lines) at the end of the simulation under random behaviour (dashed lines) and choosy behaviour (continuous lines).

#### S5.3 Population trait variance

**Figure S5.3:**
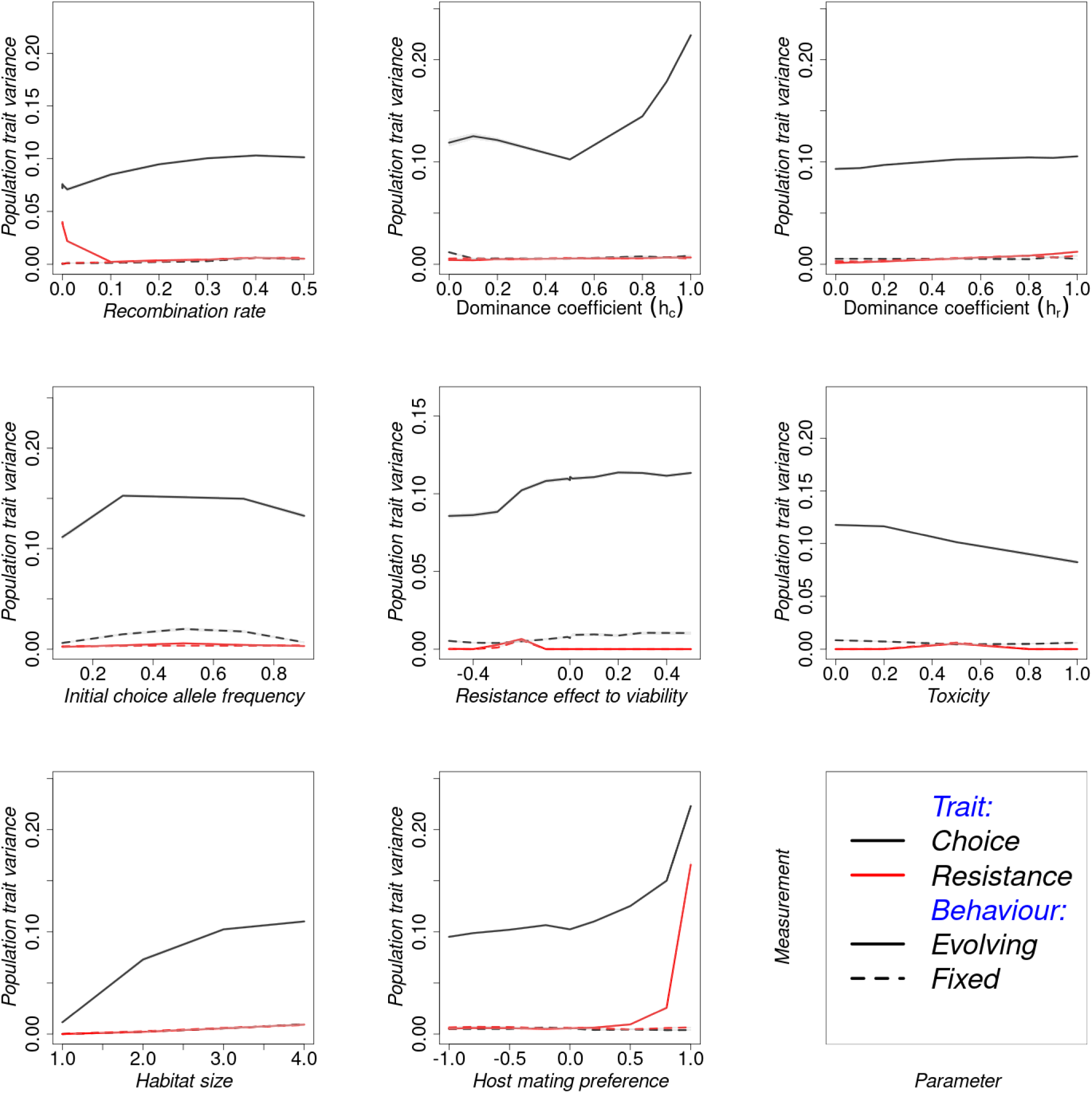
Final trait variance in large habitats, *w* = 3 but low initial choice allele frequency *p_c,_*_0_ = 0.1: Effect of model parameters on the population trait variance in resistance trait (red lines) and choice trait (black lines) between populations at the end of the simulation under random behaviour (dashed lines) and choosy behaviour (continuous lines). Plotted is the mean of trait variance across replicates.

### S6 Each species sampling

#### S6.1 Distribution of variables for each species

As described in the main text, the sampling of parameters from the species’ parameter distribution is done for each species independently. In case a given parameter was missing for a given species, a uniform distribution was used over a defined range as before. The choice index was used as the fixed choice parameter for the respective species. Prior simulations fixed this to 0.5, but with this species-level sampling, the fixed random choice parameter was also sampled.

The parameter sets used to generate the distributions in Figure S6.1 was obtained by sampling them from empirical distributions obtained by our quantitative literature survey and simulating 100 replicates of each set and each species separately. Note that species 11 is artificially constructed by combining all the species parameter distributions together. For a given variable distribution, the mean variable from the 100 replicates contributes one data point (as detailed in the methods).

From the trait distributions, the mean resistance trait evolves away from the initial resistance for most of the parameter sets under both evolving and fixed random choices. In both scenarios for the artificial species 11, more replicates fix for the susceptible allele. There were also small differences between evolving and fixed random choice, but the corresponding confidence intervals overlapped. Note that under the random choice of host plant behaviour, the choice trait in each particular population either fixed for the toxic or non-toxic host plant. Taking the averages between replicates, therefore, results in intermediate choice trait values. Populations generally lost variation in both traits. However, those under random niche choice lost more variation than those where niche choice evolved. The distribution of the rate of evolution of the resistance trait is symmetric at zero and roughly overlaps under both evolving plant choice and fixed random choice behaviours. However, the distributions of the evolution rate of choice traits are different between behaviours. A large proportion of parameter sets resulted in higher and positive rates of choice trait evolution under evolving than fixed random behaviour. The positive rates of choice evolution eventually resulted in a high choice trait for the non-toxic trait.

**Figure S6.1:**
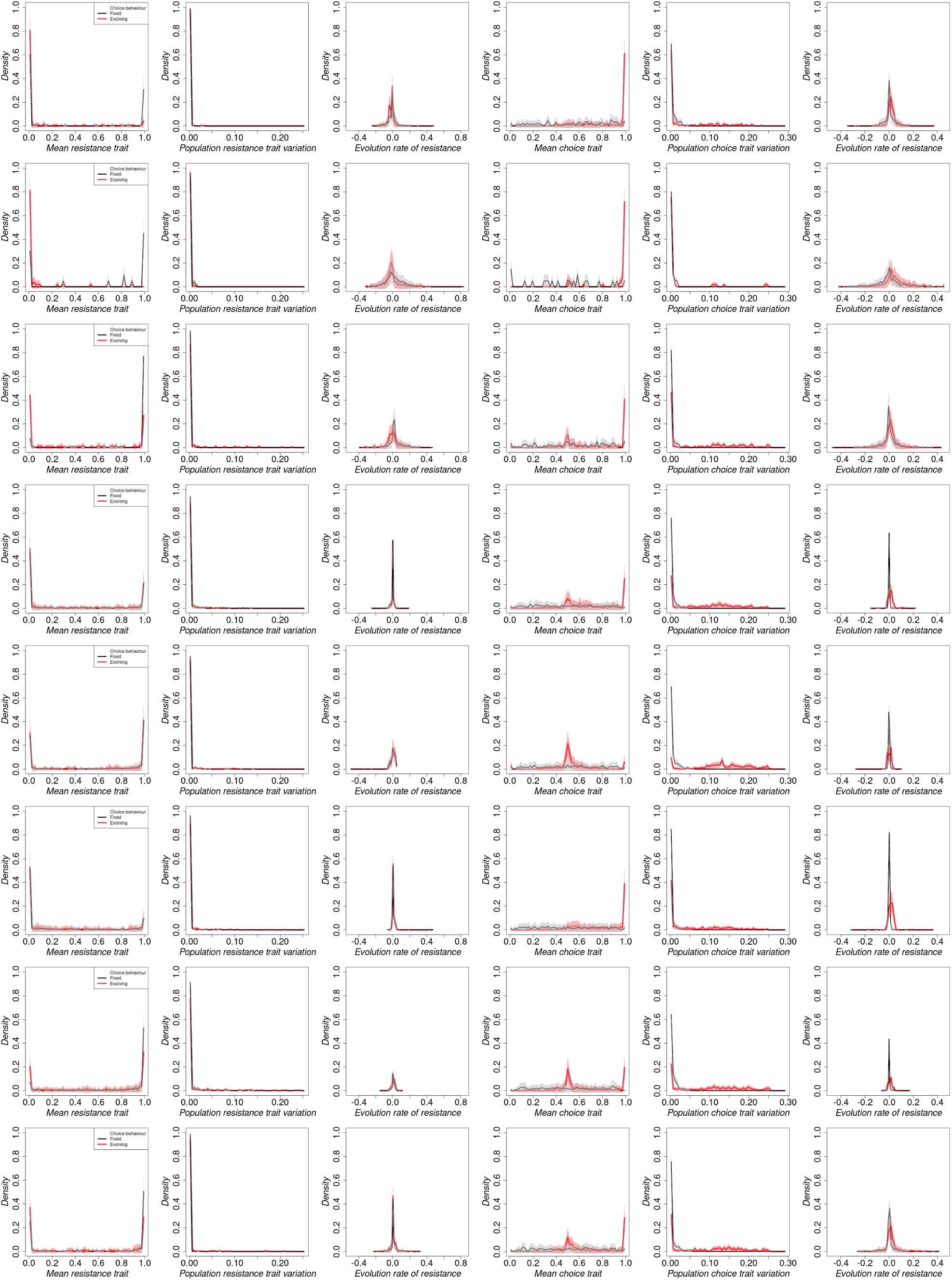
Distributions population variables at the end of 200 generations under evolving host plant choice (red lines thick) and fixed random choice (black thin lines). The mean traits were obtained as the mean of the population trait means in 100 replicate populations. The population trait variance was obtained as the mean of trait variance in 100 replicate populations. The trait evolution rate was obtained as the rate of change of mean traits from 100 replicate populations per generation. The shaded area shows the binomial confidence interval using a normal distribution approximation interval. Rows 1 to 8 correspond to species ranks 1, 2, 4, 6, 8, 9, 10, and 11, respectively. Species ranks 3, 5, and 7 are omitted because of limited data points for the trait mean and trait variation.

**Figure S6.2:**
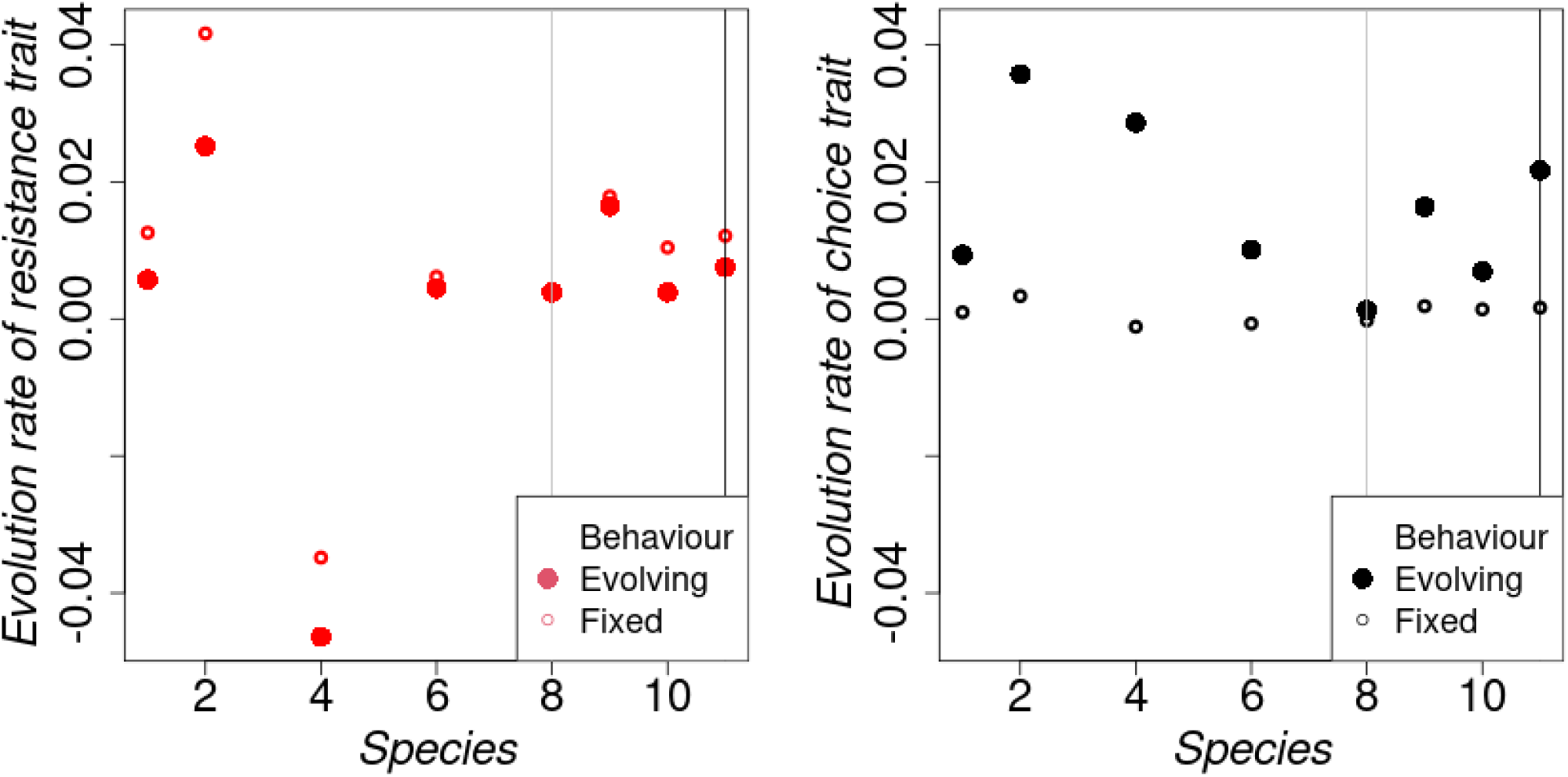
The speed of trait evolution obtained using the first 20 generations. The actual distributions from which these statistics were obtained are given in Figure S6.1. The remaining information are as given in Figure 6.

**Figure S6.3:**
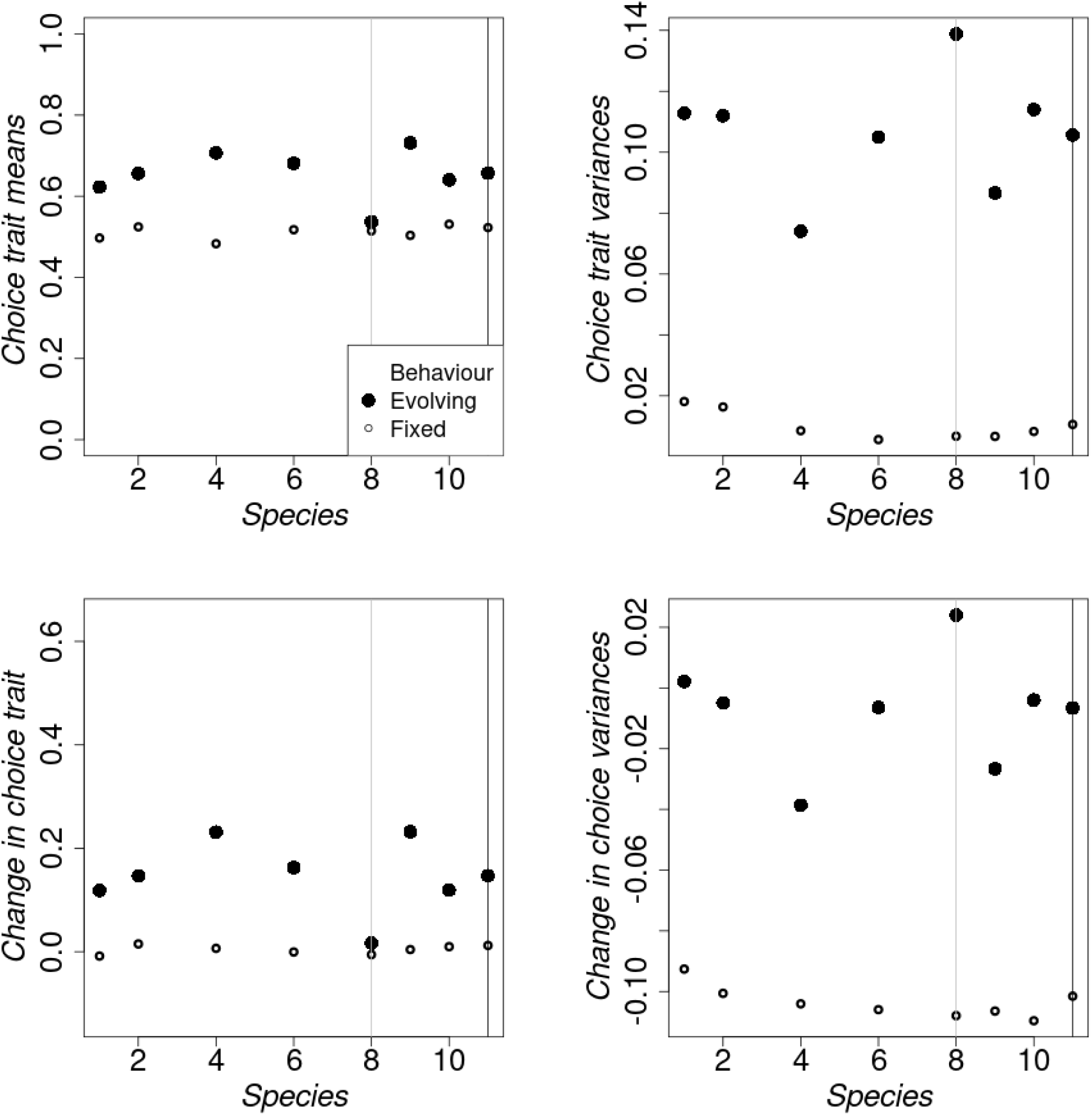
The choice trait means and trait variances obtained in generation 200 from the simulated empirical distributions. The actual distributions from which these statistics were obtained are given in Figure S6.1. The remaining information are as given in Figure 6.

### S7 Data files

Attached is the Excel file, “S7.xlsx”, with data from our empirical survey.

