## Supplementary material for "The role of evolving niche choice in herbivore adaptation to host plants": These are R scripts and data files used for generating results.: Density_Dependent_Survival_Fitting_SV.R

```
#The script does not require any other file or data.
# Script used for fitting the density-dependent survival function.

WD = "PLANTHERBIVORECHOICE"
workingdirectory = getwd()
setwd(paste0(workingdirectory,"/",WD))

density.cpb<-c(3,10, 25, 45, 150, 300); ## number of input eggs in the each niche;
Fitness<-c(1, 0.6, 0.44, 0.24, 0.006, 0); ## per capital fitness, calculated as: the number of survived adults / number of input eggs

m1<-(lm(Fitness~log(density.cpb)));
summary(m1)
m2  <- (lm(log(Fitness[-6])~density.cpb[-6]))
summary(m2)

densityvec <- 1:300

png(paste0(getwd(), "/DensityDependentSurvival.png"))
par(mar = c(5, 5, 4, 5) + 0.3)
plot(Fitness~density.cpb, pch=16, cex=2, cex.axis = 2, xlab = list("Potato beetle density", font = 3, cex = 2), ylab = list("Per capita survival fitness", font = 3, cex = 2));
#lines(seq(1:300), predict(m1, newdata=data.frame(density.cpb=seq(1:300))));
lines(densityvec, exp(m2$coefficients[1] + densityvec * m2$coefficients[2]),col="blue", lwd=5);
dev.off()

setwd(workingdirectory)
#22100000+4000000+8157000+2950000+4034000+1000000+11384500+800000
```
