## Supplementary material for "The role of evolving niche choice in herbivore adaptation to host plants": These are R scripts and data files used for generating results.: IBM_PlantHerbivore_ParameterSamplingSpeciesAnalysis_SV.R

```
#----------- New
#---------------------------
#---------------------- Empirical data distribution Simulation section ---------------
#Make sure that at the end of running this script, you run the last line of this script to reset the working directory
#Here, we upload the saved simulated data for analysis. 
#The parameter setting should be the same as in the execution and analysis scripts


WD = "PLANTHERBIVORECHOICE"
workingdirectory = getwd()
setwd(paste0(workingdirectory,"/",WD))
#----------------------------------------------
writeSuccessParametersToFile <- function(ParameterList,ts.folder,sss, reps, behaviour){
  
  outfile = paste(ts.folder,"/Parameters_successSet_choice",behaviour,".csv", sep = "")
  write.table(data.frame(t(c(unlist(ParameterList), N0rps = reps))), row.names = FALSE, col.names = sss<=1, append = sss>1, quote = FALSE, sep = ",", file = outfile)
  
}


subFolder = "DistributionSampleSimulation"
if(!dir.exists(paste0(getwd(), "/", subFolder))){
  dir.create(paste0(getwd(), "/", subFolder))
}

nnsim=5;# Number of parameter sets to be drawn. Used 500 for the manuscript
numgenerations=200; # Number of generations. Used 200 for the manuscript
nsim=5;# Number of replicates per each parameter set. Used 100 for the manuscript
targetTtimes_empirical = c(1, numgenerations)
T20 = 20# Time to measure speed
T50 = 50 # Another time to measure speed

SpRank = SETS = 1:11 # 11 species
empRresistancemean1 = empRresistancesd1 = empRresistancemean0 = empRresistancesd0 = empCchoicemean1 = empCchoicesd1 = empCchoicemean0 = empCchoicesd0 = empRresistance_popvariancemean1 = empRresistance_popvariancemean0 = empCchoice_popvariancemean1 = empCchoice_popvariancemean0 = array(NA, dim = c(length(targetTtimes_empirical), nnsim, length(SpRank)))
empnoabs_stdResSpeed1 = empnoabs_stdResSpeed0 = empnoabs_stdChoSpeed1 = empnoabs_stdChoSpeed0 = empnoabs_ResSpeed1 = empnoabs_ResSpeed0 = empnoabs_ChoSpeed1 = empnoabs_ChoSpeed0=empResSpeed1 = empResSpeed0 = empChoSpeed1 = empChoSpeed0 = array(NA, dim = c(2, nnsim, length(SpRank)))
RawPopResistance1 = RawPopResistance0 = RawPopChoice1 = RawPopChoice0 = array(NA, dim = c(length(targetTtimes_empirical), nsim, nnsim, length(SpRank)))
s0=s1=0

for(set in 1:length(SpRank)){
  #Folder = paste0("BigSimulationSpecies", set)
  Folder = paste0("BigSimulationSpecies", set)
  ParameterList = read.csv(paste0(getwd(),"/", Folder, "/Parameters_Set", set,".csv" ))[,-1]
  duplicated(ParameterList)
  ParameterList[duplicated(ParameterList), ]
  ParameterList = ParameterList[!duplicated(ParameterList), ]
  ParameterList = ParameterList[!is.na(ParameterList$SSet),]
  for(ssss in 1:nnsim){
    
    #outfilerawavoid = paste0(getwd(), "/",Folder, "/", Parameters,"_", "Avoid", "_SimPar.RData")
    # Extracting actual parameter values drawn
    
    fecunditycost = ParameterList$fc[ssss]
    scaledensity = ParameterList$hDr[ssss]
    SortMating = ParameterList$AM[ssss]
    dominanceresistance = ParameterList$DomR[ssss]
    dominancechoice = ParameterList$C[ssss]
    recombinationrate = ParameterList$rec[ssss]
    resistancecost = ParameterList$rc[ssss]
    initfreqresistance = ParameterList$fr[ssss]
    initfreqchoice = ParameterList$fch[ssss]
    toxicity = ParameterList$tox[ssss]
    N_offspring = ParameterList$lam[ssss]
    fixedChoice = ParameterList$Fxchoice[ssss]
    
    Parameters = paste0("fc", fecunditycost, "_hDr", scaledensity, "_AM",SortMating, "_DomR",dominanceresistance[1], "C", dominancechoice[1], 
                        "_rec", recombinationrate, "_rc", resistancecost, "_fr",round(initfreqresistance, 2), "fch", round(initfreqchoice, 2), 
                        "_tox", toxicity, "_lam", N_offspring, "_fxchoice", fixedChoice, "_Set",ssss )
    
    outfilerawavoid = paste0(getwd(), "/",Folder, "/", Parameters,"_", "Avoid", "_SimPar.RData")
    outfilerawnone = paste0(getwd(), "/",Folder, "/", Parameters,"_", "None", "_SimPar.RData")
    
    parameterList = list(sample = ssss, fc= fecunditycost, hDr= scaledensity, AM=SortMating, DomR=dominanceresistance[1], C = dominancechoice[1],
                         rec= recombinationrate[1], rc= resistancecost, fr=round(initfreqresistance, 2), fch= round(initfreqchoice, 2),
                         tox= toxicity, lam= N_offspring, Fxchoice = fixedChoice, SSet=set)
    
    out.choice = readRDS(outfilerawavoid)
    out.nochoice = readRDS(outfilerawnone)
    
    
    resistance_tracking<-out.choice$resistance_tracking;
    choice_tracking<-out.choice$choice_tracking;
    resistance_tracking_nochoice<-out.nochoice$resistance_tracking;
    choice_tracking_nochoice<-out.nochoice$choice_tracking;
    
    # Noting down which parameters where actually drawn.
    Rps1 = sum(!is.na(resistance_tracking[numgenerations,]))
    if(Rps1>0){
      s1=s1+1
      writeSuccessParametersToFile(ParameterList = parameterList, ts.folder = subFolder, sss=s1, reps = Rps1, behaviour = 1)
    }
    
    
    Rps0 = sum(!is.na(resistance_tracking_nochoice[numgenerations,]))
    if(Rps0>0){
      s0=s0+1
      writeSuccessParametersToFile(ParameterList = parameterList, ts.folder = subFolder, sss=s0, reps = Rps0, behaviour = 0)
    }
    
    
    
    resistance_popvar<-out.choice$resistance_popvar;
    choice_popvar<-out.choice$choice_popvar;
    resistance_popvar_nochoice<-out.nochoice$resistance_popvar;
    choice_popvar_nochoice<-out.nochoice$choice_popvar;
    
    RawPopResistance1[,,ssss,set] = resistance_tracking[targetTtimes_empirical,]
    RawPopResistance0[,,ssss,set] = resistance_tracking_nochoice[targetTtimes_empirical,]
    RawPopChoice1[,,ssss,set] = choice_tracking[targetTtimes_empirical,]
    RawPopChoice0[,,ssss,set] = choice_tracking_nochoice[targetTtimes_empirical,]
    
    resistancemean = apply(resistance_tracking[targetTtimes_empirical,],1,mean2)
    resistancesd = apply(resistance_tracking[targetTtimes_empirical,],1,FUN = function(x)sd(x, na.rm = TRUE))
    resistancemean_nochoice = apply(resistance_tracking_nochoice[targetTtimes_empirical,],1,mean2)
    resistancesd_nochoice = apply(resistance_tracking_nochoice[targetTtimes_empirical,],1,FUN = function(x)sd(x, na.rm = TRUE))
    choicemean = apply(choice_tracking[targetTtimes_empirical,],1,mean2)
    choicesd = apply(choice_tracking[targetTtimes_empirical,],1,FUN = function(x)sd(x, na.rm = TRUE))
    choicemean_nochoice = apply(choice_tracking_nochoice[targetTtimes_empirical,],1,mean2)
    choicesd_nochoice = apply(choice_tracking_nochoice[targetTtimes_empirical,],1,FUN = function(x)sd(x, na.rm = TRUE))
    
    
    resistance_popvarmean = apply(resistance_popvar[targetTtimes_empirical,],1,mean2)
    resistance_popvarmean_nochoice = apply(resistance_popvar_nochoice[targetTtimes_empirical,],1,mean2)
    choice_popvarmean = apply(choice_popvar[targetTtimes_empirical,],1,mean2)
    choice_popvarmean_nochoice = apply(choice_popvar_nochoice[targetTtimes_empirical,],1,mean2)
    
    
    empRresistancemean1[,ssss,set] = resistancemean
    empRresistancesd1[,ssss,set] = resistancesd
    empRresistancemean0[,ssss,set] = resistancemean_nochoice
    empRresistancesd0[,ssss,set] = resistancesd_nochoice
    empCchoicemean1[,ssss,set] = choicemean
    empCchoicesd1[,ssss,set] = choicesd
    empCchoicemean0[,ssss,set] = choicemean_nochoice
    empCchoicesd0[,ssss,set] = choicesd_nochoice
    
    # #----pop sd stored
    empRresistance_popvariancemean1[,ssss,set] = resistance_popvarmean
    empRresistance_popvariancemean0[,ssss,set] = resistance_popvarmean_nochoice
    empCchoice_popvariancemean1[,ssss,set] = choice_popvarmean
    empCchoice_popvariancemean0[,ssss,set] = choice_popvarmean_nochoice
    
    
    ##------------ Evolution rate ------------
    resT11 = rowMeans(resistance_tracking[c(1,T20, T50),], na.rm = TRUE)
    resT10 = rowMeans(resistance_tracking_nochoice[c(1,T20, T50),], na.rm = TRUE)
    choT11 = rowMeans(choice_tracking[c(1,T20, T50),], na.rm = TRUE)
    choT10 = rowMeans(choice_tracking_nochoice[c(1,T20, T50),], na.rm = TRUE)
    
    empResSpeed1[,ssss,set] = c(abs(resT11[2]-resT11[1])/(T20-1), abs(resT11[3]-resT11[1])/(T50-1))
    empResSpeed0[,ssss,set] = c(abs(resT10[2]-resT10[1])/(T20-1), abs(resT10[3]-resT10[1])/(T50-1))
    empChoSpeed1[,ssss,set] = c(abs(choT11[2]-choT11[1])/(T20-1), abs(choT11[3]-choT11[1])/(T50-1))
    empChoSpeed0[,ssss,set] = c(abs(choT10[2]-choT10[1])/(T20-1), abs(choT10[3]-choT10[1])/(T50-1))
    
    
    resT11 = diff(rowMeans(resistance_tracking[c(1:T50),], na.rm = TRUE))
    resT10 = diff(rowMeans(resistance_tracking_nochoice[c(1:T50),], na.rm = TRUE))
    choT11 = diff(rowMeans(choice_tracking[c(1:T50),], na.rm = TRUE))
    choT10 = diff(rowMeans(choice_tracking_nochoice[c(1:T50),], na.rm = TRUE))
    
    stdresT11 = apply(diff(resistance_tracking[c(1:T50),]),1,FUN = function(x)std_error(x))
    stdresT10 = apply(diff(resistance_tracking_nochoice[c(1:T50),]),1,FUN = function(x)std_error(x))
    stdchoT11 = apply(diff(choice_tracking[c(1:T50),]),1,FUN = function(x)std_error(x))
    stdchoT10 = apply(diff(choice_tracking_nochoice[c(1:T50),]),1,FUN = function(x)std_error(x))
    
    empResSpeed1[,ssss,set] = abs(c(mean(resT11[1:T20], na.rm = TRUE), mean(resT11[1:T50], na.rm = TRUE)))
    empResSpeed0[,ssss,set] = abs(c(mean(resT10[1:T20], na.rm = TRUE), mean(resT10[1:T50], na.rm = TRUE)))
    empChoSpeed1[,ssss,set] = abs(c(mean(choT11[1:T20], na.rm = TRUE), mean(choT11[1:T50], na.rm = TRUE)))
    empChoSpeed0[,ssss,set] = abs(c(mean(choT10[1:T20], na.rm = TRUE), mean(choT10[1:T50], na.rm = TRUE)))
    
    empnoabs_ResSpeed1[,ssss,set] = c(mean(resT11[1:T20], na.rm = TRUE), mean(resT11[1:T50], na.rm = TRUE))
    empnoabs_ResSpeed0[,ssss,set] = c(mean(resT10[1:T20], na.rm = TRUE), mean(resT10[1:T50], na.rm = TRUE))
    empnoabs_ChoSpeed1[,ssss,set] = c(mean(choT11[1:T20], na.rm = TRUE), mean(choT11[1:T50], na.rm = TRUE))
    empnoabs_ChoSpeed0[,ssss,set] = c(mean(choT10[1:T20], na.rm = TRUE), mean(choT10[1:T50], na.rm = TRUE))
    
    empnoabs_stdResSpeed1[,ssss,set] = c(mean(stdresT11[1:T20], na.rm = TRUE), mean(stdresT11[1:T50], na.rm = TRUE))
    empnoabs_stdResSpeed0[,ssss,set] = c(mean(stdresT10[1:T20], na.rm = TRUE), mean(stdresT10[1:T50], na.rm = TRUE))
    empnoabs_stdChoSpeed1[,ssss,set] = c(mean(stdchoT11[1:T20], na.rm = TRUE), mean(stdchoT11[1:T50], na.rm = TRUE))
    empnoabs_stdChoSpeed0[,ssss,set] = c(mean(stdchoT10[1:T20], na.rm = TRUE), mean(stdchoT10[1:T50], na.rm = TRUE))
   # print(c("set ", set, "ssss ", ssss))
  }
}

# Save analysed data such that you do not rerun everytime you want to visualize variables. If you have already analysed and saved, skip this step including the above loops and just go to reloading the analysed saved data

saveRDS(RawPopResistance1, file=paste0(getwd(),"/", subFolder,  "/", "RawPopResistance1.RData"))
saveRDS(RawPopResistance0, file=paste0(getwd(),"/", subFolder,  "/", "RawPopResistance0.RData"))
saveRDS(RawPopChoice1, file=paste0(getwd(),"/", subFolder,"/", "RawPopChoice1.RData"))
saveRDS(RawPopChoice0, file=paste0(getwd(),"/", subFolder,"/", "RawPopChoice0.RData"))

saveRDS(empRresistancemean1, file=paste0(getwd(),"/", subFolder,"/", "empRresistancemean1.RData"))
saveRDS(empRresistancesd1, file=paste0(getwd(),"/", subFolder,"/","empRresistancesd1.RData"))
saveRDS(empCchoicemean1, file=paste0(getwd(),"/", subFolder,"/","empCchoicemean1.RData"))
saveRDS(empCchoicesd1, file=paste0(getwd(),"/", subFolder,"/","empCchoicesd1.RData"))
saveRDS(empRresistancemean0, file=paste0(getwd(),"/", subFolder,"/", "empRresistancemean0.RData"))
saveRDS(empRresistancesd0, file=paste0(getwd(),"/", subFolder,"/","empRresistancesd0.RData"))
saveRDS(empCchoicemean0, file=paste0(getwd(),"/", subFolder,"/","empCchoicemean0.RData"))
saveRDS(empCchoicesd0, file=paste0(getwd(),"/", subFolder,"/","empCchoicesd0.RData"))

saveRDS(empRresistance_popvariancemean1, file=paste0(getwd(),"/", subFolder,"/", "empRresistance_popvariancemean1.RData"))
saveRDS(empCchoice_popvariancemean1, file=paste0(getwd(),"/", subFolder,"/","empCchoice_popvariancemean1.RData"))
saveRDS(empRresistance_popvariancemean0, file=paste0(getwd(),"/", subFolder,"/", "empRresistance_popvariancemean0.RData"))
saveRDS(empCchoice_popvariancemean0, file=paste0(getwd(),"/", subFolder,"/","empCchoice_popvariancemean0.RData"))

saveRDS(empResSpeed0, file=paste0(getwd(),"/", subFolder,"/", "empResSpeed0.RData"))
saveRDS(empResSpeed1, file=paste0(getwd(),"/", subFolder,"/","empResSpeed1.RData"))
saveRDS(empChoSpeed0, file=paste0(getwd(),"/", subFolder,"/","empChoSpeed0.RData"))
saveRDS(empChoSpeed1, file=paste0(getwd(),"/", subFolder,"/","empChoSpeed1.RData"))


saveRDS(empnoabs_ResSpeed0, file=paste0(getwd(),"/", subFolder,"/", "empnoabs_ResSpeed0.RData"))
saveRDS(empnoabs_ResSpeed1, file=paste0(getwd(),"/", subFolder,"/","empnoabs_ResSpeed1.RData"))
saveRDS(empnoabs_ChoSpeed0, file=paste0(getwd(),"/", subFolder,"/","empnoabs_ChoSpeed0.RData"))
saveRDS(empnoabs_ChoSpeed1, file=paste0(getwd(),"/", subFolder,"/","empnoabs_ChoSpeed1.RData"))

saveRDS(empnoabs_stdResSpeed0, file=paste0(getwd(),"/", subFolder,"/", "empnoabs_stdResSpeed0.RData"))
saveRDS(empnoabs_stdResSpeed1, file=paste0(getwd(),"/", subFolder,"/","empnoabs_stdResSpeed1.RData"))
saveRDS(empnoabs_stdChoSpeed0, file=paste0(getwd(),"/", subFolder,"/","empnoabs_stdChoSpeed0.RData"))
saveRDS(empnoabs_stdChoSpeed1, file=paste0(getwd(),"/", subFolder,"/","empnoabs_stdChoSpeed1.RData"))

setwd(workingdirectory)
```
