## Supplementary material for "The role of evolving niche choice in herbivore adaptation to host plants": These are R scripts and data files used for generating results.: IBM_PlantHerbivore_ParameterSamplingSpeciesExecute_SV.R

```
#Make sure that at the end of running this script, you run the last line of this script to reset the working directory
#Here, we execute the main IBM script for parameter distribution sampling simulations. 
#The parameter setting should be the same as in the execution and analysis scripts
# Requires parameter distributions for species 1 to 11 be stored in a folder named "EmpiricalData" in the current working directory

rm(list=ls()) 
library(dplyr)
library(scales)
library(parallel)
cores = detectCores()-1
WD = "PLANTHERBIVORECHOICE"
workingdirectory = getwd()
setwd(paste0(workingdirectory,"/",WD))

source("IBM_PlantHerbivore_SV.R")

#------ Default parameter values --------------------

nnsim=5;# Number of parameter sets for each species. Used 500 for the manuscript
numgenerations=200; # Number of generations. Used 200 for simulations
nsim=5;# Number of replicates per each parameter set. Used 100 for the manuscript
numresistanceloci  = 1; #number of loci underlying resistance to plant toxin
numchoiceloci  = 1; #number of loci underlying host plant choice
## Rescaling is done within the function to have the sum of effect sizes sum to 1. 
resistanceeffectsizes = rep(0.5, numresistanceloci); #vector of effect sizes; length should correspond to the number of resistance loci; Converted within the function such that the effect sizes add up to 1
choiceeffectsizes = rep(0.5, numchoiceloci);#vector of effect sizes; length should correspond to the number of choice loci; Converted within the function such that the effect sizes add up to 1
K = 1000
N0 = 20
environmentalsdresistance = 0.0
environmentalsdchoice = 0.0


#-----------------Sampled parameters--------------------------------
for(sprank in 1:11){
  set = sprank
  Folder = paste0("BigSimulationSpecies", set)
  
  if(!dir.exists(paste0(getwd(), "/",Folder))){
    dir.create(paste0(getwd(), "/",Folder))
  }
  
  
  ffixedChoice = read.csv(file = paste0(getwd(), "/EmpiricalData/ChoiceIndex", "_SpRank_", sprank, ".csv"), header = TRUE)[,2]
  if(length(ffixedChoice)<1){
    ffixedChoice = seq(0,1,0.1)
  }
  
  if(!exists("ffixedChoice")){
    ffixedChoice = seq(0,1,0.1)
  }else if(length(ffixedChoice)<1){
    ffixedChoice = seq(0,1,0.1)
  }
  
  NN_offsfring = read.csv(file = paste0(getwd(), "/EmpiricalData/Lambda", "_SpRank_", sprank, ".csv"), header = TRUE)[,2]
  if(length(NN_offsfring)<1){
    NN_offsfring = seq(1,20,1)
  }
  ssortMating = read.csv(file = paste0(getwd(), "/EmpiricalData/AssortativeMating", "_SpRank_", sprank, ".csv"), header = TRUE)[,2]
  if(length(ssortMating)<1){
    ssortMating = c(-1,-0.8,-0.5,-0.2,0,0.2,0.5,0.8,1)
  }
  
  FFecunditycost = read.csv(file = paste0(getwd(), "/EmpiricalData/FecundityCost", "_SpRank_", sprank, ".csv"), header = TRUE)[,2]
  if(length(FFecunditycost)<1){
    FFecunditycost = c(-1, -0.5, 0, 0.5, 1)
  }
  
  RResistancecost = read.csv(file = paste0(getwd(), "/EmpiricalData/ViabilityCost", "_SpRank_", sprank, ".csv"), header = TRUE)[,2]
  if(length(RResistancecost)<1){
    RResistancecost = c(0.001, 0.01, 0.1, 0.2, 0.3)
  }
  TToxicity = read.csv(file = paste0(getwd(), "/EmpiricalData/Toxicity2", "_SpRank_", sprank, ".csv"), header = TRUE)[,2]
  if(length(TToxicity)<1){
    TToxicity = c(0.0, 0.2, 0.5, 0.8, 1 )
  }
  FFr0 = read.csv(file = paste0(getwd(), "/EmpiricalData/Fr0", "_SpRank_", sprank, ".csv"), header = TRUE)[,2]
  if(length(FFr0)<1){
    FFr0 = c(0.1,0.3, 0.5,0.7, 0.9)
  }
  FFc0 = seq(0.1,0.9, length.out=9)
  DDominanceresistance = seq(0,1,length.out=11)
  DDominancechoice = seq(0,1,length.out=11)
  SScaledensity = c(1, 2, 3, 4, 5)
  RRecombinationrate = c(0.1, 0.2, 0.3, 0.4, 0.5)
  #fixedChoice = 0.5 # random choice behaviour
  for(ssss in 1:nnsim){
    fixedChoice = sample(x=ffixedChoice, size = 1) # random choice behaviour
    N_offspring = sample(x=NN_offsfring, size = 1)
    Recombinationrate = sample(x=RRecombinationrate, size = 1)
    Dominancechoice = sample(x=DDominancechoice, size = 1)
    Dominanceresistance = sample(x=DDominanceresistance, size = 1)
    sortMating = sample(x=ssortMating, size = 1)
    Fecunditycost = sample(x=FFecunditycost, size = 1)
    Resistancecost = sample(x=RResistancecost, size = 1)
    Toxicity = sample(x=TToxicity, size=1)
    Fr0 = sample(x=FFr0, size = 1)
    Fc0 = sample(x=FFc0, size = 1)
    Scaledensity = sample(x=SScaledensity, size = 1)
    hr = Dominanceresistance
    hc = Dominancechoice
    dominanceresistance = rep(hr, numresistanceloci);#dominance values for the resistance loci, should generally be between 0 and 1
    dominancechoice = rep(hc, numchoiceloci);#dominance values for the choice loci, should generally be between 0 an 1
    scaledensity = Scaledensity
    SortMating = sortMating
    toxicity = c(0,Toxicity)
    resistancecost = Resistancecost
    fecunditycost = Fecunditycost
    initfreqchoice = Fc0
    initfreqresistance = Fr0
    recrate = Recombinationrate
    recombinationrate = rep(recrate, numresistanceloci + numchoiceloci)
    
    out.nochoice<-Beetle_Bigsimulation_fcb(ts.folder = Folder, numgenerations=numgenerations, nsim=nsim, choice="None", recombinationrate = recombinationrate, 
                                           numresistanceloci  = numresistanceloci, numchoiceloci = numchoiceloci, resistanceeffectsizes = resistanceeffectsizes, 
                                           choiceeffectsizes = choiceeffectsizes, dominanceresistance = dominanceresistance, dominancechoice = dominancechoice, 
                                           scaledensity = scaledensity, K  = K, toxicity=toxicity, resistancecost=resistancecost, initfreqchoice = initfreqchoice, 
                                           initfreqresistance = initfreqresistance,SortMating=SortMating,N_offspring=N_offspring, N0=N0,
                                           environmentalsdresistance = environmentalsdresistance, environmentalsdchoice = environmentalsdchoice,
                                           fecunditycost = fecunditycost, Set = ssss, FixedChoice = fixedChoice)
    
    out.avoid<-Beetle_Bigsimulation_fcb(ts.folder = Folder, numgenerations=numgenerations, nsim=nsim, choice="Avoid", recombinationrate = recombinationrate, 
                                        numresistanceloci  = numresistanceloci, numchoiceloci = numchoiceloci, resistanceeffectsizes = resistanceeffectsizes, 
                                        choiceeffectsizes = choiceeffectsizes, dominanceresistance = dominanceresistance, dominancechoice = dominancechoice, 
                                        scaledensity = scaledensity, K  = K, toxicity=toxicity, resistancecost=resistancecost, initfreqchoice = initfreqchoice, 
                                        initfreqresistance = initfreqresistance,SortMating=SortMating,N_offspring=N_offspring, N0=N0,
                                        environmentalsdresistance = environmentalsdresistance, environmentalsdchoice = environmentalsdchoice,
                                        fecunditycost = fecunditycost, Set = ssss, FixedChoice = fixedChoice)
    
    
    
    parameterList = list(sample = ssss, fc= fecunditycost, hDr= scaledensity, AM=SortMating, DomR=dominanceresistance[1], C = dominancechoice[1], 
                         rec= recombinationrate[1], rc= resistancecost, fr=round(initfreqresistance, 2), fch= round(initfreqchoice, 2), 
                         tox= toxicity[2], lam= N_offspring, Fxchoice = fixedChoice, SSet=set)
    
    
    writeParametersToFile(ParameterList = parameterList, ts.folder = Folder, sss=ssss)
   # print(c("sss ", ssss))
  }
}
# 

setwd(workingdirectory)
```
