## Supplementary material for "The role of evolving niche choice in herbivore adaptation to host plants": These are R scripts and data files used for generating results.: IBM_PlantHerbivore_ParameterSamplingSpeciesPlotting_SV.R

```
#Make sure that at the end of running this script, you run the last line of this script to reset the working directory
#Make sure that at the end of running this script, you run the last line of this script to reset the working directory
#Here, we upload the saved analysed data for plotting. 
#The parameter setting should be the same as in the execution and analysis scripts
library(binom)
library(plotrix)

WD = "PLANTHERBIVORECHOICE"
workingdirectory = getwd()
setwd(paste0(workingdirectory,"/",WD))

subFolder = "DistributionSampleSimulation"
plotFolder = "Plots"
if(!dir.exists(paste0(getwd(), "/", subFolder, "/", plotFolder))){
  dir.create(paste0(getwd(), "/", subFolder, "/", plotFolder))
}

#--------- Generations at which speed is measured -------
T20 = 20#
T50 = 50
tTspeed = c(T20, T50)
numgenerations = 200
targetTtimes_empirical = c(1, numgenerations)
#----------- Reloading saved data for plotting. 
empnoabs_stdResSpeed0 = readRDS(paste0(getwd(),"/", subFolder,"/", "empnoabs_stdResSpeed0.RData"))
empnoabs_stdResSpeed1 = readRDS(paste0(getwd(),"/", subFolder,"/","empnoabs_stdResSpeed1.RData"))
empnoabs_stdChoSpeed0 = readRDS(paste0(getwd(),"/", subFolder,"/","empnoabs_stdChoSpeed0.RData"))
empnoabs_stdChoSpeed1 = readRDS(paste0(getwd(),"/", subFolder,"/","empnoabs_stdChoSpeed1.RData"))

empnoabs_ResSpeed0 = readRDS(paste0(getwd(),"/", subFolder,"/", "empnoabs_ResSpeed0.RData"))
empnoabs_ResSpeed1 = readRDS(paste0(getwd(),"/", subFolder,"/","empnoabs_ResSpeed1.RData"))
empnoabs_ChoSpeed0 = readRDS(paste0(getwd(),"/", subFolder,"/","empnoabs_ChoSpeed0.RData"))
empnoabs_ChoSpeed1 = readRDS(paste0(getwd(),"/", subFolder,"/","empnoabs_ChoSpeed1.RData"))

empRresistancemean1= readRDS(paste0(getwd(),"/", subFolder,"/", "empRresistancemean1.RData"))
empRresistancesd1 = readRDS(paste0(getwd(),"/", subFolder,"/","empRresistancesd1.RData"))
empCchoicemean1= readRDS(paste0(getwd(),"/", subFolder,"/","empCchoicemean1.RData"))
empCchoicesd1= readRDS(paste0(getwd(),"/", subFolder,"/","empCchoicesd1.RData"))
empRresistancemean0 = readRDS(paste0(getwd(),"/", subFolder,"/", "empRresistancemean0.RData"))
empRresistancesd0 = readRDS(paste0(getwd(),"/", subFolder,"/","empRresistancesd0.RData"))
empCchoicemean0= readRDS(paste0(getwd(),"/", subFolder,"/","empCchoicemean0.RData"))
empCchoicesd0= readRDS(paste0(getwd(),"/", subFolder,"/","empCchoicesd0.RData"))

empRresistance_popvariancemean1= readRDS(paste0(getwd(),"/", subFolder,"/", "empRresistance_popvariancemean1.RData"))
empCchoice_popvariancemean1= readRDS(paste0(getwd(),"/", subFolder,"/","empCchoice_popvariancemean1.RData"))
empRresistance_popvariancemean0 = readRDS(paste0(getwd(),"/", subFolder,"/", "empRresistance_popvariancemean0.RData"))
empCchoice_popvariancemean0= readRDS(paste0(getwd(),"/", subFolder,"/","empCchoice_popvariancemean0.RData"))


RawPopResistance1= readRDS(paste0(getwd(),"/", subFolder, "/","RawPopResistance1.RData"))
RawPopResistance0= readRDS(paste0(getwd(),"/", subFolder,"/","RawPopResistance0.RData"))
RawPopChoice1= readRDS(paste0(getwd(),"/", subFolder, "/","RawPopChoice1.RData"))
RawPopChoice0= readRDS(paste0(getwd(),"/", subFolder,"/","RawPopChoice0.RData"))

empResSpeed0= readRDS(paste0(getwd(),"/", subFolder,"/", "empResSpeed0.RData"))
empResSpeed1= readRDS(paste0(getwd(),"/", subFolder,"/","empResSpeed1.RData"))
empChoSpeed0= readRDS(paste0(getwd(),"/", subFolder,"/","empChoSpeed0.RData"))
empChoSpeed1= readRDS(paste0(getwd(),"/", subFolder,"/","empChoSpeed1.RData"))

SuccessParameters0= read.csv(paste0(getwd(),"/", subFolder,"/","Parameters_successSet_choice0.csv"))
SuccessParameters1= read.csv(paste0(getwd(),"/", subFolder,"/","Parameters_successSet_choice1.csv"))


# #------------- Successful parameter distributions ------------

#--------------------- Final measurements ----------------------


#------------------ Option with speed on a uniform x-axis for the different species -----------

plotFolder = "Plots"
if(!dir.exists(paste0(getwd(), "/", subFolder, "/", plotFolder))){
  dir.create(paste0(getwd(), "/", subFolder, "/", plotFolder))
}

# #------------- Successful parameter distributions ------------

#--------------------- Final measurements ----------------------

#------------- Speed of adaptation --------------

binsize = 60
genT = length(targetTtimes_empirical)

#--------------------- Binomial confidence interval --------------------


BinomialCI_normal = function(X,n,z=1.96){
  CI = matrix(NA, nrow = length(X), ncol = 2)
  Phat = X/n
  CI[,1] = Phat-z*sqrt(Phat*(1-Phat)/n)
  CI[,2] = Phat+z*sqrt(Phat*(1-Phat)/n)
  
  return(CI)
}

BinomialCI_Wilson = function(X,n,z){
  CI = matrix(NA, nrow = length(X), ncol = 2)
  Phat = X/n
  CI[,1] = (Phat+(z^2)/(2*n))/(1+(z^2)/n) - (z/(1+(z^2)/n))*sqrt(Phat*(1-Phat)/n + (z^2)/(4*n^2))
  CI[,2] = (Phat+(z^2)/(2*n))/(1+(z^2)/n) + (z/(1+(z^2)/n))*sqrt(Phat*(1-Phat)/n + (z^2)/(4*n^2))
  return(CI)
}     

zz=1.96

#--------------------- Final measurements ----------------------
for(sprank in 1:11){
  
  MinRes = min(c(RawPopResistance1, RawPopResistance0), na.rm = TRUE)
  MaxRes = max(c(RawPopResistance1, RawPopResistance0), na.rm = TRUE)
  if(sum(!is.na(RawPopResistance1[genT,,,sprank]))>0 & sum(!is.na(RawPopResistance0[genT,,,sprank]))>0){
    
    #------------ Mean traits --------------------
    
    MinRes = min(c(empRresistancemean1, empRresistancemean0), na.rm = TRUE)
    MaxRes = max(c(empRresistancemean1, empRresistancemean0), na.rm = TRUE)
    
    
    h1 = try(hist(empRresistancemean1[genT,,sprank], breaks = seq(MinRes, MaxRes, length.out = binsize), plot = FALSE))
    h0 = try(hist(empRresistancemean0[genT,,sprank], breaks = seq(MinRes, MaxRes, length.out = binsize), plot = FALSE))
    
    BN1 = BinomialCI_normal(X=h1$density,n=sum(h1$density),z=zz)
    BN0 = BinomialCI_normal(X=h0$density,n=sum(h0$density),z=zz)
    
    BW1 = BinomialCI_Wilson(X=h1$density,n=sum(h1$density),z=zz)
    BW0 = BinomialCI_Wilson(X=h0$density,n=sum(h0$density),z=zz)
    
    png(paste0(getwd(),"/", subFolder, "/", plotFolder,"/meanResistanceTrait_Both_BCI_Normal", "binsize_", binsize, "Species", sprank, "_gen",targetTtimes_empirical[genT], ".png"))
    par(mar = c(5, 5, 4, 5) + 0.3)
    try(plot(h1$mids,h1$density/sum(h1$density), ty='l', lwd = 5, cex.axis = 2, ylim = c(min(c(0)), max(c(1))), col = 2, lty = 1,
             xlab = list("Mean resistance trait", font = 3, cex = 2), ylab = list("Density", font = 3, cex = 2), main = ""))
    lines(h0$mids, h0$density/sum(h0$density), ty='l', lwd = 3, col = 1, lty = 1)
    polygon(c(h1$mids,rev(h1$mids)), c(BN1[,1], rev(BN1[,2])),  border=NA, col=adjustcolor("red", alpha.f = 0.3))
    polygon(c(h0$mids,rev(h0$mids)), c(BN0[,1], rev(BN0[,2])),  border=NA, col=adjustcolor("gray", alpha.f = 0.5))
    
    legend("topright", legend = list("Choice behaviour","Fixed", "Evolving"), col = c(NA,1,2), lwd = c(NA,3,5), lty = 1)
    dev.off()
    
    # png(paste0(getwd(),"/", subFolder, "/", plotFolder,"/meanResistanceTrait_Both_BCI_Wilson", "binsize_", binsize, "Species", sprank, "_gen",targetTtimes_empirical[genT], ".png"))
    # par(mar = c(5, 5, 4, 5) + 0.3)
    # try(plot(h1$mids,h1$density/sum(h1$density), ty='l', lwd = 5, cex.axis = 2, ylim = c(min(c(0)), max(c(1))), col = 2, lty = 1,
    #          xlab = list("Mean resistance trait", font = 3, cex = 2), ylab = list("Density", font = 3, cex = 2), main = ""))
    # lines(h0$mids, h0$density/sum(h0$density), ty='l', lwd = 3, col = 1, lty = 1)
    # polygon(c(h1$mids,rev(h1$mids)), c(BW1[,1], rev(BW1[,2])),  border=NA, col=adjustcolor("red", alpha.f = 0.3))
    # polygon(c(h0$mids,rev(h0$mids)), c(BW0[,1], rev(BW0[,2])),  border=NA, col=adjustcolor("gray", alpha.f = 0.5))
    # 
    # legend("topright", legend = list("Choice behaviour","Fixed", "Evolving"), col = c(NA,1,2), lwd = c(NA,3,5), lty = 1)
    # dev.off()
    
    
    MinCho = min(c(empCchoicemean1, empCchoicemean0), na.rm = TRUE)
    MaxCho = max(c(empCchoicemean1, empCchoicemean0), na.rm = TRUE)
    
    h1 = try(hist(empCchoicemean1[genT,,sprank], breaks = seq(MinCho, MaxCho, length.out = binsize), plot = FALSE))
    h0 = try(hist(empCchoicemean0[genT,,sprank], breaks = seq(MinCho, MaxCho, length.out = binsize), plot = FALSE))
    
    
    BN1 = BinomialCI_normal(X=h1$density,n=sum(h1$density),z=zz)
    BN0 = BinomialCI_normal(X=h0$density,n=sum(h0$density),z=zz)
    
    BW1 = BinomialCI_Wilson(X=h1$density,n=sum(h1$density),z=zz)
    BW0 = BinomialCI_Wilson(X=h0$density,n=sum(h0$density),z=zz)
    
    png(paste0(getwd(),"/", subFolder, "/", plotFolder,"/meanChoiceTrait_Both_BCI_Normal", "binsize_", binsize, "Species", sprank, "_gen",targetTtimes_empirical[genT], ".png"))
    par(mar = c(5, 5, 4, 5) + 0.3)
    try(plot(h1$mids,h1$density/sum(h1$density), ty='l', lwd = 5, cex.axis = 2, ylim = c(min(c(0)), max(c(1))), col = 2, lty = 1,
             xlab = list("Mean choice trait", font = 3, cex = 2), ylab = list("Density", font = 3, cex = 2), main = ""))
    lines(h0$mids, h0$density/sum(h0$density), ty='l', lwd = 3, col = 1, lty = 1)
    polygon(c(h1$mids,rev(h1$mids)), c(BN1[,1], rev(BN1[,2])),  border=NA, col=adjustcolor("red", alpha.f = 0.3))
    polygon(c(h0$mids,rev(h0$mids)), c(BN0[,1], rev(BN0[,2])),  border=NA, col=adjustcolor("gray", alpha.f = 0.5))
    
    #legend("topright", legend = list("Choice behaviour","Fixed", "Evolving"), col = c(NA,1,2), lwd = c(NA,3,5), lty = 1)
    dev.off()
    
    # png(paste0(getwd(),"/", subFolder, "/", plotFolder,"/meanChoiceTrait_Both_BCI_Wilson", "binsize_", binsize, "Species", sprank, "_gen",targetTtimes_empirical[genT], ".png"))
    # par(mar = c(5, 5, 4, 5) + 0.3)
    # try(plot(h1$mids,h1$density/sum(h1$density), ty='l', lwd = 5, cex.axis = 2, ylim = c(min(c(0)), max(c(1))), col = 2, lty = 1,
    #          xlab = list("Mean choice trait", font = 3, cex = 2), ylab = list("Density", font = 3, cex = 2), main = ""))
    # lines(h0$mids, h0$density/sum(h0$density), ty='l', lwd = 3, col = 1, lty = 1)
    # polygon(c(h1$mids,rev(h1$mids)), c(BW1[,1], rev(BW1[,2])),  border=NA, col=adjustcolor("red", alpha.f = 0.3))
    # polygon(c(h0$mids,rev(h0$mids)), c(BW0[,1], rev(BW0[,2])),  border=NA, col=adjustcolor("gray", alpha.f = 0.5))
    # 
    # #legend("topright", legend = list("Choice behaviour","Fixed", "Evolving"), col = c(NA,1,2), lwd = c(NA,3,5), lty = 1)
    # dev.off()
    
    
    #------------- sd within populations --------------
    
    MinRes = min(c(empRresistance_popvariancemean1, empRresistance_popvariancemean0), na.rm = TRUE)
    MaxRes = max(c(empRresistance_popvariancemean1, empRresistance_popvariancemean0), na.rm = TRUE)
    
    
    h1 = try(hist(empRresistance_popvariancemean1[genT,,sprank], breaks = seq(MinRes, MaxRes, length.out = binsize), plot = FALSE))
    h0 = try(hist(empRresistance_popvariancemean0[genT,,sprank], breaks = seq(MinRes, MaxRes, length.out = binsize), plot = FALSE))
    
    
    BN1 = BinomialCI_normal(X=h1$density,n=sum(h1$density),z=zz)
    BN0 = BinomialCI_normal(X=h0$density,n=sum(h0$density),z=zz)
    
    BW1 = BinomialCI_Wilson(X=h1$density,n=sum(h1$density),z=zz)
    BW0 = BinomialCI_Wilson(X=h0$density,n=sum(h0$density),z=zz)
    
    png(paste0(getwd(),"/", subFolder, "/", plotFolder,"/sdpopResistanceTrait_Both_BCI_Normal", "binsize_", binsize, "Species", sprank, "_gen",targetTtimes_empirical[genT], ".png"))
    par(mar = c(5, 5, 4, 5) + 0.3)
    try(plot(h1$mids,h1$density/sum(h1$density), ty='l', lwd = 5, cex.axis = 2, ylim = c(min(c(0)), max(c(1))), col = 2, lty = 1,
             xlab = list("Population resistance trait variation", font = 3, cex = 2), ylab = list("Density", font = 3, cex = 2), main = ""))
    lines(h0$mids, h0$density/sum(h0$density), ty='l', lwd = 3, col = 1, lty = 1)
    polygon(c(h1$mids,rev(h1$mids)), c(BN1[,1], rev(BN1[,2])),  border=NA, col=adjustcolor("red", alpha.f = 0.3))
    polygon(c(h0$mids,rev(h0$mids)), c(BN0[,1], rev(BN0[,2])),  border=NA, col=adjustcolor("gray", alpha.f = 0.5))
    
    #legend("topright", legend = list("Choice behaviour","Fixed", "Evolving"), col = c(NA,1,2), lwd = c(NA,3,5), lty = 1)
    dev.off()
    
    # png(paste0(getwd(),"/", subFolder, "/", plotFolder,"/sdpopResistanceTrait_Both_BCI_Wilson", "binsize_", binsize, "Species", sprank, "_gen",targetTtimes_empirical[genT], ".png"))
    # par(mar = c(5, 5, 4, 5) + 0.3)
    # try(plot(h1$mids,h1$density/sum(h1$density), ty='l', lwd = 5, cex.axis = 2, ylim = c(min(c(0)), max(c(1))), col = 2, lty = 1,
    #          xlab = list("Population resistance trait variation", font = 3, cex = 2), ylab = list("Density", font = 3, cex = 2), main = ""))
    # lines(h0$mids, h0$density/sum(h0$density), ty='l', lwd = 3, col = 1, lty = 1)
    # polygon(c(h1$mids,rev(h1$mids)), c(BW1[,1], rev(BW1[,2])),  border=NA, col=adjustcolor("red", alpha.f = 0.3))
    # polygon(c(h0$mids,rev(h0$mids)), c(BW0[,1], rev(BW0[,2])),  border=NA, col=adjustcolor("gray", alpha.f = 0.5))
    # 
    # #legend("topright", legend = list("Choice behaviour","Fixed", "Evolving"), col = c(NA,1,2), lwd = c(NA,3,5), lty = 1)
    # dev.off()
    
    
    MinCho = min(c(empCchoice_popvariancemean1, empCchoice_popvariancemean0), na.rm = TRUE)
    MaxCho = max(c(empCchoice_popvariancemean1, empCchoice_popvariancemean0), na.rm = TRUE)
    
    h1 = try(hist(empCchoice_popvariancemean1[genT,,sprank], breaks = seq(MinCho, MaxCho, length.out = binsize)))
    h0 = try(hist(empCchoice_popvariancemean0[genT,,sprank], breaks = seq(MinCho, MaxCho, length.out = binsize)))
    
    
    BN1 = BinomialCI_normal(X=h1$density,n=sum(h1$density),z=zz)
    BN0 = BinomialCI_normal(X=h0$density,n=sum(h0$density),z=zz)
    
    BW1 = BinomialCI_Wilson(X=h1$density,n=sum(h1$density),z=zz)
    BW0 = BinomialCI_Wilson(X=h0$density,n=sum(h0$density),z=zz)
    
    png(paste0(getwd(),"/", subFolder, "/", plotFolder,"/sdpopChoiceTrait_Both_BCI_Normal", "binsize_", binsize, "Species", sprank, "_gen",targetTtimes_empirical[genT], ".png"))
    par(mar = c(5, 5, 4, 5) + 0.3)
    try(plot(h1$mids,h1$density/sum(h1$density), ty='l', lwd = 5, cex.axis = 2, ylim = c(min(c(0)), max(c(1))), col = 2, lty = 1,
             xlab = list("Population choice trait variation", font = 3, cex = 2), ylab = list("Density", font = 3, cex = 2), main = ""))
    lines(h0$mids, h0$density/sum(h0$density), ty='l', lwd = 3, col = 1, lty = 1)
    polygon(c(h1$mids,rev(h1$mids)), c(BN1[,1], rev(BN1[,2])),  border=NA, col=adjustcolor("red", alpha.f = 0.3))
    polygon(c(h0$mids,rev(h0$mids)), c(BN0[,1], rev(BN0[,2])),  border=NA, col=adjustcolor("gray", alpha.f = 0.5))
    
    #legend("topright", legend = list("Choice behaviour","Fixed", "Evolving"), col = c(NA,1,2), lwd = c(NA,3,5), lty = 1)
    dev.off()
    
    # png(paste0(getwd(),"/", subFolder, "/", plotFolder,"/sdpopChoiceTrait_Both_BCI_Wilson", "binsize_", binsize, "Species", sprank, "_gen",targetTtimes_empirical[genT], ".png"))
    # par(mar = c(5, 5, 4, 5) + 0.3)
    # try(plot(h1$mids,h1$density/sum(h1$density), ty='l', lwd = 5, cex.axis = 2, ylim = c(min(c(0)), max(c(1))), col = 2, lty = 1,
    #          xlab = list("Population choice trait variation", font = 3, cex = 2), ylab = list("Density", font = 3, cex = 2), main = ""))
    # lines(h0$mids, h0$density/sum(h0$density), ty='l', lwd = 3, col = 1, lty = 1)
    # polygon(c(h1$mids,rev(h1$mids)), c(BW1[,1], rev(BW1[,2])),  border=NA, col=adjustcolor("red", alpha.f = 0.3))
    # polygon(c(h0$mids,rev(h0$mids)), c(BW0[,1], rev(BW0[,2])),  border=NA, col=adjustcolor("gray", alpha.f = 0.5))
    # 
    # #legend("topright", legend = list("Choice behaviour","Fixed", "Evolving"), col = c(NA,1,2), lwd = c(NA,3,5), lty = 1)
    # dev.off()
    
  }
}


#----------------- Speed: Absolute values not used for speed ----------------------
#------------- Speed of adaptation --------------
# subplotFolder = "Speed_NoAbsolute"
# if(!dir.exists(paste0(getwd(), "/", subFolder, "/", plotFolder, "/", subplotFolder))){
#   dir.create(paste0(getwd(), "/", subFolder, "/", plotFolder, "/", subplotFolder))
# }

for(sprank in 1:11){
  #for(spt in 1:length(tTspeed)){
  for(spt in 1:1){
    MminRes = min(c(empnoabs_ResSpeed1[spt,,], empnoabs_ResSpeed0[spt,,]), na.rm = TRUE)
    MmaxRes = max(c(empnoabs_ResSpeed1[spt,,], empnoabs_ResSpeed0[spt,,]), na.rm = TRUE)
    MminCho = min(c(empnoabs_ChoSpeed1[spt,,], empnoabs_ChoSpeed0[spt,,]), na.rm = TRUE)
    MmaxCho = max(c(empnoabs_ChoSpeed1[spt,,], empnoabs_ChoSpeed0[spt,,]), na.rm = TRUE)
    
    MinRes = min(c(empnoabs_ResSpeed1[spt,,sprank], empnoabs_ResSpeed0[spt,,sprank]), na.rm = TRUE)
    MaxRes = max(c(empnoabs_ResSpeed1[spt,,sprank], empnoabs_ResSpeed0[spt,,sprank]), na.rm = TRUE)
    
    if(MinRes<MaxRes){
      h1 = try(hist(empnoabs_ResSpeed1[spt,,sprank], breaks = seq(MinRes, MaxRes, length.out = binsize)))
      h0 = try(hist(empnoabs_ResSpeed0[spt,,sprank], breaks = seq(MinRes, MaxRes, length.out = binsize)))
      
      BN1 = BinomialCI_normal(X=h1$density,n=sum(h1$density),z=zz)
      BN0 = BinomialCI_normal(X=h0$density,n=sum(h0$density),z=zz)
      
      BW1 = BinomialCI_Wilson(X=h1$density,n=sum(h1$density),z=zz)
      BW0 = BinomialCI_Wilson(X=h0$density,n=sum(h0$density),z=zz)
      
      png(paste0(getwd(),"/", subFolder, "/", plotFolder,  "/speednoabsResistanceTrait_atgen",tTspeed[spt],"_Both_BCI_Normal", "binsize_", binsize, "Species", sprank, ".png"))
      par(mar = c(5, 5, 4, 5) + 0.3)
      try(plot(h1$mids,h1$density/sum(h1$density), ty='l', lwd = 5, cex.axis = 2, ylim = c(min(c(0)), max(c(1))), col = 2, lty = 1, xlim = c(MminRes, MmaxRes),
               xlab = list("Evolution rate of resistance", font = 3, cex = 2), ylab = list("Density", font = 3, cex = 2), main = ""))
      lines(h0$mids, h0$density/sum(h0$density), ty='l', lwd = 3, col = 1, lty = 1)
      polygon(c(h1$mids,rev(h1$mids)), c(BN1[,1], rev(BN1[,2])),  border=NA, col=adjustcolor("red", alpha.f = 0.3))
      polygon(c(h0$mids,rev(h0$mids)), c(BN0[,1], rev(BN0[,2])),  border=NA, col=adjustcolor("gray", alpha.f = 0.5))
      
      #legend("topright", legend = list("Choice behaviour","Fixed", "Evolving"), col = c(NA,1,2), lwd = c(NA,3,5), lty = 1)
      dev.off()
      
      # png(paste0(getwd(),"/", subFolder, "/", plotFolder,  "/speednoabsResistanceTrait_atgen",tTspeed[spt],"_Both_BCI_Wilson", "binsize_", binsize, "Species", sprank, ".png"))
      # par(mar = c(5, 5, 4, 5) + 0.3)
      # try(plot(h1$mids,h1$density/sum(h1$density), ty='l', lwd = 5, cex.axis = 2, ylim = c(min(c(0)), max(c(1))), col = 2, lty = 1, xlim = c(MminRes, MmaxRes),
      #          xlab = list("Evolution rate of resistance", font = 3, cex = 2), ylab = list("Density", font = 3, cex = 2), main = ""))
      # lines(h0$mids, h0$density/sum(h0$density), ty='l', lwd = 3, col = 1, lty = 1)
      # polygon(c(h1$mids,rev(h1$mids)), c(BW1[,1], rev(BW1[,2])),  border=NA, col=adjustcolor("red", alpha.f = 0.3))
      # polygon(c(h0$mids,rev(h0$mids)), c(BW0[,1], rev(BW0[,2])),  border=NA, col=adjustcolor("gray", alpha.f = 0.5))
      # 
      # #legend("topright", legend = list("Choice behaviour","Fixed", "Evolving"), col = c(NA,1,2), lwd = c(NA,3,5), lty = 1)
      # dev.off()
    }else{
      
      h1 = try(hist(empnoabs_ResSpeed1[spt,,sprank], breaks = binsize))
      h0 = try(hist(empnoabs_ResSpeed0[spt,,sprank], breaks = binsize))
      
      BN1 = BinomialCI_normal(X=h1$density,n=sum(h1$density),z=zz)
      BN0 = BinomialCI_normal(X=h0$density,n=sum(h0$density),z=zz)
      
      BW1 = BinomialCI_Wilson(X=h1$density,n=sum(h1$density),z=zz)
      BW0 = BinomialCI_Wilson(X=h0$density,n=sum(h0$density),z=zz)
      
      png(paste0(getwd(),"/", subFolder, "/", plotFolder,  "/speednoabsResistanceTrait_atgen",tTspeed[spt],"_Both_BCI_Normal", "binsize_", binsize, "Species", sprank, ".png"))
      par(mar = c(5, 5, 4, 5) + 0.3)
      try(plot(h1$mids,h1$density/sum(h1$density), ty='l', lwd = 5, cex.axis = 2, ylim = c(min(c(0)), max(c(1))), col = 2, lty = 1, xlim = c(MminRes, MmaxRes),
               xlab = list("Evolution rate of resistance", font = 3, cex = 2), ylab = list("Density", font = 3, cex = 2), main = ""))
      lines(h0$mids, h0$density/sum(h0$density), ty='l', lwd = 3, col = 1, lty = 1)
      polygon(c(h1$mids,rev(h1$mids)), c(BN1[,1], rev(BN1[,2])),  border=NA, col=adjustcolor("red", alpha.f = 0.3))
      polygon(c(h0$mids,rev(h0$mids)), c(BN0[,1], rev(BN0[,2])),  border=NA, col=adjustcolor("gray", alpha.f = 0.5))
      
      #legend("topright", legend = list("Choice behaviour","Fixed", "Evolving"), col = c(NA,1,2), lwd = c(NA,3,5), lty = 1)
      dev.off()
      
      # png(paste0(getwd(),"/", subFolder, "/", plotFolder,  "/speednoabsResistanceTrait_atgen",tTspeed[spt],"_Both_BCI_Wilson", "binsize_", binsize, "Species", sprank, ".png"))
      # par(mar = c(5, 5, 4, 5) + 0.3)
      # try(plot(h1$mids,h1$density/sum(h1$density), ty='l', lwd = 5, cex.axis = 2, ylim = c(min(c(0)), max(c(1))), col = 2, lty = 1, xlim = c(MminRes, MmaxRes),
      #          xlab = list("Evolution rate of resistance", font = 3, cex = 2), ylab = list("Density", font = 3, cex = 2), main = ""))
      # lines(h0$mids, h0$density/sum(h0$density), ty='l', lwd = 3, col = 1, lty = 1)
      # polygon(c(h1$mids,rev(h1$mids)), c(BW1[,1], rev(BW1[,2])),  border=NA, col=adjustcolor("red", alpha.f = 0.3))
      # polygon(c(h0$mids,rev(h0$mids)), c(BW0[,1], rev(BW0[,2])),  border=NA, col=adjustcolor("gray", alpha.f = 0.5))
      # 
      # #legend("topright", legend = list("Choice behaviour","Fixed", "Evolving"), col = c(NA,1,2), lwd = c(NA,3,5), lty = 1)
      # dev.off()
    }
    
    
    MinCho = min(c(empnoabs_ChoSpeed1[spt,,sprank], empnoabs_ChoSpeed0[spt,,sprank]), na.rm = TRUE)
    MaxCho = max(c(empnoabs_ChoSpeed1[spt,,sprank], empnoabs_ChoSpeed0[spt,,sprank]), na.rm = TRUE)
    
    if(MinCho<MaxCho){
      
      h1 = try(hist(empnoabs_ChoSpeed1[spt,,sprank], breaks = seq(MinCho, MaxCho, length.out = binsize), plot = FALSE))
      h0 = try(hist(empnoabs_ChoSpeed0[spt,,sprank], breaks = seq(MinCho, MaxCho, length.out = binsize), plot = FALSE))
      
      
      BN1 = BinomialCI_normal(X=h1$density,n=sum(h1$density),z=zz)
      BN0 = BinomialCI_normal(X=h0$density,n=sum(h0$density),z=zz)
      
      BW1 = BinomialCI_Wilson(X=h1$density,n=sum(h1$density),z=zz)
      BW0 = BinomialCI_Wilson(X=h0$density,n=sum(h0$density),z=zz)
      
      png(paste0(getwd(),"/", subFolder, "/", plotFolder,  "/speednoabsChoiceTrait_atgen",tTspeed[spt],"_Both_BCI_Normal", "binsize_", binsize, "Species", sprank, ".png"))
      par(mar = c(5, 5, 4, 5) + 0.3)
      try(plot(h1$mids,h1$density/sum(h1$density), ty='l', lwd = 5, cex.axis = 2, ylim = c(min(c(0)), max(c(1))), col = 2, lty = 1, xlim = c(MminCho, MmaxCho),
               xlab = list("Evolution rate of resistance", font = 3, cex = 2), ylab = list("Density", font = 3, cex = 2), main = ""))
      lines(h0$mids, h0$density/sum(h0$density), ty='l', lwd = 3, col = 1, lty = 1)
      polygon(c(h1$mids,rev(h1$mids)), c(BN1[,1], rev(BN1[,2])),  border=NA, col=adjustcolor("red", alpha.f = 0.3))
      polygon(c(h0$mids,rev(h0$mids)), c(BN0[,1], rev(BN0[,2])),  border=NA, col=adjustcolor("gray", alpha.f = 0.5))
      
      #legend("topright", legend = list("Choice behaviour","Fixed", "Evolving"), col = c(NA,1,2), lwd = c(NA,3,5), lty = 1)
      dev.off()
      
      # png(paste0(getwd(),"/", subFolder, "/", plotFolder,  "/speednoabsChoiceTrait_atgen",tTspeed[spt],"_Both_BCI_Wilson", "binsize_", binsize, "Species", sprank, ".png"))
      # par(mar = c(5, 5, 4, 5) + 0.3)
      # try(plot(h1$mids,h1$density/sum(h1$density), ty='l', lwd = 5, cex.axis = 2, ylim = c(min(c(0)), max(c(1))), col = 2, lty = 1, xlim = c(MminCho, MmaxCho),
      #          xlab = list("Evolution rate of resistance", font = 3, cex = 2), ylab = list("Density", font = 3, cex = 2), main = ""))
      # lines(h0$mids, h0$density/sum(h0$density), ty='l', lwd = 3, col = 1, lty = 1)
      # polygon(c(h1$mids,rev(h1$mids)), c(BW1[,1], rev(BW1[,2])),  border=NA, col=adjustcolor("red", alpha.f = 0.3))
      # polygon(c(h0$mids,rev(h0$mids)), c(BW0[,1], rev(BW0[,2])),  border=NA, col=adjustcolor("gray", alpha.f = 0.5))
      # 
      # #legend("topright", legend = list("Choice behaviour","Fixed", "Evolving"), col = c(NA,1,2), lwd = c(NA,3,5), lty = 1)
      # dev.off()
      
      png(paste0(getwd(),"/", subFolder, "/", plotFolder,  "/speednoabsChoiceTrait_atgen",tTspeed[spt],"_Both", "binsize_", binsize, "Species", sprank, ".png"))
      par(mar = c(5, 5, 4, 5) + 0.3)
      try(plot(h1$mids, h1$density, ty='l', lwd = 5, cex.axis = 2, ylim = c(min(c(h1$density, h0$density)), max(c(h1$density, h0$density))), col = 2, xlim = c(MminCho, MmaxCho),
               xlab = list("Evolution rate of choice", font = 3, cex = 2), ylab = list("Density", font = 3, cex = 2), main = ""))
      lines(h0$mids, h0$density, ty='l', lwd = 3, col = 1)
      #legend("topright", legend = list("Behaviour","Random", "Choice"), col = c(NA,1,2), lwd = c(NA,3,5), lty = 1)
      dev.off()
    }else{
      
      h1 = try(hist(empnoabs_ChoSpeed1[spt,,sprank], breaks = binsize, plot = FALSE))
      h0 = try(hist(empnoabs_ChoSpeed0[spt,,sprank], breaks = binsize, plot = FALSE))
      
      
      BN1 = BinomialCI_normal(X=h1$density,n=sum(h1$density),z=zz)
      BN0 = BinomialCI_normal(X=h0$density,n=sum(h0$density),z=zz)
      
      BW1 = BinomialCI_Wilson(X=h1$density,n=sum(h1$density),z=zz)
      BW0 = BinomialCI_Wilson(X=h0$density,n=sum(h0$density),z=zz)
      
      png(paste0(getwd(),"/", subFolder, "/", plotFolder,  "/speednoabsChoiceTrait_atgen",tTspeed[spt],"_Both_BCI_Normal", "binsize_", binsize, "Species", sprank, ".png"))
      par(mar = c(5, 5, 4, 5) + 0.3)
      try(plot(h1$mids,h1$density/sum(h1$density), ty='l', lwd = 5, cex.axis = 2, ylim = c(min(c(0)), max(c(1))), col = 2, lty = 1, xlim = c(MminCho, MmaxCho),
               xlab = list("Evolution rate of resistance", font = 3, cex = 2), ylab = list("Density", font = 3, cex = 2), main = ""))
      lines(h0$mids, h0$density/sum(h0$density), ty='l', lwd = 3, col = 1, lty = 1)
      polygon(c(h1$mids,rev(h1$mids)), c(BN1[,1], rev(BN1[,2])),  border=NA, col=adjustcolor("red", alpha.f = 0.3))
      polygon(c(h0$mids,rev(h0$mids)), c(BN0[,1], rev(BN0[,2])),  border=NA, col=adjustcolor("gray", alpha.f = 0.5))
      
      #legend("topright", legend = list("Choice behaviour","Fixed", "Evolving"), col = c(NA,1,2), lwd = c(NA,3,5), lty = 1)
      dev.off()
      
      # png(paste0(getwd(),"/", subFolder, "/", plotFolder,  "/speednoabsChoiceTrait_atgen",tTspeed[spt],"_Both_BCI_Wilson", "binsize_", binsize, "Species", sprank, ".png"))
      # par(mar = c(5, 5, 4, 5) + 0.3)
      # try(plot(h1$mids,h1$density/sum(h1$density), ty='l', lwd = 5, cex.axis = 2, ylim = c(min(c(0)), max(c(1))), col = 2, lty = 1, xlim = c(MminCho, MmaxCho),
      #          xlab = list("Evolution rate of resistance", font = 3, cex = 2), ylab = list("Density", font = 3, cex = 2), main = ""))
      # lines(h0$mids, h0$density/sum(h0$density), ty='l', lwd = 3, col = 1, lty = 1)
      # polygon(c(h1$mids,rev(h1$mids)), c(BW1[,1], rev(BW1[,2])),  border=NA, col=adjustcolor("red", alpha.f = 0.3))
      # polygon(c(h0$mids,rev(h0$mids)), c(BW0[,1], rev(BW0[,2])),  border=NA, col=adjustcolor("gray", alpha.f = 0.5))
      # 
      # #legend("topright", legend = list("Choice behaviour","Fixed", "Evolving"), col = c(NA,1,2), lwd = c(NA,3,5), lty = 1)
      # dev.off()
      
      png(paste0(getwd(),"/", subFolder, "/", plotFolder,  "/speednoabsChoiceTrait_atgen",tTspeed[spt],"_Both", "binsize_", binsize, "Species", sprank, ".png"))
      par(mar = c(5, 5, 4, 5) + 0.3)
      try(plot(h1$mids, h1$density, ty='l', lwd = 5, cex.axis = 2, ylim = c(min(c(h1$density, h0$density)), max(c(h1$density, h0$density))), col = 2, xlim = c(MminCho, MmaxCho),
               xlab = list("Evolution rate of choice", font = 3, cex = 2), ylab = list("Density", font = 3, cex = 2), main = ""))
      lines(h0$mids, h0$density, ty='l', lwd = 3, col = 1)
      #legend("topright", legend = list("Behaviour","Random", "Choice"), col = c(NA,1,2), lwd = c(NA,3,5), lty = 1)
      dev.off()
    }
  }
}


#------------------- Summary statistics -------------------------------
spmeanrepRes1=spmeanrepRes0=spmeanrepCho1=spmeanrepCho0=spsdrepRes1=spsdrepRes0=spsdrepCho1=spsdrepCho0=spmeanmeanrepRes1=spmeanmeanrepRes0=spmeanmeanrepCho1=spmeanmeanrepCho0=spsdmeanrepRes1=spsdmeanrepRes0=spsdmeanrepCho1=spsdmeanrepCho0=spmeanvarmeanbtnRes1=spmeanvarmeanbtnRes0=spmeanvarmeanbtnCho1=spmeanvarmeanbtnCho0=spmeanvarmeanwtnRes1=spmeanvarmeanwtnRes0=spmeanvarmeanwtnCho1=spmeanvarmeanwtnCho0=spsdvarmeanwtnRes1=spsdvarmeanwtnRes0=spsdvarmeanwtnCho1=spsdvarmeanwtnCho0=spsdvarmeanbtnRes1=spsdvarmeanbtnRes0=spsdvarmeanbtnCho1=spsdvarmeanbtnCho0=matrix(NA, nrow = 11, ncol = length(targetTtimes_empirical))
for(genT in 1:length(targetTtimes_empirical)){
  for(sprank in c(1:2,4,6,8,9,10,11)){
    #---- Mean from replicates ----
    spmeanrepRes1[sprank,genT]   = mean(c(RawPopResistance1[genT,,,sprank]), na.rm = TRUE)
    spmeanrepRes0[sprank,genT]   = mean(c(RawPopResistance0[genT,,,sprank]), na.rm = TRUE)
    
    
    spmeanrepCho1[sprank,genT]   = mean(c(RawPopChoice1[genT,,,sprank]), na.rm = TRUE)
    spmeanrepCho0[sprank,genT]   = mean(c(RawPopChoice0[genT,,,sprank]), na.rm = TRUE)
    
    
    # spsdrepRes1[sprank,genT]   = sd(c(RawPopResistance1[genT,,,sprank]), na.rm = TRUE)
    # spsdrepRes0[sprank,genT]   = sd(c(RawPopResistance0[genT,,,sprank]), na.rm = TRUE)
    
    spsdrepRes1[sprank,genT]   = std_error(c(RawPopResistance1[genT,,,sprank]))
    spsdrepRes0[sprank,genT]   = std_error(c(RawPopResistance0[genT,,,sprank]))
    
    
    #spsdrepCho1[sprank,genT]   = sd(c(RawPopChoice1[genT,,,sprank]), na.rm = TRUE)
    #spsdrepCho0[sprank,genT]   = sd(c(RawPopChoice0[genT,,,sprank]), na.rm = TRUE)
    
    spsdrepCho1[sprank,genT]   = std_error(c(RawPopChoice1[genT,,,sprank]))
    spsdrepCho0[sprank,genT]   = std_error(c(RawPopChoice0[genT,,,sprank]))
    
    #------------ Mean of means traits --------------------
    
    spmeanmeanrepRes1[sprank,genT]   = mean(c(empRresistancemean1[genT,,sprank]), na.rm = TRUE)
    spmeanmeanrepRes0[sprank,genT]   = mean(c(empRresistancemean0[genT,,sprank]), na.rm = TRUE)
    
    spmeanmeanrepCho1[sprank,genT]   = mean(c(empCchoicemean1[genT,,sprank]), na.rm = TRUE)
    spmeanmeanrepCho0[sprank,genT]   = mean(c(empCchoicemean0[genT,,sprank]), na.rm = TRUE)
    
    #spsdmeanrepRes1[sprank,genT]   = sd(c(empRresistancemean1[genT,,sprank]), na.rm = TRUE)
    #spsdmeanrepRes0[sprank,genT]   = sd(c(empRresistancemean0[genT,,sprank]), na.rm = TRUE)
    
    spsdmeanrepRes1[sprank,genT]   = std_error(c(empRresistancemean1[genT,,sprank]))
    spsdmeanrepRes0[sprank,genT]   = std_error(c(empRresistancemean0[genT,,sprank]))
    
    #spsdmeanrepCho1[sprank,genT]   = sd(c(empCchoicemean1[genT,,sprank]), na.rm = TRUE)
    #spsdmeanrepCho0[sprank,genT]   = sd(c(empCchoicemean0[genT,,sprank]), na.rm = TRUE)
    
    spsdmeanrepCho1[sprank,genT]   = std_error(c(empCchoicemean1[genT,,sprank]))
    spsdmeanrepCho0[sprank,genT]   = std_error(c(empCchoicemean0[genT,,sprank]))
    #--------- sd between populations -------------
    
    spmeanvarmeanbtnRes1[sprank,genT]   = mean(c(empRresistancesd1[genT,,sprank]), na.rm = TRUE)
    spmeanvarmeanbtnRes0[sprank,genT]   = mean(c(empRresistancesd0[genT,,sprank]), na.rm = TRUE)
    
    spmeanvarmeanbtnCho1[sprank,genT]   = mean(c(empCchoicesd1[genT,,sprank]), na.rm = TRUE)
    spmeanvarmeanbtnCho0[sprank,genT]   = mean(c(empCchoicesd0[genT,,sprank]), na.rm = TRUE)
    
    spsdvarmeanbtnRes1[sprank,genT]   = sd(c(empRresistancesd1[genT,,sprank]), na.rm = TRUE)
    spsdvarmeanbtnRes0[sprank,genT]   = sd(c(empRresistancesd0[genT,,sprank]), na.rm = TRUE)
    
    spsdvarmeanbtnCho1[sprank,genT]   = sd(c(empCchoicesd1[genT,,sprank]), na.rm = TRUE)
    spsdvarmeanbtnCho0[sprank,genT]   = sd(c(empCchoicesd0[genT,,sprank]), na.rm = TRUE)
    
    #------------- sd within populations --------------
    spmeanvarmeanwtnRes1[sprank,genT]   = mean(c(empRresistance_popvariancemean1[genT,,sprank]), na.rm = TRUE)
    spmeanvarmeanwtnRes0[sprank,genT]   = mean(c(empRresistance_popvariancemean0[genT,,sprank]), na.rm = TRUE)
    
    spmeanvarmeanwtnCho1[sprank,genT]   = mean(c(empCchoice_popvariancemean1[genT,,sprank]), na.rm = TRUE)
    spmeanvarmeanwtnCho0[sprank,genT]   = mean(c(empCchoice_popvariancemean0[genT,,sprank]), na.rm = TRUE)
    
    # spsdvarmeanwtnRes1[sprank,genT]   = sd(c(empRresistance_popvariancemean1[genT,,sprank]), na.rm = TRUE)
    # spsdvarmeanwtnRes0[sprank,genT]   = sd(c(empRresistance_popvariancemean0[genT,,sprank]), na.rm = TRUE)
    
    spsdvarmeanwtnRes1[sprank,genT]   = std_error(c(empRresistance_popvariancemean1[genT,,sprank]))
    spsdvarmeanwtnRes0[sprank,genT]   = std_error(c(empRresistance_popvariancemean0[genT,,sprank]))
    
    # spsdvarmeanwtnCho1[sprank,genT]   = sd(c(empCchoice_popvariancemean1[genT,,sprank]), na.rm = TRUE)
    # spsdvarmeanwtnCho0[sprank,genT]   = sd(c(empCchoice_popvariancemean0[genT,,sprank]), na.rm = TRUE)
    
    spsdvarmeanwtnCho1[sprank,genT]   = std_error(c(empCchoice_popvariancemean1[genT,,sprank]))
    spsdvarmeanwtnCho0[sprank,genT]   = std_error(c(empCchoice_popvariancemean0[genT,,sprank]))
    
  }
}


for(genT in 1:length(targetTtimes_empirical)){
  #----------------------------- mean Choice resistance traits at mean levels -------------------
  yyr1 = spmeanmeanrepRes1[,genT][!is.na(spsdmeanrepRes1[,genT])]
  sdyyr1 = spsdmeanrepRes1[,genT][!is.na(spsdmeanrepRes1[,genT])]
  xxr1 = c(1:11)[!is.na(spsdmeanrepRes1[,genT])]
  yyr0 = spmeanmeanrepRes0[,genT][!is.na(spsdmeanrepRes0[,genT])]
  sdyyr0 = spsdmeanrepRes0[,genT][!is.na(spsdmeanrepRes0[,genT])]
  xxr0 = c(1:11)[!is.na(spsdmeanrepRes0[,genT])]
  
  yyc1 = spmeanmeanrepCho1[,genT][!is.na(spsdmeanrepCho1[,genT])]
  sdyyc1 = spsdmeanrepCho1[,genT][!is.na(spsdmeanrepCho1[,genT])]
  xxc1 = c(1:11)[!is.na(spsdmeanrepCho1[,genT])]
  yyc0 = spmeanmeanrepCho0[,genT][!is.na(spsdmeanrepCho0[,genT])]
  sdyyc0 = spsdmeanrepCho0[,genT][!is.na(spsdmeanrepCho0[,genT])]
  xxc0 = c(1:11)[!is.na(spsdmeanrepCho0[,genT])]
  
  # meanResistanceTrait
  png(paste0(getwd(),"/", subFolder, "/", plotFolder, "/spsrank_meanResistanceTrait_MeansEmp", "_gen",targetTtimes_empirical[genT], "r.png"))
  par(mar = c(5, 5, 4, 5) + 0.3)
  plot(xxr1, yyr1, ty='p', lwd = 5, cex.axis = 2, ylim = c(0,1), col = "red", lty = 1, cex=2, pch=16,
       xlab = list("Species", font = 3, cex = 2), ylab = list("Resistance trait mean ", font = 3, cex = 2), main = "" )
  points(xxr0, yyr0, col=adjustcolor("red", alpha.f = 1), pch = 1, lwd=3, lty=2, ty="p")
  #polygon(c(xxr1,rev(xxr1)), c(yyr1+sdyyr1, rev(yyr1-sdyyr1)),  border=NA, col=adjustcolor("red", alpha.f = 0.3))
  #polygon(c(xxr0,rev(xxr0)), c(yyr0+sdyyr0, rev(yyr0-sdyyr0)),  border=NA, col=adjustcolor("red", alpha.f = 0.3))
  
  # points(xxc1, yyc1, col=adjustcolor("black", alpha.f = 1), pch = 16, lwd=5, lty=1, ty="p", cex=2)
  # points(xxc0, yyc0, col=adjustcolor("black", alpha.f = 1), pch = 1, lwd=3, lty=2, ty="p")
  # #polygon(c(xxc1,rev(xxc1)), c(yyc1+sdyyc1, rev(yyc1-sdyyc1)),  border=NA, col=adjustcolor("gray", alpha.f = 0.3))
  # #polygon(c(xxc0,rev(xxc0)), c(yyc0+sdyyc0, rev(yyc0-sdyyc0)),  border=NA, col=adjustcolor("gray", alpha.f = 0.3))
  abline(v=c(8,11), col=c("gray", adjustcolor("black", alpha.f = 5)))
  legend("bottomright", legend = c("Behaviour", "Evolving", "Fixed"), pt.cex = c(2,2,1), cex = 1.5, col=c(NA,"red","red"), pch = c(1,16,1))
  
  dev.off()
  
  # meanChoiceTrait
  png(paste0(getwd(),"/", subFolder, "/", plotFolder, "/spsrank_meanChoiceTrait_MeansEmp", "_gen",targetTtimes_empirical[genT], "r.png"))
  par(mar = c(5, 5, 4, 5) + 0.3)
  plot(xxr1, yyc1, ty='p', lwd = 5, cex.axis = 2, ylim = c(0,1), col = "black", lty = 1, cex=2, pch=16,
       xlab = list("Species", font = 3, cex = 2), ylab = list("Choice trait means ", font = 3, cex = 2), main = "" )
  #points(xxr0, yyr0, col=adjustcolor("red", alpha.f = 1), pch = 1, lwd=3, lty=2, ty="p")
  #polygon(c(xxr1,rev(xxr1)), c(yyr1+sdyyr1, rev(yyr1-sdyyr1)),  border=NA, col=adjustcolor("red", alpha.f = 0.3))
  #polygon(c(xxr0,rev(xxr0)), c(yyr0+sdyyr0, rev(yyr0-sdyyr0)),  border=NA, col=adjustcolor("red", alpha.f = 0.3))
  
  points(xxc1, yyc1, col=adjustcolor("black", alpha.f = 1), pch = 16, lwd=5, lty=1, ty="p", cex=2)
  points(xxc0, yyc0, col=adjustcolor("black", alpha.f = 1), pch = 1, lwd=3, lty=2, ty="p")
  # #polygon(c(xxc1,rev(xxc1)), c(yyc1+sdyyc1, rev(yyc1-sdyyc1)),  border=NA, col=adjustcolor("gray", alpha.f = 0.3))
  # #polygon(c(xxc0,rev(xxc0)), c(yyc0+sdyyc0, rev(yyc0-sdyyc0)),  border=NA, col=adjustcolor("gray", alpha.f = 0.3))
  abline(v=c(8,11), col=c("gray", adjustcolor("black", alpha.f = 5)))
  legend("bottomright", legend = c("Behaviour", "Evolving", "Fixed"), pt.cex = c(2,2,1), cex = 1.5, col=c(NA,1,1), pch = c(1,16,1))
  
  dev.off()
  
  # For calculating the difference between a given generation and initial generation
  if(genT>1){
    yyr1i = spmeanmeanrepRes1[,1][!is.na(spsdmeanrepRes1[,genT])]
    sdyyr1i = spsdmeanrepRes1[,1][!is.na(spsdmeanrepRes1[,genT])]
    xxr1i = c(1:11)[!is.na(spsdmeanrepRes1[,genT])]
    yyr0i = spmeanmeanrepRes0[,1][!is.na(spsdmeanrepRes0[,genT])]
    sdyyr0i = spsdmeanrepRes0[,1][!is.na(spsdmeanrepRes0[,genT])]
    xxr0i = c(1:11)[!is.na(spsdmeanrepRes0[,genT])]
    
    yyc1i = spmeanmeanrepCho1[,1][!is.na(spsdmeanrepCho1[,genT])]
    sdyyc1i = spsdmeanrepCho1[,1][!is.na(spsdmeanrepCho1[,genT])]
    xxc1i = c(1:11)[!is.na(spsdmeanrepCho1[,genT])]
    yyc0i = spmeanmeanrepCho0[,1][!is.na(spsdmeanrepCho0[,genT])]
    sdyyc0i = spsdmeanrepCho0[,1][!is.na(spsdmeanrepCho0[,genT])]
    xxc0i = c(1:11)[!is.na(spsdmeanrepCho0[,genT])]
    ylm = c(yyc1-yyc1i, yyc0-yyc0i,yyc1-yyc1i+sdyyc1-sdyyc1i, yyc0-yyc0i+sdyyc0-sdyyc0i,yyr1-yyr1i, yyr0-yyr0i, yyr1-yyr1i+sdyyr1-sdyyr1i, yyr0-yyr0i+sdyyr0-sdyyr0i)
    ylm = c(yyr1-yyr1i, yyr0-yyr0i, yyr1-yyr1i+sdyyr1-sdyyr1i, yyr0-yyr0i+sdyyr0-sdyyr0i)
    png(paste0(getwd(),"/", subFolder, "/", plotFolder, "/spsrank_diffmeanResistanceTrait_MeansEmp", "_gen",targetTtimes_empirical[genT], "r.png"))
    par(mar = c(5, 5, 4, 5) + 0.3)
    plot(xxr1, yyr1-yyr1i, ty='p', lwd = 5, cex.axis = 2, ylim = c(min(ylm),max(ylm)), col = "red", lty = 1, cex=2, pch = 16,
         xlab = list("Species", font = 3, cex = 2), ylab = list("Change in resistance trait", font = 3, cex = 2), main = "" )
    points(xxr0, yyr0-yyr0i, col="red", pch = 1, lwd=3, lty=2, ty="p", cex = 1)
    #polygon(c(xxr1,rev(xxr1)), c(yyr1-yyr1i+abs(sdyyr1-sdyyr1i), rev(yyr1-yyr1i-abs(sdyyr1-sdyyr1i))),  border=NA, col=adjustcolor("red", alpha.f = 0.3))
    #polygon(c(xxr0,rev(xxr0)), c(yyr0-yyr0i+abs(sdyyr0-sdyyr0i), rev(yyr0-yyr0i-abs(sdyyr0-sdyyr0i))),  border=NA, col=adjustcolor("red", alpha.f = 0.3))
    
    # points(xxc1, yyc1-yyc1i, col="black", pch = 16, lwd=5, lty=1, ty="p", cex=2)
    # points(xxc0, yyc0-yyc0i, col="black", pch = 1, lwd=3, lty=2, ty="p")
    # #polygon(c(xxc1,rev(xxc1)), c(yyc1-yyc1i+abs(sdyyc1-sdyyc1i), rev(yyc1-yyc1i-abs(sdyyc1-sdyyc1i))),  border=NA, col=adjustcolor("gray", alpha.f = 0.3))
    # #polygon(c(xxc0,rev(xxc0)), c(yyc0-yyc0i+abs(sdyyc0-sdyyc0i), rev(yyc0-yyc0i-abs(sdyyc0-sdyyc0i))),  border=NA, col=adjustcolor("gray", alpha.f = 0.3))
    abline(v=c(8,11), col=c("gray", adjustcolor("black", alpha.f = 5)))
    # legend("topright", legend = c("Behaviour", "Evolving", "Fixed"), pt.cex = c(2,2,1), cex = 1.5, col=c(NA,2,2), pch = c(1,16,1))
    
    dev.off()
    
    png(paste0(getwd(),"/", subFolder, "/", plotFolder, "/spsrank_diffmeanChoiceTrait_MeansEmp", "_gen",targetTtimes_empirical[genT], "r.png"))
    par(mar = c(5, 5, 4, 5) + 0.3)
    plot(xxr1, yyc1-yyc1i, ty='p', lwd = 5, cex.axis = 2, ylim = c(min(ylm),max(ylm)), col = "black", lty = 1, cex=2, pch = 16,
         xlab = list("Species", font = 3, cex = 2), ylab = list("Change in choice trait", font = 3, cex = 2), main = "" )
    #points(xxr0, yyr0-yyr0i, col="red", pch = 1, lwd=3, lty=2, ty="p", cex = 1)
    #polygon(c(xxr1,rev(xxr1)), c(yyr1-yyr1i+abs(sdyyr1-sdyyr1i), rev(yyr1-yyr1i-abs(sdyyr1-sdyyr1i))),  border=NA, col=adjustcolor("red", alpha.f = 0.3))
    #polygon(c(xxr0,rev(xxr0)), c(yyr0-yyr0i+abs(sdyyr0-sdyyr0i), rev(yyr0-yyr0i-abs(sdyyr0-sdyyr0i))),  border=NA, col=adjustcolor("red", alpha.f = 0.3))
    
    points(xxc1, yyc1-yyc1i, col="black", pch = 16, lwd=5, lty=1, ty="p", cex=2)
    points(xxc0, yyc0-yyc0i, col="black", pch = 1, lwd=3, lty=2, ty="p")
    # #polygon(c(xxc1,rev(xxc1)), c(yyc1-yyc1i+abs(sdyyc1-sdyyc1i), rev(yyc1-yyc1i-abs(sdyyc1-sdyyc1i))),  border=NA, col=adjustcolor("gray", alpha.f = 0.3))
    # #polygon(c(xxc0,rev(xxc0)), c(yyc0-yyc0i+abs(sdyyc0-sdyyc0i), rev(yyc0-yyc0i-abs(sdyyc0-sdyyc0i))),  border=NA, col=adjustcolor("gray", alpha.f = 0.3))
    abline(v=c(8,11), col=c("gray", adjustcolor("black", alpha.f = 5)))
    #legend("topright", legend = c("Behaviour", "Evolving", "Fixed"), pt.cex = c(2,2,1), cex = 1.5, col=c(NA,1,1), pch = c(1,16,1))
    
    dev.off()
  }
  
  
  #----------------------------- Variance within populations Choice resistance traits at mean levels -------------------
  yyr1 = spmeanvarmeanwtnRes1[,genT][!is.na(spsdvarmeanwtnRes1[,genT])]
  sdyyr1 = spsdvarmeanwtnRes1[,genT][!is.na(spsdvarmeanwtnRes1[,genT])]
  xxr1 = c(1:11)[!is.na(spsdvarmeanwtnRes1[,genT])]
  yyr0 = spmeanvarmeanwtnRes0[,genT][!is.na(spsdvarmeanwtnRes0[,genT])]
  sdyyr0 = spsdvarmeanwtnRes0[,genT][!is.na(spsdvarmeanwtnRes0[,genT])]
  xxr0 = c(1:11)[!is.na(spsdvarmeanwtnRes0[,genT])]
  
  yyc1 = spmeanvarmeanwtnCho1[,genT][!is.na(spsdvarmeanwtnCho1[,genT])]
  sdyyc1 = spsdvarmeanwtnCho1[,genT][!is.na(spsdvarmeanwtnCho1[,genT])]
  xxc1 = c(1:11)[!is.na(spsdvarmeanwtnCho1[,genT])]
  yyc0 = spmeanvarmeanwtnCho0[,genT][!is.na(spsdvarmeanwtnCho0[,genT])]
  sdyyc0 = spsdvarmeanwtnCho0[,genT][!is.na(spsdvarmeanwtnCho0[,genT])]
  xxc0 = c(1:11)[!is.na(spsdvarmeanwtnCho0[,genT])]
  ylm = c(yyc1, yyc0,yyc1+sdyyc1, yyc0+sdyyc0,yyr1, yyr0, yyr1+sdyyr1, yyr0+sdyyr0)
  ylm = c(yyr1, yyr0, yyr1+sdyyr1, yyr0+sdyyr0)
  png(paste0(getwd(),"/", subFolder, "/", plotFolder, "/spsrank_varwithinResistanceTrait", "_gen",targetTtimes_empirical[genT], "r.png"))
  par(mar = c(5, 5, 4, 5) + 0.3)
  plot(xxr1, yyr1, ty='p', lwd = 5, cex.axis = 2, ylim = c(min(ylm),max(ylm)), col = "red", lty = 1, cex=2, pch=16,
       xlab = list("Species", font = 3, cex = 2), ylab = list("Resistance trait variances ", font = 3, cex = 2), main = "" )
  points(xxr0, yyr0, col=adjustcolor("red", alpha.f = 1), pch = 1, lwd=3, lty=2, ty="p")
  #polygon(c(xxr1,rev(xxr1)), c(yyr1+sdyyr1, rev(yyr1-sdyyr1)),  border=NA, col=adjustcolor("red", alpha.f = 0.3))
  #polygon(c(xxr0,rev(xxr0)), c(yyr0+sdyyr0, rev(yyr0-sdyyr0)),  border=NA, col=adjustcolor("red", alpha.f = 0.3))
  
  # points(xxc1, yyc1, col=adjustcolor("black", alpha.f = 1), pch = 16, lwd=5, lty=1, ty="p", cex=2)
  # points(xxc0, yyc0, col=adjustcolor("black", alpha.f = 1), pch = 1, lwd=3, lty=2, ty="p")
  # #polygon(c(xxc1,rev(xxc1)), c(yyc1+sdyyc1, rev(yyc1-sdyyc1)),  border=NA, col=adjustcolor("gray", alpha.f = 0.3))
  # #polygon(c(xxc0,rev(xxc0)), c(yyc0+sdyyc0, rev(yyc0-sdyyc0)),  border=NA, col=adjustcolor("gray", alpha.f = 0.3))
  abline(v=c(8,11), col=c("gray", adjustcolor("black", alpha.f = 5)))
  #legend("topleft", legend = c("Behaviour", "Evolving", "Fixed", "Trait", "Resistance", "Choice"), pt.cex = c(2,2,1,1,2,2), cex = 1.5, col=c(NA,1,1,NA,"red","black"), pch = c(1,16,1,16,16,16))
  legend("topleft", legend = c("Behaviour", "Evolving", "Fixed"), pt.cex = c(2,2,1), cex = 1.5, col=c(NA,2,2), pch = c(1,16,1))
  
  dev.off()
  
  
  png(paste0(getwd(),"/", subFolder, "/", plotFolder, "/spsrank_varwithinChoiceTrait", "_gen",targetTtimes_empirical[genT], "r.png"))
  par(mar = c(5, 5, 4, 5) + 0.3)
  plot(xxc1, yyc1, ty='p', lwd = 5, cex.axis = 2, ylim = c(min(c(yyc1,yyc0)),max(c(yyc1,yyc0))), col = "black", lty = 1, cex=2, pch=16,
       xlab = list("Species", font = 3, cex = 2), ylab = list("Choice trait variances ", font = 3, cex = 2), main = "" )
  #points(xxr0, yyr0, col=adjustcolor("red", alpha.f = 1), pch = 1, lwd=3, lty=2, ty="p")
  #polygon(c(xxr1,rev(xxr1)), c(yyr1+sdyyr1, rev(yyr1-sdyyr1)),  border=NA, col=adjustcolor("red", alpha.f = 0.3))
  #polygon(c(xxr0,rev(xxr0)), c(yyr0+sdyyr0, rev(yyr0-sdyyr0)),  border=NA, col=adjustcolor("red", alpha.f = 0.3))
  
  points(xxc1, yyc1, col=adjustcolor("black", alpha.f = 1), pch = 16, lwd=5, lty=1, ty="p", cex=2)
  points(xxc0, yyc0, col=adjustcolor("black", alpha.f = 1), pch = 1, lwd=3, lty=2, ty="p")
  # #polygon(c(xxc1,rev(xxc1)), c(yyc1+sdyyc1, rev(yyc1-sdyyc1)),  border=NA, col=adjustcolor("gray", alpha.f = 0.3))
  # #polygon(c(xxc0,rev(xxc0)), c(yyc0+sdyyc0, rev(yyc0-sdyyc0)),  border=NA, col=adjustcolor("gray", alpha.f = 0.3))
  abline(v=c(8,11), col=c("gray", adjustcolor("black", alpha.f = 5)))
  #legend("topleft", legend = c("Behaviour", "Evolving", "Fixed", "Trait", "Resistance", "Choice"), pt.cex = c(2,2,1,1,2,2), cex = 1.5, col=c(NA,1,1,NA,"red","black"), pch = c(1,16,1,16,16,16))
  #legend("topleft", legend = c("Behaviour", "Evolving", "Fixed"), pt.cex = c(2,2,1), cex = 1.5, col=c(NA,2,2), pch = c(1,16,1))
  
  dev.off()
  
  if(genT>1){
    yyr1i = spmeanvarmeanwtnRes1[,1][!is.na(spsdvarmeanwtnRes1[,genT])]
    sdyyr1i = spsdvarmeanwtnRes1[,1][!is.na(spsdvarmeanwtnRes1[,genT])]
    xxr1i = c(1:11)[!is.na(spsdvarmeanwtnRes1[,genT])]
    yyr0i = spmeanvarmeanwtnRes0[,1][!is.na(spsdvarmeanwtnRes0[,genT])]
    sdyyr0i = spsdvarmeanwtnRes0[,1][!is.na(spsdvarmeanwtnRes0[,genT])]
    xxr0i = c(1:11)[!is.na(spsdvarmeanwtnRes0[,genT])]
    
    yyc1i = spmeanvarmeanwtnCho1[,1][!is.na(spsdvarmeanwtnCho1[,genT])]
    sdyyc1i = spsdvarmeanwtnCho1[,1][!is.na(spsdvarmeanwtnCho1[,genT])]
    xxc1i = c(1:11)[!is.na(spsdvarmeanwtnCho1[,genT])]
    yyc0i = spmeanvarmeanwtnCho0[,1][!is.na(spsdvarmeanwtnCho0[,genT])]
    sdyyc0i = spsdvarmeanwtnCho0[,1][!is.na(spsdvarmeanwtnCho0[,genT])]
    xxc0i = c(1:11)[!is.na(spsdvarmeanwtnCho0[,genT])]
    ylm = c(yyc1-yyc1i, yyc0-yyc0i,yyc1-yyc1i+sdyyc1-sdyyc1i, yyc0-yyc0i+sdyyc0-sdyyc0i,yyr1-yyr1i, yyr0-yyr0i, yyr1-yyr1i+sdyyr1-sdyyr1i, yyr0-yyr0i+sdyyr0-sdyyr0i)
    ylm = c(yyr1-yyr1i, yyr0-yyr0i, yyr1-yyr1i+sdyyr1-sdyyr1i, yyr0-yyr0i+sdyyr0-sdyyr0i)
    png(paste0(getwd(),"/", subFolder, "/", plotFolder, "/spsrank_diffvarwithinResistanceTrait", "_gen",targetTtimes_empirical[genT], "r.png"))
    par(mar = c(5, 5, 4, 5) + 0.3)
    plot(xxr1, yyr1-yyr1i, ty='p', lwd = 5, cex.axis = 2, ylim = c(min(ylm),max(ylm)), col = "red", lty = 1, cex=2, pch = 16,
         xlab = list("Species", font = 3, cex = 2), ylab = list("Change in resistance variances", font = 3, cex = 2), main = "" )
    points(xxr0, yyr0-yyr0i, col="red", pch = 1, lwd=3, lty=2, ty="p", cex = 1)
    #polygon(c(xxr1,rev(xxr1)), c(yyr1-yyr1i+abs(sdyyr1-sdyyr1i), rev(yyr1-yyr1i-abs(sdyyr1-sdyyr1i))),  border=NA, col=adjustcolor("red", alpha.f = 0.3))
    #polygon(c(xxr0,rev(xxr0)), c(yyr0-yyr0i+abs(sdyyr0-sdyyr0i), rev(yyr0-yyr0i-abs(sdyyr0-sdyyr0i))),  border=NA, col=adjustcolor("red", alpha.f = 0.3))
    
    # points(xxc1, yyc1-yyc1i, col="black", pch = 16, lwd=5, lty=1, ty="p", cex=2)
    # points(xxc0, yyc0-yyc0i, col="black", pch = 1, lwd=3, lty=2, ty="p")
    # #polygon(c(xxc1,rev(xxc1)), c(yyc1-yyc1i+abs(sdyyc1-sdyyc1i), rev(yyc1-yyc1i-abs(sdyyc1-sdyyc1i))),  border=NA, col=adjustcolor("gray", alpha.f = 0.3))
    # #polygon(c(xxc0,rev(xxc0)), c(yyc0-yyc0i+abs(sdyyc0-sdyyc0i), rev(yyc0-yyc0i-abs(sdyyc0-sdyyc0i))),  border=NA, col=adjustcolor("gray", alpha.f = 0.3))
    abline(v=c(8,11), col=c("gray", adjustcolor("black", alpha.f = 5)))
    
    dev.off()
    
    
    png(paste0(getwd(),"/", subFolder, "/", plotFolder, "/spsrank_diffvarwithinChoiceTrait", "_gen",targetTtimes_empirical[genT], "r.png"))
    par(mar = c(5, 5, 4, 5) + 0.3)
    plot(xxr1, yyc1-yyc1i, ty='p', lwd = 5, cex.axis = 2, ylim = c(min(c(yyc1-yyc1i,yyc0-yyc0i)),max(c(yyc1-yyc1i,yyc0-yyc0i))), col = "black", lty = 1, cex=2, pch = 16,
         xlab = list("Species", font = 3, cex = 2), ylab = list("Change in choice variances", font = 3, cex = 2), main = "" )
    #points(xxr0, yyr0-yyr0i, col="red", pch = 1, lwd=3, lty=2, ty="p", cex = 1)
    #polygon(c(xxr1,rev(xxr1)), c(yyr1-yyr1i+abs(sdyyr1-sdyyr1i), rev(yyr1-yyr1i-abs(sdyyr1-sdyyr1i))),  border=NA, col=adjustcolor("red", alpha.f = 0.3))
    #polygon(c(xxr0,rev(xxr0)), c(yyr0-yyr0i+abs(sdyyr0-sdyyr0i), rev(yyr0-yyr0i-abs(sdyyr0-sdyyr0i))),  border=NA, col=adjustcolor("red", alpha.f = 0.3))
    
    points(xxc1, yyc1-yyc1i, col="black", pch = 16, lwd=5, lty=1, ty="p", cex=2)
    points(xxc0, yyc0-yyc0i, col="black", pch = 1, lwd=3, lty=2, ty="p")
    # #polygon(c(xxc1,rev(xxc1)), c(yyc1-yyc1i+abs(sdyyc1-sdyyc1i), rev(yyc1-yyc1i-abs(sdyyc1-sdyyc1i))),  border=NA, col=adjustcolor("gray", alpha.f = 0.3))
    # #polygon(c(xxc0,rev(xxc0)), c(yyc0-yyc0i+abs(sdyyc0-sdyyc0i), rev(yyc0-yyc0i-abs(sdyyc0-sdyyc0i))),  border=NA, col=adjustcolor("gray", alpha.f = 0.3))
    abline(v=c(8,11), col=c("gray", adjustcolor("black", alpha.f = 5)))
    
    dev.off()
  }
}

#------------- Speed of adaptation --------------
meannoabs_speedRes1=meannoabs_speedRes0=sdnoabs_speedRes1=sdnoabs_speedRes0=meannoabs_speedCho1=meannoabs_speedCho0=sdnoabs_speedCho1=sdnoabs_speedCho0 = meanspeedRes1=meanspeedRes0=sdspeedRes1=sdspeedRes0=meanspeedCho1=meanspeedCho0=sdspeedCho1=sdspeedCho0=matrix(NA, nrow = 11, ncol = length(tTspeed))

for(sprank in c(1:2,4,6,8,9,10,11)){
  for(spt in 1:length(tTspeed)){
    #------------- Not Absolute speed -------
    
    meannoabs_speedRes1[sprank,spt] = mean(c(empnoabs_ResSpeed1[spt,,sprank]), na.rm = TRUE)
    meannoabs_speedRes0[sprank,spt] = mean(c(empnoabs_ResSpeed0[spt,,sprank]), na.rm = TRUE)
    
    meannoabs_speedCho1[sprank,spt] = mean(c(empnoabs_ChoSpeed1[spt,,sprank]), na.rm = TRUE)
    meannoabs_speedCho0[sprank,spt] = mean(c(empnoabs_ChoSpeed0[spt,,sprank]), na.rm = TRUE)
    
    # sdnoabs_speedRes1[sprank,spt] = sd(c(empnoabs_ResSpeed1[spt,,sprank]), na.rm = TRUE)
    # sdnoabs_speedRes0[sprank,spt] = sd(c(empnoabs_ResSpeed0[spt,,sprank]), na.rm = TRUE)
    # 
    # sdnoabs_speedCho1[sprank,spt] = sd(c(empnoabs_ChoSpeed1[spt,,sprank]), na.rm = TRUE)
    # sdnoabs_speedCho0[sprank,spt] = sd(c(empnoabs_ChoSpeed0[spt,,sprank]), na.rm = TRUE)
    
    sdnoabs_speedRes1[sprank,spt] = std_error(c(empnoabs_ResSpeed1[spt,,sprank]))
    sdnoabs_speedRes0[sprank,spt] = std_error(c(empnoabs_ResSpeed0[spt,,sprank]))
    
    sdnoabs_speedCho1[sprank,spt] = std_error(c(empnoabs_ChoSpeed1[spt,,sprank]))
    sdnoabs_speedCho0[sprank,spt] = std_error(c(empnoabs_ChoSpeed0[spt,,sprank]))
    #----------------------===============
    
  }
}


for(spt in 1:length(tTspeed)){
  
  
  #-----------------No absolute speed Choice resistance trait at replicate level ---------------------
  yyr1 = meannoabs_speedRes1[,spt][!is.na(sdnoabs_speedRes1[,spt])]
  sdyyr1 = sdnoabs_speedRes1[,spt][!is.na(sdnoabs_speedRes1[,spt])]
  xxr1 = c(1:11)[!is.na(sdnoabs_speedRes1[,spt])]
  yyr0 = meannoabs_speedRes0[,spt][!is.na(sdnoabs_speedRes0[,spt])]
  sdyyr0 = sdnoabs_speedRes0[,spt][!is.na(sdnoabs_speedRes0[,spt])]
  xxr0 = c(1:11)[!is.na(sdnoabs_speedRes0[,spt])]
  
  yyc1 = meannoabs_speedCho1[,spt][!is.na(sdnoabs_speedCho1[,spt])]
  sdyyc1 = sdnoabs_speedCho1[,spt][!is.na(sdnoabs_speedCho1[,spt])]
  xxc1 = c(1:11)[!is.na(sdnoabs_speedCho1[,spt])]
  yyc0 = meannoabs_speedCho0[,spt][!is.na(sdnoabs_speedCho0[,spt])]
  sdyyc0 = sdnoabs_speedCho0[,spt][!is.na(sdnoabs_speedCho0[,spt])]
  xxc0 = c(1:11)[!is.na(sdnoabs_speedCho0[,spt])]
  
  png(paste0(getwd(),"/", subFolder, "/", plotFolder,"/speednoabs_ResistanceTrait_atgen",tTspeed[spt], ".png"))
  par(mar = c(5, 5, 4, 5) + 0.3)
  plot(xxr1, yyr1, ty='p', lwd = 5, cex.axis = 2, ylim = c(min(c(yyr1,yyr0,yyc1,yyc0)),max(c(yyr1,yyr0,yyc1,yyc0))), col = "red", lty = 1, cex=2, pch=16,
       xlab = list("Species", font = 3, cex = 2), ylab = list("Evolution rate of resistance trait", font = 3, cex = 2), main = "" )
  points(xxr0, yyr0, col=adjustcolor("red", alpha.f = 1), pch = 1, lwd=3, lty=2, ty="p")
  #polygon(c(xxr1,rev(xxr1)), c(yyr1+sdyyr1, rev(yyr1-sdyyr1)),  border=NA, col=adjustcolor("red", alpha.f = 0.3))
  #polygon(c(xxr0,rev(xxr0)), c(yyr0+sdyyr0, rev(yyr0-sdyyr0)),  border=NA, col=adjustcolor("red", alpha.f = 0.3))
  
  #points(xxc1, yyc1, col=adjustcolor("black", alpha.f = 1), pch = 16, lwd=5, lty=1, ty="p", cex=2)
  #points(xxc0, yyc0, col=adjustcolor("black", alpha.f = 1), pch = 1, lwd=3, lty=2, ty="p")
  #polygon(c(xxc1,rev(xxc1)), c(yyc1+sdyyc1, rev(yyc1-sdyyc1)),  border=NA, col=adjustcolor("gray", alpha.f = 0.3))
  #polygon(c(xxc0,rev(xxc0)), c(yyc0+sdyyc0, rev(yyc0-sdyyc0)),  border=NA, col=adjustcolor("gray", alpha.f = 0.3))
  abline(v=c(8,11), col=c("gray", adjustcolor("black", alpha.f = 5)))
  
  legend("bottomright", legend = c("Behaviour", "Evolving", "Fixed"), pt.cex = c(2,2,1), cex = 1.5, col=c(NA,2,2), pch = c(1,16,1))
  
  dev.off()
  
  png(paste0(getwd(),"/", subFolder, "/", plotFolder,"/speednoabs_ChoiceTrait_atgen",tTspeed[spt], ".png"))
  par(mar = c(5, 5, 4, 5) + 0.3)
  plot(xxc1, yyc1, ty='p', lwd = 5, cex.axis = 2, ylim = c(min(c(yyr1,yyr0,yyc1,yyc0)),max(c(yyr1,yyr0,yyc1,yyc0))), col = "black", lty = 1, cex=2, pch=16,
       xlab = list("Species", font = 3, cex = 2), ylab = list("Evolution rate of choice trait", font = 3, cex = 2), main = "" )
  #points(xxr0, yyr0, col=adjustcolor("red", alpha.f = 1), pch = 1, lwd=3, lty=2, ty="p")
  #polygon(c(xxr1,rev(xxr1)), c(yyr1+sdyyr1, rev(yyr1-sdyyr1)),  border=NA, col=adjustcolor("red", alpha.f = 0.3))
  #polygon(c(xxr0,rev(xxr0)), c(yyr0+sdyyr0, rev(yyr0-sdyyr0)),  border=NA, col=adjustcolor("red", alpha.f = 0.3))
  
  points(xxc1, yyc1, col=adjustcolor("black", alpha.f = 1), pch = 16, lwd=5, lty=1, ty="p", cex=2)
  points(xxc0, yyc0, col=adjustcolor("black", alpha.f = 1), pch = 1, lwd=3, lty=2, ty="p")
  #polygon(c(xxc1,rev(xxc1)), c(yyc1+sdyyc1, rev(yyc1-sdyyc1)),  border=NA, col=adjustcolor("gray", alpha.f = 0.3))
  #polygon(c(xxc0,rev(xxc0)), c(yyc0+sdyyc0, rev(yyc0-sdyyc0)),  border=NA, col=adjustcolor("gray", alpha.f = 0.3))
  abline(v=c(8,11), col=c("gray", adjustcolor("black", alpha.f = 5)))
  
  legend("bottomright", legend = c("Behaviour", "Evolving", "Fixed"), pt.cex = c(2,2,1), cex = 1.5, col=c(NA,1,1), pch = c(1,16,1))
  
  dev.off()
  
}


setwd(workingdirectory)
```
