## Supplementary material for "The role of evolving niche choice in herbivore adaptation to host plants": These are R scripts and data files used for generating results.: IBM_PlantHerbivore_SimulationAnalysis_SV.R

```
#Make sure that at the end of running this script, you run the last line of this script to reset the working directory
#Here, we upload the saved simulated data for analysis. 
#The parameter setting should be the same as in the execution and analysis scripts
#We illustrate for 4 parameter combinations, which includes the time series. The start of a parameter combination starts by defining the folder name, "Folder", in which data is (to be) kept


library(dplyr)
library(fields)


WD = "PLANTHERBIVORECHOICE"
workingdirectory = getwd()
setwd(paste0(workingdirectory,"/",WD))

mean2  <- function(x) mean(x,na.rm=T)
sd2  <- function(x) sd(x,na.rm=T)
std_error <- function(x) sd(x, na.rm = TRUE)/sqrt(length(!is.na(x)))


numgenerations=200; # Number of generations. Used 200 for the manuscript
nsim = 5;# Number of replicates. For cluster, 1000 was used for the manuscript
numresistanceloci  = 1; #number of loci underlying resistance to plant toxin
numchoiceloci  = 1; #number of loci underlying host plant choice
## Rescaling is done within the function to have the sum of effect sizes sum to 1. 
resistanceeffectsizes = rep(0.5, numresistanceloci); #vector of effect sizes; length should correspond to the number of resistance loci; Converted within the function such that the effect sizes add up to 1
choiceeffectsizes = rep(0.5, numchoiceloci);#vector of effect sizes; length should correspond to the number of choice loci; Converted within the function such that the effect sizes add up to 1
K = 1000 # Carrying capacity
N0 = 20# Initial population size
N_offspring = 4# Mean number of offspring for a susceptible individual
environmentalsdresistance = 0.0 # Environmental noice on resistance trait # NOT USED and SET to 0
environmentalsdchoice = 0.0# Environmental noice on choice trait # NOT USED and SET to 0

#setwd("~/Dropbox/Bielefeld_University/Munster_Collaboration/SimulatedData") # Set working directory
Dominancechoice = c(0,0.1, 0.2,0.5,0.8,0.9, 1) # Dominance choice
Dominanceresistance = c(0,0.1,0.2,0.5,0.8,0.9,1) # Dominance resistance
sortMating = c(-1,-0.8,-0.5,-0.2,0,0.2,0.5,0.8,1) # Host mating preference 
Recombinationrate = c(0, 0.0001, 0.001,0.01, 0.1, 0.2, 0.3, 0.4, 0.5) # Recombination rate
Fecunditycost = c(-1, -0.5, 0, 0.5, 1) # resistance effect on fecundity
Resistancecost = c(-0.5, -0.4, -0.3, -0.2, -0.1, -0.01, 0, 0.001, 0.01, 0.1, 0.2, 0.3, 0.4, 0.5) # Resistance effect on viability
Toxicity = c(0.0, 0.2, 0.5, 0.8, 1 ) # Level of toxicity of the toxic plant
Fr0 = c(0.1,0.3, 0.5,0.7, 0.9)# Initial resistance allele frequency
Fc0 = c(0.1,0.3, 0.5,0.7, 0.9)# Initial choice allele frequency
Scaledensity = c(1, 2, 3, 4)# Habitat size
fixedChoice = 0.5 # random choice behaviour
T20 = 20 # Time to measure speed of evolution # Used 20 in the simulations
T50 = 50 # another time to measure speed of evolution # Used 50 in the simulations
tTspeed = c(T20,T50) 


targetTtimes = c(100,numgenerations)#c(1:numgenerations)
Set = 1# Unused parameter. Leave it as it is. 


Folder = "Dominance_Density_AM" # Folder for storing simulated data
FFolder = Folder

ParameterCombination = 1# Assign different numbers for different parameter combinations to help distinguish the names of plots
Parameter1 = Dominanceresistance
Parameter2 = Dominancechoice
Parameter3 = Scaledensity
Parameter4 = sortMating


#-------------------
noabs_stdResSpeed1 = noabs_stdResSpeed0 = noabs_stdChoSpeed1 = noabs_stdChoSpeed0 = noabs_ResSpeed1 = noabs_ResSpeed0 = noabs_ChoSpeed1 = noabs_ChoSpeed0= Rresistancestd1 = Rresistancestd0 = Cchoicestd1 = Cchoicestd0 = ResSpeed1 = ResSpeed0 = ChoSpeed1 = ChoSpeed0 = N1_mean1 = N1_mean0 = N1_std1 = N1_std0 = N2_mean1 = N2_mean0 = N2_std1 = N2_std0 = Rresistancemean1 = Rresistancesd1 = Rresistancemean0 = Rresistancesd0 = Cchoicemean1 = Cchoicesd1 = Cchoicemean0 = Cchoicesd0 = Rresistance_popvariancemean1 = Rresistance_popvariancestd1 = Rresistance_popvariancemean0 = Rresistance_popvariancestd0 = Cchoice_popvariancemean1 = Cchoice_popvariancestd1 = Cchoice_popvariancemean0 = Cchoice_popvariancestd0= array(NA, dim = c(length(targetTtimes), length(Parameter1), length(Parameter2), length(Parameter3), length(Parameter4)))

#--
# for(para1 in 1:length(Parameter1)){
#   for(para2 in 1:length(Parameter2)){
#     for(para3 in 1:length(Parameter3)){
#       for(para4 in 1:length(Parameter4)){
        for(para1 in c(1,4)){
          for(para2 in c(1,4)){
            for(para3 in c(1,3)){
              for(para4 in c(3,5,7)){
    
        hr = Parameter1[para1]
        hc = Parameter2[para2]
        dominanceresistance = rep(hr, numresistanceloci);#dominance values for the resistance loci, should generally be between 0 and 1
        dominancechoice = rep(hc, numchoiceloci);#dominance values for the choice loci, should generally be between 0 an 1
        scaledensity = Parameter3[para3]
        SortMating = Parameter4[para4]
        toxicity = c(0,Toxicity[3])
        resistancecost = Resistancecost[4]
        fecunditycost = Fecunditycost[3]
        initfreqchoice = Fc0[5]
        initfreqresistance = Fr0[1]
        recrate = Recombinationrate[9]
        recombinationrate = rep(recrate, numresistanceloci + numchoiceloci)
        
        Parameters = paste0("fc", fecunditycost, "_hDr", scaledensity, "_AM",SortMating, "_DomR",dominanceresistance[1], "C", dominancechoice[1],
                            "_rec", recombinationrate[1], "_rc", resistancecost, "_fr",round(initfreqresistance, 2), "fch", round(initfreqchoice, 2),
                            "_tox", toxicity[2], "_lam", N_offspring, "_fxchoice", fixedChoice, "_Set",Set )
        
          outfilerawavoid = paste0(getwd(), "/",FFolder, "/", Parameters,"_", "Avoid", "_SimPar.RData")
          outfilerawnone = paste0(getwd(), "/",FFolder, "/", Parameters,"_", "None", "_SimPar.RData")
          
          # Reloading saved data
          out.choice = readRDS(outfilerawavoid)
          out.nochoice = readRDS(outfilerawnone)
          
          # - Extracting data for different variables
          resistance_tracking = out.choice$resistance_tracking
          choice_tracking = out.choice$choice_tracking
          resistance_tracking_nochoice = out.nochoice$resistance_tracking
          choice_tracking_nochoice = out.nochoice$choice_tracking
          resistance_popvariance = out.choice$resistance_popvariance
          choice_popvariance =  out.choice$choice_popvariance
          resistance_popvariance_nochoice = out.nochoice$resistance_popvariance
          choice_popvariance_nochoice = out.nochoice$choice_popvariance
          N1_nochoice = out.nochoice$N1
          N2_nochoice = out.nochoice$N2
          N1_choice = out.choice$N1
          N2_choice = out.choice$N2
        
        #- We calculate mean, std deviation, and std error in the traits for evolving and random choice behavior. Std deviation is not used in the plots ---
        resistancemean = apply(resistance_tracking[targetTtimes,],1,mean2)
        resistancesd = apply(resistance_tracking[targetTtimes,],1,FUN = function(x)sd(x, na.rm = TRUE))
        resistancestd = apply(resistance_tracking[targetTtimes,],1,FUN = function(x)std_error(x))
        resistancemean_nochoice = apply(resistance_tracking_nochoice[targetTtimes,],1,mean2)
        resistancesd_nochoice = apply(resistance_tracking_nochoice[targetTtimes,],1,FUN = function(x)sd(x, na.rm = TRUE))
        resistancestd_nochoice = apply(resistance_tracking_nochoice[targetTtimes,],1,FUN = function(x)std_error(x))
        choicemean = apply(choice_tracking[targetTtimes,],1,mean2)
        choicesd = apply(choice_tracking[targetTtimes,],1,FUN = function(x)sd(x, na.rm = TRUE))
        choicestd = apply(choice_tracking[targetTtimes,],1,FUN = function(x)std_error(x))
        choicemean_nochoice = apply(choice_tracking_nochoice[targetTtimes,],1,mean2)
        choicesd_nochoice = apply(choice_tracking_nochoice[targetTtimes,],1,FUN = function(x)sd(x, na.rm = TRUE))
        choicestd_nochoice = apply(choice_tracking_nochoice[targetTtimes,],1,FUN = function(x)std_error(x))


        resistance_popvarmean = apply(resistance_popvariance[targetTtimes,],1,mean2)
        resistance_popvarstd = apply(resistance_popvariance[targetTtimes,],1,FUN = function(x)std_error(x))
        resistance_popvarmean_nochoice = apply(resistance_popvariance_nochoice[targetTtimes,],1,mean2)
        resistance_popvarstd_nochoice = apply(resistance_popvariance_nochoice[targetTtimes,],1,FUN = function(x)std_error(x))
        choice_popvarmean = apply(choice_popvariance[targetTtimes,],1,mean2)
        choice_popvarstd = apply(choice_popvariance[targetTtimes,],1,FUN = function(x)std_error(x))
        choice_popvarmean_nochoice = apply(choice_popvariance_nochoice[targetTtimes,],1,mean2)
        choice_popvarstd_nochoice = apply(choice_popvariance_nochoice[targetTtimes,],1,FUN = function(x)std_error(x))


        Rresistancemean1[,para1,para2,para3,para4] = resistancemean
        Rresistancesd1[,para1,para2,para3,para4] = resistancesd
        Rresistancestd1[,para1,para2,para3,para4] = resistancestd
        Rresistancemean0[,para1,para2,para3,para4] = resistancemean_nochoice
        Rresistancesd0[,para1,para2,para3,para4] = resistancesd_nochoice
        Rresistancestd0[,para1,para2,para3,para4] = resistancestd_nochoice
        Cchoicemean1[,para1,para2,para3,para4] = choicemean
        Cchoicesd1[,para1,para2,para3,para4] = choicesd
        Cchoicestd1[,para1,para2,para3,para4] = choicestd
        Cchoicemean0[,para1,para2,para3,para4] = choicemean_nochoice
        Cchoicesd0[,para1,para2,para3,para4] = choicesd_nochoice
        Cchoicestd0[,para1,para2,para3,para4] = choicestd_nochoice

        # #----pop std mean + its std error stored
        Rresistance_popvariancemean1[,para1,para2,para3,para4] = resistance_popvarmean
        Rresistance_popvariancestd1[,para1,para2,para3,para4] = resistance_popvarstd
        Rresistance_popvariancemean0[,para1,para2,para3,para4] = resistance_popvarmean_nochoice
        Rresistance_popvariancestd0[,para1,para2,para3,para4] = resistance_popvarstd_nochoice
        Cchoice_popvariancemean1[,para1,para2,para3,para4] = choice_popvarmean
        Cchoice_popvariancestd1[,para1,para2,para3,para4] = choice_popvarstd
        Cchoice_popvariancemean0[,para1,para2,para3,para4] = choice_popvarmean_nochoice
        Cchoice_popvariancestd0[,para1,para2,para3,para4] = choice_popvarstd_nochoice

        # Population size at the respective plants ---------
        
        N1mean = apply(N1_choice[targetTtimes,],1,mean2)
        N1_std = apply(N1_choice[targetTtimes,],1,FUN = function(x)std_error(x))

        N1mean_nochoice = apply(N1_nochoice[targetTtimes,],1,mean2)
        N1_std_nochoice = apply(N1_nochoice[targetTtimes,],1,FUN = function(x)std_error(x))

        N2mean = apply(N2_choice[targetTtimes,],1,mean2)
        N2_std = apply(N2_choice[targetTtimes,],1,FUN = function(x)std_error(x))

        N2mean_nochoice = apply(N2_nochoice[targetTtimes,],1,mean2)
        N2_std_nochoice = apply(N2_nochoice[targetTtimes,],1,FUN = function(x)std_error(x))

        N1_mean1[,para1,para2,para3,para4] = N1mean
        N1_mean0[,para1,para2,para3,para4] = N1mean_nochoice
        N1_std1[,para1,para2,para3,para4] = N1_std
        N1_std0[,para1,para2,para3,para4] = N1_std_nochoice
        N2_mean1[,para1,para2,para3,para4] = N2mean
        N2_mean0[,para1,para2,para3,para4] = N2mean_nochoice
        N2_std1[,para1,para2,para3,para4] = N2_std
        N2_std0[,para1,para2,para3,para4] = N2_std_nochoice
        ##---------------------
        
        ##------------ Evolution rate: mean and std error ------------
        
        resT11 = diff(rowMeans(resistance_tracking[c(1:T50),], na.rm = TRUE))
        resT10 = diff(rowMeans(resistance_tracking_nochoice[c(1:T50),], na.rm = TRUE))
        choT11 = diff(rowMeans(choice_tracking[c(1:T50),], na.rm = TRUE))
        choT10 = diff(rowMeans(choice_tracking_nochoice[c(1:T50),], na.rm = TRUE))
        
        stdresT11 = apply(diff(resistance_tracking[c(1:T50),]),1,FUN = function(x)std_error(x))
        stdresT10 = apply(diff(resistance_tracking_nochoice[c(1:T50),]),1,FUN = function(x)std_error(x))
        stdchoT11 = apply(diff(choice_tracking[c(1:T50),]),1,FUN = function(x)std_error(x))
        stdchoT10 = apply(diff(choice_tracking_nochoice[c(1:T50),]),1,FUN = function(x)std_error(x))
        
        noabs_ResSpeed1[,para1,para2,para3,para4] = c(mean(resT11[1:T20], na.rm = TRUE), mean(resT11[1:T50], na.rm = TRUE))
        noabs_ResSpeed0[,para1,para2,para3,para4] = c(mean(resT10[1:T20], na.rm = TRUE), mean(resT10[1:T50], na.rm = TRUE))
        noabs_ChoSpeed1[,para1,para2,para3,para4] = c(mean(choT11[1:T20], na.rm = TRUE), mean(choT11[1:T50], na.rm = TRUE))
        noabs_ChoSpeed0[,para1,para2,para3,para4] = c(mean(choT10[1:T20], na.rm = TRUE), mean(choT10[1:T50], na.rm = TRUE))
        
        noabs_stdResSpeed1[,para1,para2,para3,para4] = c(mean(stdresT11[1:T20], na.rm = TRUE), mean(stdresT11[1:T50], na.rm = TRUE))
        noabs_stdResSpeed0[,para1,para2,para3,para4] = c(mean(stdresT10[1:T20], na.rm = TRUE), mean(stdresT10[1:T50], na.rm = TRUE))
        noabs_stdChoSpeed1[,para1,para2,para3,para4] = c(mean(stdchoT11[1:T20], na.rm = TRUE), mean(stdchoT11[1:T50], na.rm = TRUE))
        noabs_stdChoSpeed0[,para1,para2,para3,para4] = c(mean(stdchoT10[1:T20], na.rm = TRUE), mean(stdchoT10[1:T50], na.rm = TRUE))
        
        ResSpeed1[,para1,para2,para3,para4] = abs(c(mean(resT11[1:T20], na.rm = TRUE), mean(resT11[1:T50], na.rm = TRUE)))
        ResSpeed0[,para1,para2,para3,para4] = abs(c(mean(resT10[1:T20], na.rm = TRUE), mean(resT10[1:T50], na.rm = TRUE)))
        ChoSpeed1[,para1,para2,para3,para4] = abs(c(mean(choT11[1:T20], na.rm = TRUE), mean(choT11[1:T50], na.rm = TRUE)))
        ChoSpeed0[,para1,para2,para3,para4] = abs(c(mean(choT10[1:T20], na.rm = TRUE), mean(choT10[1:T50], na.rm = TRUE)))
        
      }}}}


#-Because it takes long to get analysed data, its better to save them in a folder so that one does not have to re-run the above analysis again but just upload the saved data and proceed to plotting. 
DataAnalysis = "AnalysedData"

if(!dir.exists(paste0(getwd(), "/",Folder, "/", DataAnalysis))){
  dir.create(paste0(getwd(), "/",Folder, "/", DataAnalysis))
}

saveRDS(Rresistancemean1, file=paste0(getwd(),"/", Folder,"/", DataAnalysis, "/", "Rresistancemean1.RData"))
saveRDS(Rresistancesd1, file=paste0(getwd(),"/", Folder,"/", DataAnalysis, "/","Rresistancesd1.RData"))
saveRDS(Rresistancestd1, file=paste0(getwd(),"/", Folder,"/", DataAnalysis, "/","Rresistancestd1.RData"))
saveRDS(Cchoicemean1, file=paste0(getwd(),"/", Folder,"/", DataAnalysis, "/","Cchoicemean1.RData"))
saveRDS(Cchoicesd1, file=paste0(getwd(),"/", Folder,"/", DataAnalysis, "/","Cchoicesd1.RData"))
saveRDS(Cchoicestd1, file=paste0(getwd(),"/", Folder,"/", DataAnalysis, "/","Cchoicestd1.RData"))
saveRDS(Rresistancemean0, file=paste0(getwd(),"/", Folder,"/", DataAnalysis, "/", "Rresistancemean0.RData"))
saveRDS(Rresistancesd0, file=paste0(getwd(),"/", Folder,"/", DataAnalysis, "/","Rresistancesd0.RData"))
saveRDS(Rresistancestd0, file=paste0(getwd(),"/", Folder,"/", DataAnalysis, "/","Rresistancestd0.RData"))
saveRDS(Cchoicemean0, file=paste0(getwd(),"/", Folder,"/", DataAnalysis, "/","Cchoicemean0.RData"))
saveRDS(Cchoicesd0, file=paste0(getwd(),"/", Folder,"/", DataAnalysis, "/","Cchoicesd0.RData"))
saveRDS(Cchoicestd0, file=paste0(getwd(),"/", Folder,"/", DataAnalysis, "/","Cchoicestd0.RData"))

saveRDS(Rresistance_popvariancemean1, file=paste0(getwd(),"/", Folder,"/", DataAnalysis, "/", "Rresistance_popvariancemean1.RData"))
saveRDS(Rresistance_popvariancestd1, file=paste0(getwd(),"/", Folder,"/", DataAnalysis, "/","Rresistance_popvariancestd1.RData"))
saveRDS(Cchoice_popvariancemean1, file=paste0(getwd(),"/", Folder,"/", DataAnalysis, "/","Cchoice_popvariancemean1.RData"))
saveRDS(Cchoice_popvariancestd1, file=paste0(getwd(),"/", Folder,"/", DataAnalysis, "/","Cchoice_popvariancestd1.RData"))
saveRDS(Rresistance_popvariancemean0, file=paste0(getwd(),"/", Folder,"/", DataAnalysis, "/", "Rresistance_popvariancemean0.RData"))
saveRDS(Rresistance_popvariancestd0, file=paste0(getwd(),"/", Folder,"/", DataAnalysis, "/","Rresistance_popvariancestd0.RData"))
saveRDS(Cchoice_popvariancemean0, file=paste0(getwd(),"/", Folder,"/", DataAnalysis, "/","Cchoice_popvariancemean0.RData"))
saveRDS(Cchoice_popvariancestd0, file=paste0(getwd(),"/", Folder,"/", DataAnalysis, "/","Cchoice_popvariancestd0.RData"))


saveRDS(N1_mean1, file=paste0(getwd(),"/", Folder,"/", DataAnalysis, "/", "N1_mean1.RData"))
saveRDS(N1_mean0, file=paste0(getwd(),"/", Folder,"/", DataAnalysis, "/","N1_mean0.RData"))
saveRDS(N1_std1, file=paste0(getwd(),"/", Folder,"/", DataAnalysis, "/","N1_std1.RData"))
saveRDS(N1_std0, file=paste0(getwd(),"/", Folder,"/", DataAnalysis, "/","N1_std0.RData"))
saveRDS(N2_mean1, file=paste0(getwd(),"/", Folder,"/", DataAnalysis, "/", "N2_mean1.RData"))
saveRDS(N2_mean0, file=paste0(getwd(),"/", Folder,"/", DataAnalysis, "/","N2_mean0.RData"))
saveRDS(N2_std1, file=paste0(getwd(),"/", Folder,"/", DataAnalysis, "/","N2_std1.RData"))
saveRDS(N2_std0, file=paste0(getwd(),"/", Folder,"/", DataAnalysis, "/","N2_std0.RData"))


saveRDS(noabs_stdResSpeed0, file=paste0(getwd(),"/", Folder,"/", DataAnalysis, "/", "noabs_stdResSpeed0.RData"))
saveRDS(noabs_stdResSpeed1, file=paste0(getwd(),"/", Folder,"/", DataAnalysis, "/","noabs_stdResSpeed1.RData"))
saveRDS(noabs_stdChoSpeed0, file=paste0(getwd(),"/", Folder,"/", DataAnalysis, "/","noabs_stdChoSpeed0.RData"))
saveRDS(noabs_stdChoSpeed1, file=paste0(getwd(),"/", Folder,"/", DataAnalysis, "/","noabs_stdChoSpeed1.RData"))

saveRDS(noabs_ResSpeed0, file=paste0(getwd(),"/", Folder,"/", DataAnalysis, "/", "noabs_ResSpeed0.RData"))
saveRDS(noabs_ResSpeed1, file=paste0(getwd(),"/", Folder,"/", DataAnalysis, "/","noabs_ResSpeed1.RData"))
saveRDS(noabs_ChoSpeed0, file=paste0(getwd(),"/", Folder,"/", DataAnalysis, "/","noabs_ChoSpeed0.RData"))
saveRDS(noabs_ChoSpeed1, file=paste0(getwd(),"/", Folder,"/", DataAnalysis, "/","noabs_ChoSpeed1.RData"))

saveRDS(ResSpeed0, file=paste0(getwd(),"/", Folder,"/", DataAnalysis, "/", "ResSpeed0.RData"))
saveRDS(ResSpeed1, file=paste0(getwd(),"/", Folder,"/", DataAnalysis, "/","ResSpeed1.RData"))
saveRDS(ChoSpeed0, file=paste0(getwd(),"/", Folder,"/", DataAnalysis, "/","ChoSpeed0.RData"))
saveRDS(ChoSpeed1, file=paste0(getwd(),"/", Folder,"/", DataAnalysis, "/","ChoSpeed1.RData"))


#-------------------------------------- Illustration for costs, recombination rates, and toxicity -------------------------------------------

Folder = "Costs_Recombinationrate_Toxicity" # Folder for storing simulated data
FFolder = Folder

ParameterCombination = 2# Assign different numbers for different parameter combinations to help distinguish the names of plots
Parameter1 = Fecunditycost
Parameter2 = Resistancecost
Parameter3 = Toxicity
Parameter4 = Recombinationrate


#-------------------
noabs_stdResSpeed1 = noabs_stdResSpeed0 = noabs_stdChoSpeed1 = noabs_stdChoSpeed0 = noabs_ResSpeed1 = noabs_ResSpeed0 = noabs_ChoSpeed1 = noabs_ChoSpeed0= Rresistancestd1 = Rresistancestd0 = Cchoicestd1 = Cchoicestd0 = ResSpeed1 = ResSpeed0 = ChoSpeed1 = ChoSpeed0 = N1_mean1 = N1_mean0 = N1_std1 = N1_std0 = N2_mean1 = N2_mean0 = N2_std1 = N2_std0 = Rresistancemean1 = Rresistancesd1 = Rresistancemean0 = Rresistancesd0 = Cchoicemean1 = Cchoicesd1 = Cchoicemean0 = Cchoicesd0 = Rresistance_popvariancemean1 = Rresistance_popvariancestd1 = Rresistance_popvariancemean0 = Rresistance_popvariancestd0 = Cchoice_popvariancemean1 = Cchoice_popvariancestd1 = Cchoice_popvariancemean0 = Cchoice_popvariancestd0= array(NA, dim = c(length(targetTtimes), length(Parameter1), length(Parameter2), length(Parameter3), length(Parameter4)))

#--
for(para1 in 1:length(Parameter1)){
  for(para2 in 1:length(Parameter2)){
    for(para3 in 1:length(Parameter3)){
      for(para4 in 1:length(Parameter4)){
        
        hr = Dominanceresistance[4]
        hc = Dominancechoice[4]
        dominanceresistance = rep(hr, numresistanceloci);#dominance values for the resistance loci, should generally be between 0 and 1
        dominancechoice = rep(hc, numchoiceloci);#dominance values for the choice loci, should generally be between 0 an 1
        scaledensity = Scaledensity[3]
        SortMating = sortMating[5]
        toxicity = c(0,Parameter3[para3])
        resistancecost = Parameter[para2]
        fecunditycost = Parameter1[para1]
        initfreqchoice = Fc0[5]
        initfreqresistance = Fr0[1]
        recrate = Parameter4[para4]
        recombinationrate = rep(recrate, numresistanceloci + numchoiceloci)
        
        Parameters = paste0("fc", fecunditycost, "_hDr", scaledensity, "_AM",SortMating, "_DomR",dominanceresistance[1], "C", dominancechoice[1],
                            "_rec", recombinationrate[1], "_rc", resistancecost, "_fr",round(initfreqresistance, 2), "fch", round(initfreqchoice, 2),
                            "_tox", toxicity[2], "_lam", N_offspring, "_fxchoice", fixedChoice, "_Set",Set )
        
        outfilerawavoid = paste0(getwd(), "/",FFolder, "/", Parameters,"_", "Avoid", "_SimPar.RData")
        outfilerawnone = paste0(getwd(), "/",FFolder, "/", Parameters,"_", "None", "_SimPar.RData")
        
        # Reloading saved data
        out.choice = readRDS(outfilerawavoid)
        out.nochoice = readRDS(outfilerawnone)
        
        # - Extracting data for different variables
        resistance_tracking = out.choice$resistance_tracking
        choice_tracking = out.choice$choice_tracking
        resistance_tracking_nochoice = out.nochoice$resistance_tracking
        choice_tracking_nochoice = out.nochoice$choice_tracking
        resistance_popvariance = out.choice$resistance_popvariance
        choice_popvariance =  out.choice$choice_popvariance
        resistance_popvariance_nochoice = out.nochoice$resistance_popvariance
        choice_popvariance_nochoice = out.nochoice$choice_popvariance
        N1_nochoice = out.nochoice$N1
        N2_nochoice = out.nochoice$N2
        N1_choice = out.choice$N1
        N2_choice = out.choice$N2
        
        #- We calculate mean, std deviation, and std error in the traits for evolving and random choice behavior. Std deviation is not used in the plots ---
        resistancemean = apply(resistance_tracking[targetTtimes,],1,mean2)
        resistancesd = apply(resistance_tracking[targetTtimes,],1,FUN = function(x)sd(x, na.rm = TRUE))
        resistancestd = apply(resistance_tracking[targetTtimes,],1,FUN = function(x)std_error(x))
        resistancemean_nochoice = apply(resistance_tracking_nochoice[targetTtimes,],1,mean2)
        resistancesd_nochoice = apply(resistance_tracking_nochoice[targetTtimes,],1,FUN = function(x)sd(x, na.rm = TRUE))
        resistancestd_nochoice = apply(resistance_tracking_nochoice[targetTtimes,],1,FUN = function(x)std_error(x))
        choicemean = apply(choice_tracking[targetTtimes,],1,mean2)
        choicesd = apply(choice_tracking[targetTtimes,],1,FUN = function(x)sd(x, na.rm = TRUE))
        choicestd = apply(choice_tracking[targetTtimes,],1,FUN = function(x)std_error(x))
        choicemean_nochoice = apply(choice_tracking_nochoice[targetTtimes,],1,mean2)
        choicesd_nochoice = apply(choice_tracking_nochoice[targetTtimes,],1,FUN = function(x)sd(x, na.rm = TRUE))
        choicestd_nochoice = apply(choice_tracking_nochoice[targetTtimes,],1,FUN = function(x)std_error(x))
        
        
        resistance_popvarmean = apply(resistance_popvariance[targetTtimes,],1,mean2)
        resistance_popvarstd = apply(resistance_popvariance[targetTtimes,],1,FUN = function(x)std_error(x))
        resistance_popvarmean_nochoice = apply(resistance_popvariance_nochoice[targetTtimes,],1,mean2)
        resistance_popvarstd_nochoice = apply(resistance_popvariance_nochoice[targetTtimes,],1,FUN = function(x)std_error(x))
        choice_popvarmean = apply(choice_popvariance[targetTtimes,],1,mean2)
        choice_popvarstd = apply(choice_popvariance[targetTtimes,],1,FUN = function(x)std_error(x))
        choice_popvarmean_nochoice = apply(choice_popvariance_nochoice[targetTtimes,],1,mean2)
        choice_popvarstd_nochoice = apply(choice_popvariance_nochoice[targetTtimes,],1,FUN = function(x)std_error(x))
        
        
        Rresistancemean1[,para1,para2,para3,para4] = resistancemean
        Rresistancesd1[,para1,para2,para3,para4] = resistancesd
        Rresistancestd1[,para1,para2,para3,para4] = resistancestd
        Rresistancemean0[,para1,para2,para3,para4] = resistancemean_nochoice
        Rresistancesd0[,para1,para2,para3,para4] = resistancesd_nochoice
        Rresistancestd0[,para1,para2,para3,para4] = resistancestd_nochoice
        Cchoicemean1[,para1,para2,para3,para4] = choicemean
        Cchoicesd1[,para1,para2,para3,para4] = choicesd
        Cchoicestd1[,para1,para2,para3,para4] = choicestd
        Cchoicemean0[,para1,para2,para3,para4] = choicemean_nochoice
        Cchoicesd0[,para1,para2,para3,para4] = choicesd_nochoice
        Cchoicestd0[,para1,para2,para3,para4] = choicestd_nochoice
        
        # #----pop std mean + its std error stored
        Rresistance_popvariancemean1[,para1,para2,para3,para4] = resistance_popvarmean
        Rresistance_popvariancestd1[,para1,para2,para3,para4] = resistance_popvarstd
        Rresistance_popvariancemean0[,para1,para2,para3,para4] = resistance_popvarmean_nochoice
        Rresistance_popvariancestd0[,para1,para2,para3,para4] = resistance_popvarstd_nochoice
        Cchoice_popvariancemean1[,para1,para2,para3,para4] = choice_popvarmean
        Cchoice_popvariancestd1[,para1,para2,para3,para4] = choice_popvarstd
        Cchoice_popvariancemean0[,para1,para2,para3,para4] = choice_popvarmean_nochoice
        Cchoice_popvariancestd0[,para1,para2,para3,para4] = choice_popvarstd_nochoice
        
        # Population size at the respective plants ---------
        
        N1mean = apply(N1_choice[targetTtimes,],1,mean2)
        N1_std = apply(N1_choice[targetTtimes,],1,FUN = function(x)std_error(x))
        
        N1mean_nochoice = apply(N1_nochoice[targetTtimes,],1,mean2)
        N1_std_nochoice = apply(N1_nochoice[targetTtimes,],1,FUN = function(x)std_error(x))
        
        N2mean = apply(N2_choice[targetTtimes,],1,mean2)
        N2_std = apply(N2_choice[targetTtimes,],1,FUN = function(x)std_error(x))
        
        N2mean_nochoice = apply(N2_nochoice[targetTtimes,],1,mean2)
        N2_std_nochoice = apply(N2_nochoice[targetTtimes,],1,FUN = function(x)std_error(x))
        
        N1_mean1[,para1,para2,para3,para4] = N1mean
        N1_mean0[,para1,para2,para3,para4] = N1mean_nochoice
        N1_std1[,para1,para2,para3,para4] = N1_std
        N1_std0[,para1,para2,para3,para4] = N1_std_nochoice
        N2_mean1[,para1,para2,para3,para4] = N2mean
        N2_mean0[,para1,para2,para3,para4] = N2mean_nochoice
        N2_std1[,para1,para2,para3,para4] = N2_std
        N2_std0[,para1,para2,para3,para4] = N2_std_nochoice
        ##---------------------
        
        ##------------ Evolution rate: mean and std error ------------
        
        resT11 = diff(rowMeans(resistance_tracking[c(1:T50),], na.rm = TRUE))
        resT10 = diff(rowMeans(resistance_tracking_nochoice[c(1:T50),], na.rm = TRUE))
        choT11 = diff(rowMeans(choice_tracking[c(1:T50),], na.rm = TRUE))
        choT10 = diff(rowMeans(choice_tracking_nochoice[c(1:T50),], na.rm = TRUE))
        
        stdresT11 = apply(diff(resistance_tracking[c(1:T50),]),1,FUN = function(x)std_error(x))
        stdresT10 = apply(diff(resistance_tracking_nochoice[c(1:T50),]),1,FUN = function(x)std_error(x))
        stdchoT11 = apply(diff(choice_tracking[c(1:T50),]),1,FUN = function(x)std_error(x))
        stdchoT10 = apply(diff(choice_tracking_nochoice[c(1:T50),]),1,FUN = function(x)std_error(x))
        
        noabs_ResSpeed1[,para1,para2,para3,para4] = c(mean(resT11[1:T20], na.rm = TRUE), mean(resT11[1:T50], na.rm = TRUE))
        noabs_ResSpeed0[,para1,para2,para3,para4] = c(mean(resT10[1:T20], na.rm = TRUE), mean(resT10[1:T50], na.rm = TRUE))
        noabs_ChoSpeed1[,para1,para2,para3,para4] = c(mean(choT11[1:T20], na.rm = TRUE), mean(choT11[1:T50], na.rm = TRUE))
        noabs_ChoSpeed0[,para1,para2,para3,para4] = c(mean(choT10[1:T20], na.rm = TRUE), mean(choT10[1:T50], na.rm = TRUE))
        
        noabs_stdResSpeed1[,para1,para2,para3,para4] = c(mean(stdresT11[1:T20], na.rm = TRUE), mean(stdresT11[1:T50], na.rm = TRUE))
        noabs_stdResSpeed0[,para1,para2,para3,para4] = c(mean(stdresT10[1:T20], na.rm = TRUE), mean(stdresT10[1:T50], na.rm = TRUE))
        noabs_stdChoSpeed1[,para1,para2,para3,para4] = c(mean(stdchoT11[1:T20], na.rm = TRUE), mean(stdchoT11[1:T50], na.rm = TRUE))
        noabs_stdChoSpeed0[,para1,para2,para3,para4] = c(mean(stdchoT10[1:T20], na.rm = TRUE), mean(stdchoT10[1:T50], na.rm = TRUE))
        
        ResSpeed1[,para1,para2,para3,para4] = abs(c(mean(resT11[1:T20], na.rm = TRUE), mean(resT11[1:T50], na.rm = TRUE)))
        ResSpeed0[,para1,para2,para3,para4] = abs(c(mean(resT10[1:T20], na.rm = TRUE), mean(resT10[1:T50], na.rm = TRUE)))
        ChoSpeed1[,para1,para2,para3,para4] = abs(c(mean(choT11[1:T20], na.rm = TRUE), mean(choT11[1:T50], na.rm = TRUE)))
        ChoSpeed0[,para1,para2,para3,para4] = abs(c(mean(choT10[1:T20], na.rm = TRUE), mean(choT10[1:T50], na.rm = TRUE)))
        
      }}}}


#-Because it takes long to get analysed data, its better to save them in a folder so that one does not have to re-run the above analysis again but just upload the saved data and proceed to plotting. 
DataAnalysis = "AnalysedData"

if(!dir.exists(paste0(getwd(), "/",Folder, "/", DataAnalysis))){
  dir.create(paste0(getwd(), "/",Folder, "/", DataAnalysis))
}

saveRDS(Rresistancemean1, file=paste0(getwd(),"/", Folder,"/", DataAnalysis, "/", "Rresistancemean1.RData"))
saveRDS(Rresistancesd1, file=paste0(getwd(),"/", Folder,"/", DataAnalysis, "/","Rresistancesd1.RData"))
saveRDS(Rresistancestd1, file=paste0(getwd(),"/", Folder,"/", DataAnalysis, "/","Rresistancestd1.RData"))
saveRDS(Cchoicemean1, file=paste0(getwd(),"/", Folder,"/", DataAnalysis, "/","Cchoicemean1.RData"))
saveRDS(Cchoicesd1, file=paste0(getwd(),"/", Folder,"/", DataAnalysis, "/","Cchoicesd1.RData"))
saveRDS(Cchoicestd1, file=paste0(getwd(),"/", Folder,"/", DataAnalysis, "/","Cchoicestd1.RData"))
saveRDS(Rresistancemean0, file=paste0(getwd(),"/", Folder,"/", DataAnalysis, "/", "Rresistancemean0.RData"))
saveRDS(Rresistancesd0, file=paste0(getwd(),"/", Folder,"/", DataAnalysis, "/","Rresistancesd0.RData"))
saveRDS(Rresistancestd0, file=paste0(getwd(),"/", Folder,"/", DataAnalysis, "/","Rresistancestd0.RData"))
saveRDS(Cchoicemean0, file=paste0(getwd(),"/", Folder,"/", DataAnalysis, "/","Cchoicemean0.RData"))
saveRDS(Cchoicesd0, file=paste0(getwd(),"/", Folder,"/", DataAnalysis, "/","Cchoicesd0.RData"))
saveRDS(Cchoicestd0, file=paste0(getwd(),"/", Folder,"/", DataAnalysis, "/","Cchoicestd0.RData"))

saveRDS(Rresistance_popvariancemean1, file=paste0(getwd(),"/", Folder,"/", DataAnalysis, "/", "Rresistance_popvariancemean1.RData"))
saveRDS(Rresistance_popvariancestd1, file=paste0(getwd(),"/", Folder,"/", DataAnalysis, "/","Rresistance_popvariancestd1.RData"))
saveRDS(Cchoice_popvariancemean1, file=paste0(getwd(),"/", Folder,"/", DataAnalysis, "/","Cchoice_popvariancemean1.RData"))
saveRDS(Cchoice_popvariancestd1, file=paste0(getwd(),"/", Folder,"/", DataAnalysis, "/","Cchoice_popvariancestd1.RData"))
saveRDS(Rresistance_popvariancemean0, file=paste0(getwd(),"/", Folder,"/", DataAnalysis, "/", "Rresistance_popvariancemean0.RData"))
saveRDS(Rresistance_popvariancestd0, file=paste0(getwd(),"/", Folder,"/", DataAnalysis, "/","Rresistance_popvariancestd0.RData"))
saveRDS(Cchoice_popvariancemean0, file=paste0(getwd(),"/", Folder,"/", DataAnalysis, "/","Cchoice_popvariancemean0.RData"))
saveRDS(Cchoice_popvariancestd0, file=paste0(getwd(),"/", Folder,"/", DataAnalysis, "/","Cchoice_popvariancestd0.RData"))


saveRDS(N1_mean1, file=paste0(getwd(),"/", Folder,"/", DataAnalysis, "/", "N1_mean1.RData"))
saveRDS(N1_mean0, file=paste0(getwd(),"/", Folder,"/", DataAnalysis, "/","N1_mean0.RData"))
saveRDS(N1_std1, file=paste0(getwd(),"/", Folder,"/", DataAnalysis, "/","N1_std1.RData"))
saveRDS(N1_std0, file=paste0(getwd(),"/", Folder,"/", DataAnalysis, "/","N1_std0.RData"))
saveRDS(N2_mean1, file=paste0(getwd(),"/", Folder,"/", DataAnalysis, "/", "N2_mean1.RData"))
saveRDS(N2_mean0, file=paste0(getwd(),"/", Folder,"/", DataAnalysis, "/","N2_mean0.RData"))
saveRDS(N2_std1, file=paste0(getwd(),"/", Folder,"/", DataAnalysis, "/","N2_std1.RData"))
saveRDS(N2_std0, file=paste0(getwd(),"/", Folder,"/", DataAnalysis, "/","N2_std0.RData"))


saveRDS(noabs_stdResSpeed0, file=paste0(getwd(),"/", Folder,"/", DataAnalysis, "/", "noabs_stdResSpeed0.RData"))
saveRDS(noabs_stdResSpeed1, file=paste0(getwd(),"/", Folder,"/", DataAnalysis, "/","noabs_stdResSpeed1.RData"))
saveRDS(noabs_stdChoSpeed0, file=paste0(getwd(),"/", Folder,"/", DataAnalysis, "/","noabs_stdChoSpeed0.RData"))
saveRDS(noabs_stdChoSpeed1, file=paste0(getwd(),"/", Folder,"/", DataAnalysis, "/","noabs_stdChoSpeed1.RData"))

saveRDS(noabs_ResSpeed0, file=paste0(getwd(),"/", Folder,"/", DataAnalysis, "/", "noabs_ResSpeed0.RData"))
saveRDS(noabs_ResSpeed1, file=paste0(getwd(),"/", Folder,"/", DataAnalysis, "/","noabs_ResSpeed1.RData"))
saveRDS(noabs_ChoSpeed0, file=paste0(getwd(),"/", Folder,"/", DataAnalysis, "/","noabs_ChoSpeed0.RData"))
saveRDS(noabs_ChoSpeed1, file=paste0(getwd(),"/", Folder,"/", DataAnalysis, "/","noabs_ChoSpeed1.RData"))

saveRDS(ResSpeed0, file=paste0(getwd(),"/", Folder,"/", DataAnalysis, "/", "ResSpeed0.RData"))
saveRDS(ResSpeed1, file=paste0(getwd(),"/", Folder,"/", DataAnalysis, "/","ResSpeed1.RData"))
saveRDS(ChoSpeed0, file=paste0(getwd(),"/", Folder,"/", DataAnalysis, "/","ChoSpeed0.RData"))
saveRDS(ChoSpeed1, file=paste0(getwd(),"/", Folder,"/", DataAnalysis, "/","ChoSpeed1.RData"))


#-------------------------------------- Illustration for costs and Initial Allele frequencies

Folder = "Costs_Allelefrequency" # Folder for storing simulated data
FFolder = Folder

ParameterCombination = 3# Assign different numbers for different parameter combinations to help distinguish the names of plots
Parameter1 = Fecunditycost
Parameter2 = Resistancecost
Parameter3 = Fr0
Parameter4 = Fc0


#-------------------
noabs_stdResSpeed1 = noabs_stdResSpeed0 = noabs_stdChoSpeed1 = noabs_stdChoSpeed0 = noabs_ResSpeed1 = noabs_ResSpeed0 = noabs_ChoSpeed1 = noabs_ChoSpeed0= Rresistancestd1 = Rresistancestd0 = Cchoicestd1 = Cchoicestd0 = ResSpeed1 = ResSpeed0 = ChoSpeed1 = ChoSpeed0 = N1_mean1 = N1_mean0 = N1_std1 = N1_std0 = N2_mean1 = N2_mean0 = N2_std1 = N2_std0 = Rresistancemean1 = Rresistancesd1 = Rresistancemean0 = Rresistancesd0 = Cchoicemean1 = Cchoicesd1 = Cchoicemean0 = Cchoicesd0 = Rresistance_popvariancemean1 = Rresistance_popvariancestd1 = Rresistance_popvariancemean0 = Rresistance_popvariancestd0 = Cchoice_popvariancemean1 = Cchoice_popvariancestd1 = Cchoice_popvariancemean0 = Cchoice_popvariancestd0= array(NA, dim = c(length(targetTtimes), length(Parameter1), length(Parameter2), length(Parameter3), length(Parameter4)))

#--
for(para1 in 1:length(Parameter1)){
  for(para2 in 1:length(Parameter2)){
    for(para3 in 1:length(Parameter3)){
      for(para4 in 1:length(Parameter4)){
        
        hr = Dominanceresistance[4]
        hc = Dominancechoice[4]
        dominanceresistance = rep(hr, numresistanceloci);#dominance values for the resistance loci, should generally be between 0 and 1
        dominancechoice = rep(hc, numchoiceloci);#dominance values for the choice loci, should generally be between 0 an 1
        scaledensity = Scaledensity[3]
        SortMating = sortMating[5]
        toxicity = c(0,Toxicity[3])
        resistancecost = Parameter[para2]
        fecunditycost = Parameter1[para1]
        initfreqchoice = Parameter4[para4]
        initfreqresistance = Parameter3[para3]
        recrate = Recombinationrate[9]
        recombinationrate = rep(recrate, numresistanceloci + numchoiceloci)
        
        Parameters = paste0("fc", fecunditycost, "_hDr", scaledensity, "_AM",SortMating, "_DomR",dominanceresistance[1], "C", dominancechoice[1],
                            "_rec", recombinationrate[1], "_rc", resistancecost, "_fr",round(initfreqresistance, 2), "fch", round(initfreqchoice, 2),
                            "_tox", toxicity[2], "_lam", N_offspring, "_fxchoice", fixedChoice, "_Set",Set )
        
        outfilerawavoid = paste0(getwd(), "/",FFolder, "/", Parameters,"_", "Avoid", "_SimPar.RData")
        outfilerawnone = paste0(getwd(), "/",FFolder, "/", Parameters,"_", "None", "_SimPar.RData")
        
        # Reloading saved data
        out.choice = readRDS(outfilerawavoid)
        out.nochoice = readRDS(outfilerawnone)
        
        # - Extracting data for different variables
        resistance_tracking = out.choice$resistance_tracking
        choice_tracking = out.choice$choice_tracking
        resistance_tracking_nochoice = out.nochoice$resistance_tracking
        choice_tracking_nochoice = out.nochoice$choice_tracking
        resistance_popvariance = out.choice$resistance_popvariance
        choice_popvariance =  out.choice$choice_popvariance
        resistance_popvariance_nochoice = out.nochoice$resistance_popvariance
        choice_popvariance_nochoice = out.nochoice$choice_popvariance
        N1_nochoice = out.nochoice$N1
        N2_nochoice = out.nochoice$N2
        N1_choice = out.choice$N1
        N2_choice = out.choice$N2
        
        #- We calculate mean, std deviation, and std error in the traits for evolving and random choice behavior. Std deviation is not used in the plots ---
        resistancemean = apply(resistance_tracking[targetTtimes,],1,mean2)
        resistancesd = apply(resistance_tracking[targetTtimes,],1,FUN = function(x)sd(x, na.rm = TRUE))
        resistancestd = apply(resistance_tracking[targetTtimes,],1,FUN = function(x)std_error(x))
        resistancemean_nochoice = apply(resistance_tracking_nochoice[targetTtimes,],1,mean2)
        resistancesd_nochoice = apply(resistance_tracking_nochoice[targetTtimes,],1,FUN = function(x)sd(x, na.rm = TRUE))
        resistancestd_nochoice = apply(resistance_tracking_nochoice[targetTtimes,],1,FUN = function(x)std_error(x))
        choicemean = apply(choice_tracking[targetTtimes,],1,mean2)
        choicesd = apply(choice_tracking[targetTtimes,],1,FUN = function(x)sd(x, na.rm = TRUE))
        choicestd = apply(choice_tracking[targetTtimes,],1,FUN = function(x)std_error(x))
        choicemean_nochoice = apply(choice_tracking_nochoice[targetTtimes,],1,mean2)
        choicesd_nochoice = apply(choice_tracking_nochoice[targetTtimes,],1,FUN = function(x)sd(x, na.rm = TRUE))
        choicestd_nochoice = apply(choice_tracking_nochoice[targetTtimes,],1,FUN = function(x)std_error(x))
        
        
        resistance_popvarmean = apply(resistance_popvariance[targetTtimes,],1,mean2)
        resistance_popvarstd = apply(resistance_popvariance[targetTtimes,],1,FUN = function(x)std_error(x))
        resistance_popvarmean_nochoice = apply(resistance_popvariance_nochoice[targetTtimes,],1,mean2)
        resistance_popvarstd_nochoice = apply(resistance_popvariance_nochoice[targetTtimes,],1,FUN = function(x)std_error(x))
        choice_popvarmean = apply(choice_popvariance[targetTtimes,],1,mean2)
        choice_popvarstd = apply(choice_popvariance[targetTtimes,],1,FUN = function(x)std_error(x))
        choice_popvarmean_nochoice = apply(choice_popvariance_nochoice[targetTtimes,],1,mean2)
        choice_popvarstd_nochoice = apply(choice_popvariance_nochoice[targetTtimes,],1,FUN = function(x)std_error(x))
        
        
        Rresistancemean1[,para1,para2,para3,para4] = resistancemean
        Rresistancesd1[,para1,para2,para3,para4] = resistancesd
        Rresistancestd1[,para1,para2,para3,para4] = resistancestd
        Rresistancemean0[,para1,para2,para3,para4] = resistancemean_nochoice
        Rresistancesd0[,para1,para2,para3,para4] = resistancesd_nochoice
        Rresistancestd0[,para1,para2,para3,para4] = resistancestd_nochoice
        Cchoicemean1[,para1,para2,para3,para4] = choicemean
        Cchoicesd1[,para1,para2,para3,para4] = choicesd
        Cchoicestd1[,para1,para2,para3,para4] = choicestd
        Cchoicemean0[,para1,para2,para3,para4] = choicemean_nochoice
        Cchoicesd0[,para1,para2,para3,para4] = choicesd_nochoice
        Cchoicestd0[,para1,para2,para3,para4] = choicestd_nochoice
        
        # #----pop std mean + its std error stored
        Rresistance_popvariancemean1[,para1,para2,para3,para4] = resistance_popvarmean
        Rresistance_popvariancestd1[,para1,para2,para3,para4] = resistance_popvarstd
        Rresistance_popvariancemean0[,para1,para2,para3,para4] = resistance_popvarmean_nochoice
        Rresistance_popvariancestd0[,para1,para2,para3,para4] = resistance_popvarstd_nochoice
        Cchoice_popvariancemean1[,para1,para2,para3,para4] = choice_popvarmean
        Cchoice_popvariancestd1[,para1,para2,para3,para4] = choice_popvarstd
        Cchoice_popvariancemean0[,para1,para2,para3,para4] = choice_popvarmean_nochoice
        Cchoice_popvariancestd0[,para1,para2,para3,para4] = choice_popvarstd_nochoice
        
        # Population size at the respective plants ---------
        
        N1mean = apply(N1_choice[targetTtimes,],1,mean2)
        N1_std = apply(N1_choice[targetTtimes,],1,FUN = function(x)std_error(x))
        
        N1mean_nochoice = apply(N1_nochoice[targetTtimes,],1,mean2)
        N1_std_nochoice = apply(N1_nochoice[targetTtimes,],1,FUN = function(x)std_error(x))
        
        N2mean = apply(N2_choice[targetTtimes,],1,mean2)
        N2_std = apply(N2_choice[targetTtimes,],1,FUN = function(x)std_error(x))
        
        N2mean_nochoice = apply(N2_nochoice[targetTtimes,],1,mean2)
        N2_std_nochoice = apply(N2_nochoice[targetTtimes,],1,FUN = function(x)std_error(x))
        
        N1_mean1[,para1,para2,para3,para4] = N1mean
        N1_mean0[,para1,para2,para3,para4] = N1mean_nochoice
        N1_std1[,para1,para2,para3,para4] = N1_std
        N1_std0[,para1,para2,para3,para4] = N1_std_nochoice
        N2_mean1[,para1,para2,para3,para4] = N2mean
        N2_mean0[,para1,para2,para3,para4] = N2mean_nochoice
        N2_std1[,para1,para2,para3,para4] = N2_std
        N2_std0[,para1,para2,para3,para4] = N2_std_nochoice
        ##---------------------
        
        ##------------ Evolution rate: mean and std error ------------
        
        resT11 = diff(rowMeans(resistance_tracking[c(1:T50),], na.rm = TRUE))
        resT10 = diff(rowMeans(resistance_tracking_nochoice[c(1:T50),], na.rm = TRUE))
        choT11 = diff(rowMeans(choice_tracking[c(1:T50),], na.rm = TRUE))
        choT10 = diff(rowMeans(choice_tracking_nochoice[c(1:T50),], na.rm = TRUE))
        
        stdresT11 = apply(diff(resistance_tracking[c(1:T50),]),1,FUN = function(x)std_error(x))
        stdresT10 = apply(diff(resistance_tracking_nochoice[c(1:T50),]),1,FUN = function(x)std_error(x))
        stdchoT11 = apply(diff(choice_tracking[c(1:T50),]),1,FUN = function(x)std_error(x))
        stdchoT10 = apply(diff(choice_tracking_nochoice[c(1:T50),]),1,FUN = function(x)std_error(x))
        
        noabs_ResSpeed1[,para1,para2,para3,para4] = c(mean(resT11[1:T20], na.rm = TRUE), mean(resT11[1:T50], na.rm = TRUE))
        noabs_ResSpeed0[,para1,para2,para3,para4] = c(mean(resT10[1:T20], na.rm = TRUE), mean(resT10[1:T50], na.rm = TRUE))
        noabs_ChoSpeed1[,para1,para2,para3,para4] = c(mean(choT11[1:T20], na.rm = TRUE), mean(choT11[1:T50], na.rm = TRUE))
        noabs_ChoSpeed0[,para1,para2,para3,para4] = c(mean(choT10[1:T20], na.rm = TRUE), mean(choT10[1:T50], na.rm = TRUE))
        
        noabs_stdResSpeed1[,para1,para2,para3,para4] = c(mean(stdresT11[1:T20], na.rm = TRUE), mean(stdresT11[1:T50], na.rm = TRUE))
        noabs_stdResSpeed0[,para1,para2,para3,para4] = c(mean(stdresT10[1:T20], na.rm = TRUE), mean(stdresT10[1:T50], na.rm = TRUE))
        noabs_stdChoSpeed1[,para1,para2,para3,para4] = c(mean(stdchoT11[1:T20], na.rm = TRUE), mean(stdchoT11[1:T50], na.rm = TRUE))
        noabs_stdChoSpeed0[,para1,para2,para3,para4] = c(mean(stdchoT10[1:T20], na.rm = TRUE), mean(stdchoT10[1:T50], na.rm = TRUE))
        
        ResSpeed1[,para1,para2,para3,para4] = abs(c(mean(resT11[1:T20], na.rm = TRUE), mean(resT11[1:T50], na.rm = TRUE)))
        ResSpeed0[,para1,para2,para3,para4] = abs(c(mean(resT10[1:T20], na.rm = TRUE), mean(resT10[1:T50], na.rm = TRUE)))
        ChoSpeed1[,para1,para2,para3,para4] = abs(c(mean(choT11[1:T20], na.rm = TRUE), mean(choT11[1:T50], na.rm = TRUE)))
        ChoSpeed0[,para1,para2,para3,para4] = abs(c(mean(choT10[1:T20], na.rm = TRUE), mean(choT10[1:T50], na.rm = TRUE)))
        
      }}}}


#-Because it takes long to get analysed data, its better to save them in a folder so that one does not have to re-run the above analysis again but just upload the saved data and proceed to plotting. 
DataAnalysis = "AnalysedData"

if(!dir.exists(paste0(getwd(), "/",Folder, "/", DataAnalysis))){
  dir.create(paste0(getwd(), "/",Folder, "/", DataAnalysis))
}

saveRDS(Rresistancemean1, file=paste0(getwd(),"/", Folder,"/", DataAnalysis, "/", "Rresistancemean1.RData"))
saveRDS(Rresistancesd1, file=paste0(getwd(),"/", Folder,"/", DataAnalysis, "/","Rresistancesd1.RData"))
saveRDS(Rresistancestd1, file=paste0(getwd(),"/", Folder,"/", DataAnalysis, "/","Rresistancestd1.RData"))
saveRDS(Cchoicemean1, file=paste0(getwd(),"/", Folder,"/", DataAnalysis, "/","Cchoicemean1.RData"))
saveRDS(Cchoicesd1, file=paste0(getwd(),"/", Folder,"/", DataAnalysis, "/","Cchoicesd1.RData"))
saveRDS(Cchoicestd1, file=paste0(getwd(),"/", Folder,"/", DataAnalysis, "/","Cchoicestd1.RData"))
saveRDS(Rresistancemean0, file=paste0(getwd(),"/", Folder,"/", DataAnalysis, "/", "Rresistancemean0.RData"))
saveRDS(Rresistancesd0, file=paste0(getwd(),"/", Folder,"/", DataAnalysis, "/","Rresistancesd0.RData"))
saveRDS(Rresistancestd0, file=paste0(getwd(),"/", Folder,"/", DataAnalysis, "/","Rresistancestd0.RData"))
saveRDS(Cchoicemean0, file=paste0(getwd(),"/", Folder,"/", DataAnalysis, "/","Cchoicemean0.RData"))
saveRDS(Cchoicesd0, file=paste0(getwd(),"/", Folder,"/", DataAnalysis, "/","Cchoicesd0.RData"))
saveRDS(Cchoicestd0, file=paste0(getwd(),"/", Folder,"/", DataAnalysis, "/","Cchoicestd0.RData"))

saveRDS(Rresistance_popvariancemean1, file=paste0(getwd(),"/", Folder,"/", DataAnalysis, "/", "Rresistance_popvariancemean1.RData"))
saveRDS(Rresistance_popvariancestd1, file=paste0(getwd(),"/", Folder,"/", DataAnalysis, "/","Rresistance_popvariancestd1.RData"))
saveRDS(Cchoice_popvariancemean1, file=paste0(getwd(),"/", Folder,"/", DataAnalysis, "/","Cchoice_popvariancemean1.RData"))
saveRDS(Cchoice_popvariancestd1, file=paste0(getwd(),"/", Folder,"/", DataAnalysis, "/","Cchoice_popvariancestd1.RData"))
saveRDS(Rresistance_popvariancemean0, file=paste0(getwd(),"/", Folder,"/", DataAnalysis, "/", "Rresistance_popvariancemean0.RData"))
saveRDS(Rresistance_popvariancestd0, file=paste0(getwd(),"/", Folder,"/", DataAnalysis, "/","Rresistance_popvariancestd0.RData"))
saveRDS(Cchoice_popvariancemean0, file=paste0(getwd(),"/", Folder,"/", DataAnalysis, "/","Cchoice_popvariancemean0.RData"))
saveRDS(Cchoice_popvariancestd0, file=paste0(getwd(),"/", Folder,"/", DataAnalysis, "/","Cchoice_popvariancestd0.RData"))


saveRDS(N1_mean1, file=paste0(getwd(),"/", Folder,"/", DataAnalysis, "/", "N1_mean1.RData"))
saveRDS(N1_mean0, file=paste0(getwd(),"/", Folder,"/", DataAnalysis, "/","N1_mean0.RData"))
saveRDS(N1_std1, file=paste0(getwd(),"/", Folder,"/", DataAnalysis, "/","N1_std1.RData"))
saveRDS(N1_std0, file=paste0(getwd(),"/", Folder,"/", DataAnalysis, "/","N1_std0.RData"))
saveRDS(N2_mean1, file=paste0(getwd(),"/", Folder,"/", DataAnalysis, "/", "N2_mean1.RData"))
saveRDS(N2_mean0, file=paste0(getwd(),"/", Folder,"/", DataAnalysis, "/","N2_mean0.RData"))
saveRDS(N2_std1, file=paste0(getwd(),"/", Folder,"/", DataAnalysis, "/","N2_std1.RData"))
saveRDS(N2_std0, file=paste0(getwd(),"/", Folder,"/", DataAnalysis, "/","N2_std0.RData"))


saveRDS(noabs_stdResSpeed0, file=paste0(getwd(),"/", Folder,"/", DataAnalysis, "/", "noabs_stdResSpeed0.RData"))
saveRDS(noabs_stdResSpeed1, file=paste0(getwd(),"/", Folder,"/", DataAnalysis, "/","noabs_stdResSpeed1.RData"))
saveRDS(noabs_stdChoSpeed0, file=paste0(getwd(),"/", Folder,"/", DataAnalysis, "/","noabs_stdChoSpeed0.RData"))
saveRDS(noabs_stdChoSpeed1, file=paste0(getwd(),"/", Folder,"/", DataAnalysis, "/","noabs_stdChoSpeed1.RData"))

saveRDS(noabs_ResSpeed0, file=paste0(getwd(),"/", Folder,"/", DataAnalysis, "/", "noabs_ResSpeed0.RData"))
saveRDS(noabs_ResSpeed1, file=paste0(getwd(),"/", Folder,"/", DataAnalysis, "/","noabs_ResSpeed1.RData"))
saveRDS(noabs_ChoSpeed0, file=paste0(getwd(),"/", Folder,"/", DataAnalysis, "/","noabs_ChoSpeed0.RData"))
saveRDS(noabs_ChoSpeed1, file=paste0(getwd(),"/", Folder,"/", DataAnalysis, "/","noabs_ChoSpeed1.RData"))

saveRDS(ResSpeed0, file=paste0(getwd(),"/", Folder,"/", DataAnalysis, "/", "ResSpeed0.RData"))
saveRDS(ResSpeed1, file=paste0(getwd(),"/", Folder,"/", DataAnalysis, "/","ResSpeed1.RData"))
saveRDS(ChoSpeed0, file=paste0(getwd(),"/", Folder,"/", DataAnalysis, "/","ChoSpeed0.RData"))
saveRDS(ChoSpeed1, file=paste0(getwd(),"/", Folder,"/", DataAnalysis, "/","ChoSpeed1.RData"))


#------------- Time series plots. Here, I have selected only a few parameter combinations as it takes a lot of time to pass through many generations --------------------

Folder = "Dominance_Density_AM" # Folder for storing simulated data
FFolder = Folder
ParameterCombination = "Series"
seriesDominanceresistance = c(0,0.5,1)
seriesDominancechoice = c(0,0.5,1)
seriesScaledensity = c(1,3)
seriessortMating = c(0,0.8)

seriesTtimes = c(1:numgenerations)


#-------------------
N1_mean1_timeseries = N1_mean0_timeseries = N1_std1_timeseries = N1_std0_timeseries = N2_mean1_timeseries = N2_mean0_timeseries = N2_std1_timeseries = N2_std0_timeseries = Rresistancemean1_timeseries = Rresistancestd1_timeseries = Rresistancemean0_timeseries = Rresistancestd0_timeseries = Cchoicemean1_timeseries = Cchoicestd1_timeseries = Cchoicemean0_timeseries = Cchoicestd0_timeseries = Rresistance_popvariancemean1_timeseries = Rresistance_popvariancestd1_timeseries = Rresistance_popvariancemean0_timeseries = Rresistance_popvariancestd0_timeseries = Cchoice_popvariancemean1_timeseries = Cchoice_popvariancestd1_timeseries = Cchoice_popvariancemean0_timeseries = Cchoice_popvariancestd0_timeseries = array(NA, dim = c(length(seriesTtimes), length(seriesDominanceresistance), length(seriesDominancechoice), length(seriessortMating), length(seriesScaledensity)))
#--

for(para3 in 1:length(seriesScaledensity)){
  for(para4 in 1:length(seriessortMating)){
    for(para1 in 1:length(seriesDominanceresistance)){
      for(para2 in 1:length(seriesDominancechoice)){
        hr = seriesDominanceresistance[para1]
        hc = seriesDominancechoice[para2]
        dominanceresistance = rep(hr, numresistanceloci);#dominance values for the resistance loci, should generally be between 0 and 1
        dominancechoice = rep(hc, numchoiceloci);#dominance values for the choice loci, should generally be between 0 an 1
        scaledensity = seriesScaledensity[para3]
        SortMating = seriessortMating[para4]
        toxicity = c(0,Toxicity[3])
        resistancecost = Resistancecost[4]
        fecunditycost = Fecunditycost[3]
        initfreqchoice = Fc0[5]
        initfreqresistance = Fr0[1]
        recrate = Recombinationrate[9]
        recombinationrate = rep(recrate, numresistanceloci + numchoiceloci)
        
        Parameters = paste0("fc", fecunditycost, "_hDr", scaledensity, "_AM",SortMating, "_DomR",dominanceresistance[1], "C", dominancechoice[1],
                            "_rec", recombinationrate[1], "_rc", resistancecost, "_fr",round(initfreqresistance, 2), "fch", round(initfreqchoice, 2),
                            "_tox", toxicity[2], "_lam", N_offspring, "_fxchoice", fixedChoice, "_Set",Set )
        
        
        
        outfilerawavoid = paste0(getwd(), "/",FFolder, "/", Parameters,"_", "Avoid", "_SimPar.RData")
        outfilerawnone = paste0(getwd(), "/",FFolder, "/", Parameters,"_", "None", "_SimPar.RData")
        # Reloading saved data
        out.choice = readRDS(outfilerawavoid)
        out.nochoice = readRDS(outfilerawnone)
        
        
        resistance_tracking = out.choice$resistance_tracking
        choice_tracking = out.choice$choice_tracking
        resistance_tracking_nochoice = out.nochoice$resistance_tracking
        choice_tracking_nochoice = out.nochoice$choice_tracking
        resistance_popvariance = out.choice$resistance_popvariance
        choice_popvariance = out.choice$choice_popvariance
        resistance_popvariance_nochoice = out.nochoice$resistance_popvariance
        choice_popvariance_nochoice = out.nochoice$choice_popvariance
        N1_nochoice = out.nochoice$N1
        N2_nochoice = out.nochoice$N2
        N1_choice = out.choice$N1
        N2_choice = out.choice$N2
        
        
        resistancemean = apply(resistance_tracking[seriesTtimes,],1,mean2)
        resistancestderror = apply(resistance_tracking[seriesTtimes,],1,FUN = function(x)std_error(x))
        resistancemean_nochoice = apply(resistance_tracking_nochoice[seriesTtimes,],1,mean2)
        resistancestderror_nochoice = apply(resistance_tracking_nochoice[seriesTtimes,],1,FUN = function(x)std_error(x))
        
        choicemean = apply(choice_tracking[seriesTtimes,],1,mean2)
        choicestderror = apply(choice_tracking[seriesTtimes,],1,FUN = function(x)std_error(x))
        choicemean_nochoice = apply(choice_tracking_nochoice[seriesTtimes,],1,mean2)
        choicestderror_nochoice = apply(choice_tracking_nochoice[seriesTtimes,],1,FUN = function(x)std_error(x))
        
        
        resistance_popvarmean = apply(resistance_popvariance[seriesTtimes,],1,mean2)
        resistance_popvarstd = apply(resistance_popvariance[seriesTtimes,],1,FUN = function(x)std_error(x))
        resistance_popvarmean_nochoice = apply(resistance_popvariance_nochoice[seriesTtimes,],1,mean2)
        resistance_popvarstd_nochoice = apply(resistance_popvariance_nochoice[seriesTtimes,],1,FUN = function(x)std_error(x))
        choice_popvarmean = apply(choice_popvariance[seriesTtimes,],1,mean2)
        choice_popvarstd = apply(choice_popvariance[seriesTtimes,],1,FUN = function(x)std_error(x))
        choice_popvarmean_nochoice = apply(choice_popvariance_nochoice[seriesTtimes,],1,mean2)
        choice_popvarstd_nochoice = apply(choice_popvariance_nochoice[seriesTtimes,],1,FUN = function(x)std_error(x))
        
        
        Rresistancemean1_timeseries[,para1,para2,para3,para4] = resistancemean
        Rresistancestd1_timeseries[,para1,para2,para3,para4] = resistancestderror
        Rresistancemean0_timeseries[,para1,para2,para3,para4] = resistancemean_nochoice
        Rresistancestd0_timeseries[,para1,para2,para3,para4] = resistancestderror_nochoice
        Cchoicemean1_timeseries[,para1,para2,para3,para4] = choicemean
        Cchoicestd1_timeseries[,para1,para2,para3,para4] = choicestderror
        Cchoicemean0_timeseries[,para1,para2,para3,para4] = choicemean_nochoice
        Cchoicestd0_timeseries[,para1,para2,para3,para4] = choicestderror_nochoice
        
        # #----pop sd stored
        Rresistance_popvariancemean1_timeseries[,para1,para2,para3,para4] = resistance_popvarmean
        Rresistance_popvariancestd1_timeseries[,para1,para2,para3,para4] = resistance_popvarstd
        Rresistance_popvariancemean0_timeseries[,para1,para2,para3,para4] = resistance_popvarmean_nochoice
        Rresistance_popvariancestd0_timeseries[,para1,para2,para3,para4] = resistance_popvarstd_nochoice
        Cchoice_popvariancemean1_timeseries[,para1,para2,para3,para4] = choice_popvarmean
        Cchoice_popvariancestd1_timeseries[,para1,para2,para3,para4] = choice_popvarstd
        Cchoice_popvariancemean0_timeseries[,para1,para2,para3,para4] = choice_popvarmean_nochoice
        Cchoice_popvariancestd0_timeseries[,para1,para2,para3,para4] = choice_popvarstd_nochoice
        
        
        #
        N1mean = apply(N1_choice[seriesTtimes,],1,mean2)
        N1_std = apply(N1_choice[seriesTtimes,],1,FUN = function(x)std_error(x))
        
        N1mean_nochoice = apply(N1_nochoice[seriesTtimes,],1,mean2)
        N1_std_nochoice = apply(N1_nochoice[seriesTtimes,],1,FUN = function(x)std_error(x))
        
        N2mean = apply(N2_choice[seriesTtimes,],1,mean2)
        N2_std = apply(N2_choice[seriesTtimes,],1,FUN = function(x)std_error(x))
        
        N2mean_nochoice = apply(N2_nochoice[seriesTtimes,],1,mean2)
        N2_std_nochoice = apply(N2_nochoice[seriesTtimes,],1,FUN = function(x)std_error(x))
        
        N1_mean1_timeseries[,para1,para2,para3,para4] = N1mean
        N1_mean0_timeseries[,para1,para2,para3,para4] = N1mean_nochoice
        N1_std1_timeseries[,para1,para2,para3,para4] = N1_std
        N1_std0_timeseries[,para1,para2,para3,para4] = N1_std_nochoice
        N2_mean1_timeseries[,para1,para2,para3,para4] = N2mean
        N2_mean0_timeseries[,para1,para2,para3,para4] = N2mean_nochoice
        N2_std1_timeseries[,para1,para2,para3,para4] = N2_std
        N2_std0_timeseries[,para1,para2,para3,para4] = N2_std_nochoice
        ##---------------------
        
        
      }}}}

DataAnalysis = "AnalysedData"

if(!dir.exists(paste0(getwd(), "/",Folder, "/", DataAnalysis))){
  dir.create(paste0(getwd(), "/",Folder, "/", DataAnalysis))
}


saveRDS(Rresistancemean1_timeseries, file=paste0(getwd(),"/", Folder,"/", DataAnalysis, "/", "Rresistancemean1_timeseries.RData"))
saveRDS(Rresistancestd1_timeseries, file=paste0(getwd(),"/", Folder,"/", DataAnalysis, "/","Rresistancestd1_timeseries.RData"))
saveRDS(Cchoicemean1_timeseries, file=paste0(getwd(),"/", Folder,"/", DataAnalysis, "/","Cchoicemean1_timeseries.RData"))
saveRDS(Cchoicestd1_timeseries, file=paste0(getwd(),"/", Folder,"/", DataAnalysis, "/","Cchoicestd1_timeseries.RData"))
saveRDS(Rresistancemean0_timeseries, file=paste0(getwd(),"/", Folder,"/", DataAnalysis, "/", "Rresistancemean0_timeseries.RData"))
saveRDS(Rresistancestd0_timeseries, file=paste0(getwd(),"/", Folder,"/", DataAnalysis, "/","Rresistancestd0_timeseries.RData"))
saveRDS(Cchoicemean0_timeseries, file=paste0(getwd(),"/", Folder,"/", DataAnalysis, "/","Cchoicemean0_timeseries.RData"))
saveRDS(Cchoicestd0_timeseries, file=paste0(getwd(),"/", Folder,"/", DataAnalysis, "/","Cchoicestd0_timeseries.RData"))

saveRDS(Rresistance_popvariancemean1_timeseries, file=paste0(getwd(),"/", Folder,"/", DataAnalysis, "/", "Rresistance_popvariancemean1_timeseries.RData"))
saveRDS(Rresistance_popvariancestd1_timeseries, file=paste0(getwd(),"/", Folder,"/", DataAnalysis, "/","Rresistance_popvariancestd1_timeseries.RData"))
saveRDS(Cchoice_popvariancemean1_timeseries, file=paste0(getwd(),"/", Folder,"/", DataAnalysis, "/","Cchoice_popvariancemean1_timeseries.RData"))
saveRDS(Cchoice_popvariancestd1_timeseries, file=paste0(getwd(),"/", Folder,"/", DataAnalysis, "/","Cchoice_popvariancestd1_timeseries.RData"))
saveRDS(Rresistance_popvariancemean0_timeseries, file=paste0(getwd(),"/", Folder,"/", DataAnalysis, "/", "Rresistance_popvariancemean0_timeseries.RData"))
saveRDS(Rresistance_popvariancestd0_timeseries, file=paste0(getwd(),"/", Folder,"/", DataAnalysis, "/","Rresistance_popvariancestd0_timeseries.RData"))
saveRDS(Cchoice_popvariancemean0_timeseries, file=paste0(getwd(),"/", Folder,"/", DataAnalysis, "/","Cchoice_popvariancemean0_timeseries.RData"))
saveRDS(Cchoice_popvariancestd0_timeseries, file=paste0(getwd(),"/", Folder,"/", DataAnalysis, "/","Cchoice_popvariancestd0_timeseries.RData"))


saveRDS(N1_mean1_timeseries, file=paste0(getwd(),"/", Folder,"/", DataAnalysis, "/", "N1_mean1_timeseries.RData"))
saveRDS(N1_mean0_timeseries, file=paste0(getwd(),"/", Folder,"/", DataAnalysis, "/","N1_mean0_timeseries.RData"))
saveRDS(N1_std1_timeseries, file=paste0(getwd(),"/", Folder,"/", DataAnalysis, "/","N1_std1_timeseries.RData"))
saveRDS(N1_std0_timeseries, file=paste0(getwd(),"/", Folder,"/", DataAnalysis, "/","N1_std0_timeseries.RData"))
saveRDS(N2_mean1_timeseries, file=paste0(getwd(),"/", Folder,"/", DataAnalysis, "/", "N2_mean1_timeseries.RData"))
saveRDS(N2_mean0_timeseries, file=paste0(getwd(),"/", Folder,"/", DataAnalysis, "/","N2_mean0_timeseries.RData"))
saveRDS(N2_std1_timeseries, file=paste0(getwd(),"/", Folder,"/", DataAnalysis, "/","N2_std1_timeseries.RData"))
saveRDS(N2_std0_timeseries, file=paste0(getwd(),"/", Folder,"/", DataAnalysis, "/","N2_std0_timeseries.RData"))


setwd(workingdirectory)
```
