## Supplementary material for "The role of evolving niche choice in herbivore adaptation to host plants": These are R scripts and data files used for generating results.: IBM_PlantHerbivore_SimulationPlotting_SV.R

```
#Make sure that at the end of running this script, you run the last line of this script to reset the working directory
#Here, we upload the saved analysed data for plotting. 
#The parameter setting should be the same as in the execution and analysis scripts
#We illustrate for 4 parameter combinations, which includes the time series. The start of a parameter combination starts by defining the folder name, "Folder", in which data is (to be) kept


WD = "PLANTHERBIVORECHOICE"
workingdirectory = getwd()
setwd(paste0(workingdirectory,"/",WD))


numgenerations=200; # Number of generations. Used 200 for simulations
nsim=5;# Number of replicates. For cluster, 1000 was used
numresistanceloci  = 1; #number of loci underlying resistance to plant toxin
numchoiceloci  = 1; #number of loci underlying host plant choice
## Rescaling is done within the function to have the sum of effect sizes sum to 1. 
resistanceeffectsizes = rep(0.5, numresistanceloci); #vector of effect sizes; length should correspond to the number of resistance loci; Converted within the function such that the effect sizes add up to 1
choiceeffectsizes = rep(0.5, numchoiceloci);#vector of effect sizes; length should correspond to the number of choice loci; Converted within the function such that the effect sizes add up to 1
K = 1000 # Carrying capacity
N0 = 20# Initial population size
N_offspring = 4# Mean number of offspring for a susceptible individual
environmentalsdresistance = 0.0 # Environmental noice on resistance trait # NOT USED and SET to 0
environmentalsdchoice = 0.0# Environmental noice on choice trait # NOT USED and SET to 0

#setwd("~/Dropbox/Bielefeld_University/Munster_Collaboration/SimulatedData") # Set working directory
Dominancechoice = c(0,0.1, 0.2,0.5,0.8,0.9, 1) # Dominance choice
Dominanceresistance = c(0,0.1,0.2,0.5,0.8,0.9,1) # Dominance resistance
sortMating = c(-1,-0.8,-0.5,-0.2,0,0.2,0.5,0.8,1) # Host mating preference 
Recombinationrate = c(0, 0.0001, 0.001,0.01, 0.1, 0.2, 0.3, 0.4, 0.5) # Recombination rate
Fecunditycost = c(-1, -0.5, 0, 0.5, 1) # resistance effect on fecundity
Resistancecost = c(-0.5, -0.4, -0.3, -0.2, -0.1, -0.01, 0, 0.001, 0.01, 0.1, 0.2, 0.3, 0.4, 0.5) # Resistance effect on viability
Toxicity = c(0.0, 0.2, 0.5, 0.8, 1 ) # Level of toxicity of the toxic plant
Fr0 = c(0.1,0.3, 0.5,0.7, 0.9)# Initial resistance allele frequency
Fc0 = c(0.1,0.3, 0.5,0.7, 0.9)# Initial choice allele frequency
Scaledensity = c(1, 2, 3, 4)# Habitat size
fixedChoice = 0.5 # random choice behaviour
T20 = 20 # Time to measure speed of evolution # Used 20 in the simulations
T50 = 50 # another time to measure speed of evolution # Used 50 in the simulations
tTspeed = c(T20,T50) 


targetTtimes = c(10,numgenerations)#c(1:numgenerations)
Set = 1# Unused parameter. Leave it as it is. 


Folder = "Dominance_Density_AM" # Folder for storing simulated data
FFolder = Folder

ParameterCombination = 1# Assign different numbers for different parameter combinations to help distinguish the names of plots
Parameter1 = Dominanceresistance
Parameter2 = Dominancechoice
Parameter3 = Scaledensity
Parameter4 = sortMating


#------ This folder is for storing plots
plotFolder = "Plots"

if(!dir.exists(paste0(getwd(), "/",Folder, "/", plotFolder))){
  dir.create(paste0(getwd(), "/",Folder, "/", plotFolder))
}

#--------- Loading already analysed data for plotting--------------
DataAnalysis = "AnalysedData"

noabs_stdResSpeed0= readRDS(paste0(getwd(),"/", Folder,"/", DataAnalysis, "/", "noabs_stdResSpeed0.RData"))
noabs_stdResSpeed1= readRDS(paste0(getwd(),"/", Folder,"/", DataAnalysis, "/","noabs_stdResSpeed1.RData"))
noabs_stdChoSpeed0= readRDS(paste0(getwd(),"/", Folder,"/", DataAnalysis, "/","noabs_stdChoSpeed0.RData"))
noabs_stdChoSpeed1= readRDS(paste0(getwd(),"/", Folder,"/", DataAnalysis, "/","noabs_stdChoSpeed1.RData"))

noabs_ResSpeed0= readRDS(paste0(getwd(),"/", Folder,"/", DataAnalysis, "/", "noabs_ResSpeed0.RData"))
noabs_ResSpeed1= readRDS(paste0(getwd(),"/", Folder,"/", DataAnalysis, "/","noabs_ResSpeed1.RData"))
noabs_ChoSpeed0= readRDS(paste0(getwd(),"/", Folder,"/", DataAnalysis, "/","noabs_ChoSpeed0.RData"))
noabs_ChoSpeed1= readRDS(paste0(getwd(),"/", Folder,"/", DataAnalysis, "/","noabs_ChoSpeed1.RData"))


ResSpeed0= readRDS(paste0(getwd(),"/", Folder,"/", DataAnalysis, "/", "ResSpeed0.RData"))
ResSpeed1= readRDS(paste0(getwd(),"/", Folder,"/", DataAnalysis, "/","ResSpeed1.RData"))
ChoSpeed0= readRDS(paste0(getwd(),"/", Folder,"/", DataAnalysis, "/","ChoSpeed0.RData"))
ChoSpeed1= readRDS(paste0(getwd(),"/", Folder,"/", DataAnalysis, "/","ChoSpeed1.RData"))

Rresistancemean1= readRDS(paste0(getwd(),"/", Folder,"/", DataAnalysis, "/", "Rresistancemean1.RData"))
Rresistancesd1 = readRDS(paste0(getwd(),"/", Folder,"/", DataAnalysis, "/","Rresistancesd1.RData"))
Rresistancestd1 = readRDS(paste0(getwd(),"/", Folder,"/", DataAnalysis, "/","Rresistancestd1.RData"))
Cchoicemean1= readRDS(paste0(getwd(),"/", Folder,"/", DataAnalysis, "/","Cchoicemean1.RData"))
Cchoicesd1= readRDS(paste0(getwd(),"/", Folder,"/", DataAnalysis, "/","Cchoicesd1.RData"))
Cchoicestd1= readRDS(paste0(getwd(),"/", Folder,"/", DataAnalysis, "/","Cchoicestd1.RData"))
Rresistancemean0 = readRDS(paste0(getwd(),"/", Folder,"/", DataAnalysis, "/", "Rresistancemean0.RData"))
Rresistancesd0 = readRDS(paste0(getwd(),"/", Folder,"/", DataAnalysis, "/","Rresistancesd0.RData"))
Rresistancestd0 = readRDS(paste0(getwd(),"/", Folder,"/", DataAnalysis, "/","Rresistancestd0.RData"))
Cchoicemean0= readRDS(paste0(getwd(),"/", Folder,"/", DataAnalysis, "/","Cchoicemean0.RData"))
Cchoicesd0= readRDS(paste0(getwd(),"/", Folder,"/", DataAnalysis, "/","Cchoicesd0.RData"))
Cchoicestd0= readRDS(paste0(getwd(),"/", Folder,"/", DataAnalysis, "/","Cchoicestd0.RData"))


Rresistance_popvariancemean1= readRDS(paste0(getwd(),"/", Folder,"/", DataAnalysis, "/", "Rresistance_popvariancemean1.RData"))
Rresistance_popvariancestd1 = readRDS(paste0(getwd(),"/", Folder,"/", DataAnalysis, "/","Rresistance_popvariancestd1.RData"))
Cchoice_popvariancemean1= readRDS(paste0(getwd(),"/", Folder,"/", DataAnalysis, "/","Cchoice_popvariancemean1.RData"))
Cchoice_popvariancestd1= readRDS(paste0(getwd(),"/", Folder,"/", DataAnalysis, "/","Cchoice_popvariancestd1.RData"))
Rresistance_popvariancemean0 = readRDS(paste0(getwd(),"/", Folder,"/", DataAnalysis, "/", "Rresistance_popvariancemean0.RData"))
Rresistance_popvariancestd0 = readRDS(paste0(getwd(),"/", Folder,"/", DataAnalysis, "/","Rresistance_popvariancestd0.RData"))
Cchoice_popvariancemean0= readRDS(paste0(getwd(),"/", Folder,"/", DataAnalysis, "/","Cchoice_popvariancemean0.RData"))
Cchoice_popvariancestd0= readRDS(paste0(getwd(),"/", Folder,"/", DataAnalysis, "/","Cchoice_popvariancestd0.RData"))

N1_mean1= readRDS(paste0(getwd(),"/", Folder,"/", DataAnalysis, "/", "N1_mean1.RData"))
N1_mean0= readRDS(paste0(getwd(),"/", Folder,"/", DataAnalysis, "/","N1_mean0.RData"))
N1_std1= readRDS(paste0(getwd(),"/", Folder,"/", DataAnalysis, "/","N1_std1.RData"))
N1_std0= readRDS(paste0(getwd(),"/", Folder,"/", DataAnalysis, "/","N1_std0.RData"))
N2_mean1= readRDS(paste0(getwd(),"/", Folder,"/", DataAnalysis, "/", "N2_mean1.RData"))
N2_mean0= readRDS(paste0(getwd(),"/", Folder,"/", DataAnalysis, "/","N2_mean0.RData"))
N2_std1= readRDS(paste0(getwd(),"/", Folder,"/", DataAnalysis, "/","N2_std1.RData"))
N2_std0= readRDS(paste0(getwd(),"/", Folder,"/", DataAnalysis, "/","N2_std0.RData"))


tttt = targetTtimes[length(targetTtimes)]
ttt = targetTtimes[1]
plottargetTtimes = c(ttt, tttt)


#------------------- Preference - Resistance Population replicates 

subFolder = "Resistance_Preference" # For storing mean traits and population trait variance plots-----------

if(!dir.exists(paste0(getwd(), "/",Folder,"/",plotFolder,"/",subFolder))){
  dir.create(paste0(getwd(), "/",Folder,"/",plotFolder,"/",subFolder))
}


tT = length(targetTtimes)
{
  #for(para3 in 1:length(Parameter3)){
  para3 = 0
  for(para4 in 1:length(Parameter4)){
    for(para1 in 1:length(Parameter1)){
      for(para2 in 1:length(Parameter2)){
        
        png(paste0(getwd(), "/",Folder,"/", plotFolder, "/",subFolder,"/meanTraits_Param1",para1, "_Param3", para3, "_Param4",para4, "_Param2", para2, "_paracomb",ParameterCombination,".png"))
        
        par(mar = c(5, 5, 4, 5) + 0.3)
        
        matplot(Parameter3,Cchoicemean1[tT,para1,para2,,para4],ylim = c(0,1),cex.axis = 2, type = "n", ylab= list("Mean trait", font = 3, cex = 2),xlab=list("Parameter 3", font = 3, cex = 2))
        lines(Parameter3,Cchoicemean1[tT,para1,para2,,para4],type = 'l',col="black", lty = 1, lwd = 3);
        lines(Parameter3,Cchoicemean0[tT,para1,para2,,para4],type = 'l',col="black", lty = 2, lwd = 3);
        
        #----------------- Resistance -----------------
        lines(Parameter3,Rresistancemean1[tT,para1,para2,,para4],type = 'l',col="red", lty = 1, lwd = 3);
        lines(Parameter3,Rresistancemean0[tT,para1,para2,,para4],type = 'l',col="red", lty = 2, lwd = 3);
        #------- Add std errors ----------------
        polygon(x=c(Parameter3, rev(Parameter3)), y=c(Rresistancemean1[tT,para1,para2,,para4]-Rresistancestd1[tT,para1,para2,,para4], rev(Rresistancemean1[tT,para1,para2,,para4]+Rresistancestd1[tT,para1,para2,,para4])), border=NA, col=adjustcolor("gray", alpha.f = 0.3))
        polygon(x=c(Parameter3, rev(Parameter3)), y=c(Rresistancemean0[tT,para1,para2,,para4]-Rresistancestd0[tT,para1,para2,,para4], rev(Rresistancemean0[tT,para1,para2,,para4]+Rresistancestd0[tT,para1,para2,,para4])), border=NA, col=adjustcolor("gray", alpha.f = 0.3))
        polygon(x=c(Parameter3, rev(Parameter3)), y=c(Cchoicemean1[tT,para1,para2,,para4]-Cchoicestd1[tT,para1,para2,,para4], rev(Cchoicemean1[tT,para1,para2,,para4]+Cchoicestd1[tT,para1,para2,,para4])), border=NA, col=adjustcolor("gray", alpha.f = 0.3))
        polygon(x=c(Parameter3, rev(Parameter3)), y=c(Cchoicemean0[tT,para1,para2,,para4]-Cchoicestd0[tT,para1,para2,,para4], rev(Cchoicemean0[tT,para1,para2,,para4]+Cchoicestd0[tT,para1,para2,,para4])), border=NA, col=adjustcolor("gray", alpha.f = 0.3))
        
        legend("bottomleft", legend = c("Trait", "Choice", "Resistance"), col = c(NA,"black", "red"), text.font = 3, lwd = 3, lty=c(1,1,1))
        legend("bottomright", legend = c("Behaviour", "Evolving", "Fixed"), col = c(NA,"black", "black"), text.font = 3, lwd = 3, lty=c(1,1,2))
        dev.off()        
      }}}#}
}


{
  for(para3 in 1:length(Parameter3)){
    #for(para4 in 1:length(Parameter4)){
    para4 = 0
    for(para1 in 1:length(Parameter1)){
      for(para2 in 1:length(Parameter2)){
        
        png(paste0(getwd(), "/",Folder,"/", plotFolder, "/",subFolder,"/meanTraits_Param1",para1, "_Param3", para3, "_Param4",para4, "_Param2", para2, "_paracomb",ParameterCombination,".png"))
        
        par(mar = c(5, 5, 4, 5) + 0.3)
        
        matplot(Parameter4,Cchoicemean1[tT,para1,para2,para3,],ylim = c(0,1),cex.axis = 2, type = "n", ylab= list("Mean trait", font = 3, cex = 2),xlab=list("Parameter 4", font = 3, cex = 2))
        lines(Parameter4,Cchoicemean1[tT,para1,para2,para3,],type = 'l',col="black", lty = 1, lwd = 3);
        lines(Parameter4,Cchoicemean0[tT,para1,para2,para3,],type = 'l',col="black", lty = 2, lwd = 3);
        
        #----------------- Resistance -----------------
        lines(Parameter4,Rresistancemean1[tT,para1,para2,para3,],type = 'l',col="red", lty = 1, lwd = 3);
        lines(Parameter4,Rresistancemean0[tT,para1,para2,para3,],type = 'l',col="red", lty = 2, lwd = 3);
        
        polygon(x=c(Parameter4, rev(Parameter4)), y=c(Rresistancemean1[tT,para1,para2,para3,]-Rresistancestd1[tT,para1,para2,para3,], rev(Rresistancemean1[tT,para1,para2,para3,]+Rresistancestd1[tT,para1,para2,para3,])), border=NA, col=adjustcolor("gray", alpha.f = 0.3))
        polygon(x=c(Parameter4, rev(Parameter4)), y=c(Rresistancemean0[tT,para1,para2,para3,]-Rresistancestd0[tT,para1,para2,para3,], rev(Rresistancemean0[tT,para1,para2,para3,]+Rresistancestd0[tT,para1,para2,para3,])), border=NA, col=adjustcolor("gray", alpha.f = 0.3))
        polygon(x=c(Parameter4, rev(Parameter4)), y=c(Cchoicemean1[tT,para1,para2,para3,]-Cchoicestd1[tT,para1,para2,para3,], rev(Cchoicemean1[tT,para1,para2,para3,]+Cchoicestd1[tT,para1,para2,para3,])), border=NA, col=adjustcolor("gray", alpha.f = 0.3))
        polygon(x=c(Parameter4, rev(Parameter4)), y=c(Cchoicemean0[tT,para1,para2,para3,]-Cchoicestd0[tT,para1,para2,para3,], rev(Cchoicemean0[tT,para1,para2,para3,]+Cchoicestd0[tT,para1,para2,para3,])), border=NA, col=adjustcolor("gray", alpha.f = 0.3))
        
        # legend("bottomleft", legend = c("Choice", "Resistance"), col = c("black", "red"), text.font = 3, lwd = 3, lty=c(1,1))
        # legend("bottomright", legend = c( "Evolving", "Fixed"), col = c("black"), text.font = 3, lwd = 3, lty=c(1,2))
        dev.off()        
      }}}#}
}


{
  for(para3 in 1:length(Parameter3)){
    for(para4 in 1:length(Parameter4)){
      #for(para1 in 1:length(Parameter1)){
      para1 = 0
      for(para2 in 1:length(Parameter2)){
        
        png(paste0(getwd(), "/",Folder,"/", plotFolder, "/",subFolder,"/meanTraits_Param1",para1, "_Param3", para3, "_Param4",para4, "_Param2", para2, "_paracomb",ParameterCombination,".png"))
        
        par(mar = c(5, 5, 4, 5) + 0.3)
        
        matplot(Parameter1,Cchoicemean1[tT,,para2,para3,para4],ylim = c(0,1),cex.axis = 2, type = "n", ylab= list("Mean trait", font = 3, cex = 2),xlab=list("Parameter 1", font = 3, cex = 2))
        lines(Parameter1,Cchoicemean1[tT,,para2,para3,para4],type = 'l',col="black", lty = 1, lwd = 3);
        lines(Parameter1,Cchoicemean0[tT,,para2,para3,para4],type = 'l',col="black", lty = 2, lwd = 3);
        
        #----------------- Resistance -----------------
        lines(Parameter1,Rresistancemean1[tT,,para2,para3,para4],type = 'l',col="red", lty = 1, lwd = 3);
        lines(Parameter1,Rresistancemean0[tT,,para2,para3,para4],type = 'l',col="red", lty = 2, lwd = 3);
        
        polygon(x=c(Parameter1, rev(Parameter1)), y=c(Rresistancemean1[tT,,para2,para3,para4]-Rresistancestd1[tT,,para2,para3,para4], rev(Rresistancemean1[tT,,para2,para3,para4]+Rresistancestd1[tT,,para2,para3,para4])), border=NA, col=adjustcolor("gray", alpha.f = 0.3))
        polygon(x=c(Parameter1, rev(Parameter1)), y=c(Rresistancemean0[tT,,para2,para3,para4]-Rresistancestd0[tT,,para2,para3,para4], rev(Rresistancemean0[tT,,para2,para3,para4]+Rresistancestd0[tT,,para2,para3,para4])), border=NA, col=adjustcolor("gray", alpha.f = 0.3))
        polygon(x=c(Parameter1, rev(Parameter1)), y=c(Cchoicemean1[tT,,para2,para3,para4]-Cchoicestd1[tT,,para2,para3,para4], rev(Cchoicemean1[tT,,para2,para3,para4]+Cchoicestd1[tT,,para2,para3,para4])), border=NA, col=adjustcolor("gray", alpha.f = 0.3))
        polygon(x=c(Parameter1, rev(Parameter1)), y=c(Cchoicemean0[tT,,para2,para3,para4]-Cchoicestd0[tT,,para2,para3,para4], rev(Cchoicemean0[tT,,para2,para3,para4]+Cchoicestd0[tT,,para2,para3,para4])), border=NA, col=adjustcolor("gray", alpha.f = 0.3))
        
        # legend("bottomleft", legend = c("Choice", "Resistance"), col = c("black", "red"), text.font = 3, lwd = 3, lty=c(1,1))
        # legend("bottomright", legend = c( "Evolving", "Fixed"), col = c("black"), text.font = 3, lwd = 3, lty=c(1,2))
        dev.off()        
      }}}#}
}


{
  for(para3 in 1:length(Parameter3)){
    for(para4 in 1:length(Parameter4)){
      for(para1 in 1:length(Parameter1)){
        #for(para2 in 1:length(Parameter2)){
        para2 = 0
        png(paste0(getwd(), "/",Folder,"/", plotFolder, "/",subFolder,"/meanTraits_Param1",para1, "_Param3", para3, "_Param4",para4, "_Param2", para2, "_paracomb",ParameterCombination,".png"))
        
        par(mar = c(5, 5, 4, 5) + 0.3)
        
        matplot(Parameter2,Cchoicemean1[tT,para1,,para3,para4],ylim = c(0,1),cex.axis = 2, type = "n", ylab= list("Mean trait", font = 3, cex = 2),xlab=list("Parameter 2", font = 3, cex = 2))
        lines(Parameter2,Cchoicemean1[tT,para1,,para3,para4],type = 'l',col="black", lty = 1, lwd = 3);
        lines(Parameter2,Cchoicemean0[tT,para1,,para3,para4],type = 'l',col="black", lty = 2, lwd = 3);
        
        #----------------- Resistance -----------------
        lines(Parameter2,Rresistancemean1[tT,para1,,para3,para4],type = 'l',col="red", lty = 1, lwd = 3);
        lines(Parameter2,Rresistancemean0[tT,para1,,para3,para4],type = 'l',col="red", lty = 2, lwd = 3);
        
        polygon(x=c(Parameter2, rev(Parameter2)), y=c(Rresistancemean1[tT,para1,,para3,para4]-Rresistancestd1[tT,para1,,para3,para4], rev(Rresistancemean1[tT,para1,,para3,para4]+Rresistancestd1[tT,para1,,para3,para4])), border=NA, col=adjustcolor("gray", alpha.f = 0.3))
        polygon(x=c(Parameter2, rev(Parameter2)), y=c(Rresistancemean0[tT,para1,,para3,para4]-Rresistancestd0[tT,para1,,para3,para4], rev(Rresistancemean0[tT,para1,,para3,para4]+Rresistancestd0[tT,para1,,para3,para4])), border=NA, col=adjustcolor("gray", alpha.f = 0.3))
        polygon(x=c(Parameter2, rev(Parameter2)), y=c(Cchoicemean1[tT,para1,,para3,para4]-Cchoicestd1[tT,para1,,para3,para4], rev(Cchoicemean1[tT,para1,,para3,para4]+Cchoicestd1[tT,para1,,para3,para4])), border=NA, col=adjustcolor("gray", alpha.f = 0.3))
        polygon(x=c(Parameter2, rev(Parameter2)), y=c(Cchoicemean0[tT,para1,,para3,para4]-Cchoicestd0[tT,para1,,para3,para4], rev(Cchoicemean0[tT,para1,,para3,para4]+Cchoicestd0[tT,para1,,para3,para4])), border=NA, col=adjustcolor("gray", alpha.f = 0.3))
        
        # legend("bottomleft", legend = c("Choice", "Resistance"), col = c("black", "red"), text.font = 3, lwd = 3, lty=c(1,1))
        # legend("bottomright", legend = c( "Evolving", "Fixed"), col = c("black"), text.font = 3, lwd = 3, lty=c(1,2))
        dev.off()        
      }}}#}
}


#----------------- mean trait Variance within populations --------------
Minvar = min(c(Cchoice_popvariancemean1,Cchoice_popvariancemean0, Rresistance_popvariancemean0, Rresistance_popvariancemean1), na.rm = T)
Maxvar = max(c(Cchoice_popvariancemean1,Cchoice_popvariancemean0, Rresistance_popvariancemean0, Rresistance_popvariancemean1), na.rm = T)
{
  #for(para3 in 1:length(Parameter3)){
  para3 = 0
  for(para4 in 1:length(Parameter4)){
    for(para1 in 1:length(Parameter1)){
      for(para2 in 1:length(Parameter2)){
        
        png(paste0(getwd(), "/",Folder,"/", plotFolder, "/",subFolder,"/VarianceTraits_Param1",para1, "_Param3", para3, "_Param4",para4, "_Param2", para2, "_paracomb",ParameterCombination,".png"))
        
        par(mar = c(5, 5, 4, 5) + 0.3)
        
        matplot(Parameter3,Cchoice_popvariancemean1[tT,para1,para2,,para4],ylim = c(Minvar,Maxvar),cex.axis = 2, type = "n", ylab= list("Population trait variance", font = 3, cex = 2),xlab=list("Parameter 3", font = 3, cex = 2))
        lines(Parameter3,Cchoice_popvariancemean1[tT,para1,para2,,para4],type = 'l',col="black", lty = 1, lwd = 3);
        lines(Parameter3,Cchoice_popvariancemean0[tT,para1,para2,,para4],type = 'l',col="black", lty = 2, lwd = 3);
        
        
        #----------------- Resistance -----------------
        lines(Parameter3,Rresistance_popvariancemean1[tT,para1,para2,,para4],type = 'l',col="red", lty = 1, lwd = 3);
        lines(Parameter3,Rresistance_popvariancemean0[tT,para1,para2,,para4],type = 'l',col="red", lty = 2, lwd = 3);
        
        polygon(x=c(Parameter3, rev(Parameter3)), y=c(Rresistance_popvariancemean1[tT,para1,para2,,para4]-Rresistance_popvariancestd1[tT,para1,para2,,para4], rev(Rresistance_popvariancemean1[tT,para1,para2,,para4]+Rresistance_popvariancestd1[tT,para1,para2,,para4])), border=NA, col=adjustcolor("gray", alpha.f = 0.3))
        polygon(x=c(Parameter3, rev(Parameter3)), y=c(Rresistance_popvariancemean0[tT,para1,para2,,para4]-Rresistance_popvariancestd0[tT,para1,para2,,para4], rev(Rresistance_popvariancemean0[tT,para1,para2,,para4]+Rresistance_popvariancestd0[tT,para1,para2,,para4])), border=NA, col=adjustcolor("gray", alpha.f = 0.3))
        polygon(x=c(Parameter3, rev(Parameter3)), y=c(Cchoice_popvariancemean1[tT,para1,para2,,para4]-Cchoice_popvariancestd1[tT,para1,para2,,para4], rev(Cchoice_popvariancemean1[tT,para1,para2,,para4]+Cchoice_popvariancestd1[tT,para1,para2,,para4])), border=NA, col=adjustcolor("gray", alpha.f = 0.3))
        polygon(x=c(Parameter3, rev(Parameter3)), y=c(Cchoice_popvariancemean0[tT,para1,para2,,para4]-Cchoice_popvariancestd0[tT,para1,para2,,para4], rev(Cchoice_popvariancemean0[tT,para1,para2,,para4]+Cchoice_popvariancestd0[tT,para1,para2,,para4])), border=NA, col=adjustcolor("gray", alpha.f = 0.3))
        
        
        legend("bottomleft", legend = c("Trait", "Choice", "Resistance"), col = c(NA,"black", "red"), text.font = 3, lwd = 3, lty=c(1,1,1))
        legend("bottomright", legend = c("Behaviour", "Evolving", "Fixed"), col = c(NA,"black", "black"), text.font = 3, lwd = 3, lty=c(1,1,2))
        dev.off()        
      }}}#}
}


{
  for(para3 in 1:length(Parameter3)){
    #for(para4 in 1:length(Parameter4)){
    para4 = 0
    for(para1 in 1:length(Parameter1)){
      for(para2 in 1:length(Parameter2)){
        
        png(paste0(getwd(), "/",Folder,"/", plotFolder, "/",subFolder,"/VarianceTraits_Param1",para1, "_Param3", para3, "_Param4",para4, "_Param2", para2, "_paracomb",ParameterCombination,".png"))
        
        par(mar = c(5, 5, 4, 5) + 0.3)
        
        matplot(Parameter4,Cchoice_popvariancemean1[tT,para1,para2,para3,],ylim = c(Minvar,Maxvar),cex.axis = 2, type = "n", ylab= list("Population trait variance", font = 3, cex = 2),xlab=list("Parameter 4", font = 3, cex = 2))
        lines(Parameter4,Cchoice_popvariancemean1[tT,para1,para2,para3,],type = 'l',col="black", lty = 1, lwd = 3);
        lines(Parameter4,Cchoice_popvariancemean0[tT,para1,para2,para3,],type = 'l',col="black", lty = 2, lwd = 3);
        
        #----------------- Resistance -----------------
        lines(Parameter4,Rresistance_popvariancemean1[tT,para1,para2,para3,],type = 'l',col="red", lty = 1, lwd = 3);
        lines(Parameter4,Rresistance_popvariancemean0[tT,para1,para2,para3,],type = 'l',col="red", lty = 2, lwd = 3);
        
        polygon(x=c(Parameter4, rev(Parameter4)), y=c(Rresistance_popvariancemean1[tT,para1,para2,para3,]-Rresistance_popvariancestd1[tT,para1,para2,para3,], rev(Rresistance_popvariancemean1[tT,para1,para2,para3,]+Rresistance_popvariancestd1[tT,para1,para2,para3,])), border=NA, col=adjustcolor("gray", alpha.f = 0.3))
        polygon(x=c(Parameter4, rev(Parameter4)), y=c(Rresistance_popvariancemean0[tT,para1,para2,para3,]-Rresistance_popvariancestd0[tT,para1,para2,para3,], rev(Rresistance_popvariancemean0[tT,para1,para2,para3,]+Rresistance_popvariancestd0[tT,para1,para2,para3,])), border=NA, col=adjustcolor("gray", alpha.f = 0.3))
        polygon(x=c(Parameter4, rev(Parameter4)), y=c(Cchoice_popvariancemean1[tT,para1,para2,para3,]-Cchoice_popvariancestd1[tT,para1,para2,para3,], rev(Cchoice_popvariancemean1[tT,para1,para2,para3,]+Cchoice_popvariancestd1[tT,para1,para2,para3,])), border=NA, col=adjustcolor("gray", alpha.f = 0.3))
        polygon(x=c(Parameter4, rev(Parameter4)), y=c(Cchoice_popvariancemean0[tT,para1,para2,para3,]-Cchoice_popvariancestd0[tT,para1,para2,para3,], rev(Cchoice_popvariancemean0[tT,para1,para2,para3,]+Cchoice_popvariancestd0[tT,para1,para2,para3,])), border=NA, col=adjustcolor("gray", alpha.f = 0.3))
        
        # legend("bottomleft", legend = c("Choice", "Resistance"), col = c("black", "red"), text.font = 3, lwd = 3, lty=c(1,1))
        # legend("bottomright", legend = c( "Evolving", "Fixed"), col = c("black"), text.font = 3, lwd = 3, lty=c(1,2))
        dev.off()        
      }}}#}
}


{
  for(para3 in 1:length(Parameter3)){
    for(para4 in 1:length(Parameter4)){
      #for(para1 in 1:length(Parameter1)){
      para1 = 0
      for(para2 in 1:length(Parameter2)){
        
        png(paste0(getwd(), "/",Folder,"/", plotFolder, "/",subFolder,"/VarianceTraits_Param1",para1, "_Param3", para3, "_Param4",para4, "_Param2", para2, "_paracomb",ParameterCombination,".png"))
        par(mar = c(5, 5, 4, 5) + 0.3)
        
        matplot(Parameter1,Cchoice_popvariancemean1[tT,,para2,para3,para4],ylim = c(Minvar,Maxvar),cex.axis = 2, type = "n", ylab= list("Population trait variance", font = 3, cex = 2),xlab=list("Parameter 1", font = 3, cex = 2))
        lines(Parameter1,Cchoice_popvariancemean1[tT,,para2,para3,para4],type = 'l',col="black", lty = 1, lwd = 3);
        lines(Parameter1,Cchoice_popvariancemean0[tT,,para2,para3,para4],type = 'l',col="black", lty = 2, lwd = 3);
        
        #----------------- Resistance -----------------
        lines(Parameter1,Rresistance_popvariancemean1[tT,,para2,para3,para4],type = 'l',col="red", lty = 1, lwd = 3);
        lines(Parameter1,Rresistance_popvariancemean0[tT,,para2,para3,para4],type = 'l',col="red", lty = 2, lwd = 3);
        
        polygon(x=c(Parameter1, rev(Parameter1)), y=c(Rresistance_popvariancemean1[tT,,para2,para3,para4]-Rresistance_popvariancestd1[tT,,para2,para3,para4], rev(Rresistance_popvariancemean1[tT,,para2,para3,para4]+Rresistance_popvariancestd1[tT,,para2,para3,para4])), border=NA, col=adjustcolor("gray", alpha.f = 0.3))
        polygon(x=c(Parameter1, rev(Parameter1)), y=c(Rresistance_popvariancemean0[tT,,para2,para3,para4]-Rresistance_popvariancestd0[tT,,para2,para3,para4], rev(Rresistance_popvariancemean0[tT,,para2,para3,para4]+Rresistance_popvariancestd0[tT,,para2,para3,para4])), border=NA, col=adjustcolor("gray", alpha.f = 0.3))
        polygon(x=c(Parameter1, rev(Parameter1)), y=c(Cchoice_popvariancemean1[tT,,para2,para3,para4]-Cchoice_popvariancestd1[tT,,para2,para3,para4], rev(Cchoice_popvariancemean1[tT,,para2,para3,para4]+Cchoice_popvariancestd1[tT,,para2,para3,para4])), border=NA, col=adjustcolor("gray", alpha.f = 0.3))
        polygon(x=c(Parameter1, rev(Parameter1)), y=c(Cchoice_popvariancemean0[tT,,para2,para3,para4]-Cchoice_popvariancestd0[tT,,para2,para3,para4], rev(Cchoice_popvariancemean0[tT,,para2,para3,para4]+Cchoice_popvariancestd0[tT,,para2,para3,para4])), border=NA, col=adjustcolor("gray", alpha.f = 0.3))
        
        # legend("bottomleft", legend = c("Choice", "Resistance"), col = c("black", "red"), text.font = 3, lwd = 3, lty=c(1,1))
        # legend("bottomright", legend = c( "Evolving", "Fixed"), col = c("black"), text.font = 3, lwd = 3, lty=c(1,2))
        dev.off()        
      }}}#}
}


{
  for(para3 in 1:length(Parameter3)){
    for(para4 in 1:length(Parameter4)){
      for(para1 in 1:length(Parameter1)){
        #for(para2 in 1:length(Parameter2)){
        para2 = 0
        png(paste0(getwd(), "/",Folder,"/", plotFolder, "/",subFolder,"/VarianceTraits_Param1",para1, "_Param3", para3, "_Param4",para4, "_Param2", para2, "_paracomb",ParameterCombination,".png"))
        
        par(mar = c(5, 5, 4, 5) + 0.3)
        
        matplot(Parameter2,Cchoice_popvariancemean1[tT,para1,,para3,para4],ylim = c(Minvar,Maxvar),cex.axis = 2, type = "n", ylab= list("Population trait variance", font = 3, cex = 2),xlab=list("Parameter 2", font = 3, cex = 2))
        lines(Parameter2,Cchoice_popvariancemean1[tT,para1,,para3,para4],type = 'l',col="black", lty = 1, lwd = 3);
        lines(Parameter2,Cchoice_popvariancemean0[tT,para1,,para3,para4],type = 'l',col="black", lty = 2, lwd = 3);
        
        #----------------- Resistance -----------------
        lines(Parameter2,Rresistance_popvariancemean1[tT,para1,,para3,para4],type = 'l',col="red", lty = 1, lwd = 3);
        lines(Parameter2,Rresistance_popvariancemean0[tT,para1,,para3,para4],type = 'l',col="red", lty = 2, lwd = 3);
        
        polygon(x=c(Parameter2, rev(Parameter2)), y=c(Rresistance_popvariancemean1[tT,para1,,para3,para4]-Rresistance_popvariancestd1[tT,para1,,para3,para4], rev(Rresistance_popvariancemean1[tT,para1,,para3,para4]+Rresistance_popvariancestd1[tT,para1,,para3,para4])), border=NA, col=adjustcolor("gray", alpha.f = 0.3))
        polygon(x=c(Parameter2, rev(Parameter2)), y=c(Rresistance_popvariancemean0[tT,para1,,para3,para4]-Rresistance_popvariancestd0[tT,para1,,para3,para4], rev(Rresistance_popvariancemean0[tT,para1,,para3,para4]+Rresistance_popvariancestd0[tT,para1,,para3,para4])), border=NA, col=adjustcolor("gray", alpha.f = 0.3))
        polygon(x=c(Parameter2, rev(Parameter2)), y=c(Cchoice_popvariancemean1[tT,para1,,para3,para4]-Cchoice_popvariancestd1[tT,para1,,para3,para4], rev(Cchoice_popvariancemean1[tT,para1,,para3,para4]+Cchoice_popvariancestd1[tT,para1,,para3,para4])), border=NA, col=adjustcolor("gray", alpha.f = 0.3))
        polygon(x=c(Parameter2, rev(Parameter2)), y=c(Cchoice_popvariancemean0[tT,para1,,para3,para4]-Cchoice_popvariancestd0[tT,para1,,para3,para4], rev(Cchoice_popvariancemean0[tT,para1,,para3,para4]+Cchoice_popvariancestd0[tT,para1,,para3,para4])), border=NA, col=adjustcolor("gray", alpha.f = 0.3))
        
        
        # legend("bottomleft", legend = c("Choice", "Resistance"), col = c("black", "red"), text.font = 3, lwd = 3, lty=c(1,1))
        # legend("bottomright", legend = c( "Evolving", "Fixed"), col = c("black"), text.font = 3, lwd = 3, lty=c(1,2))
        dev.off()        
      }}}#}
}

# ------------------- Evolution rate ----------
subFolder = "SpeedResistance_Preference"

if(!dir.exists(paste0(getwd(), "/",Folder, "/", plotFolder, "/", subFolder))){
  dir.create(paste0(getwd(), "/",Folder, "/", plotFolder, "/", subFolder))
}


#----------------- Speed of adaptation --------------
Minspeed = min(c(ChoSpeed1,ChoSpeed0, ResSpeed0, ResSpeed1), na.rm = T)
Maxspeed = max(c(ChoSpeed1,ChoSpeed0, ResSpeed0, ResSpeed1), na.rm = T)
{
  #for(para3 in 1:length(Parameter3)){
  para3 = 0
  for(para4 in 1:length(Parameter4)){
    for(para1 in 1:length(Parameter1)){
      for(para2 in 1:length(Parameter2)){
        for(spt in 1:length(tTspeed)){
          png(paste0(getwd(), "/",Folder,"/", plotFolder, "/",subFolder,"/abs_SpeedEvolutionTraits_Param1",para1, "_Param3", para3, "_Param4",para4, "_Param2", para2, "_gen",tTspeed[spt], "_paracomb",ParameterCombination,".png"))
          
          par(mar = c(5, 5, 4, 5) + 0.3)
          
          matplot(Parameter3,ChoSpeed1[spt,para1,para2,,para4],ylim = c(Minspeed,Maxspeed),cex.axis = 2, type = "n", ylab= list("Evolution rate", font = 3, cex = 2),xlab=list("Parameter 3", font = 3, cex = 2))
          lines(Parameter3,ChoSpeed1[spt,para1,para2,,para4],type = 'l',col="black", lty = 1, lwd = 3);
          lines(Parameter3,ChoSpeed0[spt,para1,para2,,para4],type = 'l',col="black", lty = 2, lwd = 3);
          
          #----------------- Resistance -----------------
          lines(Parameter3,ResSpeed1[spt,para1,para2,,para4],type = 'l',col="red", lty = 1, lwd = 3);
          lines(Parameter3,ResSpeed0[spt,para1,para2,,para4],type = 'l',col="red", lty = 2, lwd = 3);
          
          legend("bottomleft", legend = c("Trait", "Choice", "Resistance"), col = c(NA,"black", "red"), text.font = 3, lwd = 3, lty=c(1,1,1))
          legend("bottomright", legend = c("Behaviour", "Evolving", "Fixed"), col = c(NA,"black", "black"), text.font = 3, lwd = 3, lty=c(1,1,2))
          dev.off()        
        }}}}#}
}


{
  for(para3 in 1:length(Parameter3)){
    #for(para4 in 1:length(Parameter4)){
    para4 = 0
    for(para1 in 1:length(Parameter1)){
      for(para2 in 1:length(Parameter2)){
        for(spt in 1:length(tTspeed)){
          png(paste0(getwd(), "/",Folder,"/", plotFolder, "/",subFolder,"/abs_SpeedEvolutionTraits_Param1",para1, "_Param3", para3, "_Param4",para4, "_Param2", para2, "_gen",tTspeed[spt], "_paracomb",ParameterCombination,".png"))
          
          par(mar = c(5, 5, 4, 5) + 0.3)
          
          matplot(Parameter4,ChoSpeed1[spt,para1,para2,para3,],ylim = c(Minspeed,Maxspeed),cex.axis = 2, type = "n", ylab= list("Evolution rate", font = 3, cex = 2),xlab=list("Parameter 4", font = 3, cex = 2))
          lines(Parameter4,ChoSpeed1[spt,para1,para2,para3,],type = 'l',col="black", lty = 1, lwd = 3);
          lines(Parameter4,ChoSpeed0[spt,para1,para2,para3,],type = 'l',col="black", lty = 2, lwd = 3);
          
          #----------------- Resistance -----------------
          lines(Parameter4,ResSpeed1[spt,para1,para2,para3,],type = 'l',col="red", lty = 1, lwd = 3);
          lines(Parameter4,ResSpeed0[spt,para1,para2,para3,],type = 'l',col="red", lty = 2, lwd = 3);
          # legend("bottomleft", legend = c("Choice", "Resistance"), col = c("black", "red"), text.font = 3, lwd = 3, lty=c(1,1))
          # legend("bottomright", legend = c( "Evolving", "Fixed"), col = c("black"), text.font = 3, lwd = 3, lty=c(1,2))
          dev.off()        
        }}}}#}
}


{
  for(para3 in 1:length(Parameter3)){
    for(para4 in 1:length(Parameter4)){
      #for(para1 in 1:length(Parameter1)){
      para1 = 0
      for(para2 in 1:length(Parameter2)){
        for(spt in 1:length(tTspeed)){
          png(paste0(getwd(), "/",Folder,"/", plotFolder, "/",subFolder,"/abs_SpeedEvolutionTraits_Param1",para1, "_Param3", para3, "_Param4",para4, "_Param2", para2, "_gen",tTspeed[spt], "_paracomb",ParameterCombination,".png"))
          par(mar = c(5, 5, 4, 5) + 0.3)
          
          matplot(Parameter1,ChoSpeed1[spt,,para2,para3,para4],ylim = c(Minspeed,Maxspeed),cex.axis = 2, type = "n", ylab= list("Evolution rate", font = 3, cex = 2),xlab=list("Parameter 1", font = 3, cex = 2))
          lines(Parameter1,ChoSpeed1[spt,,para2,para3,para4],type = 'l',col="black", lty = 1, lwd = 3);
          lines(Parameter1,ChoSpeed0[spt,,para2,para3,para4],type = 'l',col="black", lty = 2, lwd = 3);
          
          #----------------- Resistance -----------------
          lines(Parameter1,ResSpeed1[spt,,para2,para3,para4],type = 'l',col="red", lty = 1, lwd = 3);
          lines(Parameter1,ResSpeed0[spt,,para2,para3,para4],type = 'l',col="red", lty = 2, lwd = 3);
          
          # legend("bottomleft", legend = c("Choice", "Resistance"), col = c("black", "red"), text.font = 3, lwd = 3, lty=c(1,1))
          # legend("bottomright", legend = c( "Evolving", "Fixed"), col = c("black"), text.font = 3, lwd = 3, lty=c(1,2))
          dev.off()        
        }}}}#}
}


{
  for(para3 in 1:length(Parameter3)){
    for(para4 in 1:length(Parameter4)){
      for(para1 in 1:length(Parameter1)){
        #for(para2 in 1:length(Parameter2)){
        for(spt in 1:length(tTspeed)){
          para2 = 0
          png(paste0(getwd(), "/",Folder,"/", plotFolder, "/",subFolder,"/abs_SpeedEvolutionTraits_Param1",para1, "_Param3", para3, "_Param4",para4, "_Param2", para2, "_gen",tTspeed[spt], "_paracomb",ParameterCombination,".png"))
          
          par(mar = c(5, 5, 4, 5) + 0.3)
          
          matplot(Parameter2,ChoSpeed1[spt,para1,,para3,para4],ylim = c(Minspeed,Maxspeed),cex.axis = 2, type = "n", ylab= list("Evolution rate", font = 3, cex = 2),xlab=list("Parameter 2", font = 3, cex = 2))
          lines(Parameter2,ChoSpeed1[spt,para1,,para3,para4],type = 'l',col="black", lty = 1, lwd = 3);
          lines(Parameter2,ChoSpeed0[spt,para1,,para3,para4],type = 'l',col="black", lty = 2, lwd = 3);
          
          #----------------- Resistance -----------------
          lines(Parameter2,ResSpeed1[spt,para1,,para3,para4],type = 'l',col="red", lty = 1, lwd = 3);
          lines(Parameter2,ResSpeed0[spt,para1,,para3,para4],type = 'l',col="red", lty = 2, lwd = 3);
          
          # legend("bottomleft", legend = c("Choice", "Resistance"), col = c("black", "red"), text.font = 3, lwd = 3, lty=c(1,1))
          # legend("bottomright", legend = c( "Evolving", "Fixed"), col = c("black"), text.font = 3, lwd = 3, lty=c(1,2))
          dev.off()        
        }}}}#}
}


#------------------- Speed of adaptation -  Not absolute values ---------------------------
Minspeed = min(c(noabs_ChoSpeed1,noabs_ChoSpeed0, noabs_ResSpeed0, noabs_ResSpeed1), na.rm = T)
Maxspeed = max(c(noabs_ChoSpeed1,noabs_ChoSpeed0, noabs_ResSpeed0, noabs_ResSpeed1), na.rm = T)
{
  #for(para3 in 1:length(Parameter3)){
  para3 = 0
  for(para4 in 1:length(Parameter4)){
    for(para1 in 1:length(Parameter1)){
      for(para2 in 1:length(Parameter2)){
        for(spt in 1:length(tTspeed)){
          png(paste0(getwd(), "/",Folder,"/", plotFolder, "/",subFolder,"/noabs_SpeedEvolutionTraits_Param1",para1, "_Param3", para3, "_Param4",para4, "_Param2", para2, "_gen",tTspeed[spt], "_paracomb",ParameterCombination,".png"))
          
          par(mar = c(5, 5, 4, 5) + 0.3)
          
          matplot(Parameter3,noabs_ChoSpeed1[spt,para1,para2,,para4],ylim = c(Minspeed,Maxspeed),cex.axis = 2, type = "n", ylab= list("Evolution rate", font = 3, cex = 2),xlab=list("Parameter 3", font = 3, cex = 2))
          lines(Parameter3,noabs_ChoSpeed1[spt,para1,para2,,para4],type = 'l',col="black", lty = 1, lwd = 3);
          lines(Parameter3,noabs_ChoSpeed0[spt,para1,para2,,para4],type = 'l',col="black", lty = 2, lwd = 3);
          
          #----------------- Resistance -----------------
          lines(Parameter3,noabs_ResSpeed1[spt,para1,para2,,para4],type = 'l',col="red", lty = 1, lwd = 3);
          lines(Parameter3,noabs_ResSpeed0[spt,para1,para2,,para4],type = 'l',col="red", lty = 2, lwd = 3);
          
          polygon(x=c(Parameter3, rev(Parameter3)), y=c(noabs_ResSpeed1[spt,para1,para2,,para4]-noabs_stdResSpeed1[spt,para1,para2,,para4], rev(noabs_ResSpeed1[spt,para1,para2,,para4]+noabs_stdResSpeed1[spt,para1,para2,,para4])), border=NA, col=adjustcolor("gray", alpha.f = 0.3))
          polygon(x=c(Parameter3, rev(Parameter3)), y=c(noabs_ResSpeed0[spt,para1,para2,,para4]-noabs_stdResSpeed0[spt,para1,para2,,para4], rev(noabs_ResSpeed0[spt,para1,para2,,para4]+noabs_stdResSpeed0[spt,para1,para2,,para4])), border=NA, col=adjustcolor("gray", alpha.f = 0.3))
          polygon(x=c(Parameter3, rev(Parameter3)), y=c(noabs_ChoSpeed1[spt,para1,para2,,para4]-noabs_stdChoSpeed1[spt,para1,para2,,para4], rev(noabs_ChoSpeed1[spt,para1,para2,,para4]+noabs_stdChoSpeed1[spt,para1,para2,,para4])), border=NA, col=adjustcolor("gray", alpha.f = 0.3))
          polygon(x=c(Parameter3, rev(Parameter3)), y=c(noabs_ChoSpeed0[spt,para1,para2,,para4]-noabs_stdChoSpeed0[spt,para1,para2,,para4], rev(noabs_ChoSpeed0[spt,para1,para2,,para4]+noabs_stdChoSpeed0[spt,para1,para2,,para4])), border=NA, col=adjustcolor("gray", alpha.f = 0.3))
          
          legend("bottomleft", legend = c("Trait", "Choice", "Resistance"), col = c(NA,"black", "red"), text.font = 3, lwd = 3, lty=c(1,1,1))
          legend("bottomright", legend = c("Behaviour", "Evolving", "Fixed"), col = c(NA,"black", "black"), text.font = 3, lwd = 3, lty=c(1,1,2))
          dev.off()        
        }}}}#}
}


{
  for(para3 in 1:length(Parameter3)){
    #for(para4 in 1:length(Parameter4)){
    para4 = 0
    for(para1 in 1:length(Parameter1)){
      for(para2 in 1:length(Parameter2)){
        for(spt in 1:length(tTspeed)){
          png(paste0(getwd(), "/",Folder,"/", plotFolder, "/",subFolder,"/noabs_SpeedEvolutionTraits_Param1",para1, "_Param3", para3, "_Param4",para4, "_Param2", para2, "_gen",tTspeed[spt], "_paracomb",ParameterCombination,".png"))
          
          par(mar = c(5, 5, 4, 5) + 0.3)
          
          matplot(Parameter4,noabs_ChoSpeed1[spt,para1,para2,para3,],ylim = c(Minspeed,Maxspeed),cex.axis = 2, type = "n", ylab= list("Evolution rate", font = 3, cex = 2),xlab=list("Parameter 4", font = 3, cex = 2))
          lines(Parameter4,noabs_ChoSpeed1[spt,para1,para2,para3,],type = 'l',col="black", lty = 1, lwd = 3);
          lines(Parameter4,noabs_ChoSpeed0[spt,para1,para2,para3,],type = 'l',col="black", lty = 2, lwd = 3);
          
          #----------------- Resistance -----------------
          lines(Parameter4,noabs_ResSpeed1[spt,para1,para2,para3,],type = 'l',col="red", lty = 1, lwd = 3);
          lines(Parameter4,noabs_ResSpeed0[spt,para1,para2,para3,],type = 'l',col="red", lty = 2, lwd = 3);
          
          polygon(x=c(Parameter4, rev(Parameter4)), y=c(noabs_ResSpeed1[spt,para1,para2,para3,]-noabs_stdResSpeed1[spt,para1,para2,para3,], rev(noabs_ResSpeed1[spt,para1,para2,para3,]+noabs_stdResSpeed1[spt,para1,para2,para3,])), border=NA, col=adjustcolor("gray", alpha.f = 0.3))
          polygon(x=c(Parameter4, rev(Parameter4)), y=c(noabs_ResSpeed0[spt,para1,para2,para3,]-noabs_stdResSpeed0[spt,para1,para2,para3,], rev(noabs_ResSpeed0[spt,para1,para2,para3,]+noabs_stdResSpeed0[spt,para1,para2,para3,])), border=NA, col=adjustcolor("gray", alpha.f = 0.3))
          polygon(x=c(Parameter4, rev(Parameter4)), y=c(noabs_ChoSpeed1[spt,para1,para2,para3,]-noabs_stdChoSpeed1[spt,para1,para2,para3,], rev(noabs_ChoSpeed1[spt,para1,para2,para3,]+noabs_stdChoSpeed1[spt,para1,para2,para3,])), border=NA, col=adjustcolor("gray", alpha.f = 0.3))
          polygon(x=c(Parameter4, rev(Parameter4)), y=c(noabs_ChoSpeed0[spt,para1,para2,para3,]-noabs_stdChoSpeed0[spt,para1,para2,para3,], rev(noabs_ChoSpeed0[spt,para1,para2,para3,]+noabs_stdChoSpeed0[spt,para1,para2,para3,])), border=NA, col=adjustcolor("gray", alpha.f = 0.3))
          
          # legend("bottomleft", legend = c("Choice", "Resistance"), col = c("black", "red"), text.font = 3, lwd = 3, lty=c(1,1))
          # legend("bottomright", legend = c( "Evolving", "Fixed"), col = c("black"), text.font = 3, lwd = 3, lty=c(1,2))
          dev.off()        
        }}}}#}
}


{
  for(para3 in 1:length(Parameter3)){
    for(para4 in 1:length(Parameter4)){
      #for(para1 in 1:length(Parameter1)){
      para1 = 0
      for(para2 in 1:length(Parameter2)){
        for(spt in 1:length(tTspeed)){
          png(paste0(getwd(), "/",Folder,"/", plotFolder, "/",subFolder,"/noabs_SpeedEvolutionTraits_Param1",para1, "_Param3", para3, "_Param4",para4, "_Param2", para2, "_gen",tTspeed[spt], "_paracomb",ParameterCombination,".png"))
          par(mar = c(5, 5, 4, 5) + 0.3)
          
          matplot(Parameter1,noabs_ChoSpeed1[spt,,para2,para3,para4],ylim = c(Minspeed,Maxspeed),cex.axis = 2, type = "n", ylab= list("Evolution rate", font = 3, cex = 2),xlab=list("Parameter 1", font = 3, cex = 2))
          lines(Parameter1,noabs_ChoSpeed1[spt,,para2,para3,para4],type = 'l',col="black", lty = 1, lwd = 3);
          lines(Parameter1,noabs_ChoSpeed0[spt,,para2,para3,para4],type = 'l',col="black", lty = 2, lwd = 3);
          
          #----------------- Resistance -----------------
          lines(Parameter1,noabs_ResSpeed1[spt,,para2,para3,para4],type = 'l',col="red", lty = 1, lwd = 3);
          lines(Parameter1,noabs_ResSpeed0[spt,,para2,para3,para4],type = 'l',col="red", lty = 2, lwd = 3);
          
          polygon(x=c(Parameter1, rev(Parameter1)), y=c(noabs_ResSpeed1[spt,,para2,para3,para4]-noabs_stdResSpeed1[spt,,para2,para3,para4], rev(noabs_ResSpeed1[spt,,para2,para3,para4]+noabs_stdResSpeed1[spt,,para2,para3,para4])), border=NA, col=adjustcolor("gray", alpha.f = 0.3))
          polygon(x=c(Parameter1, rev(Parameter1)), y=c(noabs_ResSpeed0[spt,,para2,para3,para4]-noabs_stdResSpeed0[spt,,para2,para3,para4], rev(noabs_ResSpeed0[spt,,para2,para3,para4]+noabs_stdResSpeed0[spt,,para2,para3,para4])), border=NA, col=adjustcolor("gray", alpha.f = 0.3))
          polygon(x=c(Parameter1, rev(Parameter1)), y=c(noabs_ChoSpeed1[spt,,para2,para3,para4]-noabs_stdChoSpeed1[spt,,para2,para3,para4], rev(noabs_ChoSpeed1[spt,,para2,para3,para4]+noabs_stdChoSpeed1[spt,,para2,para3,para4])), border=NA, col=adjustcolor("gray", alpha.f = 0.3))
          polygon(x=c(Parameter1, rev(Parameter1)), y=c(noabs_ChoSpeed0[spt,,para2,para3,para4]-noabs_stdChoSpeed0[spt,,para2,para3,para4], rev(noabs_ChoSpeed0[spt,,para2,para3,para4]+noabs_stdChoSpeed0[spt,,para2,para3,para4])), border=NA, col=adjustcolor("gray", alpha.f = 0.3))
          
          # legend("bottomleft", legend = c("Choice", "Resistance"), col = c("black", "red"), text.font = 3, lwd = 3, lty=c(1,1))
          # legend("bottomright", legend = c( "Evolving", "Fixed"), col = c("black"), text.font = 3, lwd = 3, lty=c(1,2))
          dev.off()        
        }}}}#}
}


{
  for(para3 in 1:length(Parameter3)){
    for(para4 in 1:length(Parameter4)){
      for(para1 in 1:length(Parameter1)){
        #for(para2 in 1:length(Parameter2)){
        for(spt in 1:length(tTspeed)){
          para2 = 0
          png(paste0(getwd(), "/",Folder,"/", plotFolder, "/",subFolder,"/noabs_SpeedEvolutionTraits_Param1",para1, "_Param3", para3, "_Param4",para4, "_Param2", para2, "_gen",tTspeed[spt], "_paracomb",ParameterCombination,".png"))
          
          par(mar = c(5, 5, 4, 5) + 0.3)
          
          matplot(Parameter2,noabs_ChoSpeed1[spt,para1,,para3,para4],ylim = c(Minspeed,Maxspeed),cex.axis = 2, type = "n", ylab= list("Evolution rate", font = 3, cex = 2),xlab=list("Parameter 2", font = 3, cex = 2))
          lines(Parameter2,noabs_ChoSpeed1[spt,para1,,para3,para4],type = 'l',col="black", lty = 1, lwd = 3);
          lines(Parameter2,noabs_ChoSpeed0[spt,para1,,para3,para4],type = 'l',col="black", lty = 2, lwd = 3);
          
          #----------------- Resistance -----------------
          lines(Parameter2,noabs_ResSpeed1[spt,para1,,para3,para4],type = 'l',col="red", lty = 1, lwd = 3);
          lines(Parameter2,noabs_ResSpeed0[spt,para1,,para3,para4],type = 'l',col="red", lty = 2, lwd = 3);
          
          polygon(x=c(Parameter2, rev(Parameter2)), y=c(noabs_ResSpeed1[spt,para1,,para3,para4]-noabs_stdResSpeed1[spt,para1,,para3,para4], rev(noabs_ResSpeed1[spt,para1,,para3,para4]+noabs_stdResSpeed1[spt,para1,,para3,para4])), border=NA, col=adjustcolor("gray", alpha.f = 0.3))
          polygon(x=c(Parameter2, rev(Parameter2)), y=c(noabs_ResSpeed0[spt,para1,,para3,para4]-noabs_stdResSpeed0[spt,para1,,para3,para4], rev(noabs_ResSpeed0[spt,para1,,para3,para4]+noabs_stdResSpeed0[spt,para1,,para3,para4])), border=NA, col=adjustcolor("gray", alpha.f = 0.3))
          polygon(x=c(Parameter2, rev(Parameter2)), y=c(noabs_ChoSpeed1[spt,para1,,para3,para4]-noabs_stdChoSpeed1[spt,para1,,para3,para4], rev(noabs_ChoSpeed1[spt,para1,,para3,para4]+noabs_stdChoSpeed1[spt,para1,,para3,para4])), border=NA, col=adjustcolor("gray", alpha.f = 0.3))
          polygon(x=c(Parameter2, rev(Parameter2)), y=c(noabs_ChoSpeed0[spt,para1,,para3,para4]-noabs_stdChoSpeed0[spt,para1,,para3,para4], rev(noabs_ChoSpeed0[spt,para1,,para3,para4]+noabs_stdChoSpeed0[spt,para1,,para3,para4])), border=NA, col=adjustcolor("gray", alpha.f = 0.3))
          
          
          # legend("bottomleft", legend = c("Choice", "Resistance"), col = c("black", "red"), text.font = 3, lwd = 3, lty=c(1,1))
          # legend("bottomright", legend = c( "Evolving", "Fixed"), col = c("black"), text.font = 3, lwd = 3, lty=c(1,2))
          dev.off()        
        }}}}#}
}


#------------------ Plant Population size -----------------------

subFolder = "PlantPopSize"

if(!dir.exists(paste0(getwd(), "/",Folder, "/", plotFolder, "/", subFolder))){
  dir.create(paste0(getwd(), "/",Folder, "/", plotFolder, "/", subFolder))
}


{
  #for(para3 in 1:length(Parameter3)){
  para3 = 0
  for(para4 in 1:length(Parameter4)){
    for(para1 in 1:length(Parameter1)){
      for(para2 in 1:length(Parameter2)){
        
        png(paste0(getwd(), "/",Folder,"/", plotFolder, "/",subFolder,"/PlantPopsize_Param1",para1, "_Param3", para3, "_Param4",para4, "_Param2", para2, "_paracomb",ParameterCombination,".png"))
        
        par(mar = c(5, 5, 4, 4) + 0.3)
        plot(0,0,xlim = range(Parameter3),ylim = c(0,max(N1_mean0[tT,para1,para2,,para4], N2_mean0[tT,para1,para2,,para4], N1_mean1[tT,para1,para2,,para4], N2_mean1[tT,para1,para2,,para4], na.rm = TRUE)),cex.axis = 2, type = "n",
             ylab= list("Population size", font = 3, cex = 2),xlab= list("Parameter 3", font = 3, cex = 2))
        lines(Parameter3,N1_mean1[tT,para1,para2,,para4],type = 'l',col="purple",lwd = 3);
        polygon(x=c(Parameter3, rev(Parameter3)), y=c(N1_mean1[tT,para1,para2,,para4]-N1_std1[tT,para1,para2,,para4], rev(N1_mean1[tT,para1,para2,,para4]+N1_std1[tT,para1,para2,,para4])), border=NA, col=adjustcolor("gray", alpha.f = 0.3))
        lines(Parameter3,N1_mean0[tT,para1,para2,,para4],type = 'l',col="gray", lwd = 3);
        polygon(x=c(Parameter3, rev(Parameter3)), y=c(N1_mean0[tT,para1,para2,,para4]-N1_std0[tT,para1,para2,,para4], rev(N1_mean0[tT,para1,para2,,para4]+N1_std0[tT,para1,para2,,para4])), border=NA, col=adjustcolor("gray", alpha.f = 0.3))
        
        lines(Parameter3,N2_mean1[tT,para1,para2,,para4],type = 'l', lty = 2,col="purple", lwd = 3);
        polygon(x=c(Parameter3, rev(Parameter3)), y=c(N2_mean1[tT,para1,para2,,para4]-N2_std1[tT,para1,para2,,para4], rev(N2_mean1[tT,para1,para2,,para4]+N2_std1[tT,para1,para2,,para4])), border=NA, col=adjustcolor("gray", alpha.f = 0.3))
        lines(Parameter3,N2_mean0[tT,para1,para2,,para4],type = 'l',lty = 2,col="gray", lwd = 3);
        polygon(x=c(Parameter3, rev(Parameter3)), y=c(N2_mean0[tT,para1,para2,,para4]-N2_std0[tT,para1,para2,,para4], rev(N2_mean0[tT,para1,para2,,para4]+N2_std0[tT,para1,para2,,para4])), border=NA, col=adjustcolor("gray", alpha.f = 0.3))
        
        legend("bottomright", legend = c("HOST PLANT", "Non-toxic", "Toxic", "BEHAVIOUR", "Evolving",  "Fixed"), col = c(NA,"gray", "purple", NA,1,1), text.font = 3, lwd = 3, lty=c(NA,1,1,NA,1,2), cex=1.5, text.col = c("blue",1,1,"blue",1,1))
        
        dev.off()        
      }}}#}
}


{
  for(para3 in 1:length(Parameter3)){
    #for(para4 in 1:length(Parameter4)){
    para4 = 0
    for(para1 in 1:length(Parameter1)){
      for(para2 in 1:length(Parameter2)){
        
        png(paste0(getwd(), "/",Folder,"/", plotFolder, "/",subFolder,"/PlantPopsize_Param1",para1, "_Param3", para3, "_Param4",para4, "_Param2", para2, "_paracomb",ParameterCombination,".png"))
        
        par(mar = c(5, 5, 4, 4) + 0.3)
        plot(0,0,xlim = range(Parameter4),ylim = c(0,max(N1_mean0[tT,para1,para2,para3,], N2_mean0[tT,para1,para2,para3,], N1_mean1[tT,para1,para2,para3,], N2_mean1[tT,para1,para2,para3,], na.rm = TRUE)),cex.axis = 2, type = "n",
             ylab= list("Population size", font = 3, cex = 2),xlab= list("Parameter 4", font = 3, cex = 2))
        lines(Parameter4,N1_mean1[tT,para1,para2,para3,],type = 'l',col="purple",lwd = 3);
        polygon(x=c(Parameter4, rev(Parameter4)), y=c(N1_mean1[tT,para1,para2,para3,]-N1_std1[tT,para1,para2,para3,], rev(N1_mean1[tT,para1,para2,para3,]+N1_std1[tT,para1,para2,para3,])), border=NA, col=adjustcolor("gray", alpha.f = 0.3))
        lines(Parameter4,N1_mean0[tT,para1,para2,para3,],type = 'l',col="gray", lwd = 3);
        polygon(x=c(Parameter4, rev(Parameter4)), y=c(N1_mean0[tT,para1,para2,para3,]-N1_std0[tT,para1,para2,para3,], rev(N1_mean0[tT,para1,para2,para3,]+N1_std0[tT,para1,para2,para3,])), border=NA, col=adjustcolor("gray", alpha.f = 0.3))
        
        lines(Parameter4,N2_mean1[tT,para1,para2,para3,],type = 'l', lty = 2,col="purple", lwd = 3);
        polygon(x=c(Parameter4, rev(Parameter4)), y=c(N2_mean1[tT,para1,para2,para3,]-N2_std1[tT,para1,para2,para3,], rev(N2_mean1[tT,para1,para2,para3,]+N2_std1[tT,para1,para2,para3,])), border=NA, col=adjustcolor("gray", alpha.f = 0.3))
        lines(Parameter4,N2_mean0[tT,para1,para2,para3,],type = 'l',lty = 2,col="gray", lwd = 3);
        polygon(x=c(Parameter4, rev(Parameter4)), y=c(N2_mean0[tT,para1,para2,para3,]-N2_std0[tT,para1,para2,para3,], rev(N2_mean0[tT,para1,para2,para3,]+N2_std0[tT,para1,para2,para3,])), border=NA, col=adjustcolor("gray", alpha.f = 0.3))
        
        #legend("bottomright", legend = c("HOST PLANT", "Non-toxic", "Toxic", "BEHAVIOUR", "Evolving",  "Fixed"), col = c(NA,"gray", "purple", NA,1,1), text.font = 3, lwd = 3, lty=c(NA,1,1,NA,1,2), cex=1.5, text.col = c("blue",1,1,"blue",1,1))
        
        dev.off()        
      }}}#}
}


{
  for(para3 in 1:length(Parameter3)){
    for(para4 in 1:length(Parameter4)){
      
      #for(para1 in 1:length(Parameter1)){
      para1 = 0
      for(para2 in 1:length(Parameter2)){
        
        png(paste0(getwd(), "/",Folder,"/", plotFolder, "/",subFolder,"/PlantPopsize_Param1",para1, "_Param3", para3, "_Param4",para4, "_Param2", para2, "_paracomb",ParameterCombination,".png"))
        
        par(mar = c(5, 5, 4, 4) + 0.3)
        plot(0,0,xlim = range(Parameter2),ylim = c(0,max(N1_mean0[tT,,para2,para3,para4], N2_mean0[tT,,para2,para3,para4], N1_mean1[tT,,para2,para3,para4], N2_mean1[tT,,para2,para3,para4], na.rm = TRUE)),cex.axis = 2, type = "n",
             ylab= list("Population size", font = 3, cex = 2),xlab= list("Parameter 1", font = 3, cex = 2))
        lines(Parameter2,N1_mean1[tT,,para2,para3,para4],type = 'l',col="purple",lwd = 3);
        polygon(x=c(Parameter2, rev(Parameter2)), y=c(N1_mean1[tT,,para2,para3,para4]-N1_std1[tT,,para2,para3,para4], rev(N1_mean1[tT,,para2,para3,para4]+N1_std1[tT,,para2,para3,para4])), border=NA, col=adjustcolor("gray", alpha.f = 0.3))
        lines(Parameter2,N1_mean0[tT,,para2,para3,para4],type = 'l',col="gray", lwd = 3);
        polygon(x=c(Parameter2, rev(Parameter2)), y=c(N1_mean0[tT,,para2,para3,para4]-N1_std0[tT,,para2,para3,para4], rev(N1_mean0[tT,,para2,para3,para4]+N1_std0[tT,,para2,para3,para4])), border=NA, col=adjustcolor("gray", alpha.f = 0.3))
        
        lines(Parameter2,N2_mean1[tT,,para2,para3,para4],type = 'l', lty = 2,col="purple", lwd = 3);
        polygon(x=c(Parameter2, rev(Parameter2)), y=c(N2_mean1[tT,,para2,para3,para4]-N2_std1[tT,,para2,para3,para4], rev(N2_mean1[tT,,para2,para3,para4]+N2_std1[tT,,para2,para3,para4])), border=NA, col=adjustcolor("gray", alpha.f = 0.3))
        lines(Parameter2,N2_mean0[tT,,para2,para3,para4],type = 'l',lty = 2,col="gray", lwd = 3);
        polygon(x=c(Parameter2, rev(Parameter2)), y=c(N2_mean0[tT,,para2,para3,para4]-N2_std0[tT,,para2,para3,para4], rev(N2_mean0[tT,,para2,para3,para4]+N2_std0[tT,,para2,para3,para4])), border=NA, col=adjustcolor("gray", alpha.f = 0.3))
        
        #legend("bottomright", legend = c("HOST PLANT", "Non-toxic", "Toxic", "BEHAVIOUR", "Evolving",  "Fixed"), col = c(NA,"gray", "purple", NA,1,1), text.font = 3, lwd = 3, lty=c(NA,1,1,NA,1,2), cex=1.5, text.col = c("blue",1,1,"blue",1,1))
        
        dev.off()        
      }}}#}
}


{
  for(para3 in 1:length(Parameter3)){
    for(para4 in 1:length(Parameter4)){
      for(para1 in 1:length(Parameter1)){
        #for(para2 in 1:length(Parameter2)){
        para2 = 0 
        png(paste0(getwd(), "/",Folder,"/", plotFolder, "/",subFolder,"/PlantPopsize_Param1",para1, "_Param3", para3, "_Param4",para4, "_Param2", para2, "_paracomb",ParameterCombination,".png"))
        
        par(mar = c(5, 5, 4, 4) + 0.3)
        plot(0,0,xlim = range(Parameter1),ylim = c(0,max(N1_mean0[tT,para1,,para3,para4], N2_mean0[tT,para1,,para3,para4], N1_mean1[tT,para1,,para3,para4], N2_mean1[tT,para1,,para3,para4], na.rm = TRUE)),cex.axis = 2, type = "n",
             ylab= list("Population size", font = 3, cex = 2),xlab= list("Parameter 2", font = 3, cex = 2))
        lines(Parameter1,N1_mean1[tT,para1,,para3,para4],type = 'l',col="purple",lwd = 3);
        polygon(x=c(Parameter1, rev(Parameter1)), y=c(N1_mean1[tT,para1,,para3,para4]-N1_std1[tT,para1,,para3,para4], rev(N1_mean1[tT,para1,,para3,para4]+N1_std1[tT,para1,,para3,para4])), border=NA, col=adjustcolor("gray", alpha.f = 0.3))
        lines(Parameter1,N1_mean0[tT,para1,,para3,para4],type = 'l',col="gray", lwd = 3);
        polygon(x=c(Parameter1, rev(Parameter1)), y=c(N1_mean0[tT,para1,,para3,para4]-N1_std0[tT,para1,,para3,para4], rev(N1_mean0[tT,para1,,para3,para4]+N1_std0[tT,para1,,para3,para4])), border=NA, col=adjustcolor("gray", alpha.f = 0.3))
        
        lines(Parameter1,N2_mean1[tT,para1,,para3,para4],type = 'l', lty = 2,col="purple", lwd = 3);
        polygon(x=c(Parameter1, rev(Parameter1)), y=c(N2_mean1[tT,para1,,para3,para4]-N2_std1[tT,para1,,para3,para4], rev(N2_mean1[tT,para1,,para3,para4]+N2_std1[tT,para1,,para3,para4])), border=NA, col=adjustcolor("gray", alpha.f = 0.3))
        lines(Parameter1,N2_mean0[tT,para1,,para3,para4],type = 'l',lty = 2,col="gray", lwd = 3);
        polygon(x=c(Parameter1, rev(Parameter1)), y=c(N2_mean0[tT,para1,,para3,para4]-N2_std0[tT,para1,,para3,para4], rev(N2_mean0[tT,para1,,para3,para4]+N2_std0[tT,para1,,para3,para4])), border=NA, col=adjustcolor("gray", alpha.f = 0.3))
        
        #legend("bottomright", legend = c("HOST PLANT", "Non-toxic", "Toxic", "BEHAVIOUR", "Evolving",  "Fixed"), col = c(NA,"gray", "purple", NA,1,1), text.font = 3, lwd = 3, lty=c(NA,1,1,NA,1,2), cex=1.5, text.col = c("blue",1,1,"blue",1,1))
        
        dev.off()        
      }}}#}
}


Folder = "Costs_Recombinationrate_Toxicity" # Folder for storing simulated data
FFolder = Folder

ParameterCombination = 2# Assign different numbers for different parameter combinations to help distinguish the names of plots
Parameter1 = Fecunditycost
Parameter2 = Resistancecost
Parameter3 = Toxicity
Parameter4 = Recombinationrate


#------ This folder is for storing plots
plotFolder = "Plots"

if(!dir.exists(paste0(getwd(), "/",Folder, "/", plotFolder))){
  dir.create(paste0(getwd(), "/",Folder, "/", plotFolder))
}

#--------- Loading already analysed data for plotting--------------
DataAnalysis = "AnalysedData"
noabs_stdResSpeed0= readRDS(paste0(getwd(),"/", Folder,"/", DataAnalysis, "/", "noabs_stdResSpeed0.RData"))
noabs_stdResSpeed1= readRDS(paste0(getwd(),"/", Folder,"/", DataAnalysis, "/","noabs_stdResSpeed1.RData"))
noabs_stdChoSpeed0= readRDS(paste0(getwd(),"/", Folder,"/", DataAnalysis, "/","noabs_stdChoSpeed0.RData"))
noabs_stdChoSpeed1= readRDS(paste0(getwd(),"/", Folder,"/", DataAnalysis, "/","noabs_stdChoSpeed1.RData"))

noabs_ResSpeed0= readRDS(paste0(getwd(),"/", Folder,"/", DataAnalysis, "/", "noabs_ResSpeed0.RData"))
noabs_ResSpeed1= readRDS(paste0(getwd(),"/", Folder,"/", DataAnalysis, "/","noabs_ResSpeed1.RData"))
noabs_ChoSpeed0= readRDS(paste0(getwd(),"/", Folder,"/", DataAnalysis, "/","noabs_ChoSpeed0.RData"))
noabs_ChoSpeed1= readRDS(paste0(getwd(),"/", Folder,"/", DataAnalysis, "/","noabs_ChoSpeed1.RData"))


ResSpeed0= readRDS(paste0(getwd(),"/", Folder,"/", DataAnalysis, "/", "ResSpeed0.RData"))
ResSpeed1= readRDS(paste0(getwd(),"/", Folder,"/", DataAnalysis, "/","ResSpeed1.RData"))
ChoSpeed0= readRDS(paste0(getwd(),"/", Folder,"/", DataAnalysis, "/","ChoSpeed0.RData"))
ChoSpeed1= readRDS(paste0(getwd(),"/", Folder,"/", DataAnalysis, "/","ChoSpeed1.RData"))

Rresistancemean1= readRDS(paste0(getwd(),"/", Folder,"/", DataAnalysis, "/", "Rresistancemean1.RData"))
Rresistancesd1 = readRDS(paste0(getwd(),"/", Folder,"/", DataAnalysis, "/","Rresistancesd1.RData"))
Rresistancestd1 = readRDS(paste0(getwd(),"/", Folder,"/", DataAnalysis, "/","Rresistancestd1.RData"))
Cchoicemean1= readRDS(paste0(getwd(),"/", Folder,"/", DataAnalysis, "/","Cchoicemean1.RData"))
Cchoicesd1= readRDS(paste0(getwd(),"/", Folder,"/", DataAnalysis, "/","Cchoicesd1.RData"))
Cchoicestd1= readRDS(paste0(getwd(),"/", Folder,"/", DataAnalysis, "/","Cchoicestd1.RData"))
Rresistancemean0 = readRDS(paste0(getwd(),"/", Folder,"/", DataAnalysis, "/", "Rresistancemean0.RData"))
Rresistancesd0 = readRDS(paste0(getwd(),"/", Folder,"/", DataAnalysis, "/","Rresistancesd0.RData"))
Rresistancestd0 = readRDS(paste0(getwd(),"/", Folder,"/", DataAnalysis, "/","Rresistancestd0.RData"))
Cchoicemean0= readRDS(paste0(getwd(),"/", Folder,"/", DataAnalysis, "/","Cchoicemean0.RData"))
Cchoicesd0= readRDS(paste0(getwd(),"/", Folder,"/", DataAnalysis, "/","Cchoicesd0.RData"))
Cchoicestd0= readRDS(paste0(getwd(),"/", Folder,"/", DataAnalysis, "/","Cchoicestd0.RData"))


Rresistance_popvariancemean1= readRDS(paste0(getwd(),"/", Folder,"/", DataAnalysis, "/", "Rresistance_popvariancemean1.RData"))
Rresistance_popvariancestd1 = readRDS(paste0(getwd(),"/", Folder,"/", DataAnalysis, "/","Rresistance_popvariancestd1.RData"))
Cchoice_popvariancemean1= readRDS(paste0(getwd(),"/", Folder,"/", DataAnalysis, "/","Cchoice_popvariancemean1.RData"))
Cchoice_popvariancestd1= readRDS(paste0(getwd(),"/", Folder,"/", DataAnalysis, "/","Cchoice_popvariancestd1.RData"))
Rresistance_popvariancemean0 = readRDS(paste0(getwd(),"/", Folder,"/", DataAnalysis, "/", "Rresistance_popvariancemean0.RData"))
Rresistance_popvariancestd0 = readRDS(paste0(getwd(),"/", Folder,"/", DataAnalysis, "/","Rresistance_popvariancestd0.RData"))
Cchoice_popvariancemean0= readRDS(paste0(getwd(),"/", Folder,"/", DataAnalysis, "/","Cchoice_popvariancemean0.RData"))
Cchoice_popvariancestd0= readRDS(paste0(getwd(),"/", Folder,"/", DataAnalysis, "/","Cchoice_popvariancestd0.RData"))

N1_mean1= readRDS(paste0(getwd(),"/", Folder,"/", DataAnalysis, "/", "N1_mean1.RData"))
N1_mean0= readRDS(paste0(getwd(),"/", Folder,"/", DataAnalysis, "/","N1_mean0.RData"))
N1_std1= readRDS(paste0(getwd(),"/", Folder,"/", DataAnalysis, "/","N1_std1.RData"))
N1_std0= readRDS(paste0(getwd(),"/", Folder,"/", DataAnalysis, "/","N1_std0.RData"))
N2_mean1= readRDS(paste0(getwd(),"/", Folder,"/", DataAnalysis, "/", "N2_mean1.RData"))
N2_mean0= readRDS(paste0(getwd(),"/", Folder,"/", DataAnalysis, "/","N2_mean0.RData"))
N2_std1= readRDS(paste0(getwd(),"/", Folder,"/", DataAnalysis, "/","N2_std1.RData"))
N2_std0= readRDS(paste0(getwd(),"/", Folder,"/", DataAnalysis, "/","N2_std0.RData"))


tttt = targetTtimes[length(targetTtimes)]
ttt = targetTtimes[1]
plottargetTtimes = c(ttt, tttt)


#------------------- Preference - Resistance Population replicates 

subFolder = "Resistance_Preference" # For storing mean traits and population trait variance plots-----------

if(!dir.exists(paste0(getwd(), "/",Folder,"/",plotFolder,"/",subFolder))){
  dir.create(paste0(getwd(), "/",Folder,"/",plotFolder,"/",subFolder))
}


tT = length(targetTtimes)
{
  #for(para3 in 1:length(Parameter3)){
  para3 = 0
  for(para4 in 1:length(Parameter4)){
    for(para1 in 1:length(Parameter1)){
      for(para2 in 1:length(Parameter2)){
        
        png(paste0(getwd(), "/",Folder,"/", plotFolder, "/",subFolder,"/meanTraits_Param1",para1, "_Param3", para3, "_Param4",para4, "_Param2", para2, "_paracomb",ParameterCombination,".png"))
        
        par(mar = c(5, 5, 4, 5) + 0.3)
        
        matplot(Parameter3,Cchoicemean1[tT,para1,para2,,para4],ylim = c(0,1),cex.axis = 2, type = "n", ylab= list("Mean trait", font = 3, cex = 2),xlab=list("Parameter 3", font = 3, cex = 2))
        lines(Parameter3,Cchoicemean1[tT,para1,para2,,para4],type = 'l',col="black", lty = 1, lwd = 3);
        lines(Parameter3,Cchoicemean0[tT,para1,para2,,para4],type = 'l',col="black", lty = 2, lwd = 3);
        
        #----------------- Resistance -----------------
        lines(Parameter3,Rresistancemean1[tT,para1,para2,,para4],type = 'l',col="red", lty = 1, lwd = 3);
        lines(Parameter3,Rresistancemean0[tT,para1,para2,,para4],type = 'l',col="red", lty = 2, lwd = 3);
        #------- Add std errors ----------------
        polygon(x=c(Parameter3, rev(Parameter3)), y=c(Rresistancemean1[tT,para1,para2,,para4]-Rresistancestd1[tT,para1,para2,,para4], rev(Rresistancemean1[tT,para1,para2,,para4]+Rresistancestd1[tT,para1,para2,,para4])), border=NA, col=adjustcolor("gray", alpha.f = 0.3))
        polygon(x=c(Parameter3, rev(Parameter3)), y=c(Rresistancemean0[tT,para1,para2,,para4]-Rresistancestd0[tT,para1,para2,,para4], rev(Rresistancemean0[tT,para1,para2,,para4]+Rresistancestd0[tT,para1,para2,,para4])), border=NA, col=adjustcolor("gray", alpha.f = 0.3))
        polygon(x=c(Parameter3, rev(Parameter3)), y=c(Cchoicemean1[tT,para1,para2,,para4]-Cchoicestd1[tT,para1,para2,,para4], rev(Cchoicemean1[tT,para1,para2,,para4]+Cchoicestd1[tT,para1,para2,,para4])), border=NA, col=adjustcolor("gray", alpha.f = 0.3))
        polygon(x=c(Parameter3, rev(Parameter3)), y=c(Cchoicemean0[tT,para1,para2,,para4]-Cchoicestd0[tT,para1,para2,,para4], rev(Cchoicemean0[tT,para1,para2,,para4]+Cchoicestd0[tT,para1,para2,,para4])), border=NA, col=adjustcolor("gray", alpha.f = 0.3))
        
        legend("bottomleft", legend = c("Trait", "Choice", "Resistance"), col = c(NA,"black", "red"), text.font = 3, lwd = 3, lty=c(1,1,1))
        legend("bottomright", legend = c("Behaviour", "Evolving", "Fixed"), col = c(NA,"black", "black"), text.font = 3, lwd = 3, lty=c(1,1,2))
        dev.off()        
      }}}#}
}


{
  for(para3 in 1:length(Parameter3)){
    #for(para4 in 1:length(Parameter4)){
    para4 = 0
    for(para1 in 1:length(Parameter1)){
      for(para2 in 1:length(Parameter2)){
        
        png(paste0(getwd(), "/",Folder,"/", plotFolder, "/",subFolder,"/meanTraits_Param1",para1, "_Param3", para3, "_Param4",para4, "_Param2", para2, "_paracomb",ParameterCombination,".png"))
        
        par(mar = c(5, 5, 4, 5) + 0.3)
        
        matplot(Parameter4,Cchoicemean1[tT,para1,para2,para3,],ylim = c(0,1),cex.axis = 2, type = "n", ylab= list("Mean trait", font = 3, cex = 2),xlab=list("Parameter 4", font = 3, cex = 2))
        lines(Parameter4,Cchoicemean1[tT,para1,para2,para3,],type = 'l',col="black", lty = 1, lwd = 3);
        lines(Parameter4,Cchoicemean0[tT,para1,para2,para3,],type = 'l',col="black", lty = 2, lwd = 3);
        
        #----------------- Resistance -----------------
        lines(Parameter4,Rresistancemean1[tT,para1,para2,para3,],type = 'l',col="red", lty = 1, lwd = 3);
        lines(Parameter4,Rresistancemean0[tT,para1,para2,para3,],type = 'l',col="red", lty = 2, lwd = 3);
        
        polygon(x=c(Parameter4, rev(Parameter4)), y=c(Rresistancemean1[tT,para1,para2,para3,]-Rresistancestd1[tT,para1,para2,para3,], rev(Rresistancemean1[tT,para1,para2,para3,]+Rresistancestd1[tT,para1,para2,para3,])), border=NA, col=adjustcolor("gray", alpha.f = 0.3))
        polygon(x=c(Parameter4, rev(Parameter4)), y=c(Rresistancemean0[tT,para1,para2,para3,]-Rresistancestd0[tT,para1,para2,para3,], rev(Rresistancemean0[tT,para1,para2,para3,]+Rresistancestd0[tT,para1,para2,para3,])), border=NA, col=adjustcolor("gray", alpha.f = 0.3))
        polygon(x=c(Parameter4, rev(Parameter4)), y=c(Cchoicemean1[tT,para1,para2,para3,]-Cchoicestd1[tT,para1,para2,para3,], rev(Cchoicemean1[tT,para1,para2,para3,]+Cchoicestd1[tT,para1,para2,para3,])), border=NA, col=adjustcolor("gray", alpha.f = 0.3))
        polygon(x=c(Parameter4, rev(Parameter4)), y=c(Cchoicemean0[tT,para1,para2,para3,]-Cchoicestd0[tT,para1,para2,para3,], rev(Cchoicemean0[tT,para1,para2,para3,]+Cchoicestd0[tT,para1,para2,para3,])), border=NA, col=adjustcolor("gray", alpha.f = 0.3))
        
        # legend("bottomleft", legend = c("Choice", "Resistance"), col = c("black", "red"), text.font = 3, lwd = 3, lty=c(1,1))
        # legend("bottomright", legend = c( "Evolving", "Fixed"), col = c("black"), text.font = 3, lwd = 3, lty=c(1,2))
        dev.off()        
      }}}#}
}


{
  for(para3 in 1:length(Parameter3)){
    for(para4 in 1:length(Parameter4)){
      #for(para1 in 1:length(Parameter1)){
      para1 = 0
      for(para2 in 1:length(Parameter2)){
        
        png(paste0(getwd(), "/",Folder,"/", plotFolder, "/",subFolder,"/meanTraits_Param1",para1, "_Param3", para3, "_Param4",para4, "_Param2", para2, "_paracomb",ParameterCombination,".png"))
        
        par(mar = c(5, 5, 4, 5) + 0.3)
        
        matplot(Parameter1,Cchoicemean1[tT,,para2,para3,para4],ylim = c(0,1),cex.axis = 2, type = "n", ylab= list("Mean trait", font = 3, cex = 2),xlab=list("Parameter 1", font = 3, cex = 2))
        lines(Parameter1,Cchoicemean1[tT,,para2,para3,para4],type = 'l',col="black", lty = 1, lwd = 3);
        lines(Parameter1,Cchoicemean0[tT,,para2,para3,para4],type = 'l',col="black", lty = 2, lwd = 3);
        
        #----------------- Resistance -----------------
        lines(Parameter1,Rresistancemean1[tT,,para2,para3,para4],type = 'l',col="red", lty = 1, lwd = 3);
        lines(Parameter1,Rresistancemean0[tT,,para2,para3,para4],type = 'l',col="red", lty = 2, lwd = 3);
        
        polygon(x=c(Parameter1, rev(Parameter1)), y=c(Rresistancemean1[tT,,para2,para3,para4]-Rresistancestd1[tT,,para2,para3,para4], rev(Rresistancemean1[tT,,para2,para3,para4]+Rresistancestd1[tT,,para2,para3,para4])), border=NA, col=adjustcolor("gray", alpha.f = 0.3))
        polygon(x=c(Parameter1, rev(Parameter1)), y=c(Rresistancemean0[tT,,para2,para3,para4]-Rresistancestd0[tT,,para2,para3,para4], rev(Rresistancemean0[tT,,para2,para3,para4]+Rresistancestd0[tT,,para2,para3,para4])), border=NA, col=adjustcolor("gray", alpha.f = 0.3))
        polygon(x=c(Parameter1, rev(Parameter1)), y=c(Cchoicemean1[tT,,para2,para3,para4]-Cchoicestd1[tT,,para2,para3,para4], rev(Cchoicemean1[tT,,para2,para3,para4]+Cchoicestd1[tT,,para2,para3,para4])), border=NA, col=adjustcolor("gray", alpha.f = 0.3))
        polygon(x=c(Parameter1, rev(Parameter1)), y=c(Cchoicemean0[tT,,para2,para3,para4]-Cchoicestd0[tT,,para2,para3,para4], rev(Cchoicemean0[tT,,para2,para3,para4]+Cchoicestd0[tT,,para2,para3,para4])), border=NA, col=adjustcolor("gray", alpha.f = 0.3))
        
        # legend("bottomleft", legend = c("Choice", "Resistance"), col = c("black", "red"), text.font = 3, lwd = 3, lty=c(1,1))
        # legend("bottomright", legend = c( "Evolving", "Fixed"), col = c("black"), text.font = 3, lwd = 3, lty=c(1,2))
        dev.off()        
      }}}#}
}


{
  for(para3 in 1:length(Parameter3)){
    for(para4 in 1:length(Parameter4)){
      for(para1 in 1:length(Parameter1)){
        #for(para2 in 1:length(Parameter2)){
        para2 = 0
        png(paste0(getwd(), "/",Folder,"/", plotFolder, "/",subFolder,"/meanTraits_Param1",para1, "_Param3", para3, "_Param4",para4, "_Param2", para2, "_paracomb",ParameterCombination,".png"))
        
        par(mar = c(5, 5, 4, 5) + 0.3)
        
        matplot(Parameter2,Cchoicemean1[tT,para1,,para3,para4],ylim = c(0,1),cex.axis = 2, type = "n", ylab= list("Mean trait", font = 3, cex = 2),xlab=list("Parameter 2", font = 3, cex = 2))
        lines(Parameter2,Cchoicemean1[tT,para1,,para3,para4],type = 'l',col="black", lty = 1, lwd = 3);
        lines(Parameter2,Cchoicemean0[tT,para1,,para3,para4],type = 'l',col="black", lty = 2, lwd = 3);
        
        #----------------- Resistance -----------------
        lines(Parameter2,Rresistancemean1[tT,para1,,para3,para4],type = 'l',col="red", lty = 1, lwd = 3);
        lines(Parameter2,Rresistancemean0[tT,para1,,para3,para4],type = 'l',col="red", lty = 2, lwd = 3);
        
        polygon(x=c(Parameter2, rev(Parameter2)), y=c(Rresistancemean1[tT,para1,,para3,para4]-Rresistancestd1[tT,para1,,para3,para4], rev(Rresistancemean1[tT,para1,,para3,para4]+Rresistancestd1[tT,para1,,para3,para4])), border=NA, col=adjustcolor("gray", alpha.f = 0.3))
        polygon(x=c(Parameter2, rev(Parameter2)), y=c(Rresistancemean0[tT,para1,,para3,para4]-Rresistancestd0[tT,para1,,para3,para4], rev(Rresistancemean0[tT,para1,,para3,para4]+Rresistancestd0[tT,para1,,para3,para4])), border=NA, col=adjustcolor("gray", alpha.f = 0.3))
        polygon(x=c(Parameter2, rev(Parameter2)), y=c(Cchoicemean1[tT,para1,,para3,para4]-Cchoicestd1[tT,para1,,para3,para4], rev(Cchoicemean1[tT,para1,,para3,para4]+Cchoicestd1[tT,para1,,para3,para4])), border=NA, col=adjustcolor("gray", alpha.f = 0.3))
        polygon(x=c(Parameter2, rev(Parameter2)), y=c(Cchoicemean0[tT,para1,,para3,para4]-Cchoicestd0[tT,para1,,para3,para4], rev(Cchoicemean0[tT,para1,,para3,para4]+Cchoicestd0[tT,para1,,para3,para4])), border=NA, col=adjustcolor("gray", alpha.f = 0.3))
        
        # legend("bottomleft", legend = c("Choice", "Resistance"), col = c("black", "red"), text.font = 3, lwd = 3, lty=c(1,1))
        # legend("bottomright", legend = c( "Evolving", "Fixed"), col = c("black"), text.font = 3, lwd = 3, lty=c(1,2))
        dev.off()        
      }}}#}
}


#----------------- mean trait Variance within populations --------------
Minvar = min(c(Cchoice_popvariancemean1,Cchoice_popvariancemean0, Rresistance_popvariancemean0, Rresistance_popvariancemean1), na.rm = T)
Maxvar = max(c(Cchoice_popvariancemean1,Cchoice_popvariancemean0, Rresistance_popvariancemean0, Rresistance_popvariancemean1), na.rm = T)
{
  #for(para3 in 1:length(Parameter3)){
  para3 = 0
  for(para4 in 1:length(Parameter4)){
    for(para1 in 1:length(Parameter1)){
      for(para2 in 1:length(Parameter2)){
        
        png(paste0(getwd(), "/",Folder,"/", plotFolder, "/",subFolder,"/VarianceTraits_Param1",para1, "_Param3", para3, "_Param4",para4, "_Param2", para2, "_paracomb",ParameterCombination,".png"))
        
        par(mar = c(5, 5, 4, 5) + 0.3)
        
        matplot(Parameter3,Cchoice_popvariancemean1[tT,para1,para2,,para4],ylim = c(Minvar,Maxvar),cex.axis = 2, type = "n", ylab= list("Population trait variance", font = 3, cex = 2),xlab=list("Parameter 3", font = 3, cex = 2))
        lines(Parameter3,Cchoice_popvariancemean1[tT,para1,para2,,para4],type = 'l',col="black", lty = 1, lwd = 3);
        lines(Parameter3,Cchoice_popvariancemean0[tT,para1,para2,,para4],type = 'l',col="black", lty = 2, lwd = 3);
        
        
        #----------------- Resistance -----------------
        lines(Parameter3,Rresistance_popvariancemean1[tT,para1,para2,,para4],type = 'l',col="red", lty = 1, lwd = 3);
        lines(Parameter3,Rresistance_popvariancemean0[tT,para1,para2,,para4],type = 'l',col="red", lty = 2, lwd = 3);
        
        polygon(x=c(Parameter3, rev(Parameter3)), y=c(Rresistance_popvariancemean1[tT,para1,para2,,para4]-Rresistance_popvariancestd1[tT,para1,para2,,para4], rev(Rresistance_popvariancemean1[tT,para1,para2,,para4]+Rresistance_popvariancestd1[tT,para1,para2,,para4])), border=NA, col=adjustcolor("gray", alpha.f = 0.3))
        polygon(x=c(Parameter3, rev(Parameter3)), y=c(Rresistance_popvariancemean0[tT,para1,para2,,para4]-Rresistance_popvariancestd0[tT,para1,para2,,para4], rev(Rresistance_popvariancemean0[tT,para1,para2,,para4]+Rresistance_popvariancestd0[tT,para1,para2,,para4])), border=NA, col=adjustcolor("gray", alpha.f = 0.3))
        polygon(x=c(Parameter3, rev(Parameter3)), y=c(Cchoice_popvariancemean1[tT,para1,para2,,para4]-Cchoice_popvariancestd1[tT,para1,para2,,para4], rev(Cchoice_popvariancemean1[tT,para1,para2,,para4]+Cchoice_popvariancestd1[tT,para1,para2,,para4])), border=NA, col=adjustcolor("gray", alpha.f = 0.3))
        polygon(x=c(Parameter3, rev(Parameter3)), y=c(Cchoice_popvariancemean0[tT,para1,para2,,para4]-Cchoice_popvariancestd0[tT,para1,para2,,para4], rev(Cchoice_popvariancemean0[tT,para1,para2,,para4]+Cchoice_popvariancestd0[tT,para1,para2,,para4])), border=NA, col=adjustcolor("gray", alpha.f = 0.3))
        
        
        legend("bottomleft", legend = c("Trait", "Choice", "Resistance"), col = c(NA,"black", "red"), text.font = 3, lwd = 3, lty=c(1,1,1))
        legend("bottomright", legend = c("Behaviour", "Evolving", "Fixed"), col = c(NA,"black", "black"), text.font = 3, lwd = 3, lty=c(1,1,2))
        dev.off()        
      }}}#}
}


{
  for(para3 in 1:length(Parameter3)){
    #for(para4 in 1:length(Parameter4)){
    para4 = 0
    for(para1 in 1:length(Parameter1)){
      for(para2 in 1:length(Parameter2)){
        
        png(paste0(getwd(), "/",Folder,"/", plotFolder, "/",subFolder,"/VarianceTraits_Param1",para1, "_Param3", para3, "_Param4",para4, "_Param2", para2, "_paracomb",ParameterCombination,".png"))
        
        par(mar = c(5, 5, 4, 5) + 0.3)
        
        matplot(Parameter4,Cchoice_popvariancemean1[tT,para1,para2,para3,],ylim = c(Minvar,Maxvar),cex.axis = 2, type = "n", ylab= list("Population trait variance", font = 3, cex = 2),xlab=list("Parameter 4", font = 3, cex = 2))
        lines(Parameter4,Cchoice_popvariancemean1[tT,para1,para2,para3,],type = 'l',col="black", lty = 1, lwd = 3);
        lines(Parameter4,Cchoice_popvariancemean0[tT,para1,para2,para3,],type = 'l',col="black", lty = 2, lwd = 3);
        
        #----------------- Resistance -----------------
        lines(Parameter4,Rresistance_popvariancemean1[tT,para1,para2,para3,],type = 'l',col="red", lty = 1, lwd = 3);
        lines(Parameter4,Rresistance_popvariancemean0[tT,para1,para2,para3,],type = 'l',col="red", lty = 2, lwd = 3);
        
        polygon(x=c(Parameter4, rev(Parameter4)), y=c(Rresistance_popvariancemean1[tT,para1,para2,para3,]-Rresistance_popvariancestd1[tT,para1,para2,para3,], rev(Rresistance_popvariancemean1[tT,para1,para2,para3,]+Rresistance_popvariancestd1[tT,para1,para2,para3,])), border=NA, col=adjustcolor("gray", alpha.f = 0.3))
        polygon(x=c(Parameter4, rev(Parameter4)), y=c(Rresistance_popvariancemean0[tT,para1,para2,para3,]-Rresistance_popvariancestd0[tT,para1,para2,para3,], rev(Rresistance_popvariancemean0[tT,para1,para2,para3,]+Rresistance_popvariancestd0[tT,para1,para2,para3,])), border=NA, col=adjustcolor("gray", alpha.f = 0.3))
        polygon(x=c(Parameter4, rev(Parameter4)), y=c(Cchoice_popvariancemean1[tT,para1,para2,para3,]-Cchoice_popvariancestd1[tT,para1,para2,para3,], rev(Cchoice_popvariancemean1[tT,para1,para2,para3,]+Cchoice_popvariancestd1[tT,para1,para2,para3,])), border=NA, col=adjustcolor("gray", alpha.f = 0.3))
        polygon(x=c(Parameter4, rev(Parameter4)), y=c(Cchoice_popvariancemean0[tT,para1,para2,para3,]-Cchoice_popvariancestd0[tT,para1,para2,para3,], rev(Cchoice_popvariancemean0[tT,para1,para2,para3,]+Cchoice_popvariancestd0[tT,para1,para2,para3,])), border=NA, col=adjustcolor("gray", alpha.f = 0.3))
        
        # legend("bottomleft", legend = c("Choice", "Resistance"), col = c("black", "red"), text.font = 3, lwd = 3, lty=c(1,1))
        # legend("bottomright", legend = c( "Evolving", "Fixed"), col = c("black"), text.font = 3, lwd = 3, lty=c(1,2))
        dev.off()        
      }}}#}
}


{
  for(para3 in 1:length(Parameter3)){
    for(para4 in 1:length(Parameter4)){
      #for(para1 in 1:length(Parameter1)){
      para1 = 0
      for(para2 in 1:length(Parameter2)){
        
        png(paste0(getwd(), "/",Folder,"/", plotFolder, "/",subFolder,"/VarianceTraits_Param1",para1, "_Param3", para3, "_Param4",para4, "_Param2", para2, "_paracomb",ParameterCombination,".png"))
        par(mar = c(5, 5, 4, 5) + 0.3)
        
        matplot(Parameter1,Cchoice_popvariancemean1[tT,,para2,para3,para4],ylim = c(Minvar,Maxvar),cex.axis = 2, type = "n", ylab= list("Population trait variance", font = 3, cex = 2),xlab=list("Parameter 1", font = 3, cex = 2))
        lines(Parameter1,Cchoice_popvariancemean1[tT,,para2,para3,para4],type = 'l',col="black", lty = 1, lwd = 3);
        lines(Parameter1,Cchoice_popvariancemean0[tT,,para2,para3,para4],type = 'l',col="black", lty = 2, lwd = 3);
        
        #----------------- Resistance -----------------
        lines(Parameter1,Rresistance_popvariancemean1[tT,,para2,para3,para4],type = 'l',col="red", lty = 1, lwd = 3);
        lines(Parameter1,Rresistance_popvariancemean0[tT,,para2,para3,para4],type = 'l',col="red", lty = 2, lwd = 3);
        
        polygon(x=c(Parameter1, rev(Parameter1)), y=c(Rresistance_popvariancemean1[tT,,para2,para3,para4]-Rresistance_popvariancestd1[tT,,para2,para3,para4], rev(Rresistance_popvariancemean1[tT,,para2,para3,para4]+Rresistance_popvariancestd1[tT,,para2,para3,para4])), border=NA, col=adjustcolor("gray", alpha.f = 0.3))
        polygon(x=c(Parameter1, rev(Parameter1)), y=c(Rresistance_popvariancemean0[tT,,para2,para3,para4]-Rresistance_popvariancestd0[tT,,para2,para3,para4], rev(Rresistance_popvariancemean0[tT,,para2,para3,para4]+Rresistance_popvariancestd0[tT,,para2,para3,para4])), border=NA, col=adjustcolor("gray", alpha.f = 0.3))
        polygon(x=c(Parameter1, rev(Parameter1)), y=c(Cchoice_popvariancemean1[tT,,para2,para3,para4]-Cchoice_popvariancestd1[tT,,para2,para3,para4], rev(Cchoice_popvariancemean1[tT,,para2,para3,para4]+Cchoice_popvariancestd1[tT,,para2,para3,para4])), border=NA, col=adjustcolor("gray", alpha.f = 0.3))
        polygon(x=c(Parameter1, rev(Parameter1)), y=c(Cchoice_popvariancemean0[tT,,para2,para3,para4]-Cchoice_popvariancestd0[tT,,para2,para3,para4], rev(Cchoice_popvariancemean0[tT,,para2,para3,para4]+Cchoice_popvariancestd0[tT,,para2,para3,para4])), border=NA, col=adjustcolor("gray", alpha.f = 0.3))
        
        # legend("bottomleft", legend = c("Choice", "Resistance"), col = c("black", "red"), text.font = 3, lwd = 3, lty=c(1,1))
        # legend("bottomright", legend = c( "Evolving", "Fixed"), col = c("black"), text.font = 3, lwd = 3, lty=c(1,2))
        dev.off()        
      }}}#}
}


{
  for(para3 in 1:length(Parameter3)){
    for(para4 in 1:length(Parameter4)){
      for(para1 in 1:length(Parameter1)){
        #for(para2 in 1:length(Parameter2)){
        para2 = 0
        png(paste0(getwd(), "/",Folder,"/", plotFolder, "/",subFolder,"/VarianceTraits_Param1",para1, "_Param3", para3, "_Param4",para4, "_Param2", para2, "_paracomb",ParameterCombination,".png"))
        
        par(mar = c(5, 5, 4, 5) + 0.3)
        
        matplot(Parameter2,Cchoice_popvariancemean1[tT,para1,,para3,para4],ylim = c(Minvar,Maxvar),cex.axis = 2, type = "n", ylab= list("Population trait variance", font = 3, cex = 2),xlab=list("Parameter 2", font = 3, cex = 2))
        lines(Parameter2,Cchoice_popvariancemean1[tT,para1,,para3,para4],type = 'l',col="black", lty = 1, lwd = 3);
        lines(Parameter2,Cchoice_popvariancemean0[tT,para1,,para3,para4],type = 'l',col="black", lty = 2, lwd = 3);
        
        #----------------- Resistance -----------------
        lines(Parameter2,Rresistance_popvariancemean1[tT,para1,,para3,para4],type = 'l',col="red", lty = 1, lwd = 3);
        lines(Parameter2,Rresistance_popvariancemean0[tT,para1,,para3,para4],type = 'l',col="red", lty = 2, lwd = 3);
        
        polygon(x=c(Parameter2, rev(Parameter2)), y=c(Rresistance_popvariancemean1[tT,para1,,para3,para4]-Rresistance_popvariancestd1[tT,para1,,para3,para4], rev(Rresistance_popvariancemean1[tT,para1,,para3,para4]+Rresistance_popvariancestd1[tT,para1,,para3,para4])), border=NA, col=adjustcolor("gray", alpha.f = 0.3))
        polygon(x=c(Parameter2, rev(Parameter2)), y=c(Rresistance_popvariancemean0[tT,para1,,para3,para4]-Rresistance_popvariancestd0[tT,para1,,para3,para4], rev(Rresistance_popvariancemean0[tT,para1,,para3,para4]+Rresistance_popvariancestd0[tT,para1,,para3,para4])), border=NA, col=adjustcolor("gray", alpha.f = 0.3))
        polygon(x=c(Parameter2, rev(Parameter2)), y=c(Cchoice_popvariancemean1[tT,para1,,para3,para4]-Cchoice_popvariancestd1[tT,para1,,para3,para4], rev(Cchoice_popvariancemean1[tT,para1,,para3,para4]+Cchoice_popvariancestd1[tT,para1,,para3,para4])), border=NA, col=adjustcolor("gray", alpha.f = 0.3))
        polygon(x=c(Parameter2, rev(Parameter2)), y=c(Cchoice_popvariancemean0[tT,para1,,para3,para4]-Cchoice_popvariancestd0[tT,para1,,para3,para4], rev(Cchoice_popvariancemean0[tT,para1,,para3,para4]+Cchoice_popvariancestd0[tT,para1,,para3,para4])), border=NA, col=adjustcolor("gray", alpha.f = 0.3))
        
        
        # legend("bottomleft", legend = c("Choice", "Resistance"), col = c("black", "red"), text.font = 3, lwd = 3, lty=c(1,1))
        # legend("bottomright", legend = c( "Evolving", "Fixed"), col = c("black"), text.font = 3, lwd = 3, lty=c(1,2))
        dev.off()        
      }}}#}
}

# ------------------- Evolution rate ----------
subFolder = "SpeedResistance_Preference"

if(!dir.exists(paste0(getwd(), "/",Folder, "/", plotFolder, "/", subFolder))){
  dir.create(paste0(getwd(), "/",Folder, "/", plotFolder, "/", subFolder))
}


#----------------- Speed of adaptation --------------
Minspeed = min(c(ChoSpeed1,ChoSpeed0, ResSpeed0, ResSpeed1), na.rm = T)
Maxspeed = max(c(ChoSpeed1,ChoSpeed0, ResSpeed0, ResSpeed1), na.rm = T)
{
  #for(para3 in 1:length(Parameter3)){
  para3 = 0
  for(para4 in 1:length(Parameter4)){
    for(para1 in 1:length(Parameter1)){
      for(para2 in 1:length(Parameter2)){
        for(spt in 1:length(tTspeed)){
          png(paste0(getwd(), "/",Folder,"/", plotFolder, "/",subFolder,"/abs_SpeedEvolutionTraits_Param1",para1, "_Param3", para3, "_Param4",para4, "_Param2", para2, "_gen",tTspeed[spt], "_paracomb",ParameterCombination,".png"))
          
          par(mar = c(5, 5, 4, 5) + 0.3)
          
          matplot(Parameter3,ChoSpeed1[spt,para1,para2,,para4],ylim = c(Minspeed,Maxspeed),cex.axis = 2, type = "n", ylab= list("Evolution rate", font = 3, cex = 2),xlab=list("Parameter 3", font = 3, cex = 2))
          lines(Parameter3,ChoSpeed1[spt,para1,para2,,para4],type = 'l',col="black", lty = 1, lwd = 3);
          lines(Parameter3,ChoSpeed0[spt,para1,para2,,para4],type = 'l',col="black", lty = 2, lwd = 3);
          
          #----------------- Resistance -----------------
          lines(Parameter3,ResSpeed1[spt,para1,para2,,para4],type = 'l',col="red", lty = 1, lwd = 3);
          lines(Parameter3,ResSpeed0[spt,para1,para2,,para4],type = 'l',col="red", lty = 2, lwd = 3);
          
          legend("bottomleft", legend = c("Trait", "Choice", "Resistance"), col = c(NA,"black", "red"), text.font = 3, lwd = 3, lty=c(1,1,1))
          legend("bottomright", legend = c("Behaviour", "Evolving", "Fixed"), col = c(NA,"black", "black"), text.font = 3, lwd = 3, lty=c(1,1,2))
          dev.off()        
        }}}}#}
}


{
  for(para3 in 1:length(Parameter3)){
    #for(para4 in 1:length(Parameter4)){
    para4 = 0
    for(para1 in 1:length(Parameter1)){
      for(para2 in 1:length(Parameter2)){
        for(spt in 1:length(tTspeed)){
          png(paste0(getwd(), "/",Folder,"/", plotFolder, "/",subFolder,"/abs_SpeedEvolutionTraits_Param1",para1, "_Param3", para3, "_Param4",para4, "_Param2", para2, "_gen",tTspeed[spt], "_paracomb",ParameterCombination,".png"))
          
          par(mar = c(5, 5, 4, 5) + 0.3)
          
          matplot(Parameter4,ChoSpeed1[spt,para1,para2,para3,],ylim = c(Minspeed,Maxspeed),cex.axis = 2, type = "n", ylab= list("Evolution rate", font = 3, cex = 2),xlab=list("Parameter 4", font = 3, cex = 2))
          lines(Parameter4,ChoSpeed1[spt,para1,para2,para3,],type = 'l',col="black", lty = 1, lwd = 3);
          lines(Parameter4,ChoSpeed0[spt,para1,para2,para3,],type = 'l',col="black", lty = 2, lwd = 3);
          
          #----------------- Resistance -----------------
          lines(Parameter4,ResSpeed1[spt,para1,para2,para3,],type = 'l',col="red", lty = 1, lwd = 3);
          lines(Parameter4,ResSpeed0[spt,para1,para2,para3,],type = 'l',col="red", lty = 2, lwd = 3);
          # legend("bottomleft", legend = c("Choice", "Resistance"), col = c("black", "red"), text.font = 3, lwd = 3, lty=c(1,1))
          # legend("bottomright", legend = c( "Evolving", "Fixed"), col = c("black"), text.font = 3, lwd = 3, lty=c(1,2))
          dev.off()        
        }}}}#}
}


{
  for(para3 in 1:length(Parameter3)){
    for(para4 in 1:length(Parameter4)){
      #for(para1 in 1:length(Parameter1)){
      para1 = 0
      for(para2 in 1:length(Parameter2)){
        for(spt in 1:length(tTspeed)){
          png(paste0(getwd(), "/",Folder,"/", plotFolder, "/",subFolder,"/abs_SpeedEvolutionTraits_Param1",para1, "_Param3", para3, "_Param4",para4, "_Param2", para2, "_gen",tTspeed[spt], "_paracomb",ParameterCombination,".png"))
          par(mar = c(5, 5, 4, 5) + 0.3)
          
          matplot(Parameter1,ChoSpeed1[spt,,para2,para3,para4],ylim = c(Minspeed,Maxspeed),cex.axis = 2, type = "n", ylab= list("Evolution rate", font = 3, cex = 2),xlab=list("Parameter 1", font = 3, cex = 2))
          lines(Parameter1,ChoSpeed1[spt,,para2,para3,para4],type = 'l',col="black", lty = 1, lwd = 3);
          lines(Parameter1,ChoSpeed0[spt,,para2,para3,para4],type = 'l',col="black", lty = 2, lwd = 3);
          
          #----------------- Resistance -----------------
          lines(Parameter1,ResSpeed1[spt,,para2,para3,para4],type = 'l',col="red", lty = 1, lwd = 3);
          lines(Parameter1,ResSpeed0[spt,,para2,para3,para4],type = 'l',col="red", lty = 2, lwd = 3);
          
          # legend("bottomleft", legend = c("Choice", "Resistance"), col = c("black", "red"), text.font = 3, lwd = 3, lty=c(1,1))
          # legend("bottomright", legend = c( "Evolving", "Fixed"), col = c("black"), text.font = 3, lwd = 3, lty=c(1,2))
          dev.off()        
        }}}}#}
}


{
  for(para3 in 1:length(Parameter3)){
    for(para4 in 1:length(Parameter4)){
      for(para1 in 1:length(Parameter1)){
        #for(para2 in 1:length(Parameter2)){
        for(spt in 1:length(tTspeed)){
          para2 = 0
          png(paste0(getwd(), "/",Folder,"/", plotFolder, "/",subFolder,"/abs_SpeedEvolutionTraits_Param1",para1, "_Param3", para3, "_Param4",para4, "_Param2", para2, "_gen",tTspeed[spt], "_paracomb",ParameterCombination,".png"))
          
          par(mar = c(5, 5, 4, 5) + 0.3)
          
          matplot(Parameter2,ChoSpeed1[spt,para1,,para3,para4],ylim = c(Minspeed,Maxspeed),cex.axis = 2, type = "n", ylab= list("Evolution rate", font = 3, cex = 2),xlab=list("Parameter 2", font = 3, cex = 2))
          lines(Parameter2,ChoSpeed1[spt,para1,,para3,para4],type = 'l',col="black", lty = 1, lwd = 3);
          lines(Parameter2,ChoSpeed0[spt,para1,,para3,para4],type = 'l',col="black", lty = 2, lwd = 3);
          
          #----------------- Resistance -----------------
          lines(Parameter2,ResSpeed1[spt,para1,,para3,para4],type = 'l',col="red", lty = 1, lwd = 3);
          lines(Parameter2,ResSpeed0[spt,para1,,para3,para4],type = 'l',col="red", lty = 2, lwd = 3);
          
          # legend("bottomleft", legend = c("Choice", "Resistance"), col = c("black", "red"), text.font = 3, lwd = 3, lty=c(1,1))
          # legend("bottomright", legend = c( "Evolving", "Fixed"), col = c("black"), text.font = 3, lwd = 3, lty=c(1,2))
          dev.off()        
        }}}}#}
}


#------------------- Speed of adaptation -  Not absolute values ---------------------------
Minspeed = min(c(noabs_ChoSpeed1,noabs_ChoSpeed0, noabs_ResSpeed0, noabs_ResSpeed1), na.rm = T)
Maxspeed = max(c(noabs_ChoSpeed1,noabs_ChoSpeed0, noabs_ResSpeed0, noabs_ResSpeed1), na.rm = T)
{
  #for(para3 in 1:length(Parameter3)){
  para3 = 0
  for(para4 in 1:length(Parameter4)){
    for(para1 in 1:length(Parameter1)){
      for(para2 in 1:length(Parameter2)){
        for(spt in 1:length(tTspeed)){
          png(paste0(getwd(), "/",Folder,"/", plotFolder, "/",subFolder,"/noabs_SpeedEvolutionTraits_Param1",para1, "_Param3", para3, "_Param4",para4, "_Param2", para2, "_gen",tTspeed[spt], "_paracomb",ParameterCombination,".png"))
          
          par(mar = c(5, 5, 4, 5) + 0.3)
          
          matplot(Parameter3,noabs_ChoSpeed1[spt,para1,para2,,para4],ylim = c(Minspeed,Maxspeed),cex.axis = 2, type = "n", ylab= list("Evolution rate", font = 3, cex = 2),xlab=list("Parameter 3", font = 3, cex = 2))
          lines(Parameter3,noabs_ChoSpeed1[spt,para1,para2,,para4],type = 'l',col="black", lty = 1, lwd = 3);
          lines(Parameter3,noabs_ChoSpeed0[spt,para1,para2,,para4],type = 'l',col="black", lty = 2, lwd = 3);
          
          #----------------- Resistance -----------------
          lines(Parameter3,noabs_ResSpeed1[spt,para1,para2,,para4],type = 'l',col="red", lty = 1, lwd = 3);
          lines(Parameter3,noabs_ResSpeed0[spt,para1,para2,,para4],type = 'l',col="red", lty = 2, lwd = 3);
          
          polygon(x=c(Parameter3, rev(Parameter3)), y=c(noabs_ResSpeed1[spt,para1,para2,,para4]-noabs_stdResSpeed1[spt,para1,para2,,para4], rev(noabs_ResSpeed1[spt,para1,para2,,para4]+noabs_stdResSpeed1[spt,para1,para2,,para4])), border=NA, col=adjustcolor("gray", alpha.f = 0.3))
          polygon(x=c(Parameter3, rev(Parameter3)), y=c(noabs_ResSpeed0[spt,para1,para2,,para4]-noabs_stdResSpeed0[spt,para1,para2,,para4], rev(noabs_ResSpeed0[spt,para1,para2,,para4]+noabs_stdResSpeed0[spt,para1,para2,,para4])), border=NA, col=adjustcolor("gray", alpha.f = 0.3))
          polygon(x=c(Parameter3, rev(Parameter3)), y=c(noabs_ChoSpeed1[spt,para1,para2,,para4]-noabs_stdChoSpeed1[spt,para1,para2,,para4], rev(noabs_ChoSpeed1[spt,para1,para2,,para4]+noabs_stdChoSpeed1[spt,para1,para2,,para4])), border=NA, col=adjustcolor("gray", alpha.f = 0.3))
          polygon(x=c(Parameter3, rev(Parameter3)), y=c(noabs_ChoSpeed0[spt,para1,para2,,para4]-noabs_stdChoSpeed0[spt,para1,para2,,para4], rev(noabs_ChoSpeed0[spt,para1,para2,,para4]+noabs_stdChoSpeed0[spt,para1,para2,,para4])), border=NA, col=adjustcolor("gray", alpha.f = 0.3))
          
          legend("bottomleft", legend = c("Trait", "Choice", "Resistance"), col = c(NA,"black", "red"), text.font = 3, lwd = 3, lty=c(1,1,1))
          legend("bottomright", legend = c("Behaviour", "Evolving", "Fixed"), col = c(NA,"black", "black"), text.font = 3, lwd = 3, lty=c(1,1,2))
          dev.off()        
        }}}}#}
}


{
  for(para3 in 1:length(Parameter3)){
    #for(para4 in 1:length(Parameter4)){
    para4 = 0
    for(para1 in 1:length(Parameter1)){
      for(para2 in 1:length(Parameter2)){
        for(spt in 1:length(tTspeed)){
          png(paste0(getwd(), "/",Folder,"/", plotFolder, "/",subFolder,"/noabs_SpeedEvolutionTraits_Param1",para1, "_Param3", para3, "_Param4",para4, "_Param2", para2, "_gen",tTspeed[spt], "_paracomb",ParameterCombination,".png"))
          
          par(mar = c(5, 5, 4, 5) + 0.3)
          
          matplot(Parameter4,noabs_ChoSpeed1[spt,para1,para2,para3,],ylim = c(Minspeed,Maxspeed),cex.axis = 2, type = "n", ylab= list("Evolution rate", font = 3, cex = 2),xlab=list("Parameter 4", font = 3, cex = 2))
          lines(Parameter4,noabs_ChoSpeed1[spt,para1,para2,para3,],type = 'l',col="black", lty = 1, lwd = 3);
          lines(Parameter4,noabs_ChoSpeed0[spt,para1,para2,para3,],type = 'l',col="black", lty = 2, lwd = 3);
          
          #----------------- Resistance -----------------
          lines(Parameter4,noabs_ResSpeed1[spt,para1,para2,para3,],type = 'l',col="red", lty = 1, lwd = 3);
          lines(Parameter4,noabs_ResSpeed0[spt,para1,para2,para3,],type = 'l',col="red", lty = 2, lwd = 3);
          
          polygon(x=c(Parameter4, rev(Parameter4)), y=c(noabs_ResSpeed1[spt,para1,para2,para3,]-noabs_stdResSpeed1[spt,para1,para2,para3,], rev(noabs_ResSpeed1[spt,para1,para2,para3,]+noabs_stdResSpeed1[spt,para1,para2,para3,])), border=NA, col=adjustcolor("gray", alpha.f = 0.3))
          polygon(x=c(Parameter4, rev(Parameter4)), y=c(noabs_ResSpeed0[spt,para1,para2,para3,]-noabs_stdResSpeed0[spt,para1,para2,para3,], rev(noabs_ResSpeed0[spt,para1,para2,para3,]+noabs_stdResSpeed0[spt,para1,para2,para3,])), border=NA, col=adjustcolor("gray", alpha.f = 0.3))
          polygon(x=c(Parameter4, rev(Parameter4)), y=c(noabs_ChoSpeed1[spt,para1,para2,para3,]-noabs_stdChoSpeed1[spt,para1,para2,para3,], rev(noabs_ChoSpeed1[spt,para1,para2,para3,]+noabs_stdChoSpeed1[spt,para1,para2,para3,])), border=NA, col=adjustcolor("gray", alpha.f = 0.3))
          polygon(x=c(Parameter4, rev(Parameter4)), y=c(noabs_ChoSpeed0[spt,para1,para2,para3,]-noabs_stdChoSpeed0[spt,para1,para2,para3,], rev(noabs_ChoSpeed0[spt,para1,para2,para3,]+noabs_stdChoSpeed0[spt,para1,para2,para3,])), border=NA, col=adjustcolor("gray", alpha.f = 0.3))
          
          # legend("bottomleft", legend = c("Choice", "Resistance"), col = c("black", "red"), text.font = 3, lwd = 3, lty=c(1,1))
          # legend("bottomright", legend = c( "Evolving", "Fixed"), col = c("black"), text.font = 3, lwd = 3, lty=c(1,2))
          dev.off()        
        }}}}#}
}


{
  for(para3 in 1:length(Parameter3)){
    for(para4 in 1:length(Parameter4)){
      #for(para1 in 1:length(Parameter1)){
      para1 = 0
      for(para2 in 1:length(Parameter2)){
        for(spt in 1:length(tTspeed)){
          png(paste0(getwd(), "/",Folder,"/", plotFolder, "/",subFolder,"/noabs_SpeedEvolutionTraits_Param1",para1, "_Param3", para3, "_Param4",para4, "_Param2", para2, "_gen",tTspeed[spt], "_paracomb",ParameterCombination,".png"))
          par(mar = c(5, 5, 4, 5) + 0.3)
          
          matplot(Parameter1,noabs_ChoSpeed1[spt,,para2,para3,para4],ylim = c(Minspeed,Maxspeed),cex.axis = 2, type = "n", ylab= list("Evolution rate", font = 3, cex = 2),xlab=list("Parameter 1", font = 3, cex = 2))
          lines(Parameter1,noabs_ChoSpeed1[spt,,para2,para3,para4],type = 'l',col="black", lty = 1, lwd = 3);
          lines(Parameter1,noabs_ChoSpeed0[spt,,para2,para3,para4],type = 'l',col="black", lty = 2, lwd = 3);
          
          #----------------- Resistance -----------------
          lines(Parameter1,noabs_ResSpeed1[spt,,para2,para3,para4],type = 'l',col="red", lty = 1, lwd = 3);
          lines(Parameter1,noabs_ResSpeed0[spt,,para2,para3,para4],type = 'l',col="red", lty = 2, lwd = 3);
          
          polygon(x=c(Parameter1, rev(Parameter1)), y=c(noabs_ResSpeed1[spt,,para2,para3,para4]-noabs_stdResSpeed1[spt,,para2,para3,para4], rev(noabs_ResSpeed1[spt,,para2,para3,para4]+noabs_stdResSpeed1[spt,,para2,para3,para4])), border=NA, col=adjustcolor("gray", alpha.f = 0.3))
          polygon(x=c(Parameter1, rev(Parameter1)), y=c(noabs_ResSpeed0[spt,,para2,para3,para4]-noabs_stdResSpeed0[spt,,para2,para3,para4], rev(noabs_ResSpeed0[spt,,para2,para3,para4]+noabs_stdResSpeed0[spt,,para2,para3,para4])), border=NA, col=adjustcolor("gray", alpha.f = 0.3))
          polygon(x=c(Parameter1, rev(Parameter1)), y=c(noabs_ChoSpeed1[spt,,para2,para3,para4]-noabs_stdChoSpeed1[spt,,para2,para3,para4], rev(noabs_ChoSpeed1[spt,,para2,para3,para4]+noabs_stdChoSpeed1[spt,,para2,para3,para4])), border=NA, col=adjustcolor("gray", alpha.f = 0.3))
          polygon(x=c(Parameter1, rev(Parameter1)), y=c(noabs_ChoSpeed0[spt,,para2,para3,para4]-noabs_stdChoSpeed0[spt,,para2,para3,para4], rev(noabs_ChoSpeed0[spt,,para2,para3,para4]+noabs_stdChoSpeed0[spt,,para2,para3,para4])), border=NA, col=adjustcolor("gray", alpha.f = 0.3))
          
          # legend("bottomleft", legend = c("Choice", "Resistance"), col = c("black", "red"), text.font = 3, lwd = 3, lty=c(1,1))
          # legend("bottomright", legend = c( "Evolving", "Fixed"), col = c("black"), text.font = 3, lwd = 3, lty=c(1,2))
          dev.off()        
        }}}}#}
}


{
  for(para3 in 1:length(Parameter3)){
    for(para4 in 1:length(Parameter4)){
      for(para1 in 1:length(Parameter1)){
        #for(para2 in 1:length(Parameter2)){
        for(spt in 1:length(tTspeed)){
          para2 = 0
          png(paste0(getwd(), "/",Folder,"/", plotFolder, "/",subFolder,"/noabs_SpeedEvolutionTraits_Param1",para1, "_Param3", para3, "_Param4",para4, "_Param2", para2, "_gen",tTspeed[spt], "_paracomb",ParameterCombination,".png"))
          
          par(mar = c(5, 5, 4, 5) + 0.3)
          
          matplot(Parameter2,noabs_ChoSpeed1[spt,para1,,para3,para4],ylim = c(Minspeed,Maxspeed),cex.axis = 2, type = "n", ylab= list("Evolution rate", font = 3, cex = 2),xlab=list("Parameter 2", font = 3, cex = 2))
          lines(Parameter2,noabs_ChoSpeed1[spt,para1,,para3,para4],type = 'l',col="black", lty = 1, lwd = 3);
          lines(Parameter2,noabs_ChoSpeed0[spt,para1,,para3,para4],type = 'l',col="black", lty = 2, lwd = 3);
          
          #----------------- Resistance -----------------
          lines(Parameter2,noabs_ResSpeed1[spt,para1,,para3,para4],type = 'l',col="red", lty = 1, lwd = 3);
          lines(Parameter2,noabs_ResSpeed0[spt,para1,,para3,para4],type = 'l',col="red", lty = 2, lwd = 3);
          
          polygon(x=c(Parameter2, rev(Parameter2)), y=c(noabs_ResSpeed1[spt,para1,,para3,para4]-noabs_stdResSpeed1[spt,para1,,para3,para4], rev(noabs_ResSpeed1[spt,para1,,para3,para4]+noabs_stdResSpeed1[spt,para1,,para3,para4])), border=NA, col=adjustcolor("gray", alpha.f = 0.3))
          polygon(x=c(Parameter2, rev(Parameter2)), y=c(noabs_ResSpeed0[spt,para1,,para3,para4]-noabs_stdResSpeed0[spt,para1,,para3,para4], rev(noabs_ResSpeed0[spt,para1,,para3,para4]+noabs_stdResSpeed0[spt,para1,,para3,para4])), border=NA, col=adjustcolor("gray", alpha.f = 0.3))
          polygon(x=c(Parameter2, rev(Parameter2)), y=c(noabs_ChoSpeed1[spt,para1,,para3,para4]-noabs_stdChoSpeed1[spt,para1,,para3,para4], rev(noabs_ChoSpeed1[spt,para1,,para3,para4]+noabs_stdChoSpeed1[spt,para1,,para3,para4])), border=NA, col=adjustcolor("gray", alpha.f = 0.3))
          polygon(x=c(Parameter2, rev(Parameter2)), y=c(noabs_ChoSpeed0[spt,para1,,para3,para4]-noabs_stdChoSpeed0[spt,para1,,para3,para4], rev(noabs_ChoSpeed0[spt,para1,,para3,para4]+noabs_stdChoSpeed0[spt,para1,,para3,para4])), border=NA, col=adjustcolor("gray", alpha.f = 0.3))
          
          
          # legend("bottomleft", legend = c("Choice", "Resistance"), col = c("black", "red"), text.font = 3, lwd = 3, lty=c(1,1))
          # legend("bottomright", legend = c( "Evolving", "Fixed"), col = c("black"), text.font = 3, lwd = 3, lty=c(1,2))
          dev.off()        
        }}}}#}
}


#------------------ Plant Population size -----------------------

subFolder = "PlantPopSize"

if(!dir.exists(paste0(getwd(), "/",Folder, "/", plotFolder, "/", subFolder))){
  dir.create(paste0(getwd(), "/",Folder, "/", plotFolder, "/", subFolder))
}


{
  #for(para3 in 1:length(Parameter3)){
  para3 = 0
  for(para4 in 1:length(Parameter4)){
    for(para1 in 1:length(Parameter1)){
      for(para2 in 1:length(Parameter2)){
        
        png(paste0(getwd(), "/",Folder,"/", plotFolder, "/",subFolder,"/PlantPopsize_Param1",para1, "_Param3", para3, "_Param4",para4, "_Param2", para2, "_paracomb",ParameterCombination,".png"))
        
        par(mar = c(5, 5, 4, 4) + 0.3)
        plot(0,0,xlim = range(Parameter3),ylim = c(0,max(N1_mean0[tT,para1,para2,,para4], N2_mean0[tT,para1,para2,,para4], N1_mean1[tT,para1,para2,,para4], N2_mean1[tT,para1,para2,,para4], na.rm = TRUE)),cex.axis = 2, type = "n",
             ylab= list("Population size", font = 3, cex = 2),xlab= list("Parameter 3", font = 3, cex = 2))
        lines(Parameter3,N1_mean1[tT,para1,para2,,para4],type = 'l',col="purple",lwd = 3);
        polygon(x=c(Parameter3, rev(Parameter3)), y=c(N1_mean1[tT,para1,para2,,para4]-N1_std1[tT,para1,para2,,para4], rev(N1_mean1[tT,para1,para2,,para4]+N1_std1[tT,para1,para2,,para4])), border=NA, col=adjustcolor("gray", alpha.f = 0.3))
        lines(Parameter3,N1_mean0[tT,para1,para2,,para4],type = 'l',col="gray", lwd = 3);
        polygon(x=c(Parameter3, rev(Parameter3)), y=c(N1_mean0[tT,para1,para2,,para4]-N1_std0[tT,para1,para2,,para4], rev(N1_mean0[tT,para1,para2,,para4]+N1_std0[tT,para1,para2,,para4])), border=NA, col=adjustcolor("gray", alpha.f = 0.3))
        
        lines(Parameter3,N2_mean1[tT,para1,para2,,para4],type = 'l', lty = 2,col="purple", lwd = 3);
        polygon(x=c(Parameter3, rev(Parameter3)), y=c(N2_mean1[tT,para1,para2,,para4]-N2_std1[tT,para1,para2,,para4], rev(N2_mean1[tT,para1,para2,,para4]+N2_std1[tT,para1,para2,,para4])), border=NA, col=adjustcolor("gray", alpha.f = 0.3))
        lines(Parameter3,N2_mean0[tT,para1,para2,,para4],type = 'l',lty = 2,col="gray", lwd = 3);
        polygon(x=c(Parameter3, rev(Parameter3)), y=c(N2_mean0[tT,para1,para2,,para4]-N2_std0[tT,para1,para2,,para4], rev(N2_mean0[tT,para1,para2,,para4]+N2_std0[tT,para1,para2,,para4])), border=NA, col=adjustcolor("gray", alpha.f = 0.3))
        
        legend("bottomright", legend = c("HOST PLANT", "Non-toxic", "Toxic", "BEHAVIOUR", "Evolving",  "Fixed"), col = c(NA,"gray", "purple", NA,1,1), text.font = 3, lwd = 3, lty=c(NA,1,1,NA,1,2), cex=1.5, text.col = c("blue",1,1,"blue",1,1))
        
        dev.off()        
      }}}#}
}


{
  for(para3 in 1:length(Parameter3)){
    #for(para4 in 1:length(Parameter4)){
    para4 = 0
    for(para1 in 1:length(Parameter1)){
      for(para2 in 1:length(Parameter2)){
        
        png(paste0(getwd(), "/",Folder,"/", plotFolder, "/",subFolder,"/PlantPopsize_Param1",para1, "_Param3", para3, "_Param4",para4, "_Param2", para2, "_paracomb",ParameterCombination,".png"))
        
        par(mar = c(5, 5, 4, 4) + 0.3)
        plot(0,0,xlim = range(Parameter4),ylim = c(0,max(N1_mean0[tT,para1,para2,para3,], N2_mean0[tT,para1,para2,para3,], N1_mean1[tT,para1,para2,para3,], N2_mean1[tT,para1,para2,para3,], na.rm = TRUE)),cex.axis = 2, type = "n",
             ylab= list("Population size", font = 3, cex = 2),xlab= list("Parameter 4", font = 3, cex = 2))
        lines(Parameter4,N1_mean1[tT,para1,para2,para3,],type = 'l',col="purple",lwd = 3);
        polygon(x=c(Parameter4, rev(Parameter4)), y=c(N1_mean1[tT,para1,para2,para3,]-N1_std1[tT,para1,para2,para3,], rev(N1_mean1[tT,para1,para2,para3,]+N1_std1[tT,para1,para2,para3,])), border=NA, col=adjustcolor("gray", alpha.f = 0.3))
        lines(Parameter4,N1_mean0[tT,para1,para2,para3,],type = 'l',col="gray", lwd = 3);
        polygon(x=c(Parameter4, rev(Parameter4)), y=c(N1_mean0[tT,para1,para2,para3,]-N1_std0[tT,para1,para2,para3,], rev(N1_mean0[tT,para1,para2,para3,]+N1_std0[tT,para1,para2,para3,])), border=NA, col=adjustcolor("gray", alpha.f = 0.3))
        
        lines(Parameter4,N2_mean1[tT,para1,para2,para3,],type = 'l', lty = 2,col="purple", lwd = 3);
        polygon(x=c(Parameter4, rev(Parameter4)), y=c(N2_mean1[tT,para1,para2,para3,]-N2_std1[tT,para1,para2,para3,], rev(N2_mean1[tT,para1,para2,para3,]+N2_std1[tT,para1,para2,para3,])), border=NA, col=adjustcolor("gray", alpha.f = 0.3))
        lines(Parameter4,N2_mean0[tT,para1,para2,para3,],type = 'l',lty = 2,col="gray", lwd = 3);
        polygon(x=c(Parameter4, rev(Parameter4)), y=c(N2_mean0[tT,para1,para2,para3,]-N2_std0[tT,para1,para2,para3,], rev(N2_mean0[tT,para1,para2,para3,]+N2_std0[tT,para1,para2,para3,])), border=NA, col=adjustcolor("gray", alpha.f = 0.3))
        
        #legend("bottomright", legend = c("HOST PLANT", "Non-toxic", "Toxic", "BEHAVIOUR", "Evolving",  "Fixed"), col = c(NA,"gray", "purple", NA,1,1), text.font = 3, lwd = 3, lty=c(NA,1,1,NA,1,2), cex=1.5, text.col = c("blue",1,1,"blue",1,1))
        
        dev.off()        
      }}}#}
}


{
  for(para3 in 1:length(Parameter3)){
    for(para4 in 1:length(Parameter4)){
      
      #for(para1 in 1:length(Parameter1)){
      para1 = 0
      for(para2 in 1:length(Parameter2)){
        
        png(paste0(getwd(), "/",Folder,"/", plotFolder, "/",subFolder,"/PlantPopsize_Param1",para1, "_Param3", para3, "_Param4",para4, "_Param2", para2, "_paracomb",ParameterCombination,".png"))
        
        par(mar = c(5, 5, 4, 4) + 0.3)
        plot(0,0,xlim = range(Parameter2),ylim = c(0,max(N1_mean0[tT,,para2,para3,para4], N2_mean0[tT,,para2,para3,para4], N1_mean1[tT,,para2,para3,para4], N2_mean1[tT,,para2,para3,para4], na.rm = TRUE)),cex.axis = 2, type = "n",
             ylab= list("Population size", font = 3, cex = 2),xlab= list("Parameter 1", font = 3, cex = 2))
        lines(Parameter2,N1_mean1[tT,,para2,para3,para4],type = 'l',col="purple",lwd = 3);
        polygon(x=c(Parameter2, rev(Parameter2)), y=c(N1_mean1[tT,,para2,para3,para4]-N1_std1[tT,,para2,para3,para4], rev(N1_mean1[tT,,para2,para3,para4]+N1_std1[tT,,para2,para3,para4])), border=NA, col=adjustcolor("gray", alpha.f = 0.3))
        lines(Parameter2,N1_mean0[tT,,para2,para3,para4],type = 'l',col="gray", lwd = 3);
        polygon(x=c(Parameter2, rev(Parameter2)), y=c(N1_mean0[tT,,para2,para3,para4]-N1_std0[tT,,para2,para3,para4], rev(N1_mean0[tT,,para2,para3,para4]+N1_std0[tT,,para2,para3,para4])), border=NA, col=adjustcolor("gray", alpha.f = 0.3))
        
        lines(Parameter2,N2_mean1[tT,,para2,para3,para4],type = 'l', lty = 2,col="purple", lwd = 3);
        polygon(x=c(Parameter2, rev(Parameter2)), y=c(N2_mean1[tT,,para2,para3,para4]-N2_std1[tT,,para2,para3,para4], rev(N2_mean1[tT,,para2,para3,para4]+N2_std1[tT,,para2,para3,para4])), border=NA, col=adjustcolor("gray", alpha.f = 0.3))
        lines(Parameter2,N2_mean0[tT,,para2,para3,para4],type = 'l',lty = 2,col="gray", lwd = 3);
        polygon(x=c(Parameter2, rev(Parameter2)), y=c(N2_mean0[tT,,para2,para3,para4]-N2_std0[tT,,para2,para3,para4], rev(N2_mean0[tT,,para2,para3,para4]+N2_std0[tT,,para2,para3,para4])), border=NA, col=adjustcolor("gray", alpha.f = 0.3))
        
        #legend("bottomright", legend = c("HOST PLANT", "Non-toxic", "Toxic", "BEHAVIOUR", "Evolving",  "Fixed"), col = c(NA,"gray", "purple", NA,1,1), text.font = 3, lwd = 3, lty=c(NA,1,1,NA,1,2), cex=1.5, text.col = c("blue",1,1,"blue",1,1))
        
        dev.off()        
      }}}#}
}


{
  for(para3 in 1:length(Parameter3)){
    for(para4 in 1:length(Parameter4)){
      for(para1 in 1:length(Parameter1)){
        #for(para2 in 1:length(Parameter2)){
        para2 = 0 
        png(paste0(getwd(), "/",Folder,"/", plotFolder, "/",subFolder,"/PlantPopsize_Param1",para1, "_Param3", para3, "_Param4",para4, "_Param2", para2, "_paracomb",ParameterCombination,".png"))
        
        par(mar = c(5, 5, 4, 4) + 0.3)
        plot(0,0,xlim = range(Parameter1),ylim = c(0,max(N1_mean0[tT,para1,,para3,para4], N2_mean0[tT,para1,,para3,para4], N1_mean1[tT,para1,,para3,para4], N2_mean1[tT,para1,,para3,para4], na.rm = TRUE)),cex.axis = 2, type = "n",
             ylab= list("Population size", font = 3, cex = 2),xlab= list("Parameter 2", font = 3, cex = 2))
        lines(Parameter1,N1_mean1[tT,para1,,para3,para4],type = 'l',col="purple",lwd = 3);
        polygon(x=c(Parameter1, rev(Parameter1)), y=c(N1_mean1[tT,para1,,para3,para4]-N1_std1[tT,para1,,para3,para4], rev(N1_mean1[tT,para1,,para3,para4]+N1_std1[tT,para1,,para3,para4])), border=NA, col=adjustcolor("gray", alpha.f = 0.3))
        lines(Parameter1,N1_mean0[tT,para1,,para3,para4],type = 'l',col="gray", lwd = 3);
        polygon(x=c(Parameter1, rev(Parameter1)), y=c(N1_mean0[tT,para1,,para3,para4]-N1_std0[tT,para1,,para3,para4], rev(N1_mean0[tT,para1,,para3,para4]+N1_std0[tT,para1,,para3,para4])), border=NA, col=adjustcolor("gray", alpha.f = 0.3))
        
        lines(Parameter1,N2_mean1[tT,para1,,para3,para4],type = 'l', lty = 2,col="purple", lwd = 3);
        polygon(x=c(Parameter1, rev(Parameter1)), y=c(N2_mean1[tT,para1,,para3,para4]-N2_std1[tT,para1,,para3,para4], rev(N2_mean1[tT,para1,,para3,para4]+N2_std1[tT,para1,,para3,para4])), border=NA, col=adjustcolor("gray", alpha.f = 0.3))
        lines(Parameter1,N2_mean0[tT,para1,,para3,para4],type = 'l',lty = 2,col="gray", lwd = 3);
        polygon(x=c(Parameter1, rev(Parameter1)), y=c(N2_mean0[tT,para1,,para3,para4]-N2_std0[tT,para1,,para3,para4], rev(N2_mean0[tT,para1,,para3,para4]+N2_std0[tT,para1,,para3,para4])), border=NA, col=adjustcolor("gray", alpha.f = 0.3))
        
        #legend("bottomright", legend = c("HOST PLANT", "Non-toxic", "Toxic", "BEHAVIOUR", "Evolving",  "Fixed"), col = c(NA,"gray", "purple", NA,1,1), text.font = 3, lwd = 3, lty=c(NA,1,1,NA,1,2), cex=1.5, text.col = c("blue",1,1,"blue",1,1))
        
        dev.off()        
      }}}#}
}


Folder = "Costs_Allelefrequency" # Folder for storing simulated data
FFolder = Folder

ParameterCombination = 3# Assign different numbers for different parameter combinations to help distinguish the names of plots
Parameter1 = Fecunditycost
Parameter2 = Resistancecost
Parameter3 = Fr0
Parameter4 = Fc0

#------ This folder is for storing plots
plotFolder = "Plots"

if(!dir.exists(paste0(getwd(), "/",Folder, "/", plotFolder))){
  dir.create(paste0(getwd(), "/",Folder, "/", plotFolder))
}

#--------- Loading already analysed data for plotting--------------
DataAnalysis = "AnalysedData"
noabs_stdResSpeed0= readRDS(paste0(getwd(),"/", Folder,"/", DataAnalysis, "/", "noabs_stdResSpeed0.RData"))
noabs_stdResSpeed1= readRDS(paste0(getwd(),"/", Folder,"/", DataAnalysis, "/","noabs_stdResSpeed1.RData"))
noabs_stdChoSpeed0= readRDS(paste0(getwd(),"/", Folder,"/", DataAnalysis, "/","noabs_stdChoSpeed0.RData"))
noabs_stdChoSpeed1= readRDS(paste0(getwd(),"/", Folder,"/", DataAnalysis, "/","noabs_stdChoSpeed1.RData"))

noabs_ResSpeed0= readRDS(paste0(getwd(),"/", Folder,"/", DataAnalysis, "/", "noabs_ResSpeed0.RData"))
noabs_ResSpeed1= readRDS(paste0(getwd(),"/", Folder,"/", DataAnalysis, "/","noabs_ResSpeed1.RData"))
noabs_ChoSpeed0= readRDS(paste0(getwd(),"/", Folder,"/", DataAnalysis, "/","noabs_ChoSpeed0.RData"))
noabs_ChoSpeed1= readRDS(paste0(getwd(),"/", Folder,"/", DataAnalysis, "/","noabs_ChoSpeed1.RData"))


ResSpeed0= readRDS(paste0(getwd(),"/", Folder,"/", DataAnalysis, "/", "ResSpeed0.RData"))
ResSpeed1= readRDS(paste0(getwd(),"/", Folder,"/", DataAnalysis, "/","ResSpeed1.RData"))
ChoSpeed0= readRDS(paste0(getwd(),"/", Folder,"/", DataAnalysis, "/","ChoSpeed0.RData"))
ChoSpeed1= readRDS(paste0(getwd(),"/", Folder,"/", DataAnalysis, "/","ChoSpeed1.RData"))

Rresistancemean1= readRDS(paste0(getwd(),"/", Folder,"/", DataAnalysis, "/", "Rresistancemean1.RData"))
Rresistancesd1 = readRDS(paste0(getwd(),"/", Folder,"/", DataAnalysis, "/","Rresistancesd1.RData"))
Rresistancestd1 = readRDS(paste0(getwd(),"/", Folder,"/", DataAnalysis, "/","Rresistancestd1.RData"))
Cchoicemean1= readRDS(paste0(getwd(),"/", Folder,"/", DataAnalysis, "/","Cchoicemean1.RData"))
Cchoicesd1= readRDS(paste0(getwd(),"/", Folder,"/", DataAnalysis, "/","Cchoicesd1.RData"))
Cchoicestd1= readRDS(paste0(getwd(),"/", Folder,"/", DataAnalysis, "/","Cchoicestd1.RData"))
Rresistancemean0 = readRDS(paste0(getwd(),"/", Folder,"/", DataAnalysis, "/", "Rresistancemean0.RData"))
Rresistancesd0 = readRDS(paste0(getwd(),"/", Folder,"/", DataAnalysis, "/","Rresistancesd0.RData"))
Rresistancestd0 = readRDS(paste0(getwd(),"/", Folder,"/", DataAnalysis, "/","Rresistancestd0.RData"))
Cchoicemean0= readRDS(paste0(getwd(),"/", Folder,"/", DataAnalysis, "/","Cchoicemean0.RData"))
Cchoicesd0= readRDS(paste0(getwd(),"/", Folder,"/", DataAnalysis, "/","Cchoicesd0.RData"))
Cchoicestd0= readRDS(paste0(getwd(),"/", Folder,"/", DataAnalysis, "/","Cchoicestd0.RData"))


Rresistance_popvariancemean1= readRDS(paste0(getwd(),"/", Folder,"/", DataAnalysis, "/", "Rresistance_popvariancemean1.RData"))
Rresistance_popvariancestd1 = readRDS(paste0(getwd(),"/", Folder,"/", DataAnalysis, "/","Rresistance_popvariancestd1.RData"))
Cchoice_popvariancemean1= readRDS(paste0(getwd(),"/", Folder,"/", DataAnalysis, "/","Cchoice_popvariancemean1.RData"))
Cchoice_popvariancestd1= readRDS(paste0(getwd(),"/", Folder,"/", DataAnalysis, "/","Cchoice_popvariancestd1.RData"))
Rresistance_popvariancemean0 = readRDS(paste0(getwd(),"/", Folder,"/", DataAnalysis, "/", "Rresistance_popvariancemean0.RData"))
Rresistance_popvariancestd0 = readRDS(paste0(getwd(),"/", Folder,"/", DataAnalysis, "/","Rresistance_popvariancestd0.RData"))
Cchoice_popvariancemean0= readRDS(paste0(getwd(),"/", Folder,"/", DataAnalysis, "/","Cchoice_popvariancemean0.RData"))
Cchoice_popvariancestd0= readRDS(paste0(getwd(),"/", Folder,"/", DataAnalysis, "/","Cchoice_popvariancestd0.RData"))

N1_mean1= readRDS(paste0(getwd(),"/", Folder,"/", DataAnalysis, "/", "N1_mean1.RData"))
N1_mean0= readRDS(paste0(getwd(),"/", Folder,"/", DataAnalysis, "/","N1_mean0.RData"))
N1_std1= readRDS(paste0(getwd(),"/", Folder,"/", DataAnalysis, "/","N1_std1.RData"))
N1_std0= readRDS(paste0(getwd(),"/", Folder,"/", DataAnalysis, "/","N1_std0.RData"))
N2_mean1= readRDS(paste0(getwd(),"/", Folder,"/", DataAnalysis, "/", "N2_mean1.RData"))
N2_mean0= readRDS(paste0(getwd(),"/", Folder,"/", DataAnalysis, "/","N2_mean0.RData"))
N2_std1= readRDS(paste0(getwd(),"/", Folder,"/", DataAnalysis, "/","N2_std1.RData"))
N2_std0= readRDS(paste0(getwd(),"/", Folder,"/", DataAnalysis, "/","N2_std0.RData"))


tttt = targetTtimes[length(targetTtimes)]
ttt = targetTtimes[1]
plottargetTtimes = c(ttt, tttt)


#------------------- Preference - Resistance Population replicates 

subFolder = "Resistance_Preference" # For storing mean traits and population trait variance plots-----------

if(!dir.exists(paste0(getwd(), "/",Folder,"/",plotFolder,"/",subFolder))){
  dir.create(paste0(getwd(), "/",Folder,"/",plotFolder,"/",subFolder))
}


tT = length(targetTtimes)
{
  #for(para3 in 1:length(Parameter3)){
  para3 = 0
  for(para4 in 1:length(Parameter4)){
    for(para1 in 1:length(Parameter1)){
      for(para2 in 1:length(Parameter2)){
        
        png(paste0(getwd(), "/",Folder,"/", plotFolder, "/",subFolder,"/meanTraits_Param1",para1, "_Param3", para3, "_Param4",para4, "_Param2", para2, "_paracomb",ParameterCombination,".png"))
        
        par(mar = c(5, 5, 4, 5) + 0.3)
        
        matplot(Parameter3,Cchoicemean1[tT,para1,para2,,para4],ylim = c(0,1),cex.axis = 2, type = "n", ylab= list("Mean trait", font = 3, cex = 2),xlab=list("Parameter 3", font = 3, cex = 2))
        lines(Parameter3,Cchoicemean1[tT,para1,para2,,para4],type = 'l',col="black", lty = 1, lwd = 3);
        lines(Parameter3,Cchoicemean0[tT,para1,para2,,para4],type = 'l',col="black", lty = 2, lwd = 3);
        
        #----------------- Resistance -----------------
        lines(Parameter3,Rresistancemean1[tT,para1,para2,,para4],type = 'l',col="red", lty = 1, lwd = 3);
        lines(Parameter3,Rresistancemean0[tT,para1,para2,,para4],type = 'l',col="red", lty = 2, lwd = 3);
        #------- Add std errors ----------------
        polygon(x=c(Parameter3, rev(Parameter3)), y=c(Rresistancemean1[tT,para1,para2,,para4]-Rresistancestd1[tT,para1,para2,,para4], rev(Rresistancemean1[tT,para1,para2,,para4]+Rresistancestd1[tT,para1,para2,,para4])), border=NA, col=adjustcolor("gray", alpha.f = 0.3))
        polygon(x=c(Parameter3, rev(Parameter3)), y=c(Rresistancemean0[tT,para1,para2,,para4]-Rresistancestd0[tT,para1,para2,,para4], rev(Rresistancemean0[tT,para1,para2,,para4]+Rresistancestd0[tT,para1,para2,,para4])), border=NA, col=adjustcolor("gray", alpha.f = 0.3))
        polygon(x=c(Parameter3, rev(Parameter3)), y=c(Cchoicemean1[tT,para1,para2,,para4]-Cchoicestd1[tT,para1,para2,,para4], rev(Cchoicemean1[tT,para1,para2,,para4]+Cchoicestd1[tT,para1,para2,,para4])), border=NA, col=adjustcolor("gray", alpha.f = 0.3))
        polygon(x=c(Parameter3, rev(Parameter3)), y=c(Cchoicemean0[tT,para1,para2,,para4]-Cchoicestd0[tT,para1,para2,,para4], rev(Cchoicemean0[tT,para1,para2,,para4]+Cchoicestd0[tT,para1,para2,,para4])), border=NA, col=adjustcolor("gray", alpha.f = 0.3))
        
        legend("bottomleft", legend = c("Trait", "Choice", "Resistance"), col = c(NA,"black", "red"), text.font = 3, lwd = 3, lty=c(1,1,1))
        legend("bottomright", legend = c("Behaviour", "Evolving", "Fixed"), col = c(NA,"black", "black"), text.font = 3, lwd = 3, lty=c(1,1,2))
        dev.off()        
      }}}#}
}


{
  for(para3 in 1:length(Parameter3)){
    #for(para4 in 1:length(Parameter4)){
    para4 = 0
    for(para1 in 1:length(Parameter1)){
      for(para2 in 1:length(Parameter2)){
        
        png(paste0(getwd(), "/",Folder,"/", plotFolder, "/",subFolder,"/meanTraits_Param1",para1, "_Param3", para3, "_Param4",para4, "_Param2", para2, "_paracomb",ParameterCombination,".png"))
        
        par(mar = c(5, 5, 4, 5) + 0.3)
        
        matplot(Parameter4,Cchoicemean1[tT,para1,para2,para3,],ylim = c(0,1),cex.axis = 2, type = "n", ylab= list("Mean trait", font = 3, cex = 2),xlab=list("Parameter 4", font = 3, cex = 2))
        lines(Parameter4,Cchoicemean1[tT,para1,para2,para3,],type = 'l',col="black", lty = 1, lwd = 3);
        lines(Parameter4,Cchoicemean0[tT,para1,para2,para3,],type = 'l',col="black", lty = 2, lwd = 3);
        
        #----------------- Resistance -----------------
        lines(Parameter4,Rresistancemean1[tT,para1,para2,para3,],type = 'l',col="red", lty = 1, lwd = 3);
        lines(Parameter4,Rresistancemean0[tT,para1,para2,para3,],type = 'l',col="red", lty = 2, lwd = 3);
        
        polygon(x=c(Parameter4, rev(Parameter4)), y=c(Rresistancemean1[tT,para1,para2,para3,]-Rresistancestd1[tT,para1,para2,para3,], rev(Rresistancemean1[tT,para1,para2,para3,]+Rresistancestd1[tT,para1,para2,para3,])), border=NA, col=adjustcolor("gray", alpha.f = 0.3))
        polygon(x=c(Parameter4, rev(Parameter4)), y=c(Rresistancemean0[tT,para1,para2,para3,]-Rresistancestd0[tT,para1,para2,para3,], rev(Rresistancemean0[tT,para1,para2,para3,]+Rresistancestd0[tT,para1,para2,para3,])), border=NA, col=adjustcolor("gray", alpha.f = 0.3))
        polygon(x=c(Parameter4, rev(Parameter4)), y=c(Cchoicemean1[tT,para1,para2,para3,]-Cchoicestd1[tT,para1,para2,para3,], rev(Cchoicemean1[tT,para1,para2,para3,]+Cchoicestd1[tT,para1,para2,para3,])), border=NA, col=adjustcolor("gray", alpha.f = 0.3))
        polygon(x=c(Parameter4, rev(Parameter4)), y=c(Cchoicemean0[tT,para1,para2,para3,]-Cchoicestd0[tT,para1,para2,para3,], rev(Cchoicemean0[tT,para1,para2,para3,]+Cchoicestd0[tT,para1,para2,para3,])), border=NA, col=adjustcolor("gray", alpha.f = 0.3))
        
        # legend("bottomleft", legend = c("Choice", "Resistance"), col = c("black", "red"), text.font = 3, lwd = 3, lty=c(1,1))
        # legend("bottomright", legend = c( "Evolving", "Fixed"), col = c("black"), text.font = 3, lwd = 3, lty=c(1,2))
        dev.off()        
      }}}#}
}


{
  for(para3 in 1:length(Parameter3)){
    for(para4 in 1:length(Parameter4)){
      #for(para1 in 1:length(Parameter1)){
      para1 = 0
      for(para2 in 1:length(Parameter2)){
        
        png(paste0(getwd(), "/",Folder,"/", plotFolder, "/",subFolder,"/meanTraits_Param1",para1, "_Param3", para3, "_Param4",para4, "_Param2", para2, "_paracomb",ParameterCombination,".png"))
        
        par(mar = c(5, 5, 4, 5) + 0.3)
        
        matplot(Parameter1,Cchoicemean1[tT,,para2,para3,para4],ylim = c(0,1),cex.axis = 2, type = "n", ylab= list("Mean trait", font = 3, cex = 2),xlab=list("Parameter 1", font = 3, cex = 2))
        lines(Parameter1,Cchoicemean1[tT,,para2,para3,para4],type = 'l',col="black", lty = 1, lwd = 3);
        lines(Parameter1,Cchoicemean0[tT,,para2,para3,para4],type = 'l',col="black", lty = 2, lwd = 3);
        
        #----------------- Resistance -----------------
        lines(Parameter1,Rresistancemean1[tT,,para2,para3,para4],type = 'l',col="red", lty = 1, lwd = 3);
        lines(Parameter1,Rresistancemean0[tT,,para2,para3,para4],type = 'l',col="red", lty = 2, lwd = 3);
        
        polygon(x=c(Parameter1, rev(Parameter1)), y=c(Rresistancemean1[tT,,para2,para3,para4]-Rresistancestd1[tT,,para2,para3,para4], rev(Rresistancemean1[tT,,para2,para3,para4]+Rresistancestd1[tT,,para2,para3,para4])), border=NA, col=adjustcolor("gray", alpha.f = 0.3))
        polygon(x=c(Parameter1, rev(Parameter1)), y=c(Rresistancemean0[tT,,para2,para3,para4]-Rresistancestd0[tT,,para2,para3,para4], rev(Rresistancemean0[tT,,para2,para3,para4]+Rresistancestd0[tT,,para2,para3,para4])), border=NA, col=adjustcolor("gray", alpha.f = 0.3))
        polygon(x=c(Parameter1, rev(Parameter1)), y=c(Cchoicemean1[tT,,para2,para3,para4]-Cchoicestd1[tT,,para2,para3,para4], rev(Cchoicemean1[tT,,para2,para3,para4]+Cchoicestd1[tT,,para2,para3,para4])), border=NA, col=adjustcolor("gray", alpha.f = 0.3))
        polygon(x=c(Parameter1, rev(Parameter1)), y=c(Cchoicemean0[tT,,para2,para3,para4]-Cchoicestd0[tT,,para2,para3,para4], rev(Cchoicemean0[tT,,para2,para3,para4]+Cchoicestd0[tT,,para2,para3,para4])), border=NA, col=adjustcolor("gray", alpha.f = 0.3))
        
        # legend("bottomleft", legend = c("Choice", "Resistance"), col = c("black", "red"), text.font = 3, lwd = 3, lty=c(1,1))
        # legend("bottomright", legend = c( "Evolving", "Fixed"), col = c("black"), text.font = 3, lwd = 3, lty=c(1,2))
        dev.off()        
      }}}#}
}


{
  for(para3 in 1:length(Parameter3)){
    for(para4 in 1:length(Parameter4)){
      for(para1 in 1:length(Parameter1)){
        #for(para2 in 1:length(Parameter2)){
        para2 = 0
        png(paste0(getwd(), "/",Folder,"/", plotFolder, "/",subFolder,"/meanTraits_Param1",para1, "_Param3", para3, "_Param4",para4, "_Param2", para2, "_paracomb",ParameterCombination,".png"))
        
        par(mar = c(5, 5, 4, 5) + 0.3)
        
        matplot(Parameter2,Cchoicemean1[tT,para1,,para3,para4],ylim = c(0,1),cex.axis = 2, type = "n", ylab= list("Mean trait", font = 3, cex = 2),xlab=list("Parameter 2", font = 3, cex = 2))
        lines(Parameter2,Cchoicemean1[tT,para1,,para3,para4],type = 'l',col="black", lty = 1, lwd = 3);
        lines(Parameter2,Cchoicemean0[tT,para1,,para3,para4],type = 'l',col="black", lty = 2, lwd = 3);
        
        #----------------- Resistance -----------------
        lines(Parameter2,Rresistancemean1[tT,para1,,para3,para4],type = 'l',col="red", lty = 1, lwd = 3);
        lines(Parameter2,Rresistancemean0[tT,para1,,para3,para4],type = 'l',col="red", lty = 2, lwd = 3);
        
        polygon(x=c(Parameter2, rev(Parameter2)), y=c(Rresistancemean1[tT,para1,,para3,para4]-Rresistancestd1[tT,para1,,para3,para4], rev(Rresistancemean1[tT,para1,,para3,para4]+Rresistancestd1[tT,para1,,para3,para4])), border=NA, col=adjustcolor("gray", alpha.f = 0.3))
        polygon(x=c(Parameter2, rev(Parameter2)), y=c(Rresistancemean0[tT,para1,,para3,para4]-Rresistancestd0[tT,para1,,para3,para4], rev(Rresistancemean0[tT,para1,,para3,para4]+Rresistancestd0[tT,para1,,para3,para4])), border=NA, col=adjustcolor("gray", alpha.f = 0.3))
        polygon(x=c(Parameter2, rev(Parameter2)), y=c(Cchoicemean1[tT,para1,,para3,para4]-Cchoicestd1[tT,para1,,para3,para4], rev(Cchoicemean1[tT,para1,,para3,para4]+Cchoicestd1[tT,para1,,para3,para4])), border=NA, col=adjustcolor("gray", alpha.f = 0.3))
        polygon(x=c(Parameter2, rev(Parameter2)), y=c(Cchoicemean0[tT,para1,,para3,para4]-Cchoicestd0[tT,para1,,para3,para4], rev(Cchoicemean0[tT,para1,,para3,para4]+Cchoicestd0[tT,para1,,para3,para4])), border=NA, col=adjustcolor("gray", alpha.f = 0.3))
        
        # legend("bottomleft", legend = c("Choice", "Resistance"), col = c("black", "red"), text.font = 3, lwd = 3, lty=c(1,1))
        # legend("bottomright", legend = c( "Evolving", "Fixed"), col = c("black"), text.font = 3, lwd = 3, lty=c(1,2))
        dev.off()        
      }}}#}
}


#----------------- mean trait Variance within populations --------------
Minvar = min(c(Cchoice_popvariancemean1,Cchoice_popvariancemean0, Rresistance_popvariancemean0, Rresistance_popvariancemean1), na.rm = T)
Maxvar = max(c(Cchoice_popvariancemean1,Cchoice_popvariancemean0, Rresistance_popvariancemean0, Rresistance_popvariancemean1), na.rm = T)
{
  #for(para3 in 1:length(Parameter3)){
  para3 = 0
  for(para4 in 1:length(Parameter4)){
    for(para1 in 1:length(Parameter1)){
      for(para2 in 1:length(Parameter2)){
        
        png(paste0(getwd(), "/",Folder,"/", plotFolder, "/",subFolder,"/VarianceTraits_Param1",para1, "_Param3", para3, "_Param4",para4, "_Param2", para2, "_paracomb",ParameterCombination,".png"))
        
        par(mar = c(5, 5, 4, 5) + 0.3)
        
        matplot(Parameter3,Cchoice_popvariancemean1[tT,para1,para2,,para4],ylim = c(Minvar,Maxvar),cex.axis = 2, type = "n", ylab= list("Population trait variance", font = 3, cex = 2),xlab=list("Parameter 3", font = 3, cex = 2))
        lines(Parameter3,Cchoice_popvariancemean1[tT,para1,para2,,para4],type = 'l',col="black", lty = 1, lwd = 3);
        lines(Parameter3,Cchoice_popvariancemean0[tT,para1,para2,,para4],type = 'l',col="black", lty = 2, lwd = 3);
        
        
        #----------------- Resistance -----------------
        lines(Parameter3,Rresistance_popvariancemean1[tT,para1,para2,,para4],type = 'l',col="red", lty = 1, lwd = 3);
        lines(Parameter3,Rresistance_popvariancemean0[tT,para1,para2,,para4],type = 'l',col="red", lty = 2, lwd = 3);
        
        polygon(x=c(Parameter3, rev(Parameter3)), y=c(Rresistance_popvariancemean1[tT,para1,para2,,para4]-Rresistance_popvariancestd1[tT,para1,para2,,para4], rev(Rresistance_popvariancemean1[tT,para1,para2,,para4]+Rresistance_popvariancestd1[tT,para1,para2,,para4])), border=NA, col=adjustcolor("gray", alpha.f = 0.3))
        polygon(x=c(Parameter3, rev(Parameter3)), y=c(Rresistance_popvariancemean0[tT,para1,para2,,para4]-Rresistance_popvariancestd0[tT,para1,para2,,para4], rev(Rresistance_popvariancemean0[tT,para1,para2,,para4]+Rresistance_popvariancestd0[tT,para1,para2,,para4])), border=NA, col=adjustcolor("gray", alpha.f = 0.3))
        polygon(x=c(Parameter3, rev(Parameter3)), y=c(Cchoice_popvariancemean1[tT,para1,para2,,para4]-Cchoice_popvariancestd1[tT,para1,para2,,para4], rev(Cchoice_popvariancemean1[tT,para1,para2,,para4]+Cchoice_popvariancestd1[tT,para1,para2,,para4])), border=NA, col=adjustcolor("gray", alpha.f = 0.3))
        polygon(x=c(Parameter3, rev(Parameter3)), y=c(Cchoice_popvariancemean0[tT,para1,para2,,para4]-Cchoice_popvariancestd0[tT,para1,para2,,para4], rev(Cchoice_popvariancemean0[tT,para1,para2,,para4]+Cchoice_popvariancestd0[tT,para1,para2,,para4])), border=NA, col=adjustcolor("gray", alpha.f = 0.3))
        
        
        legend("bottomleft", legend = c("Trait", "Choice", "Resistance"), col = c(NA,"black", "red"), text.font = 3, lwd = 3, lty=c(1,1,1))
        legend("bottomright", legend = c("Behaviour", "Evolving", "Fixed"), col = c(NA,"black", "black"), text.font = 3, lwd = 3, lty=c(1,1,2))
        dev.off()        
      }}}#}
}


{
  for(para3 in 1:length(Parameter3)){
    #for(para4 in 1:length(Parameter4)){
    para4 = 0
    for(para1 in 1:length(Parameter1)){
      for(para2 in 1:length(Parameter2)){
        
        png(paste0(getwd(), "/",Folder,"/", plotFolder, "/",subFolder,"/VarianceTraits_Param1",para1, "_Param3", para3, "_Param4",para4, "_Param2", para2, "_paracomb",ParameterCombination,".png"))
        
        par(mar = c(5, 5, 4, 5) + 0.3)
        
        matplot(Parameter4,Cchoice_popvariancemean1[tT,para1,para2,para3,],ylim = c(Minvar,Maxvar),cex.axis = 2, type = "n", ylab= list("Population trait variance", font = 3, cex = 2),xlab=list("Parameter 4", font = 3, cex = 2))
        lines(Parameter4,Cchoice_popvariancemean1[tT,para1,para2,para3,],type = 'l',col="black", lty = 1, lwd = 3);
        lines(Parameter4,Cchoice_popvariancemean0[tT,para1,para2,para3,],type = 'l',col="black", lty = 2, lwd = 3);
        
        #----------------- Resistance -----------------
        lines(Parameter4,Rresistance_popvariancemean1[tT,para1,para2,para3,],type = 'l',col="red", lty = 1, lwd = 3);
        lines(Parameter4,Rresistance_popvariancemean0[tT,para1,para2,para3,],type = 'l',col="red", lty = 2, lwd = 3);
        
        polygon(x=c(Parameter4, rev(Parameter4)), y=c(Rresistance_popvariancemean1[tT,para1,para2,para3,]-Rresistance_popvariancestd1[tT,para1,para2,para3,], rev(Rresistance_popvariancemean1[tT,para1,para2,para3,]+Rresistance_popvariancestd1[tT,para1,para2,para3,])), border=NA, col=adjustcolor("gray", alpha.f = 0.3))
        polygon(x=c(Parameter4, rev(Parameter4)), y=c(Rresistance_popvariancemean0[tT,para1,para2,para3,]-Rresistance_popvariancestd0[tT,para1,para2,para3,], rev(Rresistance_popvariancemean0[tT,para1,para2,para3,]+Rresistance_popvariancestd0[tT,para1,para2,para3,])), border=NA, col=adjustcolor("gray", alpha.f = 0.3))
        polygon(x=c(Parameter4, rev(Parameter4)), y=c(Cchoice_popvariancemean1[tT,para1,para2,para3,]-Cchoice_popvariancestd1[tT,para1,para2,para3,], rev(Cchoice_popvariancemean1[tT,para1,para2,para3,]+Cchoice_popvariancestd1[tT,para1,para2,para3,])), border=NA, col=adjustcolor("gray", alpha.f = 0.3))
        polygon(x=c(Parameter4, rev(Parameter4)), y=c(Cchoice_popvariancemean0[tT,para1,para2,para3,]-Cchoice_popvariancestd0[tT,para1,para2,para3,], rev(Cchoice_popvariancemean0[tT,para1,para2,para3,]+Cchoice_popvariancestd0[tT,para1,para2,para3,])), border=NA, col=adjustcolor("gray", alpha.f = 0.3))
        
        # legend("bottomleft", legend = c("Choice", "Resistance"), col = c("black", "red"), text.font = 3, lwd = 3, lty=c(1,1))
        # legend("bottomright", legend = c( "Evolving", "Fixed"), col = c("black"), text.font = 3, lwd = 3, lty=c(1,2))
        dev.off()        
      }}}#}
}


{
  for(para3 in 1:length(Parameter3)){
    for(para4 in 1:length(Parameter4)){
      #for(para1 in 1:length(Parameter1)){
      para1 = 0
      for(para2 in 1:length(Parameter2)){
        
        png(paste0(getwd(), "/",Folder,"/", plotFolder, "/",subFolder,"/VarianceTraits_Param1",para1, "_Param3", para3, "_Param4",para4, "_Param2", para2, "_paracomb",ParameterCombination,".png"))
        par(mar = c(5, 5, 4, 5) + 0.3)
        
        matplot(Parameter1,Cchoice_popvariancemean1[tT,,para2,para3,para4],ylim = c(Minvar,Maxvar),cex.axis = 2, type = "n", ylab= list("Population trait variance", font = 3, cex = 2),xlab=list("Parameter 1", font = 3, cex = 2))
        lines(Parameter1,Cchoice_popvariancemean1[tT,,para2,para3,para4],type = 'l',col="black", lty = 1, lwd = 3);
        lines(Parameter1,Cchoice_popvariancemean0[tT,,para2,para3,para4],type = 'l',col="black", lty = 2, lwd = 3);
        
        #----------------- Resistance -----------------
        lines(Parameter1,Rresistance_popvariancemean1[tT,,para2,para3,para4],type = 'l',col="red", lty = 1, lwd = 3);
        lines(Parameter1,Rresistance_popvariancemean0[tT,,para2,para3,para4],type = 'l',col="red", lty = 2, lwd = 3);
        
        polygon(x=c(Parameter1, rev(Parameter1)), y=c(Rresistance_popvariancemean1[tT,,para2,para3,para4]-Rresistance_popvariancestd1[tT,,para2,para3,para4], rev(Rresistance_popvariancemean1[tT,,para2,para3,para4]+Rresistance_popvariancestd1[tT,,para2,para3,para4])), border=NA, col=adjustcolor("gray", alpha.f = 0.3))
        polygon(x=c(Parameter1, rev(Parameter1)), y=c(Rresistance_popvariancemean0[tT,,para2,para3,para4]-Rresistance_popvariancestd0[tT,,para2,para3,para4], rev(Rresistance_popvariancemean0[tT,,para2,para3,para4]+Rresistance_popvariancestd0[tT,,para2,para3,para4])), border=NA, col=adjustcolor("gray", alpha.f = 0.3))
        polygon(x=c(Parameter1, rev(Parameter1)), y=c(Cchoice_popvariancemean1[tT,,para2,para3,para4]-Cchoice_popvariancestd1[tT,,para2,para3,para4], rev(Cchoice_popvariancemean1[tT,,para2,para3,para4]+Cchoice_popvariancestd1[tT,,para2,para3,para4])), border=NA, col=adjustcolor("gray", alpha.f = 0.3))
        polygon(x=c(Parameter1, rev(Parameter1)), y=c(Cchoice_popvariancemean0[tT,,para2,para3,para4]-Cchoice_popvariancestd0[tT,,para2,para3,para4], rev(Cchoice_popvariancemean0[tT,,para2,para3,para4]+Cchoice_popvariancestd0[tT,,para2,para3,para4])), border=NA, col=adjustcolor("gray", alpha.f = 0.3))
        
        # legend("bottomleft", legend = c("Choice", "Resistance"), col = c("black", "red"), text.font = 3, lwd = 3, lty=c(1,1))
        # legend("bottomright", legend = c( "Evolving", "Fixed"), col = c("black"), text.font = 3, lwd = 3, lty=c(1,2))
        dev.off()        
      }}}#}
}


{
  for(para3 in 1:length(Parameter3)){
    for(para4 in 1:length(Parameter4)){
      for(para1 in 1:length(Parameter1)){
        #for(para2 in 1:length(Parameter2)){
        para2 = 0
        png(paste0(getwd(), "/",Folder,"/", plotFolder, "/",subFolder,"/VarianceTraits_Param1",para1, "_Param3", para3, "_Param4",para4, "_Param2", para2, "_paracomb",ParameterCombination,".png"))
        
        par(mar = c(5, 5, 4, 5) + 0.3)
        
        matplot(Parameter2,Cchoice_popvariancemean1[tT,para1,,para3,para4],ylim = c(Minvar,Maxvar),cex.axis = 2, type = "n", ylab= list("Population trait variance", font = 3, cex = 2),xlab=list("Parameter 2", font = 3, cex = 2))
        lines(Parameter2,Cchoice_popvariancemean1[tT,para1,,para3,para4],type = 'l',col="black", lty = 1, lwd = 3);
        lines(Parameter2,Cchoice_popvariancemean0[tT,para1,,para3,para4],type = 'l',col="black", lty = 2, lwd = 3);
        
        #----------------- Resistance -----------------
        lines(Parameter2,Rresistance_popvariancemean1[tT,para1,,para3,para4],type = 'l',col="red", lty = 1, lwd = 3);
        lines(Parameter2,Rresistance_popvariancemean0[tT,para1,,para3,para4],type = 'l',col="red", lty = 2, lwd = 3);
        
        polygon(x=c(Parameter2, rev(Parameter2)), y=c(Rresistance_popvariancemean1[tT,para1,,para3,para4]-Rresistance_popvariancestd1[tT,para1,,para3,para4], rev(Rresistance_popvariancemean1[tT,para1,,para3,para4]+Rresistance_popvariancestd1[tT,para1,,para3,para4])), border=NA, col=adjustcolor("gray", alpha.f = 0.3))
        polygon(x=c(Parameter2, rev(Parameter2)), y=c(Rresistance_popvariancemean0[tT,para1,,para3,para4]-Rresistance_popvariancestd0[tT,para1,,para3,para4], rev(Rresistance_popvariancemean0[tT,para1,,para3,para4]+Rresistance_popvariancestd0[tT,para1,,para3,para4])), border=NA, col=adjustcolor("gray", alpha.f = 0.3))
        polygon(x=c(Parameter2, rev(Parameter2)), y=c(Cchoice_popvariancemean1[tT,para1,,para3,para4]-Cchoice_popvariancestd1[tT,para1,,para3,para4], rev(Cchoice_popvariancemean1[tT,para1,,para3,para4]+Cchoice_popvariancestd1[tT,para1,,para3,para4])), border=NA, col=adjustcolor("gray", alpha.f = 0.3))
        polygon(x=c(Parameter2, rev(Parameter2)), y=c(Cchoice_popvariancemean0[tT,para1,,para3,para4]-Cchoice_popvariancestd0[tT,para1,,para3,para4], rev(Cchoice_popvariancemean0[tT,para1,,para3,para4]+Cchoice_popvariancestd0[tT,para1,,para3,para4])), border=NA, col=adjustcolor("gray", alpha.f = 0.3))
        
        
        # legend("bottomleft", legend = c("Choice", "Resistance"), col = c("black", "red"), text.font = 3, lwd = 3, lty=c(1,1))
        # legend("bottomright", legend = c( "Evolving", "Fixed"), col = c("black"), text.font = 3, lwd = 3, lty=c(1,2))
        dev.off()        
      }}}#}
}

# ------------------- Evolution rate ----------
subFolder = "SpeedResistance_Preference"

if(!dir.exists(paste0(getwd(), "/",Folder, "/", plotFolder, "/", subFolder))){
  dir.create(paste0(getwd(), "/",Folder, "/", plotFolder, "/", subFolder))
}


#----------------- Speed of adaptation --------------
Minspeed = min(c(ChoSpeed1,ChoSpeed0, ResSpeed0, ResSpeed1), na.rm = T)
Maxspeed = max(c(ChoSpeed1,ChoSpeed0, ResSpeed0, ResSpeed1), na.rm = T)
{
  #for(para3 in 1:length(Parameter3)){
  para3 = 0
  for(para4 in 1:length(Parameter4)){
    for(para1 in 1:length(Parameter1)){
      for(para2 in 1:length(Parameter2)){
        for(spt in 1:length(tTspeed)){
          png(paste0(getwd(), "/",Folder,"/", plotFolder, "/",subFolder,"/abs_SpeedEvolutionTraits_Param1",para1, "_Param3", para3, "_Param4",para4, "_Param2", para2, "_gen",tTspeed[spt], "_paracomb",ParameterCombination,".png"))
          
          par(mar = c(5, 5, 4, 5) + 0.3)
          
          matplot(Parameter3,ChoSpeed1[spt,para1,para2,,para4],ylim = c(Minspeed,Maxspeed),cex.axis = 2, type = "n", ylab= list("Evolution rate", font = 3, cex = 2),xlab=list("Parameter 3", font = 3, cex = 2))
          lines(Parameter3,ChoSpeed1[spt,para1,para2,,para4],type = 'l',col="black", lty = 1, lwd = 3);
          lines(Parameter3,ChoSpeed0[spt,para1,para2,,para4],type = 'l',col="black", lty = 2, lwd = 3);
          
          #----------------- Resistance -----------------
          lines(Parameter3,ResSpeed1[spt,para1,para2,,para4],type = 'l',col="red", lty = 1, lwd = 3);
          lines(Parameter3,ResSpeed0[spt,para1,para2,,para4],type = 'l',col="red", lty = 2, lwd = 3);
          
          legend("bottomleft", legend = c("Trait", "Choice", "Resistance"), col = c(NA,"black", "red"), text.font = 3, lwd = 3, lty=c(1,1,1))
          legend("bottomright", legend = c("Behaviour", "Evolving", "Fixed"), col = c(NA,"black", "black"), text.font = 3, lwd = 3, lty=c(1,1,2))
          dev.off()        
        }}}}#}
}


{
  for(para3 in 1:length(Parameter3)){
    #for(para4 in 1:length(Parameter4)){
    para4 = 0
    for(para1 in 1:length(Parameter1)){
      for(para2 in 1:length(Parameter2)){
        for(spt in 1:length(tTspeed)){
          png(paste0(getwd(), "/",Folder,"/", plotFolder, "/",subFolder,"/abs_SpeedEvolutionTraits_Param1",para1, "_Param3", para3, "_Param4",para4, "_Param2", para2, "_gen",tTspeed[spt], "_paracomb",ParameterCombination,".png"))
          
          par(mar = c(5, 5, 4, 5) + 0.3)
          
          matplot(Parameter4,ChoSpeed1[spt,para1,para2,para3,],ylim = c(Minspeed,Maxspeed),cex.axis = 2, type = "n", ylab= list("Evolution rate", font = 3, cex = 2),xlab=list("Parameter 4", font = 3, cex = 2))
          lines(Parameter4,ChoSpeed1[spt,para1,para2,para3,],type = 'l',col="black", lty = 1, lwd = 3);
          lines(Parameter4,ChoSpeed0[spt,para1,para2,para3,],type = 'l',col="black", lty = 2, lwd = 3);
          
          #----------------- Resistance -----------------
          lines(Parameter4,ResSpeed1[spt,para1,para2,para3,],type = 'l',col="red", lty = 1, lwd = 3);
          lines(Parameter4,ResSpeed0[spt,para1,para2,para3,],type = 'l',col="red", lty = 2, lwd = 3);
          # legend("bottomleft", legend = c("Choice", "Resistance"), col = c("black", "red"), text.font = 3, lwd = 3, lty=c(1,1))
          # legend("bottomright", legend = c( "Evolving", "Fixed"), col = c("black"), text.font = 3, lwd = 3, lty=c(1,2))
          dev.off()        
        }}}}#}
}


{
  for(para3 in 1:length(Parameter3)){
    for(para4 in 1:length(Parameter4)){
      #for(para1 in 1:length(Parameter1)){
      para1 = 0
      for(para2 in 1:length(Parameter2)){
        for(spt in 1:length(tTspeed)){
          png(paste0(getwd(), "/",Folder,"/", plotFolder, "/",subFolder,"/abs_SpeedEvolutionTraits_Param1",para1, "_Param3", para3, "_Param4",para4, "_Param2", para2, "_gen",tTspeed[spt], "_paracomb",ParameterCombination,".png"))
          par(mar = c(5, 5, 4, 5) + 0.3)
          
          matplot(Parameter1,ChoSpeed1[spt,,para2,para3,para4],ylim = c(Minspeed,Maxspeed),cex.axis = 2, type = "n", ylab= list("Evolution rate", font = 3, cex = 2),xlab=list("Parameter 1", font = 3, cex = 2))
          lines(Parameter1,ChoSpeed1[spt,,para2,para3,para4],type = 'l',col="black", lty = 1, lwd = 3);
          lines(Parameter1,ChoSpeed0[spt,,para2,para3,para4],type = 'l',col="black", lty = 2, lwd = 3);
          
          #----------------- Resistance -----------------
          lines(Parameter1,ResSpeed1[spt,,para2,para3,para4],type = 'l',col="red", lty = 1, lwd = 3);
          lines(Parameter1,ResSpeed0[spt,,para2,para3,para4],type = 'l',col="red", lty = 2, lwd = 3);
          
          # legend("bottomleft", legend = c("Choice", "Resistance"), col = c("black", "red"), text.font = 3, lwd = 3, lty=c(1,1))
          # legend("bottomright", legend = c( "Evolving", "Fixed"), col = c("black"), text.font = 3, lwd = 3, lty=c(1,2))
          dev.off()        
        }}}}#}
}


{
  for(para3 in 1:length(Parameter3)){
    for(para4 in 1:length(Parameter4)){
      for(para1 in 1:length(Parameter1)){
        #for(para2 in 1:length(Parameter2)){
        for(spt in 1:length(tTspeed)){
          para2 = 0
          png(paste0(getwd(), "/",Folder,"/", plotFolder, "/",subFolder,"/abs_SpeedEvolutionTraits_Param1",para1, "_Param3", para3, "_Param4",para4, "_Param2", para2, "_gen",tTspeed[spt], "_paracomb",ParameterCombination,".png"))
          
          par(mar = c(5, 5, 4, 5) + 0.3)
          
          matplot(Parameter2,ChoSpeed1[spt,para1,,para3,para4],ylim = c(Minspeed,Maxspeed),cex.axis = 2, type = "n", ylab= list("Evolution rate", font = 3, cex = 2),xlab=list("Parameter 2", font = 3, cex = 2))
          lines(Parameter2,ChoSpeed1[spt,para1,,para3,para4],type = 'l',col="black", lty = 1, lwd = 3);
          lines(Parameter2,ChoSpeed0[spt,para1,,para3,para4],type = 'l',col="black", lty = 2, lwd = 3);
          
          #----------------- Resistance -----------------
          lines(Parameter2,ResSpeed1[spt,para1,,para3,para4],type = 'l',col="red", lty = 1, lwd = 3);
          lines(Parameter2,ResSpeed0[spt,para1,,para3,para4],type = 'l',col="red", lty = 2, lwd = 3);
          
          # legend("bottomleft", legend = c("Choice", "Resistance"), col = c("black", "red"), text.font = 3, lwd = 3, lty=c(1,1))
          # legend("bottomright", legend = c( "Evolving", "Fixed"), col = c("black"), text.font = 3, lwd = 3, lty=c(1,2))
          dev.off()        
        }}}}#}
}


#------------------- Speed of adaptation -  Not absolute values ---------------------------
Minspeed = min(c(noabs_ChoSpeed1,noabs_ChoSpeed0, noabs_ResSpeed0, noabs_ResSpeed1), na.rm = T)
Maxspeed = max(c(noabs_ChoSpeed1,noabs_ChoSpeed0, noabs_ResSpeed0, noabs_ResSpeed1), na.rm = T)
{
  #for(para3 in 1:length(Parameter3)){
  para3 = 0
  for(para4 in 1:length(Parameter4)){
    for(para1 in 1:length(Parameter1)){
      for(para2 in 1:length(Parameter2)){
        for(spt in 1:length(tTspeed)){
          png(paste0(getwd(), "/",Folder,"/", plotFolder, "/",subFolder,"/noabs_SpeedEvolutionTraits_Param1",para1, "_Param3", para3, "_Param4",para4, "_Param2", para2, "_gen",tTspeed[spt], "_paracomb",ParameterCombination,".png"))
          
          par(mar = c(5, 5, 4, 5) + 0.3)
          
          matplot(Parameter3,noabs_ChoSpeed1[spt,para1,para2,,para4],ylim = c(Minspeed,Maxspeed),cex.axis = 2, type = "n", ylab= list("Evolution rate", font = 3, cex = 2),xlab=list("Parameter 3", font = 3, cex = 2))
          lines(Parameter3,noabs_ChoSpeed1[spt,para1,para2,,para4],type = 'l',col="black", lty = 1, lwd = 3);
          lines(Parameter3,noabs_ChoSpeed0[spt,para1,para2,,para4],type = 'l',col="black", lty = 2, lwd = 3);
          
          #----------------- Resistance -----------------
          lines(Parameter3,noabs_ResSpeed1[spt,para1,para2,,para4],type = 'l',col="red", lty = 1, lwd = 3);
          lines(Parameter3,noabs_ResSpeed0[spt,para1,para2,,para4],type = 'l',col="red", lty = 2, lwd = 3);
          
          polygon(x=c(Parameter3, rev(Parameter3)), y=c(noabs_ResSpeed1[spt,para1,para2,,para4]-noabs_stdResSpeed1[spt,para1,para2,,para4], rev(noabs_ResSpeed1[spt,para1,para2,,para4]+noabs_stdResSpeed1[spt,para1,para2,,para4])), border=NA, col=adjustcolor("gray", alpha.f = 0.3))
          polygon(x=c(Parameter3, rev(Parameter3)), y=c(noabs_ResSpeed0[spt,para1,para2,,para4]-noabs_stdResSpeed0[spt,para1,para2,,para4], rev(noabs_ResSpeed0[spt,para1,para2,,para4]+noabs_stdResSpeed0[spt,para1,para2,,para4])), border=NA, col=adjustcolor("gray", alpha.f = 0.3))
          polygon(x=c(Parameter3, rev(Parameter3)), y=c(noabs_ChoSpeed1[spt,para1,para2,,para4]-noabs_stdChoSpeed1[spt,para1,para2,,para4], rev(noabs_ChoSpeed1[spt,para1,para2,,para4]+noabs_stdChoSpeed1[spt,para1,para2,,para4])), border=NA, col=adjustcolor("gray", alpha.f = 0.3))
          polygon(x=c(Parameter3, rev(Parameter3)), y=c(noabs_ChoSpeed0[spt,para1,para2,,para4]-noabs_stdChoSpeed0[spt,para1,para2,,para4], rev(noabs_ChoSpeed0[spt,para1,para2,,para4]+noabs_stdChoSpeed0[spt,para1,para2,,para4])), border=NA, col=adjustcolor("gray", alpha.f = 0.3))
          
          legend("bottomleft", legend = c("Trait", "Choice", "Resistance"), col = c(NA,"black", "red"), text.font = 3, lwd = 3, lty=c(1,1,1))
          legend("bottomright", legend = c("Behaviour", "Evolving", "Fixed"), col = c(NA,"black", "black"), text.font = 3, lwd = 3, lty=c(1,1,2))
          dev.off()        
        }}}}#}
}


{
  for(para3 in 1:length(Parameter3)){
    #for(para4 in 1:length(Parameter4)){
    para4 = 0
    for(para1 in 1:length(Parameter1)){
      for(para2 in 1:length(Parameter2)){
        for(spt in 1:length(tTspeed)){
          png(paste0(getwd(), "/",Folder,"/", plotFolder, "/",subFolder,"/noabs_SpeedEvolutionTraits_Param1",para1, "_Param3", para3, "_Param4",para4, "_Param2", para2, "_gen",tTspeed[spt], "_paracomb",ParameterCombination,".png"))
          
          par(mar = c(5, 5, 4, 5) + 0.3)
          
          matplot(Parameter4,noabs_ChoSpeed1[spt,para1,para2,para3,],ylim = c(Minspeed,Maxspeed),cex.axis = 2, type = "n", ylab= list("Evolution rate", font = 3, cex = 2),xlab=list("Parameter 4", font = 3, cex = 2))
          lines(Parameter4,noabs_ChoSpeed1[spt,para1,para2,para3,],type = 'l',col="black", lty = 1, lwd = 3);
          lines(Parameter4,noabs_ChoSpeed0[spt,para1,para2,para3,],type = 'l',col="black", lty = 2, lwd = 3);
          
          #----------------- Resistance -----------------
          lines(Parameter4,noabs_ResSpeed1[spt,para1,para2,para3,],type = 'l',col="red", lty = 1, lwd = 3);
          lines(Parameter4,noabs_ResSpeed0[spt,para1,para2,para3,],type = 'l',col="red", lty = 2, lwd = 3);
          
          polygon(x=c(Parameter4, rev(Parameter4)), y=c(noabs_ResSpeed1[spt,para1,para2,para3,]-noabs_stdResSpeed1[spt,para1,para2,para3,], rev(noabs_ResSpeed1[spt,para1,para2,para3,]+noabs_stdResSpeed1[spt,para1,para2,para3,])), border=NA, col=adjustcolor("gray", alpha.f = 0.3))
          polygon(x=c(Parameter4, rev(Parameter4)), y=c(noabs_ResSpeed0[spt,para1,para2,para3,]-noabs_stdResSpeed0[spt,para1,para2,para3,], rev(noabs_ResSpeed0[spt,para1,para2,para3,]+noabs_stdResSpeed0[spt,para1,para2,para3,])), border=NA, col=adjustcolor("gray", alpha.f = 0.3))
          polygon(x=c(Parameter4, rev(Parameter4)), y=c(noabs_ChoSpeed1[spt,para1,para2,para3,]-noabs_stdChoSpeed1[spt,para1,para2,para3,], rev(noabs_ChoSpeed1[spt,para1,para2,para3,]+noabs_stdChoSpeed1[spt,para1,para2,para3,])), border=NA, col=adjustcolor("gray", alpha.f = 0.3))
          polygon(x=c(Parameter4, rev(Parameter4)), y=c(noabs_ChoSpeed0[spt,para1,para2,para3,]-noabs_stdChoSpeed0[spt,para1,para2,para3,], rev(noabs_ChoSpeed0[spt,para1,para2,para3,]+noabs_stdChoSpeed0[spt,para1,para2,para3,])), border=NA, col=adjustcolor("gray", alpha.f = 0.3))
          
          # legend("bottomleft", legend = c("Choice", "Resistance"), col = c("black", "red"), text.font = 3, lwd = 3, lty=c(1,1))
          # legend("bottomright", legend = c( "Evolving", "Fixed"), col = c("black"), text.font = 3, lwd = 3, lty=c(1,2))
          dev.off()        
        }}}}#}
}


{
  for(para3 in 1:length(Parameter3)){
    for(para4 in 1:length(Parameter4)){
      #for(para1 in 1:length(Parameter1)){
      para1 = 0
      for(para2 in 1:length(Parameter2)){
        for(spt in 1:length(tTspeed)){
          png(paste0(getwd(), "/",Folder,"/", plotFolder, "/",subFolder,"/noabs_SpeedEvolutionTraits_Param1",para1, "_Param3", para3, "_Param4",para4, "_Param2", para2, "_gen",tTspeed[spt], "_paracomb",ParameterCombination,".png"))
          par(mar = c(5, 5, 4, 5) + 0.3)
          
          matplot(Parameter1,noabs_ChoSpeed1[spt,,para2,para3,para4],ylim = c(Minspeed,Maxspeed),cex.axis = 2, type = "n", ylab= list("Evolution rate", font = 3, cex = 2),xlab=list("Parameter 1", font = 3, cex = 2))
          lines(Parameter1,noabs_ChoSpeed1[spt,,para2,para3,para4],type = 'l',col="black", lty = 1, lwd = 3);
          lines(Parameter1,noabs_ChoSpeed0[spt,,para2,para3,para4],type = 'l',col="black", lty = 2, lwd = 3);
          
          #----------------- Resistance -----------------
          lines(Parameter1,noabs_ResSpeed1[spt,,para2,para3,para4],type = 'l',col="red", lty = 1, lwd = 3);
          lines(Parameter1,noabs_ResSpeed0[spt,,para2,para3,para4],type = 'l',col="red", lty = 2, lwd = 3);
          
          polygon(x=c(Parameter1, rev(Parameter1)), y=c(noabs_ResSpeed1[spt,,para2,para3,para4]-noabs_stdResSpeed1[spt,,para2,para3,para4], rev(noabs_ResSpeed1[spt,,para2,para3,para4]+noabs_stdResSpeed1[spt,,para2,para3,para4])), border=NA, col=adjustcolor("gray", alpha.f = 0.3))
          polygon(x=c(Parameter1, rev(Parameter1)), y=c(noabs_ResSpeed0[spt,,para2,para3,para4]-noabs_stdResSpeed0[spt,,para2,para3,para4], rev(noabs_ResSpeed0[spt,,para2,para3,para4]+noabs_stdResSpeed0[spt,,para2,para3,para4])), border=NA, col=adjustcolor("gray", alpha.f = 0.3))
          polygon(x=c(Parameter1, rev(Parameter1)), y=c(noabs_ChoSpeed1[spt,,para2,para3,para4]-noabs_stdChoSpeed1[spt,,para2,para3,para4], rev(noabs_ChoSpeed1[spt,,para2,para3,para4]+noabs_stdChoSpeed1[spt,,para2,para3,para4])), border=NA, col=adjustcolor("gray", alpha.f = 0.3))
          polygon(x=c(Parameter1, rev(Parameter1)), y=c(noabs_ChoSpeed0[spt,,para2,para3,para4]-noabs_stdChoSpeed0[spt,,para2,para3,para4], rev(noabs_ChoSpeed0[spt,,para2,para3,para4]+noabs_stdChoSpeed0[spt,,para2,para3,para4])), border=NA, col=adjustcolor("gray", alpha.f = 0.3))
          
          # legend("bottomleft", legend = c("Choice", "Resistance"), col = c("black", "red"), text.font = 3, lwd = 3, lty=c(1,1))
          # legend("bottomright", legend = c( "Evolving", "Fixed"), col = c("black"), text.font = 3, lwd = 3, lty=c(1,2))
          dev.off()        
        }}}}#}
}


{
  for(para3 in 1:length(Parameter3)){
    for(para4 in 1:length(Parameter4)){
      for(para1 in 1:length(Parameter1)){
        #for(para2 in 1:length(Parameter2)){
        for(spt in 1:length(tTspeed)){
          para2 = 0
          png(paste0(getwd(), "/",Folder,"/", plotFolder, "/",subFolder,"/noabs_SpeedEvolutionTraits_Param1",para1, "_Param3", para3, "_Param4",para4, "_Param2", para2, "_gen",tTspeed[spt], "_paracomb",ParameterCombination,".png"))
          
          par(mar = c(5, 5, 4, 5) + 0.3)
          
          matplot(Parameter2,noabs_ChoSpeed1[spt,para1,,para3,para4],ylim = c(Minspeed,Maxspeed),cex.axis = 2, type = "n", ylab= list("Evolution rate", font = 3, cex = 2),xlab=list("Parameter 2", font = 3, cex = 2))
          lines(Parameter2,noabs_ChoSpeed1[spt,para1,,para3,para4],type = 'l',col="black", lty = 1, lwd = 3);
          lines(Parameter2,noabs_ChoSpeed0[spt,para1,,para3,para4],type = 'l',col="black", lty = 2, lwd = 3);
          
          #----------------- Resistance -----------------
          lines(Parameter2,noabs_ResSpeed1[spt,para1,,para3,para4],type = 'l',col="red", lty = 1, lwd = 3);
          lines(Parameter2,noabs_ResSpeed0[spt,para1,,para3,para4],type = 'l',col="red", lty = 2, lwd = 3);
          
          polygon(x=c(Parameter2, rev(Parameter2)), y=c(noabs_ResSpeed1[spt,para1,,para3,para4]-noabs_stdResSpeed1[spt,para1,,para3,para4], rev(noabs_ResSpeed1[spt,para1,,para3,para4]+noabs_stdResSpeed1[spt,para1,,para3,para4])), border=NA, col=adjustcolor("gray", alpha.f = 0.3))
          polygon(x=c(Parameter2, rev(Parameter2)), y=c(noabs_ResSpeed0[spt,para1,,para3,para4]-noabs_stdResSpeed0[spt,para1,,para3,para4], rev(noabs_ResSpeed0[spt,para1,,para3,para4]+noabs_stdResSpeed0[spt,para1,,para3,para4])), border=NA, col=adjustcolor("gray", alpha.f = 0.3))
          polygon(x=c(Parameter2, rev(Parameter2)), y=c(noabs_ChoSpeed1[spt,para1,,para3,para4]-noabs_stdChoSpeed1[spt,para1,,para3,para4], rev(noabs_ChoSpeed1[spt,para1,,para3,para4]+noabs_stdChoSpeed1[spt,para1,,para3,para4])), border=NA, col=adjustcolor("gray", alpha.f = 0.3))
          polygon(x=c(Parameter2, rev(Parameter2)), y=c(noabs_ChoSpeed0[spt,para1,,para3,para4]-noabs_stdChoSpeed0[spt,para1,,para3,para4], rev(noabs_ChoSpeed0[spt,para1,,para3,para4]+noabs_stdChoSpeed0[spt,para1,,para3,para4])), border=NA, col=adjustcolor("gray", alpha.f = 0.3))
          
          
          # legend("bottomleft", legend = c("Choice", "Resistance"), col = c("black", "red"), text.font = 3, lwd = 3, lty=c(1,1))
          # legend("bottomright", legend = c( "Evolving", "Fixed"), col = c("black"), text.font = 3, lwd = 3, lty=c(1,2))
          dev.off()        
        }}}}#}
}


#------------------ Plant Population size -----------------------

subFolder = "PlantPopSize"

if(!dir.exists(paste0(getwd(), "/",Folder, "/", plotFolder, "/", subFolder))){
  dir.create(paste0(getwd(), "/",Folder, "/", plotFolder, "/", subFolder))
}


{
  #for(para3 in 1:length(Parameter3)){
  para3 = 0
  for(para4 in 1:length(Parameter4)){
    for(para1 in 1:length(Parameter1)){
      for(para2 in 1:length(Parameter2)){
        
        png(paste0(getwd(), "/",Folder,"/", plotFolder, "/",subFolder,"/PlantPopsize_Param1",para1, "_Param3", para3, "_Param4",para4, "_Param2", para2, "_paracomb",ParameterCombination,".png"))
        
        par(mar = c(5, 5, 4, 4) + 0.3)
        plot(0,0,xlim = range(Parameter3),ylim = c(0,max(N1_mean0[tT,para1,para2,,para4], N2_mean0[tT,para1,para2,,para4], N1_mean1[tT,para1,para2,,para4], N2_mean1[tT,para1,para2,,para4], na.rm = TRUE)),cex.axis = 2, type = "n",
             ylab= list("Population size", font = 3, cex = 2),xlab= list("Parameter 3", font = 3, cex = 2))
        lines(Parameter3,N1_mean1[tT,para1,para2,,para4],type = 'l',col="purple",lwd = 3);
        polygon(x=c(Parameter3, rev(Parameter3)), y=c(N1_mean1[tT,para1,para2,,para4]-N1_std1[tT,para1,para2,,para4], rev(N1_mean1[tT,para1,para2,,para4]+N1_std1[tT,para1,para2,,para4])), border=NA, col=adjustcolor("gray", alpha.f = 0.3))
        lines(Parameter3,N1_mean0[tT,para1,para2,,para4],type = 'l',col="gray", lwd = 3);
        polygon(x=c(Parameter3, rev(Parameter3)), y=c(N1_mean0[tT,para1,para2,,para4]-N1_std0[tT,para1,para2,,para4], rev(N1_mean0[tT,para1,para2,,para4]+N1_std0[tT,para1,para2,,para4])), border=NA, col=adjustcolor("gray", alpha.f = 0.3))
        
        lines(Parameter3,N2_mean1[tT,para1,para2,,para4],type = 'l', lty = 2,col="purple", lwd = 3);
        polygon(x=c(Parameter3, rev(Parameter3)), y=c(N2_mean1[tT,para1,para2,,para4]-N2_std1[tT,para1,para2,,para4], rev(N2_mean1[tT,para1,para2,,para4]+N2_std1[tT,para1,para2,,para4])), border=NA, col=adjustcolor("gray", alpha.f = 0.3))
        lines(Parameter3,N2_mean0[tT,para1,para2,,para4],type = 'l',lty = 2,col="gray", lwd = 3);
        polygon(x=c(Parameter3, rev(Parameter3)), y=c(N2_mean0[tT,para1,para2,,para4]-N2_std0[tT,para1,para2,,para4], rev(N2_mean0[tT,para1,para2,,para4]+N2_std0[tT,para1,para2,,para4])), border=NA, col=adjustcolor("gray", alpha.f = 0.3))
        
        legend("bottomright", legend = c("HOST PLANT", "Non-toxic", "Toxic", "BEHAVIOUR", "Evolving",  "Fixed"), col = c(NA,"gray", "purple", NA,1,1), text.font = 3, lwd = 3, lty=c(NA,1,1,NA,1,2), cex=1.5, text.col = c("blue",1,1,"blue",1,1))
        
        dev.off()        
      }}}#}
}


{
  for(para3 in 1:length(Parameter3)){
    #for(para4 in 1:length(Parameter4)){
    para4 = 0
    for(para1 in 1:length(Parameter1)){
      for(para2 in 1:length(Parameter2)){
        
        png(paste0(getwd(), "/",Folder,"/", plotFolder, "/",subFolder,"/PlantPopsize_Param1",para1, "_Param3", para3, "_Param4",para4, "_Param2", para2, "_paracomb",ParameterCombination,".png"))
        
        par(mar = c(5, 5, 4, 4) + 0.3)
        plot(0,0,xlim = range(Parameter4),ylim = c(0,max(N1_mean0[tT,para1,para2,para3,], N2_mean0[tT,para1,para2,para3,], N1_mean1[tT,para1,para2,para3,], N2_mean1[tT,para1,para2,para3,], na.rm = TRUE)),cex.axis = 2, type = "n",
             ylab= list("Population size", font = 3, cex = 2),xlab= list("Parameter 4", font = 3, cex = 2))
        lines(Parameter4,N1_mean1[tT,para1,para2,para3,],type = 'l',col="purple",lwd = 3);
        polygon(x=c(Parameter4, rev(Parameter4)), y=c(N1_mean1[tT,para1,para2,para3,]-N1_std1[tT,para1,para2,para3,], rev(N1_mean1[tT,para1,para2,para3,]+N1_std1[tT,para1,para2,para3,])), border=NA, col=adjustcolor("gray", alpha.f = 0.3))
        lines(Parameter4,N1_mean0[tT,para1,para2,para3,],type = 'l',col="gray", lwd = 3);
        polygon(x=c(Parameter4, rev(Parameter4)), y=c(N1_mean0[tT,para1,para2,para3,]-N1_std0[tT,para1,para2,para3,], rev(N1_mean0[tT,para1,para2,para3,]+N1_std0[tT,para1,para2,para3,])), border=NA, col=adjustcolor("gray", alpha.f = 0.3))
        
        lines(Parameter4,N2_mean1[tT,para1,para2,para3,],type = 'l', lty = 2,col="purple", lwd = 3);
        polygon(x=c(Parameter4, rev(Parameter4)), y=c(N2_mean1[tT,para1,para2,para3,]-N2_std1[tT,para1,para2,para3,], rev(N2_mean1[tT,para1,para2,para3,]+N2_std1[tT,para1,para2,para3,])), border=NA, col=adjustcolor("gray", alpha.f = 0.3))
        lines(Parameter4,N2_mean0[tT,para1,para2,para3,],type = 'l',lty = 2,col="gray", lwd = 3);
        polygon(x=c(Parameter4, rev(Parameter4)), y=c(N2_mean0[tT,para1,para2,para3,]-N2_std0[tT,para1,para2,para3,], rev(N2_mean0[tT,para1,para2,para3,]+N2_std0[tT,para1,para2,para3,])), border=NA, col=adjustcolor("gray", alpha.f = 0.3))
        
        #legend("bottomright", legend = c("HOST PLANT", "Non-toxic", "Toxic", "BEHAVIOUR", "Evolving",  "Fixed"), col = c(NA,"gray", "purple", NA,1,1), text.font = 3, lwd = 3, lty=c(NA,1,1,NA,1,2), cex=1.5, text.col = c("blue",1,1,"blue",1,1))
        
        dev.off()        
      }}}#}
}


{
  for(para3 in 1:length(Parameter3)){
    for(para4 in 1:length(Parameter4)){
      
      #for(para1 in 1:length(Parameter1)){
      para1 = 0
      for(para2 in 1:length(Parameter2)){
        
        png(paste0(getwd(), "/",Folder,"/", plotFolder, "/",subFolder,"/PlantPopsize_Param1",para1, "_Param3", para3, "_Param4",para4, "_Param2", para2, "_paracomb",ParameterCombination,".png"))
        
        par(mar = c(5, 5, 4, 4) + 0.3)
        plot(0,0,xlim = range(Parameter2),ylim = c(0,max(N1_mean0[tT,,para2,para3,para4], N2_mean0[tT,,para2,para3,para4], N1_mean1[tT,,para2,para3,para4], N2_mean1[tT,,para2,para3,para4], na.rm = TRUE)),cex.axis = 2, type = "n",
             ylab= list("Population size", font = 3, cex = 2),xlab= list("Parameter 1", font = 3, cex = 2))
        lines(Parameter2,N1_mean1[tT,,para2,para3,para4],type = 'l',col="purple",lwd = 3);
        polygon(x=c(Parameter2, rev(Parameter2)), y=c(N1_mean1[tT,,para2,para3,para4]-N1_std1[tT,,para2,para3,para4], rev(N1_mean1[tT,,para2,para3,para4]+N1_std1[tT,,para2,para3,para4])), border=NA, col=adjustcolor("gray", alpha.f = 0.3))
        lines(Parameter2,N1_mean0[tT,,para2,para3,para4],type = 'l',col="gray", lwd = 3);
        polygon(x=c(Parameter2, rev(Parameter2)), y=c(N1_mean0[tT,,para2,para3,para4]-N1_std0[tT,,para2,para3,para4], rev(N1_mean0[tT,,para2,para3,para4]+N1_std0[tT,,para2,para3,para4])), border=NA, col=adjustcolor("gray", alpha.f = 0.3))
        
        lines(Parameter2,N2_mean1[tT,,para2,para3,para4],type = 'l', lty = 2,col="purple", lwd = 3);
        polygon(x=c(Parameter2, rev(Parameter2)), y=c(N2_mean1[tT,,para2,para3,para4]-N2_std1[tT,,para2,para3,para4], rev(N2_mean1[tT,,para2,para3,para4]+N2_std1[tT,,para2,para3,para4])), border=NA, col=adjustcolor("gray", alpha.f = 0.3))
        lines(Parameter2,N2_mean0[tT,,para2,para3,para4],type = 'l',lty = 2,col="gray", lwd = 3);
        polygon(x=c(Parameter2, rev(Parameter2)), y=c(N2_mean0[tT,,para2,para3,para4]-N2_std0[tT,,para2,para3,para4], rev(N2_mean0[tT,,para2,para3,para4]+N2_std0[tT,,para2,para3,para4])), border=NA, col=adjustcolor("gray", alpha.f = 0.3))
        
        #legend("bottomright", legend = c("HOST PLANT", "Non-toxic", "Toxic", "BEHAVIOUR", "Evolving",  "Fixed"), col = c(NA,"gray", "purple", NA,1,1), text.font = 3, lwd = 3, lty=c(NA,1,1,NA,1,2), cex=1.5, text.col = c("blue",1,1,"blue",1,1))
        
        dev.off()        
      }}}#}
}


{
  for(para3 in 1:length(Parameter3)){
    for(para4 in 1:length(Parameter4)){
      for(para1 in 1:length(Parameter1)){
        #for(para2 in 1:length(Parameter2)){
        para2 = 0 
        png(paste0(getwd(), "/",Folder,"/", plotFolder, "/",subFolder,"/PlantPopsize_Param1",para1, "_Param3", para3, "_Param4",para4, "_Param2", para2, "_paracomb",ParameterCombination,".png"))
        
        par(mar = c(5, 5, 4, 4) + 0.3)
        plot(0,0,xlim = range(Parameter1),ylim = c(0,max(N1_mean0[tT,para1,,para3,para4], N2_mean0[tT,para1,,para3,para4], N1_mean1[tT,para1,,para3,para4], N2_mean1[tT,para1,,para3,para4], na.rm = TRUE)),cex.axis = 2, type = "n",
             ylab= list("Population size", font = 3, cex = 2),xlab= list("Parameter 2", font = 3, cex = 2))
        lines(Parameter1,N1_mean1[tT,para1,,para3,para4],type = 'l',col="purple",lwd = 3);
        polygon(x=c(Parameter1, rev(Parameter1)), y=c(N1_mean1[tT,para1,,para3,para4]-N1_std1[tT,para1,,para3,para4], rev(N1_mean1[tT,para1,,para3,para4]+N1_std1[tT,para1,,para3,para4])), border=NA, col=adjustcolor("gray", alpha.f = 0.3))
        lines(Parameter1,N1_mean0[tT,para1,,para3,para4],type = 'l',col="gray", lwd = 3);
        polygon(x=c(Parameter1, rev(Parameter1)), y=c(N1_mean0[tT,para1,,para3,para4]-N1_std0[tT,para1,,para3,para4], rev(N1_mean0[tT,para1,,para3,para4]+N1_std0[tT,para1,,para3,para4])), border=NA, col=adjustcolor("gray", alpha.f = 0.3))
        
        lines(Parameter1,N2_mean1[tT,para1,,para3,para4],type = 'l', lty = 2,col="purple", lwd = 3);
        polygon(x=c(Parameter1, rev(Parameter1)), y=c(N2_mean1[tT,para1,,para3,para4]-N2_std1[tT,para1,,para3,para4], rev(N2_mean1[tT,para1,,para3,para4]+N2_std1[tT,para1,,para3,para4])), border=NA, col=adjustcolor("gray", alpha.f = 0.3))
        lines(Parameter1,N2_mean0[tT,para1,,para3,para4],type = 'l',lty = 2,col="gray", lwd = 3);
        polygon(x=c(Parameter1, rev(Parameter1)), y=c(N2_mean0[tT,para1,,para3,para4]-N2_std0[tT,para1,,para3,para4], rev(N2_mean0[tT,para1,,para3,para4]+N2_std0[tT,para1,,para3,para4])), border=NA, col=adjustcolor("gray", alpha.f = 0.3))
        
        #legend("bottomright", legend = c("HOST PLANT", "Non-toxic", "Toxic", "BEHAVIOUR", "Evolving",  "Fixed"), col = c(NA,"gray", "purple", NA,1,1), text.font = 3, lwd = 3, lty=c(NA,1,1,NA,1,2), cex=1.5, text.col = c("blue",1,1,"blue",1,1))
        
        dev.off()        
      }}}#}
}


#------------- Time series plots. Here, I have selected only a few parameter combinations as it takes a lot of time to pass through many generations --------------------

Folder = "Dominance_Density_AM" # Folder for storing simulated data
FFolder = Folder
ParameterCombination = "Series"
seriesDominanceresistance = c(0,0.5,1)
seriesDominancechoice = c(0,0.5,1)
seriesScaledensity = c(1,3)
seriessortMating = c(0,0.8)

seriesTtimes = c(1:numgenerations)

#-------- Loading data 
DataAnalysis = "AnalysedData"#------ This folder is for storing plots

Rresistancemean1_timeseries= readRDS(paste0(getwd(),"/", Folder,"/", DataAnalysis, "/", "Rresistancemean1_timeseries.RData"))
Rresistancestd1_timeseries = readRDS(paste0(getwd(),"/", Folder,"/", DataAnalysis, "/","Rresistancestd1_timeseries.RData"))
Cchoicemean1_timeseries= readRDS(paste0(getwd(),"/", Folder,"/", DataAnalysis, "/","Cchoicemean1_timeseries.RData"))
Cchoicestd1_timeseries= readRDS(paste0(getwd(),"/", Folder,"/", DataAnalysis, "/","Cchoicestd1_timeseries.RData"))
Rresistancemean0_timeseries = readRDS(paste0(getwd(),"/", Folder,"/", DataAnalysis, "/", "Rresistancemean0_timeseries.RData"))
Rresistancestd0_timeseries = readRDS(paste0(getwd(),"/", Folder,"/", DataAnalysis, "/","Rresistancestd0_timeseries.RData"))
Cchoicemean0_timeseries= readRDS(paste0(getwd(),"/", Folder,"/", DataAnalysis, "/","Cchoicemean0_timeseries.RData"))
Cchoicestd0_timeseries= readRDS(paste0(getwd(),"/", Folder,"/", DataAnalysis, "/","Cchoicestd0_timeseries.RData"))

Rresistance_popvariancemean1_timeseries= readRDS(paste0(getwd(),"/", Folder,"/", DataAnalysis, "/", "Rresistance_popvariancemean1_timeseries.RData"))
Rresistance_popvariancestd1_timeseries = readRDS(paste0(getwd(),"/", Folder,"/", DataAnalysis, "/","Rresistance_popvariancestd1_timeseries.RData"))
Cchoice_popvariancemean1_timeseries= readRDS(paste0(getwd(),"/", Folder,"/", DataAnalysis, "/","Cchoice_popvariancemean1_timeseries.RData"))
Cchoice_popvariancestd1_timeseries= readRDS(paste0(getwd(),"/", Folder,"/", DataAnalysis, "/","Cchoice_popvariancestd1_timeseries.RData"))
Rresistance_popvariancemean0_timeseries = readRDS(paste0(getwd(),"/", Folder,"/", DataAnalysis, "/", "Rresistance_popvariancemean0_timeseries.RData"))
Rresistance_popvariancestd0_timeseries = readRDS(paste0(getwd(),"/", Folder,"/", DataAnalysis, "/","Rresistance_popvariancestd0_timeseries.RData"))
Cchoice_popvariancemean0_timeseries= readRDS(paste0(getwd(),"/", Folder,"/", DataAnalysis, "/","Cchoice_popvariancemean0_timeseries.RData"))
Cchoice_popvariancestd0_timeseries= readRDS(paste0(getwd(),"/", Folder,"/", DataAnalysis, "/","Cchoice_popvariancestd0_timeseries.RData"))

N1_mean1_timeseries= readRDS(paste0(getwd(),"/", Folder,"/", DataAnalysis, "/", "N1_mean1_timeseries.RData"))
N1_mean0_timeseries= readRDS(paste0(getwd(),"/", Folder,"/", DataAnalysis, "/","N1_mean0_timeseries.RData"))
N1_std1_timeseries= readRDS(paste0(getwd(),"/", Folder,"/", DataAnalysis, "/","N1_std1_timeseries.RData"))
N1_std0_timeseries= readRDS(paste0(getwd(),"/", Folder,"/", DataAnalysis, "/","N1_std0_timeseries.RData"))
N2_mean1_timeseries= readRDS(paste0(getwd(),"/", Folder,"/", DataAnalysis, "/", "N2_mean1_timeseries.RData"))
N2_mean0_timeseries= readRDS(paste0(getwd(),"/", Folder,"/", DataAnalysis, "/","N2_mean0_timeseries.RData"))
N2_std1_timeseries= readRDS(paste0(getwd(),"/", Folder,"/", DataAnalysis, "/","N2_std1_timeseries.RData"))
N2_std0_timeseries= readRDS(paste0(getwd(),"/", Folder,"/", DataAnalysis, "/","N2_std0_timeseries.RData"))


plotFolder = "Plots"

if(!dir.exists(paste0(getwd(), "/",Folder, "/", plotFolder))){
  dir.create(paste0(getwd(), "/",Folder, "/", plotFolder))
}


subFolder = "Timeseries"

if(!dir.exists(paste0(getwd(), "/",Folder, plotFolder, subFolder))){
  dir.create(paste0(getwd(), "/",Folder, plotFolder, subFolder))
}


{
  for(para3 in 1:length(seriesScaledensity)){
    for(para4 in 1:length(seriessortMating)){
      for(para1 in 1:length(seriesDominanceresistance)){
        for(para2 in 1:length(seriesDominancechoice)){
          
          png(paste0(getwd(), "/",Folder,"/",plotFolder,"/",subFolder,"/Timeseries_meanTraits_Param1",para1, "_Param3", para3, "_Param4",para4, "_Param2", para2, "_paracomb",ParameterCombination,".png"))
          
          par(mar = c(5, 5, 4, 5) + 0.3)
          plot(0,0,xlim = range(seriesTtimes),ylim = c(0,1),cex.axis = 2, type = "n", ylab= list("Mean trait", font = 3, cex = 2),xlab=list("Generations", font = 3, cex = 2))
          lines(seriesTtimes,Cchoicemean1_timeseries[,para1,para2,para4,para3],type = 'l',col="black", lty = 1, lwd = 3);
          polygon(x=c(seriesTtimes, rev(seriesTtimes)), y=c(Cchoicemean1_timeseries[,para1,para2,para4,para3]-Cchoicestd1_timeseries[,para1,para2,para4,para3], rev(Cchoicemean1_timeseries[,para1,para2,para4,para3]+Cchoicestd1_timeseries[,para1,para2,para4,para3])), border=NA, col=adjustcolor("gray", alpha.f = 0.3))
          lines(seriesTtimes,Cchoicemean0_timeseries[,para1,para2,para4,para3],type = 'l',col="black", lty = 2, lwd = 3);
          polygon(x=c(seriesTtimes, rev(seriesTtimes)), y=c(Cchoicemean0_timeseries[,para1,para2,para4,para3]-Cchoicestd0_timeseries[,para1,para2,para4,para3], rev(Cchoicemean0_timeseries[,para1,para2,para4,para3]+Cchoicestd0_timeseries[,para1,para2,para4,para3])), border=NA, col=adjustcolor("gray", alpha.f = 0.3))
          
          #----------------- Resistance -----------------
          lines(seriesTtimes,Rresistancemean1_timeseries[,para1,para2,para4,para3],type = 'l',col="red", lty = 1, lwd = 3);
          polygon(x=c(seriesTtimes, rev(seriesTtimes)), y=c(Rresistancemean1_timeseries[,para1,para2,para4,para3] - Rresistancestd1_timeseries[,para1,para2,para4,para3], rev(Rresistancemean1_timeseries[,para1,para2,para4,para3] + Rresistancestd1_timeseries[,para1,para2,para4,para3])), border=NA, col=adjustcolor("gray", alpha.f = 0.3))
          
          lines(seriesTtimes,Rresistancemean0_timeseries[,para1,para2,para4,para3],type = 'l',col="red", lty = 2, lwd = 3);
          polygon(x=c(seriesTtimes, rev(seriesTtimes)), y=c(Rresistancemean0_timeseries[,para1,para2,para4,para3] - Rresistancestd0_timeseries[,para1,para2,para4,para3], rev(Rresistancemean0_timeseries[,para1,para2,para4,para3] + Rresistancestd0_timeseries[,para1,para2,para4,para3])), border=NA, col=adjustcolor("gray", alpha.f = 0.3))
          
          #legend("bottomright", legend = c("Trait", "Choice", "Resistance"), col = c(NA,"black", "red"), text.font = 3, lwd = 3, lty=c(1,1), cex=1.5)
          legend("bottomright", legend = c("TRAIT", "Choice", "Resistance"), col = c(NA,"black", "red"), text.font = 3, lwd = 3, lty=c(1,1), cex=1.5, text.col = c("blue", "black", "black"))
          
          legend("left", legend = c("Behaviour", "Evolving", "Fixed"), col = c("black"), text.font = 3, lwd = 3, lty=c(NA,1,2), cex=1.5)
          dev.off()        
        }}}}
}


#-------------- Plant sizes

{
  for(para3 in 1:length(seriesScaledensity)){
    for(para4 in 1:length(seriessortMating)){
      for(para1 in 1:length(seriesDominanceresistance)){
        for(para2 in 1:length(seriesDominancechoice)){
          
          png(paste0(getwd(), "/",Folder,"/",plotFolder,"/",subFolder,"/TimeseriesPlantPopsize_Param1",para1, "_Param3", para3, "_Param4",para4, "_Param2", para2, "_paracomb",ParameterCombination,".png"))
          
          par(mar = c(5, 5, 4, 4) + 0.3)
          plot(0,0,xlim = range(seriesTtimes),ylim = c(0,max(N1_mean0_timeseries[,para1,para2,para4,para3], N2_mean0_timeseries[,para1,para2,para4,para3], N1_mean1_timeseries[,para1,para2,para4,para3], N2_mean1_timeseries[,para1,para2,para4,para3], na.rm = TRUE)),cex.axis = 2, type = "n",ylab= list("Population size", font = 3, cex = 2),xlab= list("Generations", font = 3, cex = 2))
          lines(seriesTtimes,N1_mean1_timeseries[,para1,para2,para4,para3],type = 'l',col="gray", lwd = 3);
          polygon(x=c(seriesTtimes, rev(seriesTtimes)), y=c(N1_mean1_timeseries[,para1,para2,para4,para3]-N1_std1_timeseries[,para1,para2,para4,para3], rev(N1_mean1_timeseries[,para1,para2,para4,para3]+N1_std1_timeseries[,para1,para2,para4,para3])), border=NA, col=adjustcolor("gray", alpha.f = 0.3))
          lines(seriesTtimes,N1_mean0_timeseries[,para1,para2,para4,para3],type = 'l',col="gray", lty = 2, lwd = 3);
          polygon(x=c(seriesTtimes, rev(seriesTtimes)), y=c(N1_mean0_timeseries[,para1,para2,para4,para3]-N1_std0_timeseries[,para1,para2,para4,para3], rev(N1_mean0_timeseries[,para1,para2,para4,para3]+N1_std0_timeseries[,para1,para2,para4,para3])), border=NA, col=adjustcolor("gray", alpha.f = 0.3))
          
          lines(seriesTtimes,N2_mean1_timeseries[,para1,para2,para4,para3],type = 'l', lty = 1,col="purple", lwd = 3);
          polygon(x=c(seriesTtimes, rev(seriesTtimes)), y=c(N2_mean1_timeseries[,para1,para2,para4,para3]-N2_std1_timeseries[,para1,para2,para4,para3], rev(N2_mean1_timeseries[,para1,para2,para4,para3]+N2_std1_timeseries[,para1,para2,para4,para3])), border=NA, col=adjustcolor("gray", alpha.f = 0.3))
          lines(seriesTtimes,N2_mean0_timeseries[,para1,para2,para4,para3],type = 'l',lty = 2,col="purple", lwd = 3);
          polygon(x=c(seriesTtimes, rev(seriesTtimes)), y=c(N2_mean0_timeseries[,para1,para2,para4,para3]-N2_std0_timeseries[,para1,para2,para4,para3], rev(N2_mean0_timeseries[,para1,para2,para4,para3]+N2_std0_timeseries[,para1,para2,para4,para3])), border=NA, col=adjustcolor("gray", alpha.f = 0.3))
          
          #legend("bottomright", legend = c("HOST PLANT", "Non-toxic", "Toxic","BEHAVIOUR", "Evolving", "Fixed"), col = c(NA,"gray", "purple",NA,"black","black"), text.font = 3, lwd = 3, lty=c(NA,1,1,NA,1,2), cex=1.5)
          legend("bottomright", legend = c("HOST PLANT", "Non-toxic", "Toxic","BEHAVIOUR", "Evolving", "Fixed"), col = c(NA,"gray", "purple",NA,"black","black"), text.font = 3, lwd = 3, lty=c(NA,1,1,NA,1,2), cex=1.5, text.col = c("blue","black", "black", "blue","black", "black"))
          
          #legend("bottomright", legend = c("Behaviour", "Evolving", "Fixed"), col = c("black"), text.font = 3, lwd = 3, lty=c(NA,1,2),cex=1.5)
          
          
          dev.off()        
        }}}}
}


minvalue = min(Cchoice_popvariancemean1_timeseries, Cchoice_popvariancemean0_timeseries, Rresistance_popvariancemean1_timeseries, Rresistance_popvariancemean0_timeseries, na.rm = TRUE)
maxvalue = max(Cchoice_popvariancemean1_timeseries, Cchoice_popvariancemean0_timeseries, Rresistance_popvariancemean1_timeseries, Rresistance_popvariancemean0_timeseries, na.rm = TRUE)

{
  for(para3 in 1:length(seriesScaledensity)){
    for(para4 in 1:length(seriessortMating)){
      for(para1 in 1:length(seriesDominanceresistance)){
        for(para2 in 1:length(seriesDominancechoice)){
          
          png(paste0(getwd(), "/",Folder,"/",plotFolder,"/",subFolder,"/Timeseries_varianceTraits_Param1",para1, "_Param3", para3, "_Param4",para4, "_Param2", para2, "_paracomb",ParameterCombination,".png"))
          
          par(mar = c(5, 5, 4, 5) + 0.3)
          plot(0,0,xlim = range(seriesTtimes),ylim = c(minvalue,maxvalue),cex.axis = 2, type = "n", ylab= list("Population trait variance", font = 3, cex = 2),xlab=list("Generations", font = 3, cex = 2))
          lines(seriesTtimes,Cchoice_popvariancemean1_timeseries[,para1,para2,para4,para3],type = 'l',col="black", lty = 1, lwd = 3);
          polygon(x=c(seriesTtimes, rev(seriesTtimes)), y=c(Cchoice_popvariancemean1_timeseries[,para1,para2,para4,para3]-Cchoice_popvariancestd1_timeseries[,para1,para2,para4,para3], rev(Cchoice_popvariancemean1_timeseries[,para1,para2,para4,para3]+Cchoice_popvariancestd1_timeseries[,para1,para2,para4,para3])), border=NA, col=adjustcolor("gray", alpha.f = 0.3))
          lines(seriesTtimes,Cchoice_popvariancemean0_timeseries[,para1,para2,para4,para3],type = 'l',col="black", lty = 2, lwd = 3);
          polygon(x=c(seriesTtimes, rev(seriesTtimes)), y=c(Cchoice_popvariancemean0_timeseries[,para1,para2,para4,para3]-Cchoice_popvariancestd0_timeseries[,para1,para2,para4,para3], rev(Cchoice_popvariancemean0_timeseries[,para1,para2,para4,para3]+Cchoice_popvariancestd0_timeseries[,para1,para2,para4,para3])), border=NA, col=adjustcolor("gray", alpha.f = 0.3))
          
          #----------------- Resistance -----------------
          lines(seriesTtimes,Rresistance_popvariancemean1_timeseries[,para1,para2,para4,para3],type = 'l',col="red", lty = 1, lwd = 3);
          polygon(x=c(seriesTtimes, rev(seriesTtimes)), y=c(Rresistance_popvariancemean1_timeseries[,para1,para2,para4,para3] - Rresistance_popvariancestd1_timeseries[,para1,para2,para4,para3], rev(Rresistance_popvariancemean1_timeseries[,para1,para2,para4,para3] + Rresistance_popvariancestd1_timeseries[,para1,para2,para4,para3])), border=NA, col=adjustcolor("gray", alpha.f = 0.3))
          
          lines(seriesTtimes,Rresistance_popvariancemean0_timeseries[,para1,para2,para4,para3],type = 'l',col="red", lty = 2, lwd = 3);
          polygon(x=c(seriesTtimes, rev(seriesTtimes)), y=c(Rresistance_popvariancemean0_timeseries[,para1,para2,para4,para3] - Rresistance_popvariancestd0_timeseries[,para1,para2,para4,para3], rev(Rresistance_popvariancemean0_timeseries[,para1,para2,para4,para3] + Rresistance_popvariancestd0_timeseries[,para1,para2,para4,para3])), border=NA, col=adjustcolor("gray", alpha.f = 0.3))
          
          #legend("topright", legend = c("Behaviour", "Evolving", "Fixed", "Trait", "Choice", "Resistance"), col = c(NA,"black","black",NA,"black", "red"), text.font = 3, lwd = 3, lty=c(NA,1,2,NA,1,1), cex=1.5)
          
          #legend("topleft", legend = c("Trait", "Choice", "Resistance"), col = c(NA,"black", "red"), text.font = 3, lwd = 3, lty=c(1,1), cex=1.5)
          #legend("topright", legend = c("Behaviour", "Evolving", "Fixed"), col = c("black"), text.font = 3, lwd = 3, lty=c(NA,1,2),cex=1.5)
          
          dev.off()        
        }}}}
}


setwd(workingdirectory)
```
