## Supplementary material for "The role of evolving niche choice in herbivore adaptation to host plants": These are R scripts and data files used for generating results.: IBM_PlantHerbivore_SV.R

```
# Script contating main IBM model and other functions

mean2  <- function(x) mean(x,na.rm=T)
sd2  <- function(x) sd(x,na.rm=T)
std_error <- function(x) sd(x, na.rm = TRUE)/sqrt(length(!is.na(x)))

## Function for genotype to phenotype mapping
genotype2phenotype <- function(genotype,effectsizes,dominance,environmentaleffect){
  homozygote11 <- which(genotype == 2)
  heterozygotes <- which(genotype ==1)
  traitgenetic <- 0
  if (length(homozygote11)){
    traitgenetic <- traitgenetic + sum((genotype*effectsizes)[homozygote11])/2#MW: there was an error by a factor 2 because I had divided both the homozygotes and the heterozygotes by 2, but only the homozygotes should be divided by 2
  }
  if (length(heterozygotes)){
    traitgenetic <- traitgenetic + sum((genotype*effectsizes*dominance)[heterozygotes])
  }
  phenotype <- max(min(traitgenetic + rnorm(1,0,environmentaleffect),1),0)
  return(phenotype)
}


fecundityresistanceTradeoffs = function(fecunditycost, resistancetrait, fecundity0){
  offspring = (1+fecunditycost*resistancetrait)*fecundity0
  return(offspring)
}


writeParametersToFile <- function(ParameterList,ts.folder,sss){
  
  outfile = paste(ts.folder,"/Parameters_Set", set,".csv", sep = "")
  write.table(data.frame(t(unlist(ParameterList))), row.names = FALSE, col.names = sss<=1, append = sss>1, quote = FALSE, sep = ",", file = outfile)
  
}

# Main function

Beetle_Bigsimulation_fcb<-function(ts.folder, # The ts.folder is a folder supplied to store simulated RData in your current working folder
                                   Set=1,# to specify the set of simulation run under what parameters. This was introduced to enable split of simulations for same parameters when the number of replicates are too many to run at ago.
                                   FixedChoice = 0.5, # 0.5 is for random choice of plants/random behavior/no evolution of choice here
                                   numgenerations=200,#number of generations for which the simulation should run
                                   nsim=100,#number of replicates
                                   toxicity=c(0,0.5),#toxicity level of host plant 1 and 2
                                   initfreqresistance = 0.1,#initial frequency of more resistant alleles at the resistance loci
                                   initfreqchoice = 0.9,#initial frequency of host plant 2 choice at the choice loci
                                   resistancecost=-0.2,#will be multiplied by resistance level to get reduction in survival probability
                                   fecunditycost = 0, # between -1 and 1. Positive values imply resistance benefit, negative values imply resistance cost on fecundity
                                   choice="Avoid",
                                   SortMating=0.0,
                                   N_offspring=4,
                                   numresistanceloci  = 1, #number of loci underlying resistance to plant toxin
                                   numchoiceloci  = 1, #number of loci underlying host plant choice
                                   resistanceeffectsizes = c(1), #vector of effect sizes; length should correspond to the number of resistance loci; should add up to 1
                                   choiceeffectsizes = c(1),#vector of effect sizes; length should correspond to the number of choice loci; should add up to 1
                                   dominanceresistance = c(0.5),#dominance values for the resistance loci, should generally be between 0 and 1; length should correspond to the number of resistance loci
                                   dominancechoice = c(0.5),#dominance values for the choice loci, should generally be between 0 an 1; length should correspond to the number of choice loci
                                   environmentalsdresistance = 0.0,
                                   environmentalsdchoice = 0.0,
                                   plantfreqs  = c(0.5,0.5),#Should be of length 2. Here these are for toxic and non-toxic plants respectively
                                   scaledensity = 3, # to scale density dependent survival. Higher values above 1 implies higher survival rate
                                   recombinationrate = c(0.5), # Should be a vector of values between 0.5 and 1 and should be of length total loci minus 1
                                   N0  = 20, #initial population size
                                   K  = 1000 #global upper population size cap (population will be sampled down to this value at the end of every generation)#frequency of the two plants in the experimental setup
) {
  
  # Folder for saving data from simulated populations
  # We check if folder supplied is alreaded created, if yes, we just get a warning 
  if(!dir.exists(paste0(getwd(), "/",ts.folder))){
    dir.create(paste0(getwd(), "/",ts.folder))
    dir.create(paste0(getwd(), "/",ts.folder, "/", "Plots"))
  }else{
    #warning(paste0("Directory ", getwd(), "/",ts.folder, " already exists"))
  }
  
  # We avoid overwriting existing data.
  Parameters = paste0("fc", fecunditycost, "_hDr", scaledensity, "_AM",SortMating, "_DomR",dominanceresistance[1], "C", dominancechoice[1], 
                      "_rec", recombinationrate[1], "_rc", resistancecost, "_fr",round(initfreqresistance, 2), "fch", round(initfreqchoice, 2), 
                      "_tox", toxicity[2], "_lam", N_offspring, "_fxchoice", FixedChoice, "_Set",Set )
  
  outfileraw = paste0(getwd(), "/",ts.folder, "/", Parameters,"_", choice, "_SimPar.RData")
  if(file.exists(outfileraw)){
    print("Overwriting existing file\n")
    stop(paste0("file ", Parameters,"_", choice, "_SimPar.RData", " already exists in this directory"))
  }
  
  # Another checking!
  stopifnot(length(resistanceeffectsizes)==numresistanceloci | length(choiceeffectsizes)==numchoiceloci | length(dominancechoice)==numchoiceloci | length(dominanceresistance)==numresistanceloci)
  
  
  ## INPUT: a and b are two numeric vectors
  ## OUTPUT: a vector that at each index i contains the maximum of a[i] and b[i]
  max2 <- function(a,b){
    ifelse(a>b,a,b)
  }
  
  min2 <- function(a,b){
    ifelse(a<b,a,b)
  }
  
  lambda  <- N_offspring#average number of offspring per individual
  
  ##We assume that there are two alleles at each locus.
  ##Genetic parameters
  ##Number of loci 
  choiceeffectsizes <- choiceeffectsizes/sum(choiceeffectsizes)
  resistanceeffectsizes <- resistanceeffectsizes/sum(resistanceeffectsizes)
  totalnumloci  <- numresistanceloci+numchoiceloci
  
  #for(sim in 1:nsim){
  SimPar = mclapply(1:nsim, FUN = function(sim){
    
    resistance_tracking<-rep(NA, numgenerations+1)# Stores mean resistance trait
    choice_tracking<-rep(NA, numgenerations+1) # Stores mean choice/preference trait
    resistance_popvariance<-rep(NA, numgenerations+1) # Stores variance in resistance trait within the population
    choice_popvariance<-rep(NA, numgenerations+1)# Stores variance in preference trait within the population
    populationsizes_tracking<-rep(0, numgenerations+1) # Stores total population size
    N1 <- rep(NA, numgenerations+1)# Stores population size only at plant 1, the non-toxic host plant
    N2 <- rep(NA, numgenerations+1)# Stores total population size only at plant 2, the toxic host plant
    
    ##draw initial alleles from a Bernoulli distribution
    initialresistancealleles  <- (as.numeric(runif(2*numresistanceloci*N0) < initfreqresistance))
    initialchoicealleles  <- (as.numeric(runif(2*numchoiceloci*N0) < initfreqchoice))
    
    
    # ##initialize population matrix; the matrix has one column per locus
    populationarray = array(NA, dim = c(N0, numresistanceloci+numchoiceloci, 2) ) 
    populationarray[,1:numresistanceloci,] = initialresistancealleles
    populationarray[,(numresistanceloci+1):(numresistanceloci+numchoiceloci),] = initialchoicealleles
    
   
    ##prepare vectors for output and fill in the first entry
    populationsizes_tracking[1] = N0
    ##main generation loop
    for (t in 1:numgenerations){
      Ntotal  <- nrow(populationarray) #current total population size
      if (Ntotal > 0){
        # Convert genetypes/alleles to phenotypes/traits
        if (Ntotal > 1){
          preferences  <- apply(populationarray[,(numresistanceloci+1):(totalnumloci),1, drop = FALSE]+populationarray[,(numresistanceloci+1):(totalnumloci),2, drop = FALSE],1,function(x) genotype2phenotype(x,choiceeffectsizes,dominancechoice,environmentalsdchoice)) #the preference is the weighted average across choice loci, with the weight corresponding to the locus effect size
          resistance  <- apply(populationarray[,1:numresistanceloci,1, drop = FALSE]+populationarray[,1:numresistanceloci,2, drop = FALSE],1,function(x) genotype2phenotype(x,resistanceeffectsizes,dominanceresistance,environmentalsdresistance))
        } else {
          preferences  <- genotype2phenotype(populationarray[1,(numresistanceloci+1):(totalnumloci),1, drop = FALSE]+populationarray[1,(numresistanceloci+1):(totalnumloci),2, drop = FALSE],choiceeffectsizes,dominancechoice,environmentalsdchoice)
          resistance  <- genotype2phenotype(populationarray[,1:numresistanceloci,1, drop = FALSE]+populationarray[,1:numresistanceloci,2, drop = FALSE],resistanceeffectsizes,dominanceresistance,environmentalsdresistance)
        }
        
        
        resistance_tracking[t]<-c(mean(resistance)); 
        choice_tracking[t]<-c(mean(preferences));
        resistance_popvariance[t]<-c(var(resistance)); 
        choice_popvariance[t]<-c(var(preferences));
       
        # Niche choice behaviour implementation. Either we have avoidance, seeking, or random.
        if (plantfreqs[1]==0){#niche can only be chosen if it has non-zero frequency
          plant <- rep(2,Ntotal)
          N2[t] = length(plant)
          N1[t] = 0
        } else if (plantfreqs[2]==0){
          plant <- rep(1,Ntotal)
          N1[t] = length(plant)
          N2[t] = 0
        } else if (choice == "Avoid") { # to model avoidance behavior to the toxic host (#2);
          plant  <- (runif(Ntotal)<preferences)+1;
          plant<-abs(plant-3);
          N1[t] = length(plant[which(plant==1)])
          N2[t] = length(plant[which(plant==2)])
        } else if (choice == "Seek"){ # to model seeking behavior to the toxic host (#2);
          plant  <- (runif(Ntotal)<preferences)+1
          N1[t] = length(plant[which(plant==1)])
          N2[t] = length(plant[which(plant==2)])
        } else{                #let eggs be randomly distributed (manipulation treatment)
          plant  <- (runif(Ntotal)<FixedChoice)+1 #MW: replaced the second runif(Ntotal) by 0.5, same result but runs a bit faster because just one random number is created per individual
          N1[t] = length(plant[which(plant==1)])
          N2[t] = length(plant[which(plant==2)])
        }
        
        #- Reproduction -------------------
        Expected_offspring = fecundityresistanceTradeoffs(fecunditycost = fecunditycost, resistancetrait = resistance, fecundity0 = lambda)
        Actual_offspring = rpois(n = Ntotal, lambda = Expected_offspring)
        numoffspring = sum(Actual_offspring)
        mothervector = rep(1:Ntotal, Actual_offspring)
        
        if(sum(Actual_offspring)>K){
          mothervector = sample(mothervector, size = K, replace = FALSE)
          numoffspring = K
        }
        
        if (numoffspring > 0){
          ##initialize population matrix for next generation
          populationarraynew  <- array(NA,dim=c(numoffspring,totalnumloci,2));
          plant_offspring <- rep(NA,numoffspring)
          
          for(i in 1:numoffspring){
            Matedecision = runif(1)
            mother<-mothervector[i];
            mother.plant<-plant[mother]; # determine the host position:
            plant_offspring[i] = mother.plant
            
            if(Matedecision<=abs(SortMating) & SortMating>0){
              father = sample(which(plant==mother.plant), 1)
              
            }else if(Matedecision<=abs(SortMating) & SortMating<0 & N1[t]>0 & N2[t]>0){
              father = sample(which(plant!=mother.plant), 1)# different plant
              
            }else{ # the others are random;
              father<-sample(1:Ntotal,1); #randomly sample one of the surviving individuals as father
              
            }
            
            #Linkage included in the recombination rates
            ss = c(1,2)#sample(c(1,2), size = 2, replace = FALSE)
            if(length(unique(recombinationrate))<2){
              select.f  <- cbind(rep(father,totalnumloci),1:totalnumloci, sample(rep(ss,totalnumloci),totalnumloci,prob = c(rbind(recombinationrate, 1-recombinationrate)), replace=TRUE))
              select.m  <- cbind(rep(mother,totalnumloci),1:totalnumloci, sample(rep(ss,totalnumloci),totalnumloci,prob = c(rbind(recombinationrate, 1-recombinationrate)), replace=TRUE))
              
              
            }else{
              alleles_m = alleles_f = rep(NA, totalnumloci)
              for(l in 1:totalnumloci){
                alleles_m[l] = sample(ss,1,prob = c(recombinationrate[l], 1-recombinationrate[l]),replace=TRUE)
                alleles_f[l] = sample(ss,1,prob = c(recombinationrate[l], 1-recombinationrate[l]),replace=TRUE)
              }
              select.f  <- cbind(rep(father,totalnumloci),1:totalnumloci, alleles_f)
              select.m  <- cbind(rep(mother,totalnumloci),1:totalnumloci, alleles_m)
            }
            # ##pass on chosen allele copy at each locus according to recombination rate
            populationarraynew[i,,1] <- populationarray[select.f];
            populationarraynew[i,,2] <- populationarray[select.m];
          }
          
          if (nrow(populationarraynew) > 1){
            resistanceOffspring  <- apply(populationarraynew[,1:numresistanceloci,1, drop = FALSE]+populationarraynew[,1:numresistanceloci,2, drop = FALSE],1,function(x) genotype2phenotype(x,resistanceeffectsizes,dominanceresistance,environmentalsdresistance))
          } else {
            resistanceOffspring  <- genotype2phenotype(populationarraynew[,1:numresistanceloci,1, drop = FALSE]+populationarraynew[,1:numresistanceloci,2, drop = FALSE],resistanceeffectsizes,dominanceresistance,environmentalsdresistance)
          }
          ##determine how crowded each plant type is
          densitydependentsurvival  <- min2(max2(exp(0.01148851 - 0.03404447 * c(sum(plant_offspring==1),sum(plant_offspring==2))/scaledensity), 0),1) #from density dependence data, this expression is between 0 and 1 for all densities >= 1
          #densitydependentsurvival = c(1,1)
          
          ##the probability of dying depends on the plant's toxicity and on the individual's resistance, as well as on how crowded the plant is
          survivalprob  <- (1-toxicity[plant_offspring] * (1+resistanceOffspring))*densitydependentsurvival[plant_offspring]*max2(1-resistancecost*resistanceOffspring,0)
          survival  <- runif(nrow(populationarraynew)) < survivalprob #vector of booleans stating whether or not an individual survives
          numsurviving  <- sum(survival)
          survivors  <- which(survival)
          
          if(length(survivors)>0){
            plant = plant_offspring[survivors]
            populationarray = populationarraynew[survivors,,, drop=FALSE]
            
            # # Track population size
            populationsizes_tracking[t+1]<-nrow(populationarray)
            
          }else{
            # Update population array
            populationarray = array(0, dim = c(0,totalnumloci,2))
          }
        }else{
          populationarray = array(0, dim = c(0,totalnumloci,2))
        }
        
      }else{
        break
      }
      
    }
    
    return(list(resistance_tracking = resistance_tracking, choice_tracking = choice_tracking,
                resistance_popvariance = resistance_popvariance, choice_popvariance = choice_popvariance,
                populationsizes_tracking = populationsizes_tracking, N1 = N1, N2 = N2 ))
    
  }
  #}
  , mc.preschedule = FALSE, mc.cores = cores)
  # # Extracting and rearranging data from the function, SimPar 
 
  N1 = N2 = resistance_tracking = choice_tracking = resistance_popvariance = choice_popvariance = populationsizes_tracking = matrix(NA, nrow = (numgenerations+1), ncol = nsim)
  for(sim in 1:nsim){
   
    resistance_tracking[,sim] = SimPar[[sim]]$resistance_tracking
    choice_tracking[,sim] = SimPar[[sim]]$choice_tracking
    
    resistance_popvariance[,sim] = SimPar[[sim]]$resistance_popvariance
    choice_popvariance[,sim] = SimPar[[sim]]$choice_popvariance
    
    populationsizes_tracking[,sim] = SimPar[[sim]]$populationsizes_tracking
    N1[,sim] <- SimPar[[sim]]$N1
    N2[,sim] <- SimPar[[sim]]$N2
  }
  
  OUTPUT = list(resistance_tracking = resistance_tracking, choice_tracking = choice_tracking, resistance_popvariance = resistance_popvariance, choice_popvariance = choice_popvariance, populationsizes_tracking = populationsizes_tracking, N1 = N1, N2 = N2)
 
  # Save data as RData objects
  
  saveRDS(OUTPUT, file=outfileraw)
  
  return(OUTPUT)
  
}
```
