## Supplementary material for "The role of evolving niche choice in herbivore adaptation to host plants": These are R scripts and data files used for generating results.: MetaAnalysisData_EachSpecies_SV.R

```
#Make sure that at the end of running this script, you run the last line of this script to reset the working directory
# This script uploads the data from the Excell file, analyses it, makes plots and saves them, and saves the parameter distributions that will be used for sampling simulations. 

# Clear memory
rm(list=ls())
pacman::p_load(openxlsx, dplyr, tidyr, ggplot2, patchwork) 

workingdirectory = getwd()
WD = "PLANTHERBIVORECHOICE"
setwd(paste0(workingdirectory,"/",WD))

##
##
##### Script aim: #####
### 1. To calculate average values of all parameters used in the model in Nabutanyi et. al. 2023
### 2. To plot the distributions of those parameters 
##
##

##
##### Libraries & data #####
##

ParameterDataFolder = "EmpiricalData"

if(!dir.exists(paste0(getwd(), "/", ParameterDataFolder))){
  dir.create(paste0(getwd(), "/", ParameterDataFolder))
}


PlotsParameterDistributionFolder = "PlotsParameterDistributions"

if(!dir.exists(paste0(getwd(), "/", PlotsParameterDistributionFolder))){
  dir.create(paste0(getwd(), "/", PlotsParameterDistributionFolder))
}


#- Convert habitat mating preference to model interpretation range--
ConversionFunc_AM = function(xvalue, OldMax, OldMin, NewMax, NewMin){
  OldRange = (OldMax - OldMin)  
  NewRange = (NewMax - NewMin)  
  NewValue = (((xvalue - OldMin) * NewRange) / OldRange) + NewMin
  return(NewValue)
}

# Convert the choice interval from (-1,1) to (0,1)
ConversionFunc_Choice = function(xvalue, OldMax, OldMin, NewMax, NewMin){
  OldRange = (OldMax - OldMin)  
  NewRange = (NewMax - NewMin)  
  NewValue = (((xvalue - OldMin) * NewRange) / OldRange) + NewMin
  return(NewValue)
}

##
## Divide the sheets of the workbook into five dataframes 
##

#-- Loading data from excel sheets
{
df1  <- read.xlsx("S7.xlsx",
                  colNames=T,
                  sheet = 1)
df2 <- read.xlsx("S7.xlsx",
                 colNames=T,
                 sheet = 2)
df3 <- read.xlsx("S7.xlsx",
                 colNames=T,
                 sheet = 3)
df4 <- read.xlsx("S7.xlsx",
                 colNames=T,
                 sheet = 4)
df5 <- read.xlsx("S7.xlsx",
                 colNames=T,
                 sheet = 5)
df6 <- read.xlsx("S7.xlsx",
                 colNames=T,
                 sheet = 6)
## initial data exploration 
#
}
ranki = 1
binsize = 20
for(ranki in 1:10){
  
  # # exploring T_Mortality
  # table(df1$T_Mortality_Mean[df1$SpeciesRank==ranki])
  # table(is.na(df1$T_Mortality_Mean[df1$SpeciesRank==ranki])) 
  
  df1S <- df1 %>% filter(df1$Resistance == "S",df1$SpeciesRank==ranki)
  
  df1S2 <- df1 %>% filter(df1$Resistance == "S", df1$SpeciesRank==ranki, df1$C_Mortality_Mean != "NA")
  
  df1R <- df1 %>% filter(df1$Resistance == "R", df1$SpeciesRank==ranki) 
  df1R2 <- df1 %>% filter(df1$Resistance == "R", df1$SpeciesRank==ranki, df1$C_Mortality_Mean != "NA")
  
  
  ### Aim 1. To calculate average values of all parameters
  
  ## From sheet 1: survival / resistance data
  
  ## A) Calculating toxicity level of plant 2 (toxic)
  
  T_sd<-sd(df1S$T_Mortality_Mean, na.rm = T)
  tau2<-c(df1S$T_Mortality_Mean)
  tau2 <- tau2[!is.na(tau2)] #vector with all toxicity levels of plant 2 (toxic)
  #try(hist(tau2))
  
  ## B) Calculating toxicity level of plant 1 (nontoxic)
  
  C_sd<-sd(df1S$C_Mortality_Mean, na.rm = T)
  tau1<-c(as.numeric(df1S$C_Mortality_Mean))
  tau1 <- tau1[!is.na(tau1)] #vector with all toxicity levels of plant 1 (nontoxic)
  
  ## Correcting for control mortality using Abbott's Formula
  df1corrected <- df1 %>% filter(df1$T_Mortality_Mean != "NA" & df1$C_Mortality_Mean != "NA", df1$SpeciesRank==ranki)
  
  df1corrected$T_Corrected_Mortality<- (df1corrected$T_Mortality_Mean-as.numeric(df1corrected$C_Mortality_Mean))*100/(100-as.numeric(df1corrected$C_Mortality_Mean))
  
  df1ScorrectedS <- df1corrected %>% filter(df1corrected$Resistance == "S")
  
  new_T_sd<-sd(df1ScorrectedS$T_Corrected_Mortality, na.rm = T)
  newtau2<-c(df1ScorrectedS$T_Corrected_Mortality)
  newtau2 <- newtau2[!is.na(newtau2)] #vector with all toxicity levels of plant 2 (toxic)
 
  
  ## C) Calculating resistance cost
  
  #To calculate cost, creating modified dataframes containing only papers with mortalities for a susceptible AND a resistant strain
  df1S2$PaperWise_SerialNo<-with(df1S2,ave(as.character(PaperID),PaperID,FUN=seq_along))
  df1S2$PaperIDnew<-paste0(df1S2$PaperID, df1S2$PaperWise_SerialNo)
  df1R2$PaperWise_SerialNo<-with(df1R2,ave(as.character(PaperID),PaperID,FUN=seq_along))
  df1R2$PaperIDnew<-paste0(df1R2$PaperID, df1R2$PaperWise_SerialNo)
  newdf<-merge(df1S2, df1R2, by = "PaperIDnew")
  
  
  newdf$cost<-1-((100-as.numeric(newdf$C_Mortality_Mean.x))/(100-as.numeric(newdf$C_Mortality_Mean.y)))
  #newdf<-newdf[-c(79), ]  ##removed -Inf row
  mean(newdf$cost, na.rm = T)
  cost_sd<-sd(newdf$cost, na.rm = T)
  res_cost<-newdf$cost #vector with all resistance costs
  Rc_model = 1/(1-res_cost) - 1#((as.numeric(newdf$C_Mortality_Mean.x))/100)/(-1+as.numeric(res_cost))
  Rc_model1 = round(Rc_model[is.finite(Rc_model)],2)
  
  ## From sheet 2: choice data
  
  choice_sd<-sd(df2$CI[df2$SpeciesRank==ranki], na.rm = T)
  choice<-df2$CI[df2$SpeciesRank==ranki] #vector with all choice indices: -1 means a preference for nontoxic plants
  
  
  ## From sheet 3: Mating choice data
  
  mating_sd<-sd(df3$Same_prop[df3$SpeciesRank==ranki], na.rm = T)
  Mating<-df3$Same_prop[df3$SpeciesRank==ranki] #vector with all mating choices: 0 is random mating
  
  
  ## From sheet 4: Genetics data
  
  res_alleles<-df4$Freq_Res_alleles[df4$SpeciesRank==ranki] #vector with all frequencies of resistance loci
  res_loci<-df4$n_Res_loci[df4$SpeciesRank==ranki] #vector with all frequencies of resistance loci
  
  
  ## From sheet 5: Fecundity data
  df5S <- df5 %>% filter(df5$Resistance == "S", df5$Plant == "C", df5$SpeciesRank==ranki)
  df5R <- df5 %>% filter(df5$Resistance == "R", df5$Plant == "C",df5$SpeciesRank==ranki) 
  
  #To calculate fecundity cost, creating modified dataframes containing only papers with fecundities for a susceptible AND a resistant strain
  df5S$PaperWise_SerialNo<-with(df5S,ave(as.character(PaperID),PaperID,FUN=seq_along))
  df5S$PaperIDnew<-paste0(df5S$PaperID, df5S$PaperWise_SerialNo)
  df5R$PaperWise_SerialNo<-with(df5R,ave(as.character(PaperID),PaperID,FUN=seq_along))
  df5R$PaperIDnew<-paste0(df5R$PaperID, df5R$PaperWise_SerialNo)
  newdf5<-merge(df5S, df5R, by = "PaperIDnew")
  
  newdf5$cost<-1-(newdf5$`Eggs/female.x`/newdf5$`Eggs/female.y`)
  sd_Fcost<-sd(newdf5$cost, na.rm = T)
  
  fec_cost<-df5R$cost
  
  fec_sd<-sd(df5S$`Eggs/female`, na.rm = T)
  lambda<-df5S$`Eggs/female`
  
    #-------------------------------------Plots ---------
  
  (fc_cost = newdf5$cost)#df5R2$cost)
  nresistanceloci = df4$n_Res_loci[df4$SpeciesRank==ranki]
  freq_res_loci = round(df4$Freq_Res_alleles[df4$SpeciesRank==ranki],2)
  
  png(paste0(getwd(),"/" ,PlotsParameterDistributionFolder,"/","ChoiceIndex", "_SpRank_", ranki, ".png"))
  par(mar = c(5, 5, 4, 2) + 0.3)
  try(hist(choice, cex.axis = 2, xlab = list(expression(Fixed ~plant ~choice ~z["c,fixed"]), font = 3, cex = 2), ylab = list("Count", font = 3, cex = 2), main = "", xlim = c(-1,1),breaks = seq(-1, 1, length.out = binsize)))
  dev.off()
  
  png(paste0(getwd(),"/" ,PlotsParameterDistributionFolder,"/","Toxicity1", "_SpRank_", ranki, ".png"))
  par(mar = c(5, 5, 4, 2) + 0.3)
  try(hist(tau1/100, cex.axis = 2, xlab = list(expression(Plant ~1 ~toxicity~tau[1]), font = 3, cex = 2), ylab = list("Count", font = 3, cex = 2), main = "", xlim = c(0,1),breaks = seq(0, 1, length.out = binsize)))
  dev.off()
  
  png(paste0(getwd(),"/" ,PlotsParameterDistributionFolder,"/","Toxicity2", "_SpRank_", ranki, ".png"))
  par(mar = c(5, 5, 4, 2) + 0.3)
  try(hist(tau2/100, cex.axis = 2, xlab = list(expression(Plant ~2 ~toxicity~tau[2]), font = 3, cex = 2), ylab = list("Count", font = 3, cex = 2), main = "", xlim = c(0,1),breaks = seq(0, 1, length.out = binsize)))
  dev.off()
  
  png(paste0(getwd(),"/" ,PlotsParameterDistributionFolder,"/","ResistanceCost", "_SpRank_", ranki, ".png"))
  par(mar = c(5, 5, 4, 2) + 0.3)
  try(hist(res_cost, cex.axis = 2, xlab = list(expression(Resistance ~ effect ~ to ~ viability ~r[c]), font = 3, cex = 2), ylab = list("Count", font = 3, cex = 2), main = "", xlim = c(-3,1.5),breaks = seq(-3, 1.5, length.out = binsize)))
  dev.off()
  
  png(paste0(getwd(),"/" ,PlotsParameterDistributionFolder,"/","FecundityCost", "_SpRank_", ranki, ".png"))
  par(mar = c(5, 5, 4, 2) + 0.3)
  try(hist(fc_cost, cex.axis = 2, xlab = list(expression(Resistance ~ effect ~ to ~ fecundity~f[c]), font = 3, cex = 2), ylab = list("Count", font = 3, cex = 2), main = "", xlim = c(-5,1.5),breaks = seq(-5, 1.5, length.out = binsize)))
  dev.off()
  
  png(paste0(getwd(),"/" ,PlotsParameterDistributionFolder,"/","Eggs", "_SpRank_", ranki, ".png"))
  par(mar = c(5, 5, 4, 2) + 0.3)
  try(hist(lambda, cex.axis = 2, xlab = list(expression(Mean ~ number ~ of ~ eggs), font = 3, cex = 2), ylab = list("Count", font = 3, cex = 2), main = "", xlim = c(0,3000),breaks = seq(0, 3000, length.out = binsize)))
  dev.off()
  
  png(paste0(getwd(),"/" ,PlotsParameterDistributionFolder,"/","Eggs_converted", "_SpRank_", ranki, ".png"))
  par(mar = c(5, 5, 4, 2) + 0.3)
  try(hist(lambda/100, cex.axis = 2, xlab = list(expression(Mean ~ number ~ of ~ offspring~Lambda), font = 3, cex = 2), ylab = list("Count", font = 3, cex = 2), main = "", xlim = c(0,30),breaks = seq(0, 30, length.out = binsize)))
  dev.off()
  
  png(paste0(getwd(),"/" ,PlotsParameterDistributionFolder,"/","AssortativeMating", "_SpRank_", ranki, ".png"))
  par(mar = c(5, 5, 4, 2) + 0.3)
  try(hist(Mating, cex.axis = 2, xlab = list(expression(Host ~ mating ~ preference ~ M[a]), font = 3, cex = 2), ylab = list("Count", font = 3, cex = 2), main = "", xlim = c(0,1),breaks = seq(0, 1, length.out = binsize)))
  dev.off()
  
  png(paste0(getwd(),"/" ,PlotsParameterDistributionFolder,"/","FrequencyResistance", "_SpRank_", ranki, ".png"))
  par(mar = c(5, 5, 4, 2) + 0.3)
  try(hist(freq_res_loci, cex.axis = 2, xlab = list(expression(Resistant ~allele~ frequency ~p[r0]), font = 3, cex = 2), ylab = list("Count", font = 3, cex = 2), main = "", xlim = c(0,1),breaks = seq(0, 1, length.out = binsize)))
  dev.off()
  
  png(paste0(getwd(),"/" ,PlotsParameterDistributionFolder,"/","nResistanceLoci", "_SpRank_", ranki, ".png"))
  par(mar = c(5, 5, 4, 2) + 0.3)
  try(hist(nresistanceloci, cex.axis = 2, xlab = list(expression(Resistance~ loci ~ n[r]), font = 3, cex = 2), ylab = list("Count", font = 3, cex = 2), main = "", xlim = c(0,5),breaks = seq(0, 5, length.out = binsize)))
  dev.off()
  
  
  
  ## From sheet 2: oviposition choice data
  
 # try(hist(df2$CI_Mean[df2$SpeciesRank==ranki]))
  Choice<-df2$CI_Mean[df2$SpeciesRank==ranki] #vector with all mating choices: 0 is random mating
  New_Choice = round(ConversionFunc_Choice(xvalue = df2$CI_Mean[df2$SpeciesRank==ranki], OldMax = 1, OldMin=-1, NewMax=1, NewMin=0),2)
  
  ## From sheet 3: Mating choice data
  
  #try(hist(df3$Same_prop[df3$SpeciesRank==ranki]))
  Mating<-df3$Same_prop[df3$SpeciesRank==ranki] #vector with all mating choices: 0 is random mating
  New_AM = round(ConversionFunc_AM(xvalue = df3$Same_prop[df3$SpeciesRank==ranki], OldMax = 1, OldMin=0, NewMax=1, NewMin=-1),2)
  
  png(paste0(getwd(),"/" ,PlotsParameterDistributionFolder,"/","ChoiceIndexConverted", "_SpRank_", ranki, ".png"))
  par(mar = c(5, 5, 4, 2) + 0.3)
  try(hist(1-New_Choice, cex.axis = 2, xlab = list(expression(Fixed ~plant ~choice ~z["c,fixed"]), font = 3, cex = 2), ylab = list("Count", font = 3, cex = 2), main = "", xlim = c(0,1),breaks = seq(0, 1, length.out = binsize)))
  dev.off()
  
  png(paste0(getwd(),"/" ,PlotsParameterDistributionFolder,"/","AssortativeMatingConverted", "_SpRank_", ranki, ".png"))
  par(mar = c(5, 5, 4, 2) + 0.3)
  try(hist(New_AM, cex.axis = 2, xlab = list(expression(Host~ mating ~ preference ~ M[a]), font = 3, cex = 2), ylab = list("Count", font = 3, cex = 2), main = "", xlim = c(-1,1),breaks = seq(-1, 1, length.out = binsize)))
  dev.off()
  
  #-------------- Toxicity at plant 2 - Relative
  #------------ Get relative toxicity at the toxic plant. This assumes non-toxic plant is completely non-toxic
  Tau1 = as.numeric(df1$C_Mortality_Mean[which(df1$T_Mortality_Mean != "NA" & df1$C_Mortality_Mean != "NA" & df1$Resistance=="S" & df1$SpeciesRank==ranki)]) # Only cases provided for toxicity at both plants
  Tau2 = as.numeric(df1$T_Mortality_Mean[which(df1$T_Mortality_Mean != "NA" & df1$C_Mortality_Mean != "NA" & df1$Resistance=="S" & df1$SpeciesRank==ranki)])
  Tau_1 = 100-Tau1[which(!is.na(Tau1) & !is.na(Tau2))]# First calculate survival at plant 1
  Tau_2 = 100-Tau2[which(!is.na(Tau1) & !is.na(Tau2))]# First calculate survival at plant 2
  #sum(Tau_1>Tau_2)
  
  Ttau1 = round(1-(Tau_1/Tau_1)[Tau_1>=Tau_2],2)# Now toxicity at plant 1 is 0 but remove those that have more toxicity at plant 1
  Ttau2 = round(1-(Tau_2/Tau_1)[Tau_1>=Tau_2], 2) # Relative toxicity of plant 2
  
  png(paste0(getwd(),"/" ,PlotsParameterDistributionFolder,"/","Toxicity2_Relative", "_SpRank_", ranki, ".png"))
  par(mar = c(5, 5, 4, 2) + 0.3)
  try(hist(Ttau2, cex.axis = 2, xlab = list(expression(Plant ~2 ~relative ~ toxicity~tau[2]), font = 3, cex = 2), ylab = list("Count", font = 3, cex = 2), main = "", xlim = c(0,1),breaks = seq(0, 1, length.out = binsize)))
  dev.off()
  
  
  #------------ Fecundity cost ----------------------
  # Change fecundity costs/benefits to model range
  fc_cost = newdf5$cost
  Fc_model = round((1/(1-fc_cost))-1,2)
  Fc_model1 = Fc_model[is.finite(Fc_model)]
  fc_cost1 = fc_cost[is.finite(fc_cost)]
  
  png(paste0(getwd(),"/" ,PlotsParameterDistributionFolder,"/","FecundityCost_Model", "_SpRank_", ranki, ".png"))
  par(mar = c(5, 5, 4, 2) + 0.3)
  try(hist(Fc_model1, cex.axis = 2, xlab = list(expression(Resistance ~ effect ~ to ~ fecundity ~f[c]), font = 3, cex = 2), ylab = list("Count", font = 3, cex = 2), main = "", xlim = c(-1,1),breaks = seq(-1, 1, length.out = binsize)))
  dev.off()
  
  #--------------- Resistance cost -------
  #- Change viability costs/benefits to model range
  
  newdf$cost<-1-((100-as.numeric(newdf$C_Mortality_Mean.x))/(100-as.numeric(newdf$C_Mortality_Mean.y)))
  #newdf<-newdf[-c(79), ]  ##removed -Inf row
  mean(newdf$cost, na.rm = T)
  cost_sd<-sd(newdf$cost, na.rm = T)
  res_cost<-newdf$cost #vector with all resistance costs
  Rc_model = 1/(1-res_cost) - 1#((as.numeric(newdf$C_Mortality_Mean.x))/100)/(-1+as.numeric(res_cost))
  Rc_model1 = round(Rc_model[is.finite(Rc_model)],2)
  
  
  
  png(paste0(getwd(),"/" ,PlotsParameterDistributionFolder,"/","ResistanceCost_Model", "_SpRank_", ranki, ".png"))
  par(mar = c(5, 5, 4, 2) + 0.3)
  try(hist(Rc_model1, cex.axis = 2, xlab = list(expression(Resistance ~ effect ~ to ~ viability ~ r[c]), font = 3, cex = 2), ylab = list("Count", font = 3, cex = 2), main = "", xlim = c(-1,1),breaks = seq(-2, 2, length.out = binsize)))
  dev.off()
  
  write.csv(1-New_Choice[is.finite(New_Choice)], file = paste0(getwd(), "/", ParameterDataFolder,"/","ChoiceIndex", "_SpRank_", ranki, ".csv"), sep = "", quote = TRUE )
  write.csv(Rc_model1, file = paste0(getwd(), "/", ParameterDataFolder,"/","ViabilityCost", "_SpRank_", ranki, ".csv"), sep = "", quote = TRUE )
  write.csv(Fc_model1, file = paste0(getwd(), "/", ParameterDataFolder,"/","FecundityCost", "_SpRank_", ranki, ".csv"), sep = "", quote = FALSE)
  write.csv(Ttau2[is.finite(Ttau2)], file = paste0(getwd(), "/", ParameterDataFolder,"/","Toxicity2", "_SpRank_", ranki, ".csv"), sep = "", quote = FALSE)
  write.csv(round((lambda/100)[is.finite(lambda)],2), file = paste0(getwd(), "/", ParameterDataFolder,"/","Lambda", "_SpRank_", ranki, ".csv"), sep = "", quote = FALSE)
  write.csv(New_AM[is.finite(New_AM)], file = paste0(getwd(), "/", ParameterDataFolder,"/","AssortativeMating", "_SpRank_", ranki, ".csv"), sep = "", quote = FALSE)
  write.csv(freq_res_loci[is.finite(freq_res_loci)], file = paste0(getwd(), "/", ParameterDataFolder,"/","Fr0", "_SpRank_", ranki, ".csv"), sep = "", quote = FALSE)
}


#------------ Species 11: Artificially created with all data combined --------------------------

{
# Parameter distributions put together

df1S <- df1 %>% filter(df1$Resistance == "S")
df1S2 <- df1 %>% filter(df1$Resistance == "S", df1$C_Mortality_Mean != "NA")

df1R <- df1 %>% filter(df1$Resistance == "R") 
df1R2 <- df1 %>% filter(df1$Resistance == "R", df1$C_Mortality_Mean != "NA")


### Aim 1. To calculate average values of all parameters

## From sheet 1: survival / resistance data

## A) Calculating toxicity level of plant 2 (toxic)

# summary(df1S$T_Mortality_Mean, na.rm= T)
# mean(df1S$T_Mortality_Mean, na.rm = T)
T_sd<-sd(df1S$T_Mortality_Mean, na.rm = T)
#colSums(!is.na(df1S))
#T_sd/sqrt(267)
tau2<-c(df1S$T_Mortality_Mean)
tau2 <- tau2[!is.na(tau2)] #vector with all toxicity levels of plant 2 (toxic)
#hist(tau2)

## B) Calculating toxicity level of plant 1 (nontoxic)

#summary(as.numeric(df1S$C_Mortality_Mean, na.rm= T))
#mean(as.numeric(df1S$C_Mortality_Mean, na.rm = T))
C_sd<-sd(df1S$C_Mortality_Mean, na.rm = T)
#colSums(!is.na(df1S))
#C_sd/sqrt(230)
tau1<-c(as.numeric(df1S$C_Mortality_Mean))
tau1 <- tau1[!is.na(tau1)] #vector with all toxicity levels of plant 1 (nontoxic)
#hist(tau1)

## Correcting for control mortality using Abbott's Formula
df1corrected <- df1 %>% filter(df1$T_Mortality_Mean != "NA" & df1$C_Mortality_Mean != "NA")

df1corrected$T_Corrected_Mortality<- (df1corrected$T_Mortality_Mean-as.numeric(df1corrected$C_Mortality_Mean))*100/(100-as.numeric(df1corrected$C_Mortality_Mean))

df1ScorrectedS <- df1corrected %>% filter(df1corrected$Resistance == "S")

#summary(df1ScorrectedS$T_Corrected_Mortality, na.rm= T)
#mean(df1ScorrectedS$T_Corrected_Mortality, na.rm = T)
new_T_sd<-sd(df1ScorrectedS$T_Corrected_Mortality, na.rm = T)
#colSums(!is.na(df1ScorrectedS))
#new_T_sd/sqrt(169)
newtau2<-c(df1ScorrectedS$T_Corrected_Mortality)
newtau2 <- newtau2[!is.na(newtau2)] #vector with all toxicity levels of plant 2 (toxic)
#hist(newtau2)

## C) Calculating resistance cost

#To calculate cost, creating modified dataframes containing only papers with mortalities for a susceptible AND a resistant strain
df1S2$PaperWise_SerialNo<-with(df1S2,ave(as.character(PaperID),PaperID,FUN=seq_along))
df1S2$PaperIDnew<-paste0(df1S2$PaperID, df1S2$PaperWise_SerialNo)
df1R2$PaperWise_SerialNo<-with(df1R2,ave(as.character(PaperID),PaperID,FUN=seq_along))
df1R2$PaperIDnew<-paste0(df1R2$PaperID, df1R2$PaperWise_SerialNo)
newdf<-merge(df1S2, df1R2, by = "PaperIDnew")


newdf$cost<-1-((100-as.numeric(newdf$C_Mortality_Mean.x))/(100-as.numeric(newdf$C_Mortality_Mean.y)))
#newdf<-newdf[-c(79), ]  ##removed -Inf row
mean(newdf$cost, na.rm = T)
cost_sd<-sd(newdf$cost, na.rm = T)
res_cost<-newdf$cost #vector with all resistance costs
Rc_model = 1/(1-res_cost) - 1#((as.numeric(newdf$C_Mortality_Mean.x))/100)/(-1+as.numeric(res_cost))
Rc_model1 = round(Rc_model[is.finite(Rc_model)],2)

#hist(Rc_model)

## From sheet 2: choice data

#head(df2)
#summary(df2$CI, na.rm= T)
#mean(df2$CI, na.rm = T)
choice_sd<-sd(df2$CI, na.rm = T)
#colSums(!is.na(df2))
#choice_sd/sqrt(171)
#hist(df2$CI)
choice<-df2$CI #vector with all choice indices: -1 means a preference for nontoxic plants


## From sheet 3: Mating choice data

#head(df3)
#summary(df3$Same_prop)
#mean(df3$Same_prop, na.rm = T)
mating_sd<-sd(df3$Same_prop, na.rm = T)
#colSums(!is.na(df3))
#mating_sd/sqrt(36)
#hist(df3$Same_prop)
Mating<-df3$Same_prop #vector with all mating choices: 0 is random mating


## From sheet 4: Genetics data

#hist(df4$Freq_Res_alleles)
res_alleles<-df4$Freq_Res_alleles #vector with all frequencies of resistance loci

## From sheet 5: Fecundity data

#head(df5)
df5S <- df5 %>%
  filter(df5$Resistance == "S", df5$Plant == "C")
df5R <- df5 %>%
  filter(df5$Resistance == "R", df5$Plant == "C") 

#To calculate fecundity cost, creating modified dataframes containing only papers with fecundities for a susceptible AND a resistant strain
df5S$PaperWise_SerialNo<-with(df5S,ave(as.character(PaperID),PaperID,FUN=seq_along))
df5S$PaperIDnew<-paste0(df5S$PaperID, df5S$PaperWise_SerialNo)
df5R$PaperWise_SerialNo<-with(df5R,ave(as.character(PaperID),PaperID,FUN=seq_along))
df5R$PaperIDnew<-paste0(df5R$PaperID, df5R$PaperWise_SerialNo)
newdf5<-merge(df5S, df5R, by = "PaperIDnew")

newdf5$cost<-1-(newdf5$`Eggs/female.x`/newdf5$`Eggs/female.y`)
#summary(newdf5$cost, na.rm= T)
#mean(newdf5$cost, na.rm = T)
sd_Fcost<-sd(newdf5$cost, na.rm = T)
#colSums(!is.na(newdf5))
#sd_Fcost/sqrt(120)

#hist(newdf5$cost)
fec_cost<-df5R$cost

#summary(df5S$`Eggs/female`, na.rm= T)
#mean(df5S$`Eggs/female`, na.rm = T)
fec_sd<-sd(df5S$`Eggs/female`, na.rm = T)
#colSums(!is.na(df5S))
#fec_sd/sqrt(169)
#hist(df5S$`Eggs/female`)
lambda<-df5S$`Eggs/female`


res_loci = df4$n_Res_loci


#-------------------------------------Plots ---------

(fc_cost = newdf5$cost)#df5R2$cost)
nresistanceloci = df4$n_Res_loci
freq_res_loci = round(df4$Freq_Res_alleles,2)


png(paste0(getwd(),"/" ,PlotsParameterDistributionFolder,"/","ChoiceIndex.png"))
par(mar = c(5, 5, 4, 2) + 0.3)
try(hist(choice, cex.axis = 2, xlab = list(expression(Fixed ~plant ~choice ~z["c,fixed"]), font = 3, cex = 2), ylab = list("Count", font = 3, cex = 2), main = ""))
dev.off()

png(paste0(getwd(),"/" ,PlotsParameterDistributionFolder,"/","Toxicity1.png"))
par(mar = c(5, 5, 4, 2) + 0.3)
hist(tau1/100, cex.axis = 2, xlab = list(expression(Plant ~1 ~toxicity~tau[1]), font = 3, cex = 2), ylab = list("Count", font = 3, cex = 2), main = "")
dev.off()

png(paste0(getwd(),"/" ,PlotsParameterDistributionFolder,"/","Toxicity2.png"))
par(mar = c(5, 5, 4, 2) + 0.3)
hist(tau2/100, cex.axis = 2, xlab = list(expression(Plant ~2 ~toxicity~tau[2]), font = 3, cex = 2), ylab = list("Count", font = 3, cex = 2), main = "")
dev.off()

png(paste0(getwd(),"/" ,PlotsParameterDistributionFolder,"/","ResistanceCost.png"))
par(mar = c(5, 5, 4, 2) + 0.3)
hist(res_cost, cex.axis = 2, xlab = list(expression(Resistance ~ effect ~ to ~ viability ~r[c]), font = 3, cex = 2), ylab = list("Count", font = 3, cex = 2), main = "")
dev.off()

png(paste0(getwd(),"/" ,PlotsParameterDistributionFolder,"/","FecundityCost.png"))
par(mar = c(5, 5, 4, 2) + 0.3)
hist(fc_cost, cex.axis = 2, xlab = list(expression(Resistance ~ effect ~ to ~ fecundity~f[c]), font = 3, cex = 2), ylab = list("Count", font = 3, cex = 2), main = "")
dev.off()

png(paste0(getwd(),"/" ,PlotsParameterDistributionFolder,"/","Eggs_converted.png"))
par(mar = c(5, 5, 4, 2) + 0.3)
hist(lambda/100, cex.axis = 2, xlab = list(expression(Mean ~ number ~ of ~ offspring~Lambda), font = 3, cex = 2), ylab = list("Count", font = 3, cex = 2), main = "")
dev.off()

png(paste0(getwd(),"/" ,PlotsParameterDistributionFolder,"/","AssortativeMating.png"))
par(mar = c(5, 5, 4, 2) + 0.3)
hist(Mating, cex.axis = 2, xlab = list(expression(Host ~ mating ~ preference ~ M[a]), font = 3, cex = 2), ylab = list("Count", font = 3, cex = 2), main = "")
dev.off()

png(paste0(getwd(),"/" ,PlotsParameterDistributionFolder,"/","FrequencyResistance.png"))
par(mar = c(5, 5, 4, 2) + 0.3)
hist(freq_res_loci, cex.axis = 2, xlab = list(expression(Resistant ~allele~ frequency ~p[r0]), font = 3, cex = 2), ylab = list("Count", font = 3, cex = 2), main = "")
dev.off()

png(paste0(getwd(),"/" ,PlotsParameterDistributionFolder,"/","nResistanceLoci.png"))
par(mar = c(5, 5, 4, 2) + 0.3)
hist(nresistanceloci, cex.axis = 2, xlab = list(expression(Resistance~ loci ~ n[r]), font = 3, cex = 2), ylab = list("Count", font = 3, cex = 2), main = "")
dev.off()

## From sheet 2: oviposition choice data
Choice<-df2$CI_Mean #vector with all mating choices: 0 is random mating
New_Choice = round(ConversionFunc_Choice(xvalue = df2$CI_Mean, OldMax = 1, OldMin=-1, NewMax=1, NewMin=0),2)


## From sheet 3: Mating choice data
Mating<-df3$Same_prop #vector with all mating choices: 0 is random mating
New_AM = round(ConversionFunc_AM(xvalue = df3$Same_prop, OldMax = 1, OldMin=0, NewMax=1, NewMin=-1),2)

png(paste0(getwd(),"/" ,PlotsParameterDistributionFolder,"/","ChoiceIndexConverted.png"))
par(mar = c(5, 5, 4, 2) + 0.3)
try(hist(1-New_Choice, cex.axis = 2, xlab = list(expression(Fixed ~plant ~choice ~z["c,fixed"]), font = 3, cex = 2), ylab = list("Count", font = 3, cex = 2), main = ""))
dev.off()

png(paste0(getwd(),"/" ,PlotsParameterDistributionFolder,"/","AssortativeMatingConverted.png"))
par(mar = c(5, 5, 4, 2) + 0.3)
hist(New_AM, cex.axis = 2, xlab = list(expression(Host~ mating ~ preference ~ M[a]), font = 3, cex = 2), ylab = list("Count", font = 3, cex = 2), main = "")
dev.off()

#-------------- Toxicity at plant 2 - Relative
Tau1 = as.numeric(df1$C_Mortality_Mean[which(df1$T_Mortality_Mean != "NA" & df1$C_Mortality_Mean != "NA" & df1$Resistance=="S")]) # Only cases provided for toxicity at both plants
Tau2 = as.numeric(df1$T_Mortality_Mean[which(df1$T_Mortality_Mean != "NA" & df1$C_Mortality_Mean != "NA" & df1$Resistance=="S")])
Tau_1 = 100-Tau1[which(!is.na(Tau1) & !is.na(Tau2))]# First calculate survival at plant 1
Tau_2 = 100-Tau2[which(!is.na(Tau1) & !is.na(Tau2))]# First calculate survival at plant 2
#sum(Tau_1>Tau_2)

Ttau1 = round(1-(Tau_1/Tau_1)[Tau_1>=Tau_2],2)# Now toxicity at plant 1 is 0 but remove those that have more toxicity at plant 1
Ttau2 = round(1-(Tau_2/Tau_1)[Tau_1>=Tau_2], 2) # Relative toxicity of plant 2


png(paste0(getwd(),"/" ,PlotsParameterDistributionFolder,"/","Toxicity2_Relative.png"))
par(mar = c(5, 5, 4, 2) + 0.3)
hist(Ttau2, cex.axis = 2, xlab = list(expression(Plant ~2 ~relative ~ toxicity~tau[2]), font = 3, cex = 2), ylab = list("Count", font = 3, cex = 2), main = "")
dev.off()


#------------ Fecundity cost ----------------------
fc_cost = newdf5$cost
Fc_model = round((1/(1-fc_cost))-1,2)
Fc_model1 = Fc_model[is.finite(Fc_model)]
fc_cost1 = fc_cost[is.finite(fc_cost)]


png(paste0(getwd(),"/" ,PlotsParameterDistributionFolder,"/","FecundityCost_Model.png"))
par(mar = c(5, 5, 4, 2) + 0.3)
hist(Fc_model1, cex.axis = 2, xlab = list(expression(Resistance ~ effect ~ to ~ fecundity ~f[c]), font = 3, cex = 2), ylab = list("Count", font = 3, cex = 2), main = "")
dev.off()

#--------------- Resistance cost -------
newdf$cost<-1-((100-as.numeric(newdf$C_Mortality_Mean.x))/(100-as.numeric(newdf$C_Mortality_Mean.y)))
#newdf<-newdf[-c(79), ]  ##removed -Inf row
mean(newdf$cost, na.rm = T)
cost_sd<-sd(newdf$cost, na.rm = T)
res_cost<-newdf$cost #vector with all resistance costs
Rc_model = 1/(1-res_cost) - 1#((as.numeric(newdf$C_Mortality_Mean.x))/100)/(-1+as.numeric(res_cost))
Rc_model1 = round(Rc_model[is.finite(Rc_model)],2)


png(paste0(getwd(),"/" ,PlotsParameterDistributionFolder,"/","ResistanceCost_Model.png"))
par(mar = c(5, 5, 4, 2) + 0.3)
hist(Rc_model1, cex.axis = 2, xlab = list(expression(Resistance ~ effect ~ to ~ viability ~ r[c]), font = 3, cex = 2), ylab = list("Count", font = 3, cex = 2), main = "")
dev.off()

write.csv(1-New_Choice[is.finite(New_Choice)], file = paste0(getwd(), "/", ParameterDataFolder,"/","ChoiceIndex_SpRank_11.csv"), sep = "", quote = TRUE )
# write.csv(Rc_model1[is.finite(Rc_model1)], file = paste0(getwd(), "/", ParameterDataFolder,"/","ViabilityCost_SpRank_11.csv"), sep = "", quote = TRUE )
# write.csv(Fc_model[is.finite(Fc_model)], file = paste0(getwd(), "/", ParameterDataFolder,"/","FecundityCost_SpRank_11.csv"), sep = "", quote = FALSE)
write.csv(Rc_model1, file = paste0(getwd(), "/", ParameterDataFolder,"/","ViabilityCost_SpRank_11.csv"), sep = "", quote = TRUE )
write.csv(Fc_model1, file = paste0(getwd(), "/", ParameterDataFolder,"/","FecundityCost_SpRank_11.csv"), sep = "", quote = FALSE)
write.csv(Ttau2[is.finite(Ttau2)], file = paste0(getwd(), "/", ParameterDataFolder,"/","Toxicity2_SpRank_11.csv"), sep = "", quote = FALSE)
write.csv(round((lambda/100)[is.finite(lambda)],2), file = paste0(getwd(), "/", ParameterDataFolder,"/","Lambda_SpRank_11.csv"), sep = "", quote = FALSE)
write.csv(New_AM[is.finite(New_AM)], file = paste0(getwd(), "/", ParameterDataFolder,"/","AssortativeMating_SpRank_11.csv"), sep = "", quote = FALSE)
write.csv(freq_res_loci[is.finite(freq_res_loci)], file = paste0(getwd(), "/", ParameterDataFolder,"/","Fr0_SpRank_11.csv"), sep = "", quote = FALSE)
}


setwd(workingdirectory)
```
