## Supplementary material for "The role of evolving niche choice in herbivore adaptation to host plants": These are R scripts and data files used for generating results.: READMEFIRST.pdf

We provide the R scripts for simulating and analysing the individual-based model "The role of evolving niche choice in herbivore adaptation to host plants". We also provide R scripts for analysing the metadata, showing how parameter distributions were for model simulations.

To run any of the files, **FIRST CREATE A FOLDER NAMED** "PLANTHERBIVORECHOICE" in your working directory folder and **place the data file, ``S7.xlsx", and all the R files into this folder**. The R files include:

"IBM_PlantHerbivore_ParameterSamplingSpeciesAnalysis_SV.R",
"IBM_PlantHerbivore_ParameterSamplingSpeciesExecute_SV.R",
"IBM_PlantHerbivore_ParameterSamplingSpeciesPlotting_SV.R"
"IBM_PlantHerbivore_SimulationAnalysis_SV.R",
"IBM_PlantHerbivore_SimulationExecution_SV.R",
"IBM_PlantHerbivore_SimulationPlotting_SV.R",
"IBM_PlantHerbivore_SV.R",
"MetaAnalysisData_EachSpecies_SV.R", and
"Density_Dependent_Survival_Fitting_SV.R".

The script "MetaAnalysisData_EachSpecies_SV.R" extracts data from the Excel sheets, analyses the extracted data, and produces plots used in the manuscript. It also produces and saves (as CSV files) parameter distributions used during the sampling simulations. Therefore, **THIS SCRIPT MUST BE BEFORE THE** "IBM_PlantHerbivore_ParameterSamplingSpeciesExecute_SV.R" SCRIPT. The density-dependent survival function is obtained from the "Density_Dependent_Survival_Fitting_SV.R" script. This script contains all the data and the code for fitting. Note that there could be warning/error messages when producing figures from this script. It is OK as some plots in the middle of a for-loop have no data points and we avoided manually excluding such. So, no worries about the warning messages as the code will still run to complete the loop.

The main function for the individual-based model is given in the R script "IBM_PlantHerbivore_SV.R". This script is executed using "IBM_PlantHerbivore_SimulationExecution_SV.R" and "IBM_PlantHerbivore_ParameterSamplingSpeciesExecute_SV.R" scripts for parameter exploration and parameter distribution sampling simulations respectively.

The data produced by "IBM_PlantHerbivore_SimulationExecution_SV.R" is analysed using the "IBM_PlantHerbivore_SimulationAnalysis_SV.R" script. After the analysis, the plots are obtained by running the "IBM_PlantHerbivore_SimulationPlotting_SV.R" script.

Similarly, for the empirical data sampling, we use the scripts "IBM_PlantHerbivore_ParameterSamplingSpeciesExecute_SV.R", "IBM_PlantHerbivore_ParameterSamplingSpeciesAnalysis_SV.R", and "IBM_PlantHerbivore_ParameterSamplingSpeciesPlotting_SV.R" for data simulation, data analysis, and plotting respectively. The scripts should be run in this order: execution, analysis, and plotting.

Note, however, that to get the figures in the manuscript, one has to pick plots produced using specified parameters and combine the single plots to produce the whole figure supplied in the manuscript.

It is easy to isolate required plots. For example, for scenarios where we explore the effect of one parameter while keeping others constant, one has to note down the index for default parameters, and the index for the focal parameter of interest is 0 (the one on the x-axis). Form the search string and search in the specific plot folder. For means, the name string contains "mean" or "var" for variance. All in all, one has to pay attention to the name strings used when running the plotting scripts. Note also that, for a given parameter combination, default parameters are already fixed within the loops.

Within the scripts, all the parameters are described in the comments. The denotation/symbols of parameters in the scripts differ from those used in the manuscript partly because they are generic functions within R. Therefore, we strongly advise reading through the parameter description provided within the script to determine the right parameter combinations. Also, note that other parameters not used in our manuscript are clearly identified via comments.

Note that the number of simulations/parameter sets indicated in the scripts is much lower than that used to produce results. **We used 1000 replicates under parameter exploration simulations and 500 parameter sets with 100 replicates per parameter set for the parameter distribution sampling scenarios. However, we have indicated only 5 replicates/parameter sets in all cases.** We also give the actual values in the comment within the R script. Also, in the plot scripts, the axes limits and legends may have to be manually changed to suit the specific parameter combination(s). Other parameters are indicated as used in the production of the actual results.

WARNING! The actual parameters used to produce the figures in the manuscript may need a lot of time, ranging from hours to weeks even when using a node on cluster with 20 cores! All the execution scripts are parallelized to run on multiple cores. So, it is more suitable to use a computer or cluster with more cores. For replicates, we either used 1000 or 500, as indicated above. To avoid long waiting times on cluster for a node of 20 cores, these replicates were divided during the actual simulations and later combined. In addition, different parameter combinations were also run separately. In the execution codes provided, we assume all replicates will be run in one script. You are advised to run one parameter combination at a time and not all in one script. This recommendation applies specifically to the "IBM_PlantHerbivore_SimulationExecution_SV.R" script. Similarly, simulations for parameter distribution sampling can be split such that one does not run all the 500 parameter sets/100 replicates or 11 species in the for loop but runs say 50 parameter sets/10 replicates or 2 species at a time and later combines the data. This comment applies to the "IBM_PlantHerbivore_ParameterSamplingSpeciesExecute_SV.R" script. In addition, one can reduce the length of the parameter vectors so that fewer parameter values are run. However, you have to adjust the indices of default parameters supplied within the for loops accordingly.

These scripts were run on a Linux computing cluster (Ubuntu 22.04.3). The produced data was analysed in the Linux operating system (Release: 22.04 ), with R version 4.3.2 (2023-10-31), and also tested on the Ubuntu 20.04 release.

END
